# PI3Kδ inhibition potentiates glucocorticoids in B-lymphoblastic leukemia by decreasing receptor phosphorylation and enhancing gene regulation

**DOI:** 10.1101/2023.02.10.527869

**Authors:** Jessica A.O. Zimmerman, Mimi Fang, Miles A. Pufall

**Affiliations:** Division of Pediatric Hematology/Oncology, Stead Family Department of Pediatrics, Carver College of Medicine, University of Iowa, Iowa City, IA, USA; Holden Comprehensive Cancer Center, University of Iowa, Iowa City, IA, USA; Department of Biochemistry and Molecular Biology, Carver College of Medicine, University of Iowa, Iowa City, IA, USA

**Author notes:** **Corresponding author:** Miles A. Pufall, PhD, University of Iowa Carver College of Medicine, 51 Newton Rd, 4-430 BSB, Iowa City, IA 52242.

## Abstract

Glucocorticoids, including dexamethasone and prednisone, are the cornerstone of B-lymphoblastic leukemia (B-ALL) therapy. Because response to glucocorticoids alone predicts overall outcomes for B-ALL, enhancing glucocorticoid potency is a route to improving outcomes. However, systematic toxicities prevent the use of higher dose and more potent glucocorticoids. We therefore took a functional genomic approach to identify targets to enhance glucocorticoid activity specifically in B-ALL cells. Here we show that inhibition of the lymphoid-restricted PI3K*δ*, signaling through the RAS/MAPK pathway, enhances both prednisone and dexamethasone activity in almost all *ex vivo* B-ALL specimens tested. This potentiation is most synergistic at sub-saturating doses of glucocorticoids, approaching the EC50. Potentiation correlates with global enhancement of glucocorticoid-induced gene regulation, including regulation of effector genes that drive B-ALL cell death. Idelalisib reduces phosphorylation of the glucocorticoid receptor (GR) at MAPK1/ERK2 targets S203 and S226, and ablation of these phospho-acceptor sites enhances glucocorticoid potency. We further show that phosphorylation of S226 reduces the affinity of GR for DNA *in vitro*, which impairs DNA binding. We therefore propose that PI3K*δ* inhibition improves glucocorticoid efficacy in B-ALL in part by decreasing GR phosphorylation, increasing DNA binding affinity, and enhancing downstream gene regulation. The overall enhancement of GR function suggests that idelalisib will provide benefit to most patients with B-ALL by improving outcomes for patients whose disease is less responsive to glucocorticoid-based therapy, including high-risk disease, and allowing less toxic glucocorticoid-sparing strategies for patients with standard-risk disease.

## INTRODUCTION

Glucocorticoids, including dexamethasone and prednisone, are the cornerstone of chemotherapy regimens for B-lymphoblastic leukemia (B-ALL), the most common childhood cancer (1). Although about 90% of patients with standard-risk B-ALL are cured with glucocorticoid-based therapies, only about 75% of patients with high-risk B-ALL are as fortunate. Because response to glucocorticoid therapy alone predicts overall outcomes for patients with B-ALL (2,3), enhancing GR activity is an attractive mechanism for improving outcomes, particularly in high-risk disease. The use of more potent or higher dose glucocorticoids is prevented by systemic toxicities, including osteonecrosis, myopathy, and pancreatitis, which significantly impact quality of life and may be life-threatening (4-8). Because glucocorticoids work cell autonomously to kill B-ALL cells, one solution is to enhance GR activity specifically in B lymphocytes. We therefore sought to enhance both prednisone, which is frequently used in high-risk B-ALL induction regimens, and dexamethasone by targeting B-ALL-restricted proteins.

We, and others (9), identified PI3K*δ* as one such promising target. PI3K*δ* is a leukocyte-restricted kinase and a key component of the B-cell receptor (BCR) and interleukin-7 receptor (IL-7R) pathways in B cells. Using functional genomics, we found that knockdown or inhibition of PI3K*δ* sensitizes B-ALL cell to glucocorticoids, likely by blocking signaling through the RAS/MAPK pathway (10). Although PI3K*δ* can activate the AKT pathway, which restrains glucocorticoid potency in some T-ALL (11), we found that AKT had no effect on glucocorticoid sensitivity in B-ALL (10). These findings suggested that inhibition of PI3K*δ* would be an effective way to enhance glucocorticoid potency specifically in B-ALL without increasing systemic potency and toxicity.

Idelalisib is a first-in-class, isoform specific PI3K*δ* inhibitor used in the treatment of relapsed chronic lymphocytic leukemia. Though effective, long-term use of PI3K*δ* inhibitors has been associated with significant immune-mediated adverse effects, primarily through inhibition of regulatory T-cells (12). Intermittent dosing has been shown to improve adverse effects while maintaining anti-tumor activity (12) and could therefore be used during glucocorticoid courses in B-ALL treatment regimens. Using a limited number of patient specimens and cell lines, we found that idelalisib synergistically enhances dexamethasone-induced B-ALL cell death (10), underscoring its potential. It is not known whether idelalisib also enhances prednisone potency, particularly in high-risk B-ALL.

Glucocorticoids work through the glucocorticoid receptor (GR), a ligand-activated transcription factor, to regulate thousands of genes, including those that induce cell death in B-ALL. Glucocorticoids regulate ∼80 effector genes – genes whose regulation contribute to cell death. Effector genes include those involved in apoptosis (suppression of *BCL2* and upregulation of *BCL2L11* (13,14)) and B-cell development (10). Testing a handful of these effectors, we found that idelalisib enhanced regulation of some, but not all, of these effector genes with different effects in different cell lines (10). This suggested the model that idelalisib enhances glucocorticoid potency different ways in different types of B-ALL. However, a more systematic analysis of the effect of idelalisib on glucocorticoid gene regulation in primary specimens is needed to test this model.

The potentiation of glucocorticoids by PI3K*δ* inhibition could be a result of blocking GR phosphorylation. Consistent with our model, phosphorylation by the BCR/RAS/MAPK pathway modulates GR activity in a cell-type and gene specific manner (15-17) and has been associated with glucocorticoid resistance (18). ERK2 (MAPK1), the terminal kinase in the RAS/MAPK pathway, has been shown to phosphorylate GR at multiple sites, including S203, S211 or S226 (15,16), in other systems. Treatment of B-ALL cells with idelalisib is accompanied by a decrease in phosphorylation of GR at S203 (10), but may affect other sites as well. It is not clear which sites of MAPK phosphorylation affect GR activity in B-ALL and how they might do so in a cell or gene specific manner.

Here we show that, as with dexamethasone (10), idelalisib enhances prednisolone-induced cell death in B-ALL cell lines and most primary patient specimens, particularly at concentrations near the EC50 of dexamethasone and prednisolone. Counter to our previous model, we show that idelalisib enhances glucocorticoid-induced regulation of virtually all genes across almost all specimens tested. Consistent with this global effect on gene regulation, we show that phosphorylation of S226 decreases the affinity of GR for DNA. We therefore propose that idelalisib enhances GR function in part by blocking the phosphorylation of S226 and enhancing DNA binding.

## METHODS

### Cell viability assays

B-ALL cell lines (NALM6 and RS4;11) and NALM6 phospho-GR mutants (GR-S203A and GR-S226A) were tested with combinations of prednisolone (Acros Organics, #449470250) and the PI3K*δ* inhibitor idelalisib (Gilead). Viability was measured using PrestoBlue (ThermoFisher, A13262). Synergy was evaluated using the Bliss synergy model in SynergyFinder 2.0 (19).

Primary specimens from children with newly diagnosed or relapsed B-ALL were obtained after receiving informed consent (University of Iowa IRB protocol #201707711). Cells were isolated by Histopaque density gradient separation.

NALM6, SUP-B15, and RCH-ACV cells were tested with dexamethasone (Sigma, D4902-1g) in combination with ERK1/2 inhibitor SCH772984 (SelleckChem, #S7101).

### Gene expression analysis of NALM6 cells and primary specimens

NALM6 cells were treated with vehicle, dexamethasone (5 nM or 50 nM), idelalisib (250 nM), or dexamethasone and idelalisib. Seven primary patient specimens (MAP010, MAP014, MAP015, MAP016, MAP019, MAP020, MAP031) were treated for 24 hours with vehicle, dexamethasone (25 or 50 nM), prednisolone (25 or 50 nM), idelalisib (500 nM), combinations of dexamethasone and idelalisib, or combinations of prednisolone and idelalisib. Sequencing data was processed using R/Bioconductor and DESeq2 (20) with RUVSeq (21). Code for the analysis is available in the supplemental file.

### Protein expression and purification

The human GR AF1-DBD (27-506) polypeptide, containing most of the N-terminal AF1 region and the DNA Binding Domain (DBD), was expressed and purified as described (22). ERK2 was expressed and purified as described (23).

### Phosphorylation of GR-AF1-DBD and purification

GR-AF1-DBD was phosphorylated with ERK2 (30 minutes at 30°C). GR-AF1-DBD phosphorylated species were separated on a 5/5 MonoQ column (Cytiva) and run over a size exclusion column (Cytiva, Superdex 200).

### Mass Spectrometry

GR-AF1-DBD +/-phosphorylation samples were reduced, alkylated, digested with trypsin, and purified with C18 stage tips (Pierce, #87781) (24,25). Peptides were separated by LC and analyzed on a QExactive HF Orbitrap (ThermoFisher) mass spectrometer. Fully phosphorylated GR and singly phosphorylated GR were compared to unmodified GR-AF1-DBD using Scaffold 5.1.2 (Proteome Software Inc.). Phospho-fragment frequencies were compared to the control to determine the level of phosphorylation at each site.

### Electrophoretic Mobility Shift Assays (EMSA)

The dissociation constants for unmodified and phosphorylated GR-AF1-DBD fragments were measured by EMSA as described (22) using a consensus GR site (5’-GTAC**GGAACA**TCG**TGTACT**GTAC – 3’).

### Phospho-GR western blotting

NALM6 cells were treated with vehicle, dexamethasone (5 nM or 1 μM), idelalisib 250 nM, or dexamethasone plus idelalisib for 24 hours. Western blotting was performed as described (10) using GR-S203P or GR-S226P rabbit polyclonal antibody (generously provided by the Garabedian Lab) or GR IA-1 (10). Changes in GR phosphorylation were determined as the ratio of phospho-GR to total GR compared to controls.

### Phospho-GR mutants by CRISPR

Cas9-RNPs were transfected into cells by electroporation (SF Cell Line 4D-Nucleofector™ X Kit S (Lonza, #V4XC-2032)). After 48-72 hours, editing efficiency was checked by T7EI digest (IDT, #1075931). Cells were single-cell sorted (Becton Dickinson Aria II) into 96 well plates. Positive clones were identified by extracting genomic DNA, PCR amplifying the region, Sanger sequencing of control, experimental, and reference PCR products, and analyzing by TIDER (26).

Additional details are available in the supplement.

### Data availability statement

RNA sequencing data are available at Gene Expression Omnibus (NALM6 cells: GSE215385) and dbGaP (patient specimens: phs003085.v1.p1). Other original data are available upon request from the corresponding author.

## RESULTS

### Inhibition of PI3Kδ increases prednisolone sensitivity in B-ALL cell lines and primary patient specimens

To determine whether idelalisib synergizes with prednisolone to induce B-ALL cell death similar to dexamethasone, we titrated both into two B-ALL cell lines (RS4;11, NALM6) and measured viability. RS4;11 cells were more sensitive to prednisolone (EC50 = 25 nM) than NALM6 cells (EC50 = 80 nM). To quantify the impact of the addition of idelalisib, we evaluated both overall and peak (i.e. highlighting the doses with strongest synergy) Bliss scores with SynergyFinder (19). When combined with idelalisib, RS4;11 and NALM6 cells exhibited additivity based on overall Bliss score (RS4;11: 5.826 ± 0.88; NALM6: 3.055 ± 0.56), but both were highly synergistic at some prednisolone concentrations (peak Bliss score 23.30 for RS4;11; 23.79 for NALM6). The idelalisib concentration at peak synergy is similar in both cell lines (∼1.5 μM), but the prednisolone concentration is around half of the EC50 (RS4;11 ∼10 nM; NALM6 ∼40 nM) (**Figure 1A-B**). This suggested that idelalisib is synergistic at prednisolone concentrations near the EC50 and lower than peak clinical concentrations (∼4.5 µM for 60 mg/m^2^ prednisone dose).

**Figure 1.**
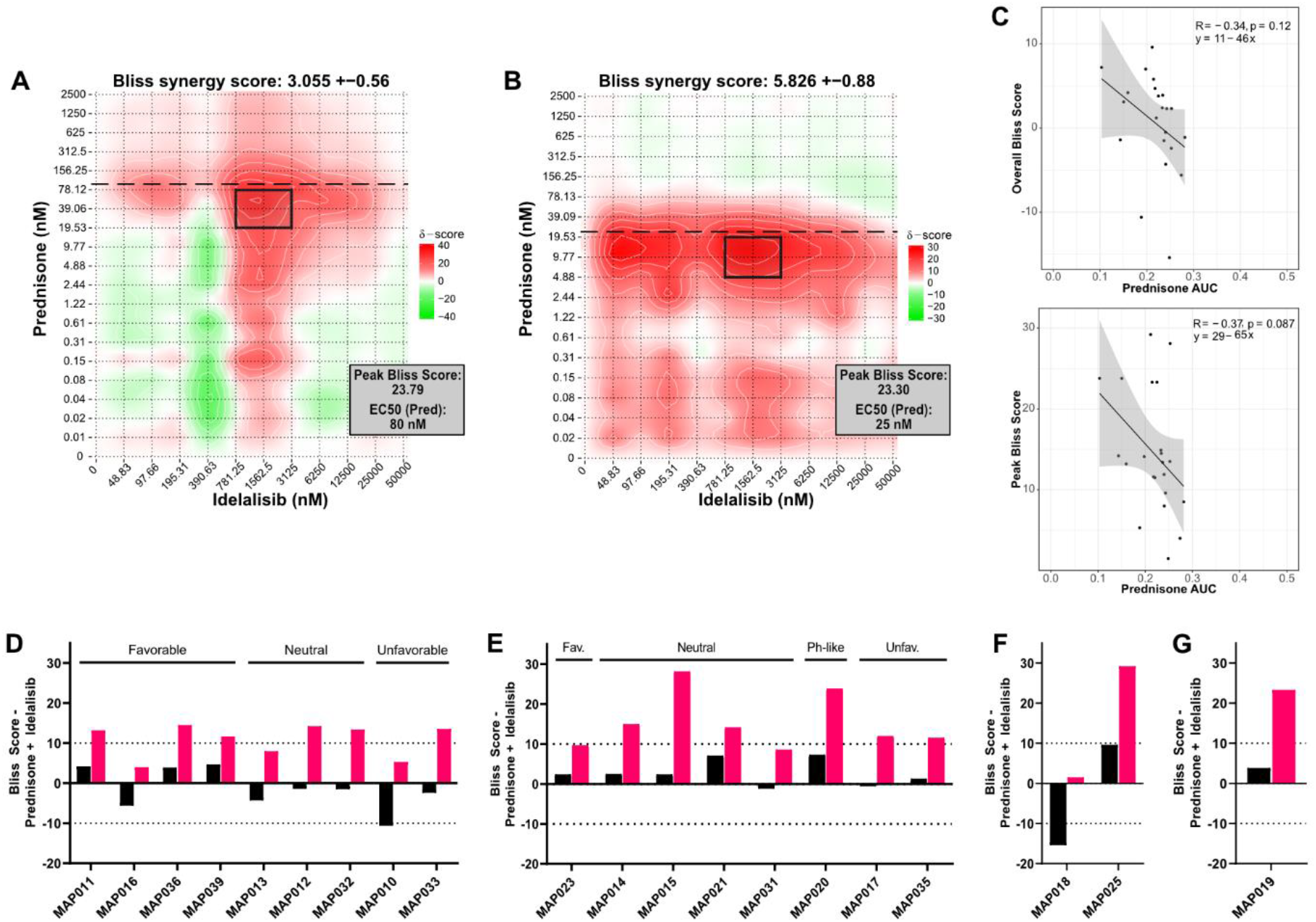
Prednisolone and idelalisib synergistically induce cell death in B-ALL cell lines and primary patient specimens in vitro. Evaluation of synergy in (**A**) NALM6 cells and (**B**) RS4;11 cells treated with the combination of prednisolone and idelalisib. A Bliss score greater than 10 indicates synergy, 10 to −10 indicates additivity, and less than −10 indicates antagonism. Overall score is at the top of each plot, and the peak Bliss score from the black outlined area is given in the box at the top right corner of each plot. Horizontal dashed line indicates the EC50 for prednisolone. (**C**) Correlations (Pearson coefficient R) of prednisolone AUC for patient specimens, NALM6, and RS4;11 cells with overall Bliss score (left) and peak Bliss score (right) demonstrate a trend toward decreased prednisolone sensitivity correlating with increased synergy with idelalisib, although neither has a *p* ≤ 0.05. (**D-G**) Overall (black bars) and peak (pink bars) Bliss scores for the combination treatment of prednisolone plus idelalisib in; (**D**) NCI standard risk specimens, (**E**) NCI high risk specimens, (**F**) infant specimens, and (**G**) a relapsed specimen do not reveal an association between synergy and risk grouping or cytogenetic features. Cytogenetic features of these specimens are favorable/fav (*ETV6::RUNX1* or double trisomy), unfavorable/unfav (*KMT2A* rearrangement, iAMP21, or hypodiploidy), Ph-like (*P2RY8::CRLF2*), or neutral (all other cytogenetic features).

We then evaluated prednisolone/idelalisib synergy by testing 20 primary B-ALL specimens (**Supplemental Table 1**). Most primary specimens were sensitive to prednisolone (EC50 = 6.5-71 nM). Three specimens were glucocorticoid resistant, including one with no response to glucocorticoids (MAP010: 105 nM; MAP021: 250 nM; MAP020 no response). We then compared the EC50 of each specimen to the viability of the specimen at the maximum concentration of prednisolone (10 μM) (**Supplemental Figure 1**). Fourteen specimens had <30% viability, with the majority (12/14) from patients with negative end of induction minimal residual disease (MRD). Of the six specimens with >50% viability, four were MRD positive, one was MRD negative, and one was unknown. Thus, consistent with the literature (2,27), we find that prednisolone response correlates with patient response, particularly with respect to reduction in viability (**Supplemental Figure 1**).

To account for both prednisolone EC50 and viability of each specimen, we calculated area under the curve (AUC) and compared these to overall and peak Bliss scores. Although no correlation reached statistical significance, a trend emerged with higher prednisolone AUC (i.e. higher glucocorticoid sensitivity) associated with lower overall and peak Bliss scores (R = −0.34 and - 0.37, respectively; **Figure 1C**). This suggests that specimens which are more glucocorticoid resistant might demonstrate higher levels of synergy with the addition of idelalisib.

We next attempted to identify a pattern predicting synergy between glucocorticoids and idelalisib, particularly through National Cancer Institute (NCI) risk grouping or cytogenetic features. An additive response of idelalisib with prednisolone was observed in 90% (18/20) of specimens based on overall Bliss score, including both NCI standard risk and NCI high risk specimens with favorable, neutral, and unfavorable cytogenetics. Overall antagonism was evident in the two remaining specimens with notable cytogenetic features – MAP010 (overall Bliss score −10.6 ± 1.1) with near haploid cytogenetics, and MAP018 (overall Bliss score −15.4 ± 2.4) with *BCR::ABL1*. Fourteen specimens exhibited a peak synergistic Bliss score (**Figure 1D-G, Supplemental Figure 2**). As in the cell lines, the peak synergistic area is near the prednisolone EC50 in all specimens except for three specimens with >50% viability (MAP012, MAP020, MAP025) and the relapsed B-ALL specimen (MAP019). Unlike cell line models, peak synergy in primary patient specimens does not occur at a consistent idelalisib concentration. We were also able to treat seven specimens with dexamethasone in combination with idelalisib, producing similar results (**Supplemental Figure 3**). Thus, synergy between idelalisib and glucocorticoids was evident across NCI risk and cytogenetic groups, including high risk and relapsed specimens, though without a consistent pattern.

**Figure 2.**
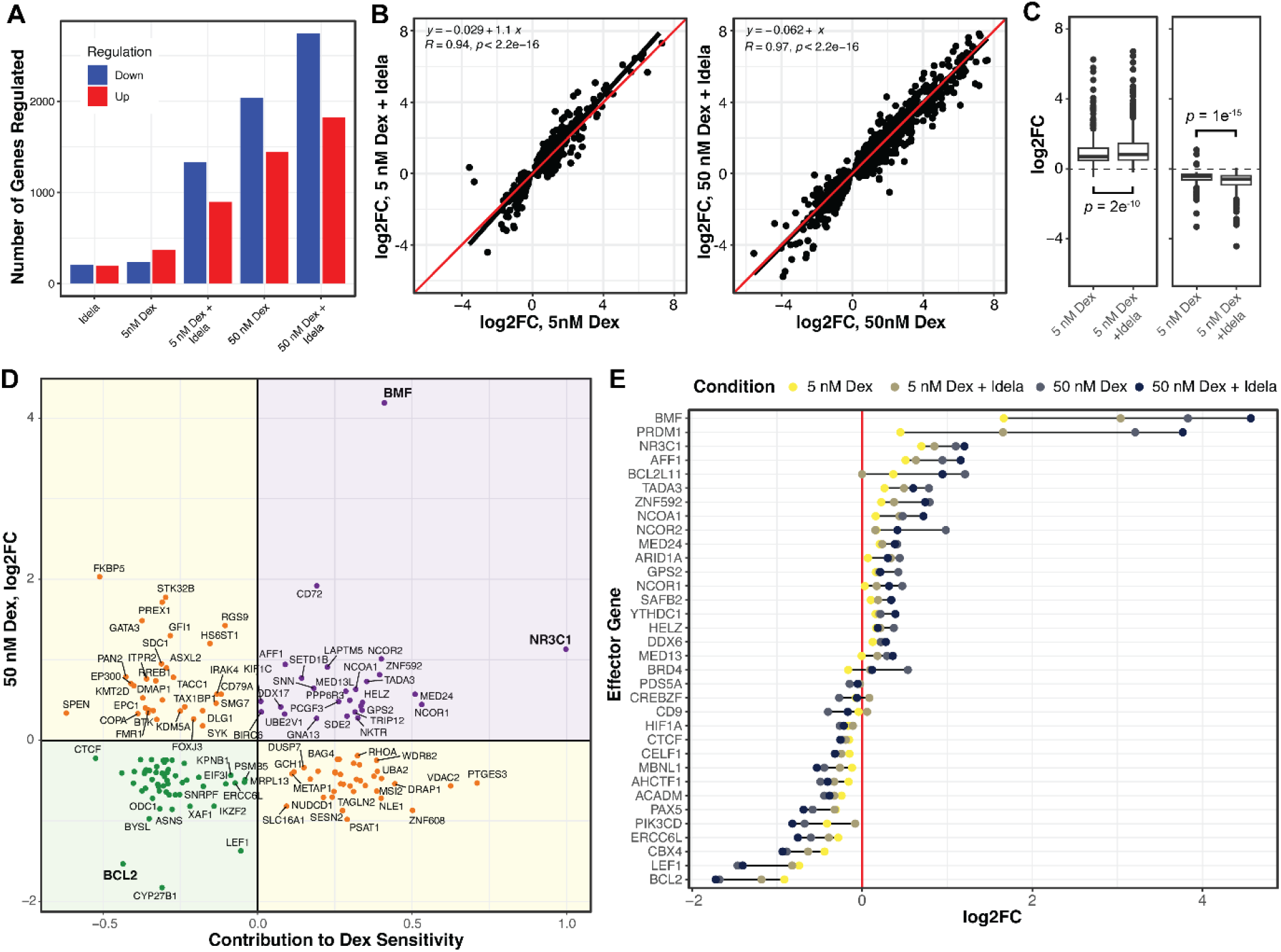
Idelalisib enhances dexamethasone regulation of effector genes in NALM6 cells. (**A**) The numbers of genes up- and down-regulated (*p* ≤ 0.01) for NALM6 cells treated with combinations of dexamethasone (dex) and idelalisib (idela) for 24 hours. (**B**) The log2 fold change in genes regulated by 5 nM dexamethasone is enhanced by idelalisib (left), whereas the enhancement by idelalisib at 50 nM dexamethasone is less pronounced for most genes. The linear regression fit is a black line compared to the red line which would be no effect. (**C**) Box plots of up-regulation (left) and down-regulation (right) by 5 nM dexamethasone show significantly enhanced regulation by the addition of idelalisib. (**D**) Plot of the effect of each gene on dex-sensitivity (x-axis) versus regulation by 50 nM dexamethasone (y-axis). Positive effector (purple) and negative effector (green) genes are those whose regulation contributes to dex-induced NALM6 cell death, whereas buffering genes (yellow) oppose dex-induced cell death. (**E**) The log2 fold change of effector genes in response to combinations of dexamethasone and idelalisib. In 2B and 2C, R = Pearson Correlation coefficient. The concentration of idelalisib used in all the experiments was 250 nM.

**Figure 3.**
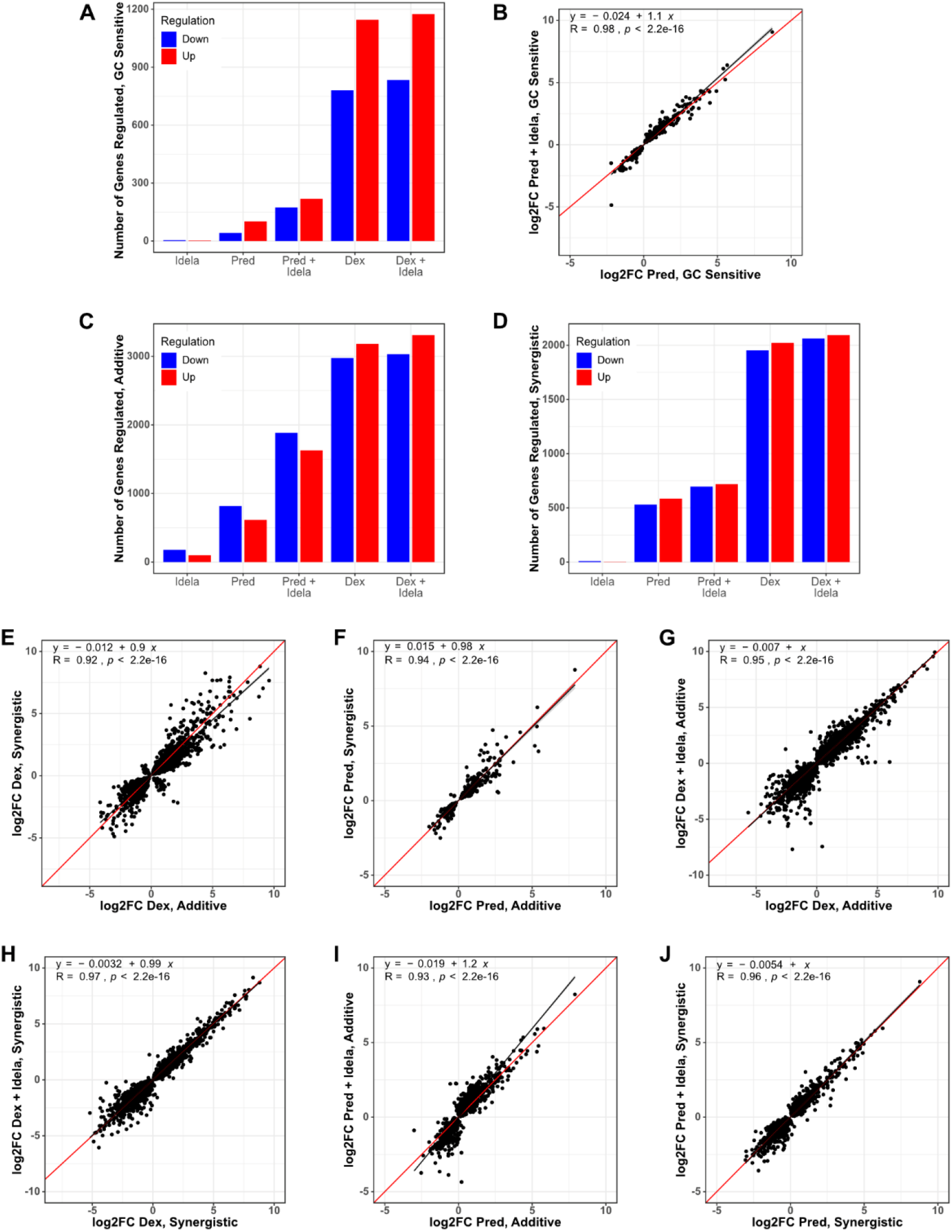
Idelalisib enhances prednisolone-induced gene regulation in glucocorticoid-sensitive primary patient specimens. (**A**) Overall number of genes upregulated (red) and downregulated (blue) in five glucocorticoid-sensitive primary patient specimens treated with idelalisib only (idela), prednisolone only (pred), prednisolone + idelalisib (pred+idela), dexamethasone only (dex), or dexamethasone + idelalisib (dex+idela). (**B**) Comparison of gene expression with prednisolone (x-axis) versus prednisolone + idelalisib (y-axis) in glucocorticoid-sensitive specimens. (**C-D**) Overall number of genes regulated in (**C**) two primary patient specimens with an additive response (MAP014 Bliss score 3 and MAP031 Bliss score −5) and (**D**) two primary patient specimens with a synergistic response (MAP015 Bliss score 22 and MAP019 Bliss score 14) to combination pred+idela treatment in viability assays at the concentrations used for RNA-seq. (**E-F**) Comparison of gene expression with (**E**) dexamethasone only or (**F**) prednisolone only in primary patient specimens with additive (x-axis) vs. synergistic (y-axis) responses to pred+idela treatment in viability assays. (**G-H**) Comparison of gene expression with dexamethasone only (x-axis) versus dexamethasone plus idelalisib (y-axis) in (**G**) primary patient specimens with an additive response or (**H**) with a synergistic response to pred+idela treatment in viability assays. (**I-J**) Comparison of gene expression with prednisolone only (x-axis) versus prednisolone plus idelalisib (y-axis) in (**I**) primary patient specimens with an additive response or (**J**) with a synergistic response to pred+idela treatment in viability assays. For all scatterplots, Pearson correlation and regression equations are reported.

### PI3Kδ inhibition induces global enhancement of glucocorticoid-induced gene regulation in NALM6 cells

To understand the mechanism of idelalisib-enhanced cell death, we measured changes in gene regulation with dexamethasone and idelalisib in NALM6 cells by RNA-seq. We first identified differentially regulated genes (adjp ≤ 0.01) in response to low dexamethasone (5 nM, ∼EC50), high dexamethasone (50 nM), idelalisib (250 nM), and the combination of low and high dexamethasone with idelalisib. Idelalisib induced up- and down-regulation of 418 genes (**Figure 2A**). Low dexamethasone induced regulation of fewer genes (649) than high dexamethasone (3779). The combination caused a greater than additive effect in the number of genes regulated, particularly with low dexamethasone (2398 combination vs. 1067 separately) compared to high dexamethasone (4965 combination vs. 4197 separately). This indicates that the combination either induces regulation of new genes relative to either drug alone or enhances dexamethasone-induced gene regulation.

To distinguish between these models, we performed linear regression on genes regulated by dexamethasone plus idelalisib (**Figure 2B-C**). For genes regulated by low dexamethasone, idelalisib significantly enhances both up-regulation (p = 5e^-9^) and down-regulation (p = 1e^-7^) with an average enhancement of 18% (p < 2e^-12^). There is a more modest, but significant, effect of idelalisib with high dexamethasone (3%, p = 1.6e^-5^). This supports the model that idelalisib better enhances gene regulation at glucocorticoid concentrations closer to the EC50 than at high concentrations.

To determine whether adding idelalisib causes regulation of new genes, we incorporated an interaction term in the differential gene expression model. Of 2398 genes regulated by low dexamethasone plus idelalisib, only 18 exhibited a greater than additive effect. Of 4965 genes regulated by high dexamethasone and idelalisib, 72 showed a greater than additive effect. This indicates that a minority of genes are synergistically or newly regulated by the combination, which we evaluate below.

### Idelalisib potentiates glucocorticoid-induced cell death by enhancing effector gene regulation

We first sought to determine whether idelalisib potentiates regulation of effector genes. We identify effector genes by integrating gene regulation data with the results of two large-scale gene knockdown screens in NALM-6 cells previously performed by our lab to identify genes impacting glucocorticoid-mediated cell death (10,28) (**Figure 2D, Supplemental Table 2**). Positive effectors are genes that contribute to dexamethasone-induced cell death (as measured in the screens) and are upregulated by dexamethasone, thereby enhancing cell death. Negative effectors are genes which impair dexamethasone-induced cell death but are down-regulated by dexamethasone, also enhancing cell death. Our highest confidence effector genes are significant in both versions of the screen (10,28) (p < 0.01).

As with most dexamethasone-regulated genes, idelalisib significantly enhances up- and down-regulation of effector genes with low dexamethasone (19%, p = 0.007) but only sporadically enhances genes with high dexamethasone (**Figure 2E**). Only *LSS* and *MED13L* were newly or synergistically regulated upon addition of idelalisib. This indicates that the primary mechanism of glucocorticoid potentiation by idelalisib is global enhancement of gene regulation, not regulation of new genes relative to either drug alone.

### PI3Kδ inhibition enhances prednisolone-induced gene regulation in primary patient specimens

To test whether idelalisib enhances glucocorticoid potency by globally enhancing gene regulation in a more clinically relevant context, we performed RNA-seq in seven freshly isolated primary B-ALL specimens. Because the glucocorticoid sensitivity of each specimen was unknown at the time of treatment, a relatively high standard dexamethasone concentration (25 or 50 nM) was used. The same prednisolone concentration was used, which is 6-10-fold less potent than dexamethasone. Dexamethasone regulated 1346 genes (adjp ≤ 0.01), and prednisolone regulated fewer genes (99). Idelalisib alone regulated only 7 genes and did not enhance the gene-regulating effects of dexamethasone or prednisolone (**Supplemental Figure 4**). This is likely because two specimens (MAP010, MAP020) are resistant to glucocorticoids (>50% viability at 10 µM prednisolone), exhibiting regulation of very few genes (dexamethasone 41, prednisolone 8, adjp < 0.01).

**Figure 4.**
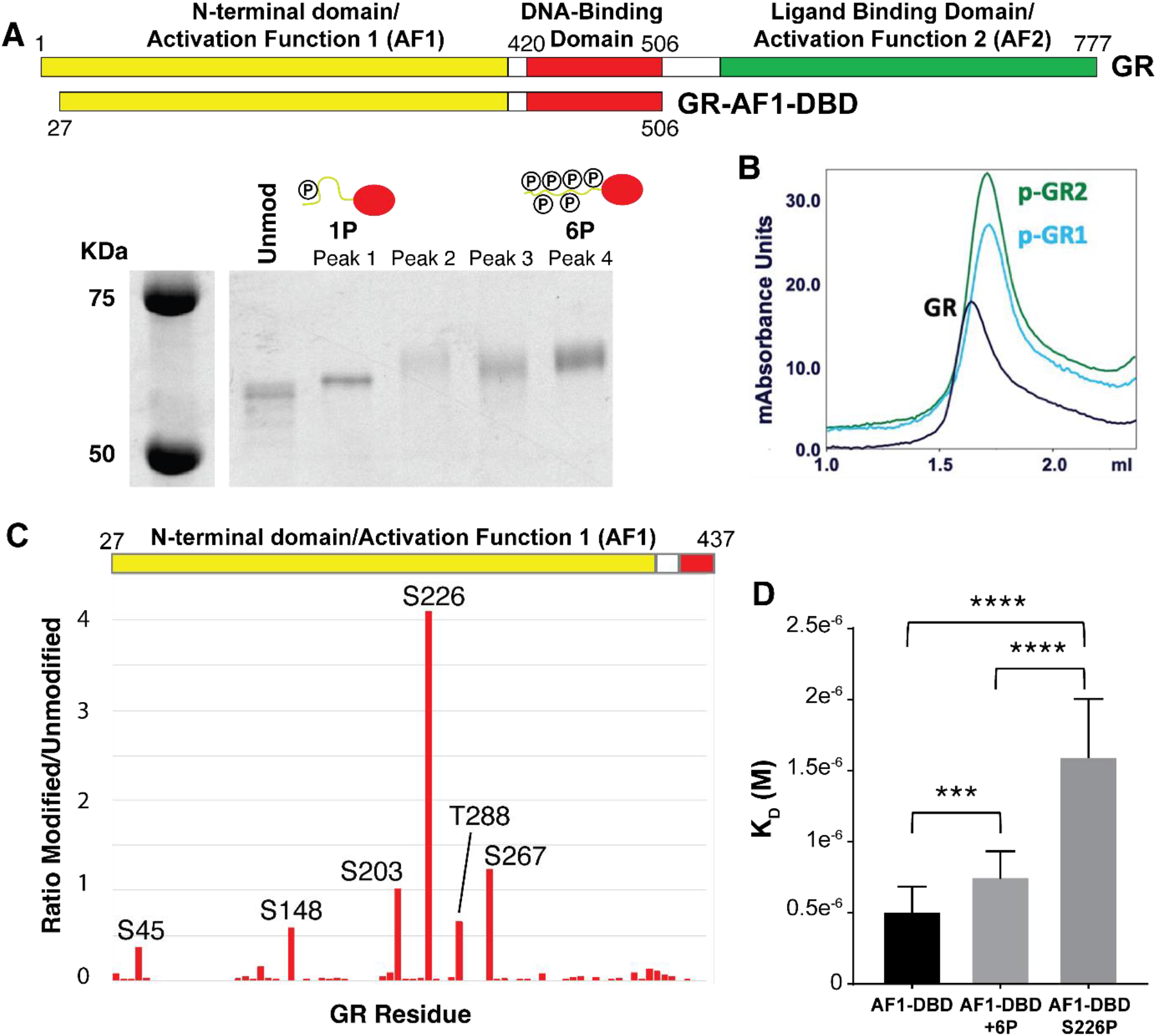
Phosphorylation of GR at S226 inhibits DNA binding. (**A**) Purified GR-AF1-DBD was expressed and phosphorylated with ERK2. Phosphorylated species of GR-AF1-DBD were separated by strong anion exchange, isolating species with primarily one (1P) and six (6P) phosphates. (**B**) Size exclusion chromatography indicated that unmodified GR (black) eluted at a lower volume than two independent samples of GR-AF1-DBD-1P (blue, green). (**C**) Mass spectrometry of phosphorylated GR-AF1-DBD-6P maps phosphorylation at six residues after ERK2 phosphorylation and isolation of GR-6P, including S203, with S226 being the most prevalent. (**D**) Unmodified GR-AF1-DBD binds with higher affinity than 6P, but when GR-1P (S226P) binding is more strongly inhibited. Dissociation constants (KD) were measured by electrophoretic mobility shift assays. Adjusted p-values for one-way ANOVA are 0.0036 (***) and <0.0001 (****).

We therefore tested whether idelalisib enhanced regulation in glucocorticoid-responsive specimens (MAP014, MAP015, MAP016, MAP019, MAP031). In these specimens, dexamethasone induced regulation of 1951 genes and prednisolone 146 genes (**Figure 3A**). Idelalisib alone only regulated 6 genes. These specimens showed enhanced gene regulation upon addition of idelalisib to prednisolone (**Figure 3B**), similar to the combination of idelalisib and low dexamethasone in NALM6 cells. However, idelalisib did not enhance gene regulation with dexamethasone (**Supplemental Figure 4**). Effector genes in these specimens also showed similar patterns of enhanced expression as NALM6 cells (**Supplemental Figure 5**).

**Figure 5.**
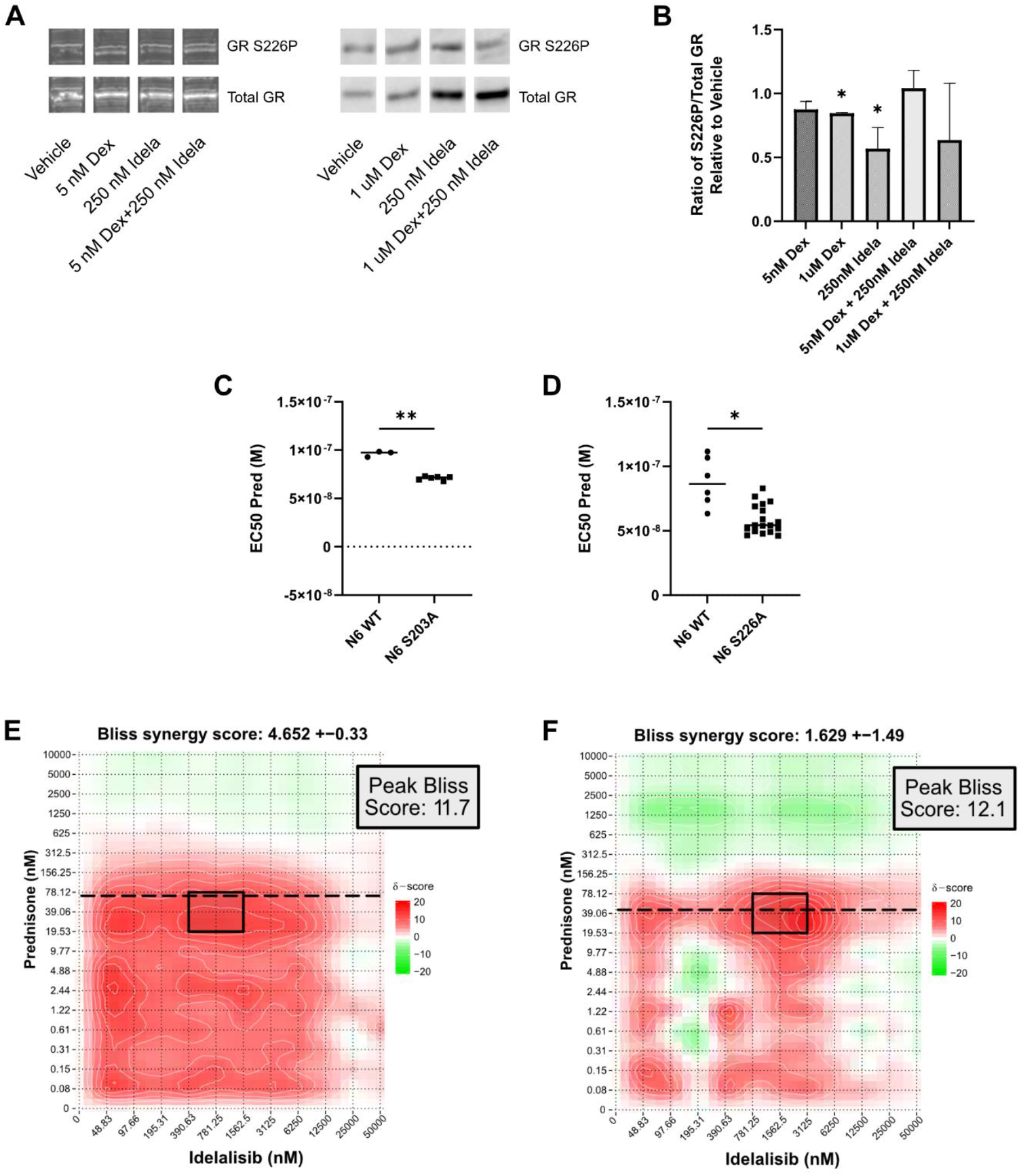
Decreased phosphorylation of GR at S203 or S226 increases glucocorticoid sensitivity. (**A**) Bands from representative western blots illustrating phosphorylated S226 of GR in NALM6 cells treated with vehicle, dexamethasone (Dex) only, idelalisib (Idela) only, and dexamethasone with idelalisib (Dex+Idela) for 24 hours. Both low dose dexamethasone (5 nM, left) and high dose dexamethasone (1 μM, right) were used. Phosphorylated S226 and total GR were blotted on the same membrane for each set of treatment conditions. (**B**) Quantification of the ratio of phosphorylated S226 to total GR normalized to vehicle control. Two biological replicates were performed for each condition except for idelalisib alone, which had 4 biological replicates. Phosphorylation of S226 is significantly reduced with both 1 μM Dex (*p* = 0.02) and 250 nM Idela (*p* = 0.01) compared to a theoretical mean of 1. (**C**) EC50 of prednisolone for CRISPR mutants with GR S203A compared to wild-type (WT) NALM6 cells. Two GR S203A clones were evaluated, with 3 biological replicates for each cell type. Welch’s t test *p* = 0.0010 (**). (**D**) EC50 of prednisolone for CRISPR mutants with GR S226A compared to wild-type (WT) NALM6 cells. Three GR S226A clones were evaluated, with 3 biological replicates per cell type. Welch’s t test *p* = 0.0111 (*). (**E-F**) Representative Bliss synergy plots for (**E**) NALM6 GR S203A cells tested with prednisolone and idelalisib and (**F**) NALM6 GR S226A cells tested with prednisolone and idelalisib. The peak Bliss score area is outlined by the black rectangle on each synergy plot, with that score given in the box to the right. The EC50 of prednisolone for each CRISPR mutant is indicated by the horizontal dashed line, demonstrating that peak synergy occurs around the prednisolone EC50. The S203A mutation appears to shift the peak synergy to a lower dose of idelalisib while the S226A mutation decreases the overall Bliss synergy score in comparison to wild-type NALM6 cells.

We then evaluated whether idelalisib better enhanced gene regulation in specimens that responded synergistically (MAP015, MAP019) compared to those that responded additively (MAP014, MAP031) at the treatment concentrations. Counterintuitively, dexamethasone or prednisolone alone regulated more genes in additive compared to synergistic specimens (dexamethasone 6195 vs. 4015; prednisolone 1447 vs. 1115; adjp ≤ 0.01). Idelalisib alone regulated genes in additive specimens (298) but few in synergistic specimens (10) (**Figure 3C-D**). When comparing glucocorticoid-regulated genes shared by both groups, additive specimens demonstrated stronger gene regulation with dexamethasone, and to a lesser extent with prednisolone, compared to synergistic specimens (**Figure 3E-F**). Additive specimens were also more sensitive to prednisolone alone based on viability compared to synergistic specimens (additive: MAP014 13%, MAP031 8%; synergistic: MAP015 25%, MAP019 27%). Since we had observed a possible correlation between lower glucocorticoid sensitivity and greater synergy, we hypothesized that synergistic specimens may be more amenable to glucocorticoid potentiation than additive specimens.

To evaluate this model, we examined how the addition of idelalisib changed glucocorticoid regulation of genes in the additive and synergistic specimens. Idelalisib did not enhance regulation of genes in either the additive or synergistic specimens compared to dexamethasone alone (**Figure 3G-H**). Counter to the model, idelalisib did enhance prednisolone gene regulation in additive specimens (**Figure 3I**), but not in synergistic specimens (**Figure 3J**), possibly due a stronger effect on gene regulation by idelalisib alone in additive specimens. We identified effector genes in the additive and synergistic specimens, and none of these were specifically regulated in the synergistic specimens to explain synergy (**Supplemental Table 3**). This lack of a clear pattern of enhanced gene regulation in synergistic specimens may be because enhanced gene regulation appears to be concentration dependent. Since we prioritized testing freshly isolated specimens, precluding optimization of drug concentrations to achieve maximum synergy in each specimen, the doses used may mask differences in enhancement in additive and synergistic specimens.

### GR is phosphorylated by ERK2 at six sites, most prominently S226

To understand the mechanism of idelalisib-enhanced gene regulation, we examined how idelalisib affects GR phosphorylation. We previously mapped MAPK1/ERK2, a key kinase in B-ALL transformation (29), downstream of PI3K*δ* in the B-cell receptor pathway (10). Inhibition of the RAS/MAPK pathway, and not the AKT/mTOR pathway, appeared to be the main driver of increased glucocorticoid sensitivity with PI3K*δ* inhibition (10). To test this, we measured synergy between an ERK1/2 inhibitor (SCH772984) and dexamethasone in three B-ALL cell lines (NALM6, SUP-B15, and RCH-ACV). The overall and peak Bliss scores were consistent with combination dexamethasone and idelalisib (**Supplemental Figures 6-8**), supporting the model that ERK2 lies downstream of PI3K*δ*.

**Figure 6.**
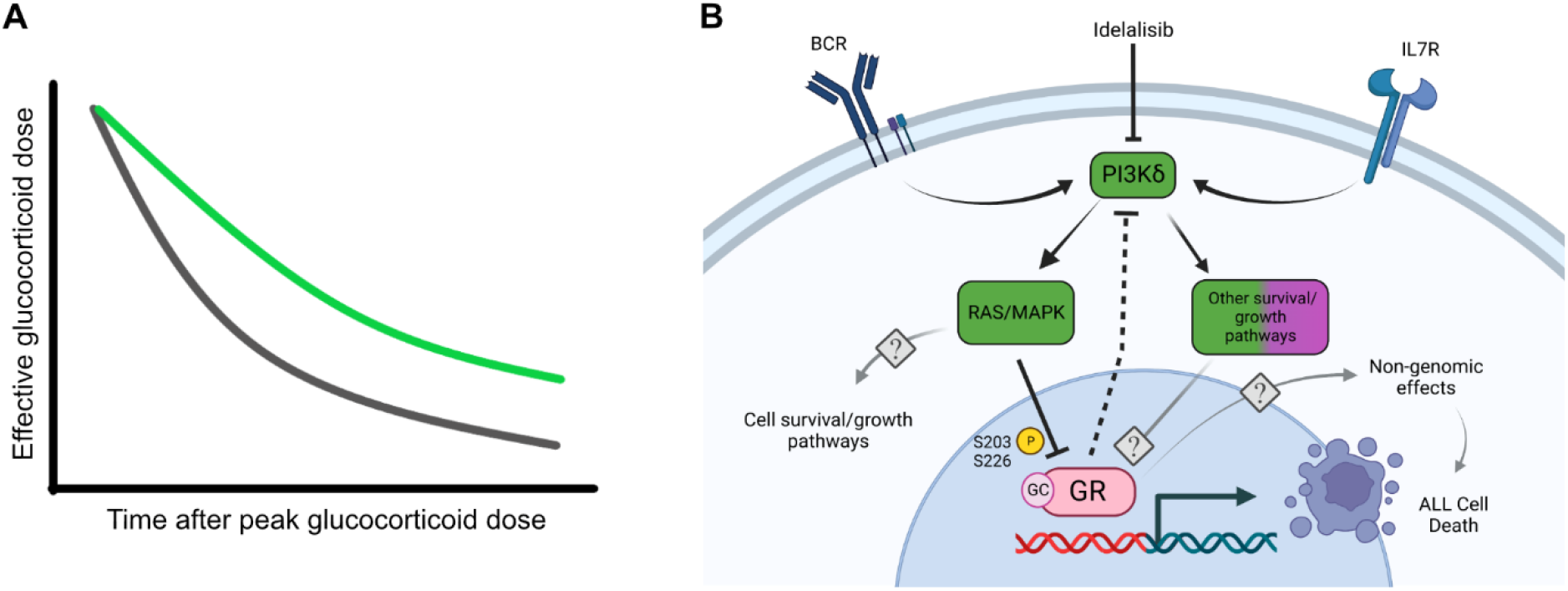
Proposed model of idelalisib-induced glucocorticoid potentiation. (**A**) Schematic of the potential impact on the effective glucocorticoid dose by adding idelalisib to glucocorticoids (green line) compared to glucocorticoids alone (gray line). (**B**) Idelalisib inhibits PI3K*δ*, which subsequently reduces the inhibitory phosphorylation of GR at S203 and S226. BCR = B-cell receptor. IL7R = Interleukin 7 receptor. GR = glucocorticoid receptor. GC = glucocorticoid. Created with BioRender.com.

We then examined how direct phosphorylation of GR by ERK2 (30) affects its activity. To identify relevant sites of ERK2 phosphorylation on GR, we expressed and purified a fragment of GR composed of the N-terminus and DNA binding domain (GR-AF1-DBD), which recapitulates the binding of full-length GR and retains the most commonly phosphorylated sites, and phosphorylated it with ERK2. This resulted in a hazy band shifted upward on an SDS-PAGE gel, indicating that GR-AF1-DBD was phosphorylated but as a mixture of species. We separated the differently phosphorylated species using strong anion exchange (MonoQ) into a high and low mobility species (**Figure 4A**). Interestingly, the highly phosphorylated form of GR-AF1-DBD (GR-6P) eluted in the same volume as the unmodified GR-AF1-DBD over a size exclusion column, but the mono-phosphorylated form (GR-1P) eluted at a later volume. This indicates that GR-1P adopts a more compact conformation than the unmodified or GR-6P species (**Figure 4B**).

Phosphopeptide mapping by mass spectrometry of GR-6P indicated that it is likely a mixture of species with six predominant sites of phosphorylation (**Figure 4C**). Two sites (S203 and S226) were previously reported as sites of ERK2 phosphorylation, but the others are rarely observed (S45, S267, and T288) or previously unreported (S148). The low mobility species (GR-1P) was predominantly phosphorylated at a single site - S226.

We tested the effect of phosphorylation on GR activity by measuring the DNA-binding affinity by electrophoretic mobility shift assay (EMSA) for a consensus GR site (5’-GTAC**GGAACA**TCG**TGTACT**GTAC – 3’). The affinity of GR-6P was modestly (50%) but significantly (p = 0.004) inhibited compared to the unmodified GR. Surprisingly, GR-1P was both substantially (∼3x) and significantly (p < 0.0001) inhibited compared to unmodified and GR-6P (**Figure 4D**). This indicates that phosphorylation of GR by ERK2 not only inhibits GR binding to DNA, but the pattern of phosphorylation can have an important effect, with S226 phosphorylation as a key modification for directly regulating GR affinity.

### Blocking phosphorylation of GR S203 or S226 increases glucocorticoid sensitivity and contributes to idelalisib-induced glucocorticoid potentiation

To validate the importance of GR phosphorylation, we tested the effect of idelalisib on S226 phosphorylation in NALM6 cells (**Figure 5A**). Similar to previous studies showing that idelalisib reduced S203 phosphorylation induced by high (1 µM) dexamethasone (10), idelalisib significantly reduced S226 phosphorylation. However, reduced S226 phosphorylation was only observed with idelalisib alone (p = 0.01) and not in combination with dexamethasone (**Figure 5B**). This indicates that idelalisib reduces phosphorylation of GR at both S203 and S226 but in different ways.

To test the importance of both S226 and S203, which have been shown in other systems to inhibit GR function when phosphorylated (31,32), we generated phospho-acceptor mutants (GR-S203A and GR-S226A) in NALM6 cell lines using CRISPR. Blocking phosphorylation at S203 (**Figure 5C**) and S226 (**Figure 5D**) increased the sensitivity of NALM6 cells to prednisolone (EC50s: wild-type ∼80 nM, GR-S203A ∼70 nM, GR-S226A ∼60 nM), indicating that phosphorylation at either site attenuates GR activity.

We then evaluated both phospho-acceptor mutants with combination prednisolone and idelalisib treatment to determine the impact of blocked phosphorylation on synergy. Interestingly, the GR-S203A clones had similar overall Bliss scores to wild-type NALM6 cells, but peak Bliss scores shifted towards lower idelalisib concentrations (**Figure 5E**). The GR-S226A clones showed decreased overall Bliss scores compared to wild-type NALM6 cells, but peak Bliss scores remained at similar idelalisib concentrations (**Figure 5F**). This suggests that S203 and S226 both contribute to glucocorticoid potentiation with differing and incomplete effects. Since some degree of synergy persists in both the GR-S203A and GR-S226A cells, this also indicates that other PI3K*δ* targets likely impact idelalisib-induced glucocorticoid potentiation.

## DISCUSSION

Our previous work (10) identified PI3K*δ* inhibition as a promising strategy for glucocorticoid potentiation in B-ALL for two main reasons: 1) small molecule inhibitors against PI3K*δ* are available for clinical use, and 2) *PIK3CD* expression is restricted to leukocytes, targeting glucocorticoid potentiation to these cells without increasing off-tumor toxicities. In this work, we show that the PI3K*δ* inhibitor idelalisib potentiates both dexamethasone and prednisone.

Prednisone is 6-10 times less potent than dexamethasone and is less effective in younger children with high-risk B-ALL, even adjusting for dose (33). Patients over 10 years of age with high-risk B-ALL, who are more prone to relapse, receive prednisone during induction to reduce systemic toxicities (33). Thus, the addition of idelalisib during induction has the potential to be well tolerated and improve outcomes for patients with high-risk disease.

Our data also indicate that idelalisib may be most synergistic in B-ALL that is less sensitive to glucocorticoids and predicts poorer outcomes for patients (28,34,35). For high-risk B-ALL, glucocorticoids are administered in doses (10 mg/m2 for dexamethasone or 60 mg/m2 for prednisone) that result in a systemic concentration of ∼700 nM for dexamethasone or ∼ 4 µM for prednisone. We found that idelalisib is most synergistic as the glucocorticoid concentration approaches the EC50 and becomes more additive further from the EC50. As glucocorticoids are metabolized, their concentrations may approach the EC50 for B-ALL cells which are less sensitive to glucocorticoids. Idelalisib would have the most synergy with glucocorticoids in these B-ALL cells, increasing the chances of triggering cell death. It also suggests that the effective dose of glucocorticoids might be maintained over a longer period of time as the glucocorticoid is cleared (**Figure 6A**). Importantly, because PI3K*δ* expression is leukocyte-restricted, this effect is unlikely to cause a significant increase in off-tumor toxicity.

Our previous studies suggested that idelalisib might work differently in different B-ALL backgrounds, perhaps limiting its use to only certain subtypes (10). This was based on measuring synergy in a few cell lines and primary specimens and monitoring cell death effector genes using qPCR. When we measured the effect of idelalisib on the regulation of all genes in more specimens by RNA-seq, we found that idelalisib enhances the regulation of virtually all glucocorticoid-regulated genes rather than select glucocorticoid effector genes (**Figure 2A,C,E**). This enhancement of gene regulation is accompanied by reduced phosphorylation of GR, including at S226, which increases the affinity of GR for DNA. These data indicate that idelalisib is a general potentiator of glucocorticoid activity in B-ALL, and the mechanism of idelalisib is to increase the potency of GR in regulating all genes, including cell death effectors, in part by increasing DNA affinity (**Figure 6B**). This mechanism of generalized GR potentiation is consistent with the enhanced glucocorticoid potency observed in nearly all specimens with measurable glucocorticoid activity. We therefore propose that idelalisib can improve outcomes by potentiating glucocorticoids in high-risk B-ALL and may allow the reduction of glucocorticoid doses and their accompanying off-tumor toxicities without compromising outcomes.

We envision PI3K*δ* inhibitors being combined with glucocorticoids for the treatment of B-ALL in two ways. First, a PI3K*δ* inhibitor like idelalisib could be added to induction glucocorticoid therapy. During the first week of induction, idelalisib could be tested in combination with glucocorticoids using patient specimens *ex vivo*, an approach studied in multiple hematologic malignancies (36-40). If idelalisib potentiates glucocorticoids *ex vivo*, idelalisib could be added for the remaining week(s) of induction glucocorticoid therapy. Second, idelalisib could be added to a lower dose of dexamethasone during the delayed intensification phase. This would enable the maintenance of on-target glucocorticoid potency while decreasing debilitating off-tumor effects of glucocorticoids, particularly the development of osteonecrosis. Both of these strategies could improve glucocorticoid potency during key phases of therapy while avoiding chronic administration of idelalisib, which has proven to be toxic (12).

The direct effects of idelalisib on GR phosphorylation at S203 and S226 do not account for all of its synergistic activity. Although B-ALL cells with GR phospho-acceptor mutations S203A or S226A are more sensitive to glucocorticoids, they retain some synergy with idelalisib. This may be due to some or all of the other 4 sites of ERK2 phosphorylation, including one (S148) that has not been reported to be phosphorylated by any kinase. Testing each phosphorylation site individually and in combination is now feasible using CRISPR and could be performed to assess the importance of the patterns of GR phosphorylation on its activity. Unfortunately, although we were able to purify GR phosphorylated homogeneously at S226, isolation of other phosphorylated species is more challenging. This will likely prevent *in vitro* study of other GR phosphoforms.

PI3K*δ* inhibition may exert its effects on glucocorticoid sensitivity by altering other downstream targets of ERK2 or by signaling through other pathways. For example, the PI3K/AKT pathway phosphorylates GR at S134, inhibiting its activity in T-ALL (11). AKT also phosphorylates and inhibits FOXO transcription factors, which regulate apoptotic and anti-proliferative genes in B-cells (41-43). We do not observe an effect on glucocorticoid sensitivity by knocking down AKT in NALM6 cells (10), but it may play a role in other B-ALL backgrounds. Further study of the downstream effects of idelalisib is needed to fully understand the complete mechanism of synergy with glucocorticoids.

## ACKNOWLEDGEMENTS

The authors acknowledge the work of Maria Nunez Hernandez, who performed the ERK2 phosphorylation experiments in this manuscript but was unable to be contacted to approve submission as a co-author. We also gratefully acknowledge Michael R. Rebagliati, PhD, of the Scientific Editing and Research Communication Core at the University of Iowa Carver College of Medicine for critical reading of the manuscript.

