## Supplemental Methods, Tables, and Figures for "PI3Kδ inhibition potentiates glucocorticoids in B-lymphoblastic leukemia by decreasing receptor phosphorylation and enhancing gene regulation"

##### *Cell viability assays*

Cells were grown in RPMI1640 + 10% FBS at 37°C with 5% $CO_2$, diluted to 625,000 cells per mL, and seeded at 9500 cells (15.2 µL) per well in 384 well plates. Dexamethasone (Sigma, D4902-1g) and prednisolone were diluted in DMSO (2500 µM to 20 nM) and then diluted 1:500 in RPMI1640 + 10% FBS. Idelalisib was also diluted in DMSO (50 mM to 48.8 µM) and then diluted 1:100 in RPMI1640 + 10% FBS. Dexamethasone/prednisolone and idelalisib dilutions were added to cells for a final drug to cell volume ratio of 1:1 for dexamethasone/prednisolone (19 µL) and 1:10 for idelalisib (3.8 µL), with $final_{[DMSO]}$ = 0.2%. Three replicates for each combination of drug concentrations and cell line were tested. Dexamethasone/prednisolone alone and idelalisib alone were also tested in triplicate. After 72-hour incubation at 37°C, the fraction of cells surviving was measured by adding a resazurin-based reagent (PrestoBlue, ThermoFisher, cat# A13261) in a 1:10 ratio of reagent to specimen and scanned for fluorescence (excitation 560 nm, emission 590 nm) on a Biotek NEO. EC50 of prednisolone alone and idelalisib alone was calculated using GraphPad Prism (4-parameter fit) for each replicate. AUC was calculated using the PharmacoGx package in R[1]. Bliss synergy scores were calculated using SynergyFinder 2.0[2], with the Bliss reference model quantifying the multiplicative effect of single drugs as if they acted independently. Bliss synergy scores greater than 10 indicate synergy, scores between -10 to 10 indicate additivity, and scores less than -10 indicate antagonism.

B-ALL cells were isolated from the primary patient specimens using Histopaque-1077 (Sigma-Aldrich) for density gradient centrifugation, and freshly isolated B-ALL cells were tested similarly to cell lines with a few modifications. Primary cells were diluted to ~5 million cells per mL in RPMI1640 + 10% FBS and seeded at ~75,000 cells (15.2 µL) per well in 384 well plates. Since glucocorticoid sensitivity was unknown at the time of treatment, prednisolone dilutions in DMSO were from 10 mM to 80 nM prior to diluting 1:500 in RPMI1640 + 10% FBS. The same idelalisib dilutions were used as with the cell lines. Two replicates for each combination of drug concentrations, prednisolone alone, and idelalisib alone were tested for each primary specimen.

NALM6, SUP-B15, and RCH-ACV cells (all obtained from DSMZ) were tested with dexamethasone and ERK1/2 inhibitor SCH772984 (SelleckChem, cat# S7101) similar to the prednisolone/idelalisib combinations. Dexamethasone was diluted 2500 µM to 10 nM in EtOH for NALM6 and SUP-B15 cells and 10000 µM to 80 nM in EtOH for RCH-ACV cells; these were then diluted 1:500 in RPMI1640 + 10% FBS. SCH772984 was diluted 20mM to 312.5 µM in DMSO and then diluted 1:500 in the previous 1:500 dexamethasone dilutions to create all combinations of dexamethasone/SCH772984 concentrations. These dilutions were added 1:1 to cells for $final_{[DMSO]}$ = 0.1% and $final_{[EtOH]}$ = 0.1%. Cells were also treated for 72 hours prior to assessing their viability.

##### *Gene expression analysis of NALM6 cells with dexamethasone and idelalisib*

NALM6 cells were seeded at 7.5e5 cells/mL in 4 mL RPMI1640 + 10% FBS per well of 6 well plates. Cells were allowed to incubate overnight before treatment with vehicle, dexamethasone (5 nM or 50 nM), idelalisib (250 nM), or combination of dexamethasone (both concentrations) and idelalisib for 3 biological replicates. RNA was extracted after 24-hour treatment using the miRNeasy Mini Kit (Qiagen cat# 217004) per the manufacturer's protocol with on column DNase digestion using Qiagen RNase-free DNase Set (cat# 79254). RNA quality was assessed using Agilent Bioanalyzer. Libraries were prepared using Epicentre ScriptSeq™ Complete Gold Kit (Human/Mouse/Rat)–Low Input (Cat# SCL24G). Sequencing was performed by the Iowa Institute of Human Genetics Genomics Division. Sequencing data was processed using R/Bioconductor and the DESeq2 package. Alpha = 0.01 was used for analysis unless otherwise specified.

***Gene expression analysis of primary specimens with glucocorticoids and idelalisib***

Seven primary patient specimens (MAP010, MAP014, MAP015, MAP016, MAP019, MAP020, MAP031) with at least 30 million cells remaining after evaluation of glucocorticoid sensitivity were treated to evaluate glucocorticoid related gene regulation. Freshly isolated primary B-ALL cells were diluted to 1.1 million cells per mL in RPMI1640 + 10% FBS and seeded at 2 million cells per well in 24 well plates. Cells were then treated with vehicle, dexamethasone (final concentration 25-50 nM), prednisolone (25-50 nM), idelalisib (500 nM), combination dexamethasone and idelalisib, or combination prednisolone and idelalisib. After 24-hour incubation, treated cells were resuspended, centrifuged, resuspended in 700 µL QIAzol lysis reagent (Qiagen, cat# 79306), and stored at -80°C until immediately prior to RNA extraction.

For RNA extraction, specimens were thawed at room temperature and homogenized using QIAshredder columns (Qiagen cat# 79654). RNA extraction was performed as described above. RNA was eluted using 35 µL RNase free water. RNA was cleaned using the Zymo RNA Clean & Concentrator-5 kit (cat# R1013) prior to assessing quality using Agilent Bioanalyzer. Libraries were then prepared by the Iowa Institute of Human Genetics Genomics Division using the Illumina TruSeq Stranded mRNA Library Prep kit. Sequencing was performed on Illumina NovaSeq 6000. One replicate of each treatment condition was evaluated for MAP010, MAP016, and MAP020. Two biological replicates of each treatment condition were evaluated for MAP014 (except for dexamethasone + idelalisib which had 1 replicate fail library preparation), MAP015, MAP019, and MAP031.

Sequencing data was processed using R/Bioconductor and the DESeq2 package. Primary specimens were analyzed first altogether, then glucocorticoid sensitive vs. resistant, and later by comparing those found to have an additive response to combination treatment at the concentrations used for this treatment (MAP014 and MAP031) versus those found to have a synergistic response to combination treatment at the concentrations used for this treatment (MAP015 and MAP019). Alpha = 0.01 was used for analysis unless otherwise specified.

***Protein expression and purification***

The human GR AF1-DBD (27-506), which contains most of the N-terminal AF1 region and the DBD excluding the hinge, was cloned into a his6-tag containing vector (pET28a). BL21 Gold (DE3) E. coli were transformed with the vector. A single colony was picked and grown at 37˚C in 50 ml of standard LB broth (50 µg/mL kanamycin) to an OD600 ~1. One liter LB cultures (50 µg/mL Kan) supplemented with 10 µM $ZnCl_2$ were inoculated with 10 ml of the starter, grown at 37˚C to an OD600 of 0.3-0.4 before shifting the temperature to 23˚C. Once the OD600 reached 0.8-1, protein expression was induced by adding isopropyl-β-D-thiogalactopyranoside (IPTG, 0.5 mM) for 8 hours. Cells were pelleted by centrifugation (4,000g for 15 minutes in a fixed angle rotor), resuspending in Ni2+ loading buffer (25 mM Tris, pH 7.5, 500 mM NaCl, 15 mM Imidazole), snap frozen, and stored at -80˚C.

Pellets of GR AF1-DBD were thawed, adjusted in volume to 20 mL/L culture Ni2+ loading buffer, and lysed by passing three times through an Emulsiflex C3. Lysate from up to 6 L of culture was then loaded onto a 5ml HisTrap column (Cytiva), washed, and eluted with a gradient from 15 mM Imidazole to 500 mM Imidazole over 40 mL. GR AF1-DBD fractions were pooled and incubated with 10U Thrombin/mg protein (T4648) in buffer adjusted to contain 2.5 mM CaCl2 overnight while dialyzing into S-column loading buffer (20 mM Tris, pH 7.5, 50 mM NaCl, 1 mM DTT). In the morning, the protein solution is cleared by spinning (10,000 g for 10 minutes) and then syringe filtering (0.45 µm) before loading onto an HP Sepharose column (Cytiva) and eluting with a gradient from 50 mM to 500 mM NaCl. Fractions are run on a gel and the purest fractions are concentrated and run on a Sepharose 200 column in 20 mM HEPES, pH 7.4, 100 mM NaCl, 1mM DTT, 10% glycerol (Cytiva) to further purify and get rid of soluble aggregates. A concentration is taken by $A_{260}$ (ε = 44810), aliquoted, snap frozen, and stored at -80˚C.

Erk2 was expressed and purified largely as described[3]. pETHis$_6$ ERK2+MEK2 R4F plasmid (Addgene #39212) was transformed into BL21 Gold cells and plated on ampicillin (100 mg/mL) containing LB-agar. A single colony was picked into a 50ml starter culture (100 mg/mL ampicillin) and then grown overnight. 15ml of the starter was inoculated into 1L of LB (100 mg/mL ampicillin), grown to an $OD_{600}$ = 0.8 at 37˚C, then reduced to 30˚C and induced with 0.25 mM IPTG for 14 hours. Cells were then pelleted, resuspended in Ni2+ column loading buffer (20 mL/L culture, 50 mM sodium phosphate, pH 8.0, 0.3 M NaCl, 0.5 mM phenylmethylsulfonyl fluoride (PMSF), 1 µg/ml leupeptin, 1 µM pepstatin, 1 mM benzamidine), frozen in liquid nitrogen, and stored at -80˚C. Resuspended pellets were thawed slowly and lysed by passing three times over an Emulsiflex C3. Lysate was cleared by ultracentrifugation for 60 minutes at 40,000 rpm (Ti45 rotor). Cleared lysate was loaded onto a 5 ml HisTrap FF column (Cytiva), washed, and eluted with a gradient to 500 mM Imidazole. Fractions from the elution peak were pooled and dialyzed into MonoQ loading buffer (20 mM Tris-HCl, pH 7.5, 50 mM NaCl, 1 mM dithiothreitol, 1 mM EGTA, 10 mM benzamidine, 0.2 µM pepstatin, 0.5 mM PMSF, and 20% glycerol). The protein solution was diluted 1:1 with loading buffer containing no glycerol (10% glycerol final) and loaded onto a 5/5 MonoQ column (Cytiva), washed, and eluted with a gradient to 500 mM NaCl. Two major peaks were observed, the second of which (elution NaCl ~250 mM) has been shown to be more active. Fractions from this peak were pooled and then run over a gel filtration column in the same buffer to get rid of high molecular weight contaminants. The concentration was calculated by $A_{260}$ (ε = 44810), and aliquots were snap frozen in liquid nitrogen and storage at -80˚C for future use.

##### *Phosphorylation of GR-AF1-DBD and purification*

GR-AF1-DBD was phosphorylated by Erk2 in 20 mM HEPES, pH 7.4, 10 mM MgCl2, 1 mM DTT, 200 µM ATP for 30 minutes at 30˚C. To favor more highly phosphorylated products, we would phosphorylate 20 µM GR-AF1-DBD with 0.2 µM Erk2 (100:1). To favor less phosphorylated products, the ratio would be 10,000:1 (20 µM GR-AF1-DBD:0.002 µM Erk2). Reactions were then diluted 10:1 in MonoQ loading buffer (20 mM Tris-HCl, pH 8.0, 25 mM NaCl, 10% glycerol, 1 mM DTT), loaded onto a 5/5 MonoQ column, and then eluted with a gradient to 250 mM NaCl. Singly phosphorylated GR-AF1-DBD eluted ~130 mM NaCl (12.2 mS/cm), with highly phosphorylated species eluting ~155 mM NaCl (~14.2 mS/cm). Phosphorylated species were then concentrated (Millipore Amicon Ultra, 10 K MWCO), snap frozen, and stored at -80˚C.

##### *Mass Spectrometry*

*In Solution Trypsin Digestion:* Five micrograms of each protein sample (GR-AF1-DBD unmodified, phosphorylated, and singly phosphorylated and purified) were reduced at 56˚C for 1 hr. in 50 µl buffer (20 mM Tris-HCL, 150 mM KCl, 5 mM $MgCl_2$, 10 mM DTT, and 1 mM EDTA) and alkylated with 56 mM chloroacetamide (CAA) at RT for 30 min (covered). Sequencing grade Trypsin Gold (Promega) in 50mM in $NH_4HCO_3$ was then added to a final concentration of 10 ng/µL and digested overnight at 37˚C. Digested samples were acidified to pH 2-3 with 50% trifluoroacetic acid (TFA) and centrifuged at 20,000 x g for 15min to pellet insoluble material. The supernatant peptides were desalted with C18 Stage Tips (Pierce, #87781)[4, 5] and eluted in 200µl 70% acetonitrile and 0.1% formic acid, were concentrated by lyophilization and reconstituted in 15µL of Mobile Phase A (MPA, 0.1% formic acid with 3% acetonitrile) for LC-MS/MS analysis.

*LC-MS/MS:* 6 µL of peptide digests were auto-loaded by an EZ-nano 1200 UPLC (ThermoFisher) onto a nanocapillary flow path with a 75 µm id x 2 cm trap (ThermoFisher, #164535) coupled to a 75um x 50 cm analytical column (ThermoFisher, #164570) via a S.S. micro-tee (Valco) hosting a split line to waste. Both trap and column are packed with Acclaim PepMap 3 um diameter C-18 coated particles with 100 Å pores. While channeled to waste, the trap was loaded at 2 µL/min and washed with 8 trap volumes of MPA and eluted with a linear gradient to MPB (90% CAN 0.1% formic acid) as follows: 300nL/min for 53min to 35% MPB, 10 min to 60% MPB, 8 min to 98%. Following a 7 min purge, MPA was reset to 97% for column reconditioning. A cyclical routine automated on a Q Exactive HF Orbitrap LC-MS/MS System (ThermoFisher) acquiring one MS1 survey scan (380-1800 Th, 60K resolution, AGC 3E06, IT 100ms) followed by 1.2 Th windowed isolations on up to 16 precursor ions. After HCD activation at 28 NCE, fragment spectra are acquired in centroid mode (30K res, 1E5 AGC, 80ms IT) and each precursor was excluded from the cycle for 30 sec.

*Data Analysis:* Data sets were analyzed using the Proteome Discoverer Search engine with a human protein database downloaded from Uniprot (Nov 9, 2018). Sequences were concatenated with a reversed version as a decoy and searched using a 10 ppm mass error for MS1 and MS2 at 2% False Discovery Rate (FDR). Search settings assumed uniform carbamidomethyl alkylation of Cys residues (+57) and variable modifications including phosphorylation of Ser, Thr, Tyr; oxidation of Met; and rare carbamidomethylation of N-terminal peptide sites. The searches were then combined and rescored using Scaffold Q+S (ProteomeSoftware) implementing Percolator with a 1% FDR.

***Electrophoretic Mobility Shift Assays (EMSA)***

The dissociation constants for unmodified and phosphorylated GR fragments were measured by electrophoretic mobility shift assay (EMSA). A Cy5-labeled oligo (Integrated DNA Technologies) containing a high-affinity GR binding sequence (5'- GTAC GGAACA TCG TGTACT GTAC -3') and its complement were resuspended in water (100 µM final) and mixed 10:1 with 10x annealing buffer (200 nM HEPES, pH 7.4, 1 M NaCl, 50 mM $MgCl_2$). The oligos were then heated (95˚C, 5 minutes) and slow annealed (to 23˚C over 3 hours) for a final duplex concentration of 10 µM. The dsDNA was then diluted to 10 nM in binding buffer (20 mM Tris-HCl, pH 8.0, 150 mM KCl, 10% Glycerol, 5 mM $MgCl_2$, 1 mM DTT, 1 mM EDTA, 200 ng/µL BSA, 40 ng/µL Salmon Sperm) and dispensed in 10 µL aliquots into a row of a 96-well plate. In a separate row, GR protein, diluted from stock with binding buffer to 10 µM, was added to the first well of a new row. GR protein was then serial diluted at a 1:2 ratio along the row, with no protein in the last well. 10 µL of protein were then added to the DNA row (1:1 ratio) and incubated for 1 hour on ice. The EMSA was run on a 4-20% native PAGE (19:1 Acrylamide/bis-Acrylamide) in 1X TG buffer (25 mM Tris-Cl, 250 mM glycine), visualized (GE LAS4010), and quantified (ImageJ). Binding curves were then fit to fraction bound with hill coefficient (Fraction Bound = $(B_{max}*[GR]^h/(K_d^h + [GR]^h))$). Each EMSA was performed ≥ 3 repeats, with phosphorylated species compared to unmodified species by t-test.

***Phospho-GR western blotting***

NALM6 cells were split into four T75 flasks at a density of 0.6 million cells per mL 18-24 hours prior to treatment. Immediately prior to treatment, dexamethasone and idelalisib dilutions were prepared with final concentrations of dexamethasone 5 nM, dexamethasone 1 µM, and idelalisib 250 nM ($final_{[DMSO]}$ = 0.1%). One flask was treated with DMSO (vehicle) only, one with dexamethasone only, one with idelalisib only, and one with dexamethasone plus idelalisib. Low dexamethasone (5 nM) and high dexamethasone (1 µM) blots were performed separately. Samples (4 mL) were removed from each treated flask 24 hours after treatment.

At each treatment timepoint, cells were centrifuged and medium removed from the cell pellet. Cells were lysed in 200 µL lysis buffer supplemented with protease and phosphatase inhibitors (50 mM HEPES pH 7.5, 150 mM NaCl, 1 mM EDTA, 1 mM EGTA, 1 mM NaF, 1% Triton X-100, 10% glycerol, 20 mM β-glycerophosphate, 8 mM sodium pyrophosphate, 1 mM PMSF, 1:500 Calbiochem protease inhibitor cocktail III), vortexed, and incubated on ice for 10 minutes. Lysates were then transferred to 1.5 mL tubes, vortexed again, and centrifuged at 12000 rpm for 15 minutes at 4°C. Supernatants were transferred to new 1.5 mL microcentrifuge tubes. Total protein was quantified based on the Bradford method (Bio-Rad cat# 5000006). SDS sample buffer was added, and each sample was boiled for 3 minutes. Samples were either used immediately or stored at -20°C for future use.

For western blot analysis, 5 µg (for 1 µM dexamethasone blots) or 10 µg (for 5 nM dexamethasone blots) of each sample was loaded in a 15-well Novex Tris-Glycine 4-20% gel (ThermoFisher, cat# XP04205BOX) and electrophoresed at 200V for 45 min in 1X Tris/Glycine/SDS buffer. Gels were transferred to low fluorescence PVDF membrane (Millipore Immobilon, cat# IPFL00010), using fresh, chilled 1X Tris/glycine buffer for wet transfer at 0.25A for 90 minutes. Each membrane was rinsed with 1X TBS-Tween (0.1%).

Blots with 1 µM dexamethasone and blocked with 5% milk in TBS for 1 hour at room temperature. These blots were probed for actin using 1:50000 StrepTactin-HRP (BioRad, #1610381) and imaged on a GE ImageQuant LAS 4000. Blots with 5 nM dexamethasone were stained with Revert total protein (LI-COR, cat# 926-11021) and imaged on an Odyssey Fc. These membranes were then stripped with Revert reversal solution (0.1 M NaOH, 30% methanol) before blocking with Li-COR blocking solution (cat# 927-50000).

Both sets of blots were incubated with primary antibody (1:10000 GR-S203P rabbit polyclonal antibody or 1:1000 GR-S226P rabbit polyclonal antibody, both provided by the Garabedian lab, or 1:500 GR IA-1 rabbit polyclonal antibody, purified at 1 µg/µL) overnight at 4°C. Incubation with secondary antibody (1:10000 donkey-anti-rabbit-HRP, GE Healthcare, for 1 µM dexamethasone blots and 1:10000 anti-rabbit-680, ThermoFisher, cat#35568 for 5 nM dexamethasone blots) was performed for 1 hour at room temperature. Blots were imaged on GE ImageQuant LAS 4000 (1 µM dexamethasone) or Odyssey Fc (5 nM dexamethasone). Quantification of actin/total protein, total GR, and phospho-GR was performed using Image Studio Lite v5.2.

***Phospho-GR mutants by CRISPR***

All oligos for CRISPR were obtained from Integrated DNA Technologies with sequences below:

gRNA:

S203A_1: GAGTTTTCTTCTGGGTCCCC

S203A_2: CTCATTCGTCTCTTTACCTG

S226A_1: AAACTGTTTGCTTTCTCCTC

S226A_2: GAATCGTCTTCTCCCGCCAG

HDR templates:

S203A_1_donor+:
GGTCTGATCTCCAAGGACTCTCATTCGTCTCTTTACCTGGAGCCCCAGAAGAAAACTCCAAATCCTGCAAAATGTCAAAGGTG

S203A_1_donor-:
CACCTTTGACATTTTGCAGGATTTGGAGTTTTCTTCTGGGGCTCCAGGTAAAGAGACGAATGAGAGTCCTTGGAGATCAGACC

S203A_2_donor+:
AACAGGTCTGATCTCCAAGGACTCTCATTCGTCTCTTTACCTGGAGCCCCAGAAGAAAACTCCAAATCCTGCAAAATGTCAAAGGTG

S203A_2_donor-:
CACCTTTGACATTTTGCAGGATTTGGAGTTTTCTTCTGGGGCTCCAGGTAAAGAGACGAATGAGAGTCCTTGGAGATCAGACCTGTT

S226A_1_donor+:
TTTCCTTCCAAAAGGAATGAATCGTCTTCTCCCGCCAGAGGGGCAAGCAAACAGTTTTCATCTATCAACAGGTCTGATCTCCAA

S226A_2_donor+:
TCGAGTTTCCTTCCAAAAGGAATGAATCGTCTTCTCCCGCCAGAGGGGCAAGCAAACAGTTTTCATCTATCAACAGGTCTGATCTCCAA

PCR primers:

S203A_F1a= 5' AGAGAACCCCAAGAGTTCAG 3'

S203A_R1b= 5' GATCCTTGGCACCTATTCCA 3'

S226A_F1a= 5' ACTCTGATGTATCTTCAGAACAGC 3'

S226A_R1b= 5' TAGCCATTAGAAAAAACTGTTCGAC 3'

Cas9 (final concentration 25 µM) was combined with gRNA (final concentration 30 µM each) and mixed before incubating for 10-20 minutes at room temperature. Nucleofector and supplement were combined in 20 µL cuvettes at room temperature. NALM6 cells were resuspended in the solution at 0.2e6 cells per 20 µL. Cells, HDR donor oligo (final concentration 4 µM each), and electroporation enhancer (final concentration 4 µM) were added to the Cas9 mixture and mixed gently. Cells were electroporated with the DS-142 program. Electroporated cells were resuspended in 75 µL of warmed media (RPMI1640+10% FBS+1% pen/strep) for total volume of 100 µL and transferred into one well of a

96 well plate containing 100 µL warmed media with HDR V2 enhancer (final concentration 1 µM). Final cell density was 1e6 cells per mL. Cells were incubated at 37°C for 12-24 hours before media was changed to remove HDR V2 enhancer.

After 48-72 hours, editing efficiency was checked with T7EI digest (Integrated DNA Technologies, cat#1075931). PCR reactions were prepared using 5X Phusion HF Buffer (New England Biolabs, cat#B0518S), dNTPs (ThermoFisher, cat#R1122, diluted to final concentration 200 µM), PCR primers (final concentration 0.5 µM each), and Phusion polymerase (1U/50µL). PCR products were used in the T7EI digestion according to the manufacturer's protocol. After the reaction, products were run on a 10% (29:1 acrylamide/bis-acrylamide) 1X Tris/Glycine PAGE gel for 60 min at 180V. Gel was stained with ethidium bromide and visualized on GE LAS4000 imager with UV box. Edited cells were single cell sorted on Becton Dickinson Aria II into 96 well plates. Clones were allowed to grow prior to extracting genomic DNA. Genomic DNA was then PCR amplified for the edited region using the same protocol as prior to T7EI digest. PCR products were purified with Qiagen MinElute PCR purification kit (cat#28004) following the manufacturer's protocol. Eluted DNA was Sanger sequenced, with control, experimental, and reference PCR products sequenced in parallel for TIDER analysis.

| Sample ID | NCI Risk Group | Karyotype | Fusion Genes Identified (if any) | Cytogenetics Grouping (non-infants/relapse) |
|---|---|---|---|---|
| MAP010 | SR | 29,XY,+4,+8,+10,+18,+21[20] | | Unfavorable |
| MAP011 | SR | 56,XY,+X,+Y,+4,+6,+10,+14,+17,+18,der(19) t(1;19)(q23;p13.3),+21,+21[12] | TCF3::PBX1 | Favorable |
| MAP012 | SR | 46,XY,t(6;8)(q25;q11.2)[8] | | Neutral |
| MAP013 | SR | 47,XY,dic(9;20)(p13;q11.2),+10,+21[4]/ 48,idem,+21[5]/48,idem,+X[3]/49,idem,+X,+ 21[2] | | Neutral |
| MAP014 | HR | 46,XX,i(9)(q10),t(12;13)(p13;q12)[23] | ETV6::FLT3 | Neutral |
| MAP015 | HR | 46,XY,del(9)(p13)x2,der(10)t(9;10)(p22;p15), der(12)t(9;12)(p22;q24.3),dup(16)(q13q11.2) [19] | | Neutral |
| MAP016 | SR | 46,XY[20] | ETV6::RUNX1 | Favorable |
| MAP017 | HR | 46,XX,der(2)t(2;9)(p11.2;q13),der(14)t(2;14) (p11.2;q32),dup(21)(q22)iAMP21[11] | | Unfavorable |
| MAP018 | N/A (infant) | 47,XX,add(2)(q33),der(3)t(3;5)(p26;q11.2),+6 ,del(10)(q24)[15] | BCR::ABL1 | N/A |
| MAP019 | N/A (relapse) | 49~50,Y,dup(X)(p11.2p22.1),+dup(X) (p11.2p22.1),t(1;7;5)(q21;p11.2;q11.2),del(6) (q23q27),+8,+9,+10,i(21)(q10),+mar[9] | | N/A |
| MAP020 | HR | 46,XY,i(7)(q10)[14] | P2RY8::CRLF2 | Ph-like |
| MAP021 | HR | 46,XY,der(19)t(1;19)(q23;p13.3)[10] | TCF3::PBX1 | Neutral |
| MAP023 | HR | 46,XX,add(12)(p11.2)[4] | ETV6::RUNX1 | Favorable |
| MAP025 | N/A (infant) | 48,XX,+X,t(4;11)(q21;q23),+21[17] | KMT2A::AFF1 | N/A |
| MAP031 | HR | 46,XY,t(7;15)(q22;q13)[18] | | Neutral |
| MAP032 | SR | 54,XY,+X,+4,+6,+14,+17,+18,+21,+21[13] | | Neutral |
| MAP033 | SR | 46,XY,dup(21)(q22)iAMP21[4]/46,XY,add(9) (p13),dup(21)(q22)iAMP21[8] | | Unfavorable |
| MAP035 | HR | 46,XX,t(4;11)(q21;q23)[10]/47,XX,+X,t(4;11) (q21;q23)[6]/47,XX,+X,t(4;11)(q21;q23), del(17)(p11.2)[5]/47,XX,+X,t(4;11)(q21;q23), i(17)(q10)[5] | KMT2A::AFF1 | Unfavorable |
| MAP036 | SR | 47,XX,+10[4] | ETV6::RUNX1 | Favorable |
| MAP039 | SR | 46,XY,t(1;3)(p34;q29)[24] | ETV6::RUNX1 | Favorable |

**Supplemental Table 1. Characteristics of primary patient specimens.** SR = standard risk. HR = high risk. Ph-like = Philadelphia chromosome-like ALL.

| Gene | Log2 Fold Change with 50nM Dex | Screen Phenotype |
|---|---|---|
| **NR3C1** | 1.129947279 | 0.998019174 |
| **NCOR2** | 1.0103191 | 0.400499412 |
| **TCF3** | 0.212719084 | 0.531101476 |
| **TRIP12** | 0.350989806 | 0.315524534 |
| **KAT6A** | 0.264583814 | 0.455455254 |
| **NCOR1** | 0.444274357 | 0.531481962 |
| **GPS2** | 0.423298552 | 0.334240629 |
| **NCOA1** | 0.632939634 | 0.3178194 |
| **DDX17** | 0.413690919 | 0.074999559 |
| **ZNF592** | 0.813638405 | 0.394779842 |
| **BCL2L11** | 1.114891533 | 0.265009516 |
| **ARID1A** | 0.371997379 | 0.338414391 |
| **NKTR** | 0.27800393 | 0.32458471 |
| **SETD1B** | 0.772636417 | 0.142520354 |
| **UBE2V1** | 0.326098713 | 0.087849676 |
| **HELZ** | 0.469395109 | 0.33976174 |
| **KIF1C** | 0.484704339 | 0.010654748 |
| **SNN** | 0.645638026 | 0.182118532 |
| **RAB5C** | 0.361174876 | 0.21392928 |
| **MED13L** | 0.608900741 | 0.285803227 |
| **ERLIN2** | 0.42217714 | 0.116513464 |
| **BMF** | 4.18793177 | 0.410481624 |
| **AFF1** | 0.941777597 | 0.089355658 |
| **PPP6R3** | 0.496508431 | 0.298601121 |
| **BIRC6** | 0.3529629 | 0.011858095 |
| **PCGF3** | 0.481364293 | 0.26222794 |
| **LAPTM5** | 0.908834258 | 0.225892787 |
| **TADA3** | 0.730576961 | 0.353623977 |
| **MED24** | 0.571802056 | 0.509794036 |
| **GNA13** | 0.271357985 | 0.191153257 |
| **USP19** | 0.452384901 | 0.176758223 |
| **DDX6** | 0.221815231 | 0.353756627 |
| **ZMYM4** | 0.325900338 | 0.178021651 |
| **ITPKB** | 0.430737367 | 0.156258067 |
| **SDE2** | 0.299691746 | 0.289492286 |
| **CD72** | 1.917252565 | 0.191689426 |
| **YTHDC1** | 0.31670777 | 0.114965401 |
| **GOLGA8A** | 0.483173555 | 0.267896256 |

| | | |
|---|---|---|
| **MT2A** | 0.518741673 | 0.277936592 |
| **UTRN** | 0.414901764 | 0.114246787 |
| **CD74** | 0.897726599 | 0.319921401 |
| **ARID4A** | 0.557281039 | 0.079339525 |
| **MDM4** | 0.400079606 | 0.137919589 |
| **DISP1** | 0.773541095 | 0.273947063 |
| **GLUL** | 1.127190419 | 0.283281814 |
| **TNNI2** | 5.357278771 | 0.360487963 |
| **PIAS1** | 0.51801577 | 0.082559794 |
| **NKAIN4** | 0.857235345 | 0.256362001 |
| **PLEKHA2** | 0.267818889 | 0.289528096 |
| **SPATA9** | 3.126715033 | 0.217078653 |
| **MPPE1** | 0.692653649 | 0.197709268 |
| **BECN1** | 0.238043795 | 0.217708769 |
| **WDR26** | 0.351572713 | 0.089585714 |
| **DMXL1** | 0.481571681 | 0.277516238 |
| **NAV1** | 0.273705643 | 0.227496439 |
| **CYP3A5** | 0.701376725 | 0.34684587 |
| **YIPF3** | 0.805642605 | 0.069816296 |
| **RASSF4** | 5.136331026 | 0.149637146 |
| **ATP11A** | 0.406236676 | 0.257377136 |
| **CPS1** | 0.425654452 | 0.071545292 |
| **MAGED4** | 3.026490011 | 0.174426836 |
| **PAG1** | 0.846559528 | 0.246944725 |
| **TTLL3** | 0.538275787 | 0.235482694 |
| **PRDM1** | 3.331907026 | 0.282467001 |
| **TCN2** | 2.281704137 | 0.125541603 |
| **DDHD1** | 0.387916231 | 0.065333776 |
| **TTC14** | 0.455355461 | 0.159652684 |
| **DOK4** | 1.746715046 | 0.211037827 |
| **VPS45** | 0.396547183 | 0.09589402 |
| **DNASE1** | 0.48640795 | 0.330198434 |
| **LRRC39** | 0.727109605 | 0.102499616 |
| **HIGD1B** | 6.437713053 | 0.048719912 |
| **SLFN5** | 1.998482062 | 0.046640726 |
| **AKIRIN2** | 0.547317461 | 0.099056326 |
| **YBX3** | 1.418937979 | 0.133789957 |
| **SAFB2** | 0.348149422 | 0.44797162 |
| **MFSD11** | 0.655926064 | 0.234709071 |

| | | |
|---|---|---|
| **PHF8** | 1.04238251 | 0.319835758 |
| **ASB3** | 0.231103662 | 0.203886243 |
| **PAXIP1** | 0.844566872 | 0.132193044 |
| **KLC4** | 0.547006107 | 0.329912981 |
| **RNF31** | 0.405856872 | 0.086177552 |
| **CHKB** | 0.859020692 | 0.334847227 |
| **PRDM11** | 1.610806427 | 0.246001806 |
| **DIP2C** | 1.522337071 | 0.322916977 |
| **TMEM106A** | 0.794987474 | 0.278557211 |
| **TMEM214** | 0.496873543 | 0.231038321 |
| **TGOLN2** | 0.220634229 | 0.057713929 |

**Supplemental Table 2A: Positive Effector Genes.** This table shows the intersection of the set of genes upregulated (adjp ≤ 0.05) in NALM6 cells treated with higher dose dexamethasone (50 nM Dex) with the set of genes with a positive phenotype (meaning contributing to dexamethasone-induced cell death) from our genome-wide shRNA screen. Upregulation of each gene is modeled to contribute to dexamethasone-induced cell death. Genes are ranked by significance of the phenotype (not shown).

| Gene | Log2 Fold Change with 50nM Dex | Screen Phenotype |
|---|---|---|
| **MBNL1** | -0.38873347 | -0.399080764 |
| **NUP153** | -0.405485208 | -0.439408595 |
| **CTCF** | -0.223221101 | -0.524249171 |
| **TCERG1** | -0.240320954 | -0.37893135 |
| **AHCTF1** | -0.487793069 | -0.341400418 |
| **CS** | -0.2506042 | -0.305489563 |
| **ZBTB33** | -0.240803232 | -0.343004389 |
| **DAZAP1** | -0.261479268 | -0.356056635 |
| **PAX5** | -0.627942746 | -0.374076749 |
| **U2AF2** | -0.548074341 | -0.323044549 |
| **BCL2** | -1.534811903 | -0.435740423 |
| **THAP11** | -0.345644396 | -0.269877259 |
| **EIF3J** | -0.384247945 | -0.364122208 |
| **PIK3CD** | -0.628700797 | -0.318161018 |
| **MEF2C** | -0.401521777 | -0.293278915 |
| **NCAPH** | -0.218318498 | -0.188847578 |
| **CCND3** | -0.596595347 | -0.362963361 |
| **EIF3I** | -0.541658314 | -0.102781228 |
| **KPNB1** | -0.435286351 | -0.086582065 |

| | | |
|---|---|---|
| **LEF1** | -1.372405174 | -0.054216607 |
| **SLC25A5** | -0.245704403 | -0.27667 |
| **ODC1** | -0.850244198 | -0.316348471 |
| **PSMB5** | -0.488030291 | -0.040513361 |
| **TOMM34** | -0.459833707 | -0.19016298 |
| **CREG1** | -0.571697457 | -0.320354864 |
| **CYP27B1** | -1.828541699 | -0.3092194 |
| **XAF1** | -0.819516192 | -0.218857998 |
| **LSM6** | -0.362324682 | -0.38803014 |
| **CFDP1** | -0.323089647 | -0.282424653 |
| **DEK** | -0.367694861 | -0.092970654 |
| **UBA1** | -0.244270723 | -0.295517172 |
| **PSMD13** | -0.272755068 | -0.301532593 |
| **IARS2** | -0.408579983 | -0.308634027 |
| **NAF1** | -0.511875069 | -0.36694396 |
| **IKZF2** | -0.818885042 | -0.140759446 |
| **ASNS** | -0.856984439 | -0.276440346 |
| **NANP** | -0.316155453 | -0.109209177 |
| **SNRPF** | -0.571737774 | -0.175515023 |
| **U2AF1** | -0.266898837 | -0.284413745 |
| **NAA25** | -0.539785666 | -0.290499593 |
| **PSME3** | -0.36396081 | -0.28910883 |
| **PSMA2** | -0.412881734 | -0.366351523 |
| **ETF1** | -0.439875811 | -0.26536202 |
| **TSR1** | -0.524879852 | -0.243294346 |
| **EIF5A** | -0.719875685 | -0.33999652 |
| **ZFP36L1** | -0.677647948 | -0.266019918 |
| **SMC2** | -0.333281408 | -0.400463058 |
| **LRPPRC** | -0.383267905 | -0.342400654 |
| **PSMD1** | -0.284612403 | -0.298361164 |
| **PSMA7** | -0.462816575 | -0.31175686 |
| **TFAM** | -0.533270294 | -0.401943021 |
| **WDR43** | -0.367911541 | -0.360655645 |
| **BYSL** | -0.973889829 | -0.350896884 |
| **POLR1B** | -0.678306709 | -0.285790958 |
| **CEBPZ** | -0.259967912 | -0.084534457 |
| **CARM1** | -0.673801562 | -0.252156164 |
| **HNRNPF** | -0.399803212 | -0.328283678 |
| **SET** | -0.449350454 | -0.300927743 |

| | | |
|---|---|---|
| **DDX18** | -0.494961678 | -0.22918775 |
| **TMEM11** | -0.375698363 | -0.349879557 |
| **ERCC6L** | -0.530244306 | -0.072627543 |
| **PNO1** | -0.641936265 | -0.295685961 |
| **POLD3** | -0.286757618 | -0.237216633 |
| **MRPL13** | -0.520519625 | -0.042322619 |
| **DPY30** | -0.500385699 | -0.292875924 |
| **PRIM1** | -0.565544086 | -0.30498711 |
| **HDAC2** | -0.327067292 | -0.326280038 |
| **DDX50** | -0.301940801 | -0.12188307 |
| **ACADM** | -0.35270397 | -0.275305436 |
| **ZFR** | -0.299197759 | -0.132933201 |
| **SFXN1** | -0.481939426 | -0.187508845 |
| **GTF3A** | -0.25343591 | -0.283577371 |
| **TP53RK** | -0.243226062 | -0.244165188 |
| **ACSL4** | -0.339685249 | -0.096621128 |
| **BRCA2** | -0.432215062 | -0.347950242 |
| **ESD** | -0.371077113 | -0.250512906 |
| **C1QBP** | -0.608815913 | -0.128796941 |
| **XRCC2** | -0.732430207 | -0.239096266 |
| **KDM1A** | -0.379880399 | -0.281137519 |
| **TEAD4** | -0.958246261 | -0.227342731 |
| **RPS27L** | -0.44197907 | -0.2800026 |
| **TUFM** | -0.275400663 | -0.403280033 |
| **BHLHE40** | -0.969779825 | -0.308311663 |
| **CBFB** | -0.286946045 | -0.105476109 |
| **HIF1A** | -0.275205247 | -0.154125302 |
| **AKAP1** | -0.467881829 | -0.328694129 |
| **MSH2** | -0.414150603 | -0.056791155 |
| **DDX39A** | -0.334569189 | -0.104735623 |
| **NUP107** | -0.200895813 | -0.311933203 |
| **PWP1** | -0.322065118 | -0.06561147 |
| **ANP32A** | -0.300411397 | -0.000925732 |
| **CCNA2** | -0.401240221 | -0.258833711 |
| **SNAPC1** | -0.611020929 | -0.073791913 |
| **AIP** | -0.365159536 | -0.248705445 |
| **LGALS3BP** | -1.031720771 | -0.102449586 |
| **AMMECR1** | -1.252138893 | -0.269326102 |
| **NCAPG** | -0.293099604 | -0.383112125 |

| | | |
|---|---|---|
| **PUF60** | -0.542155456 | -0.053419241 |
| **MAPKAPK5** | -0.409034543 | -0.093171906 |
| **CBR1** | -0.926342149 | -0.099168652 |
| **SRSF2** | -0.245976816 | -0.20285831 |
| **RUVBL1** | -0.489236797 | -0.094602185 |
| **PA2G4** | -0.63507859 | -0.231584862 |
| **FANCB** | -0.649610925 | -0.173557611 |
| **RCC1** | -0.714798295 | -0.369181294 |
| **SFXN3** | -0.426765857 | -0.344642711 |
| **HAUS1** | -0.216276033 | -0.263290897 |
| **CGREF1** | -0.648438926 | -0.08779445 |
| **TMEM126A** | -0.537203696 | -0.224218224 |
| **PSMA3** | -0.741745347 | -0.226243312 |
| **DIS3L** | -0.382866164 | -0.081470744 |
| **PARPBP** | -0.434224179 | -0.33520505 |
| **SNRPE** | -0.528817915 | -0.031686721 |
| **CBX4** | -0.706002663 | -0.354953337 |
| **PRR3** | -0.579798052 | -0.165591202 |
| **MYO19** | -0.340452902 | -0.060605128 |
| **HDDC2** | -0.339180143 | -0.098717465 |
| **MRPS27** | -0.342411091 | -0.144749788 |
| **PPP2R1B** | -0.415345087 | -0.226312483 |
| **WDR61** | -0.386915296 | -0.084596546 |
| **SRSF7** | -0.524938444 | -0.344866239 |
| **BLNK** | -0.374984773 | -0.072907524 |
| **MDM2** | -0.622620157 | -0.306767959 |
| **MILR1** | -0.586388886 | -0.24488968 |
| **CD9** | -0.386514185 | -0.298034931 |
| **TDG** | -0.283585408 | -0.305784243 |
| **PIK3C2B** | -0.748072484 | -0.373648286 |
| **HNRNPU** | -0.450377669 | -0.09718176 |
| **BRCA1** | -0.303466226 | -0.259040509 |
| **NOP10** | -0.405639265 | -0.259386648 |
| **MATR3** | -0.30352306 | -0.048991735 |
| **CD19** | -0.394774021 | -0.079173995 |
| **PRDX3** | -0.470683733 | -0.053819522 |
| **PSMA5** | -0.482298669 | -0.05672641 |
| **CSTF2** | -0.453408418 | -0.083154227 |
| **LIN7C** | -0.264296931 | -0.214370835 |

| | | |
|---|---|---|
| **BOP1** | -0.517115171 | -0.297535637 |
| **POLR1A** | -0.284075524 | -0.155418588 |
| **ZNF43** | -0.366877005 | -0.369779214 |
| **PCNP** | -0.298111407 | -0.248840637 |
| **MYBBP1A** | -0.63510918 | -0.329606394 |
| **CYC1** | -0.467386008 | -0.142080423 |
| **IRF2** | -0.479184043 | -0.352041062 |
| **SNRPD3** | -0.472651757 | -0.323950497 |
| **LRFN1** | -1.207668718 | -0.364654442 |
| **TRIB3** | -1.582777284 | -0.133805856 |
| **MME** | -0.835898779 | -0.412681241 |
| **ECHS1** | -0.527356745 | -0.19675946 |

**Supplemental Table 2B: Negative Effector Genes.** This table shows the intersection of the set of genes upregulated (adjp ≤ 0.05) in NALM6 cells treated with higher dose dexamethasone (50 nM Dex) with the set of genes with a positive phenotype (meaning contributing to dexamethasone-induced cell death) from our genome-wide shRNA screen. Downregulation of each gene is modeled to contribute to dexamethasone-induced cell death. Genes are ranked by significance of the phenotype (not shown).

| Additive Effectors - Dexamethasone | Synergistic Effectors - Dexamethasone | Overlapping Effectors - Dexamethasone | Additive Effectors - Prednisolone | Synergistic Effectors - Prednisolone | Overlapping Effectors - Prednisolone |
|---|---|---|---|---|---|
| ACADM | AFF1 | AFF1 | ACADM | AFF1 | AFF1 |
| ADNP | ANKRD11 | ARID1A | AFF1 | BCOR | EP300 |
| AFF1 | ARID1A | BCL2L11 | ARID1A | C17orf49 | IRAK4 |
| ARID1A | BBX | BCOR | BCL2 | EP300 | MBNL1 |
| BCL2 | BCL2L11 | BMF | BMF | ETV6 | PAX5 |
| BCL2L11 | BCOR | BRD2 | BOP1 | IRAK4 | POU2F1 |
| BCOR | BMF | BRD4 | CARM1 | LEF1 | PRR12 |
| BIRC5 | BRD2 | C17orf49 | CHAMP1 | MBNL1 | RUVBL1 |
| BMF | BRD4 | CD79A | CREBBP | MTMR4 | SPEN |
| BOP1 | C17orf49 | CHAMP1 | DLGAP5 | NCOA1 | SRRM1 |
| BRD2 | CARS2 | EHMT2 | DOLPP1 | NUP214 | SSRP1 |
| BRD4 | CD79A | EP300 | EHMT2 | PAX5 | |
| C17orf49 | CDC42 | ETV6 | EIF3I | POU2F1 | |
| CARM1 | CHAMP1 | GPS2 | EP300 | PRR12 | |
| CD79A | CNOT2 | GSK3A | IRAK4 | RUVBL1 | |
| CELF1 | CPEB3 | HIF1A | LARP1 | SPEN | |
| CHAMP1 | CREBBP | IRAK4 | MBNL1 | SPI1 | |
| CTCF | EBF1 | KAT6A | NLE1 | SRRM1 | |
| DLGAP5 | EHMT2 | MBNL1 | NOL6 | SSRP1 | |
| DOLPP1 | EIF4E2 | MEF2A | NR3C1 | SYK | |
| EHMT2 | EP300 | MMP14 | PAX5 | ZNF608 | |
| EIF2B1 | ETV6 | MSI2 | PDCD5 | | |
| EIF3I | GPS2 | NCK1 | PIK3CD | | |
| EIF3L | GSK3A | NCOR2 | PLAGL2 | | |
| EP300 | HIF1A | PAX5 | POU2F1 | | |
| ETV6 | IRAK4 | PIK3CD | PPP5C | | |
| GPS2 | KAT6A | POLG | PRDM1 | | |
| GSK3A | LEF1 | POU2F1 | PREX1 | | |
| HIF1A | MBNL1 | PRDM1 | PRR12 | | |
| IRAK4 | MED23 | PREX1 | PTBP1 | | |
| ITPKB | MEF2A | PRR12 | RRP12 | | |
| KAT6A | MMP14 | RRP12 | RUVBL1 | | |
| LARP1 | MSI2 | RUVBL1 | SAFB2 | | |
| MAML2 | MTMR4 | SAFB2 | SPEN | | |
| MAPK1 | NCK1 | SPEN | SRRM1 | | |
| MBNL1 | NCOA1 | SRRM1 | SSRP1 | | |
| MED11 | NCOR2 | SSRP1 | SUPT16H | | |
| MED13 | NUP214 | ZMIZ1 | YTHDC1 | | |
| MEF2A | PARD6B | ZNF592 | ZNF320 | | |
| MMP14 | PAX5 | | ZNF638 | | |

| | | | |
|---|---|---|---|
| MSI2 | PHF6 | | ZNF671 |
| NCK1 | PIK3CD | | |
| NCOA2 | POLG | | |
| NCOR2 | POU2F1 | | |
| NELFCD | PRDM1 | | |
| NLE1 | PREX1 | | |
| NOL6 | PRKAB1 | | |
| NR3C1 | PRR12 | | |
| PAX5 | RGS9 | | |
| PDCD5 | RRP12 | | |
| PHC3 | RUVBL1 | | |
| PIK3CD | SAFB2 | | |
| PLAGL2 | SESN3 | | |
| POLG | SPEN | | |
| POU2F1 | SPI1 | | |
| PPP1R12A | SRRM1 | | |
| PPP5C | SSRP1 | | |
| PRC1 | SYK | | |
| PRDM1 | TAF3 | | |
| PREX1 | ZMIZ1 | | |
| PRR12 | ZNF592 | | |
| PTBP1 | ZNF608 | | |
| RASSF4 | | | |
| RAVER1 | | | |
| RBMX2 | | | |
| RRP12 | | | |
| RUVBL1 | | | |
| SAFB | | | |
| SAFB2 | | | |
| SETD1A | | | |
| SPEN | | | |
| SRRM1 | | | |
| SSRP1 | | | |
| SUPT16H | | | |
| TADA3 | | | |
| THOC2 | | | |
| WIZ | | | |
| YTHDC1 | | | |
| ZBED4 | | | |
| ZMIZ1 | | | |
| ZMYM4 | | | |
| ZMYND8 | | | |
| ZNF320 | | | |

| |
|---|
| ZNF592 |
| ZNF638 |
| ZNF671 |

**Supplemental Table 3: Effector Genes in Primary Patient Specimens.** Genes significantly regulated (adjp <= 0.01) in primary patient specimens treated with dexamethasone or prednisolone were intersected with genes which had a significant phenotype (p value < 0.05) in our genome-wide shRNA screens. These effector genes were identified for both dexamethasone and prednisolone in primary specimens defined as having an additive response to prednisolone with idelalisib in cell viability assays (Additive Effectors) and in specimens with a synergistic response in viability assays (Synergistic Effectors). Overlapping effectors were determined by intersecting the additive effectors and synergistic effectors for dexamethasone or prednisolone.

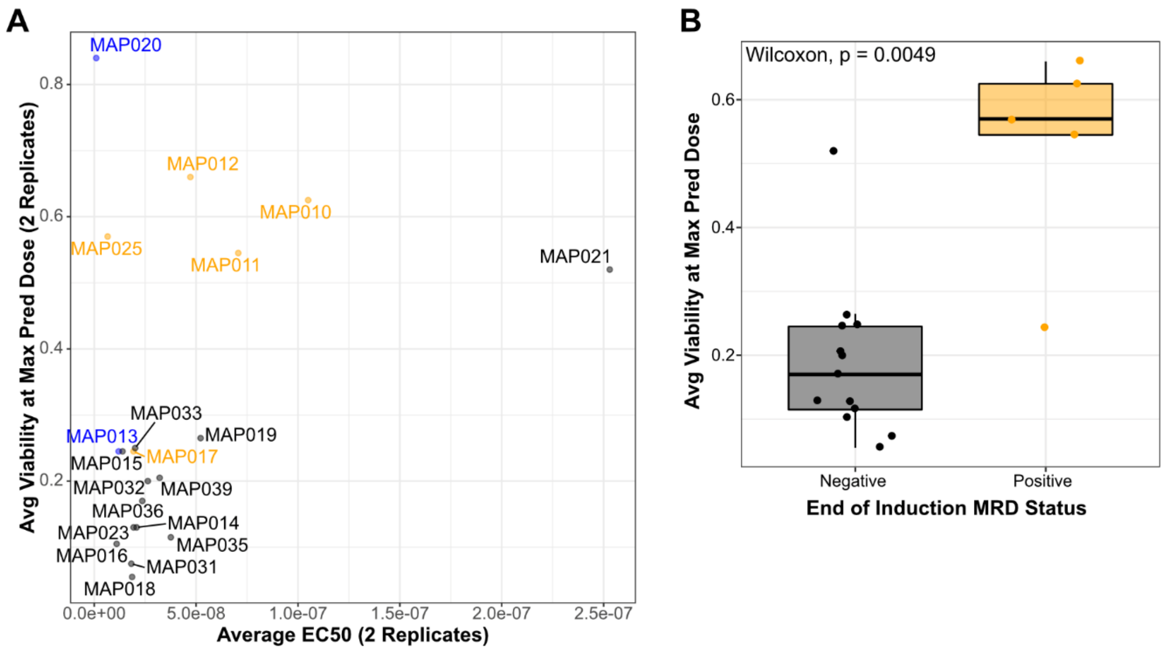

**Supplemental Figure 1.** ***In vitro*** **specimen viability after prednisolone treatment correlates with patient response.** (**A**) Response of primary specimens by EC50 of prednisolone (x-axis) and cell viability fraction at maximum tested dose of prednisolone (10 µM) (y-axis). Specimens with positive end of induction minimal residual disease (MRD) are shown in orange, and specimens with unknown end of induction MRD are in blue. The remaining specimens in black were MRD negative. (**B**) Boxplots comparing the average viability of primary patient specimens treated with the maximum dose of prednisolone (10 uM) *in vitro* for specimens obtained from patients who had negative end of induction minimal residual disease (MRD) status (left, black) or positive end of induction MRD status (right, orange).

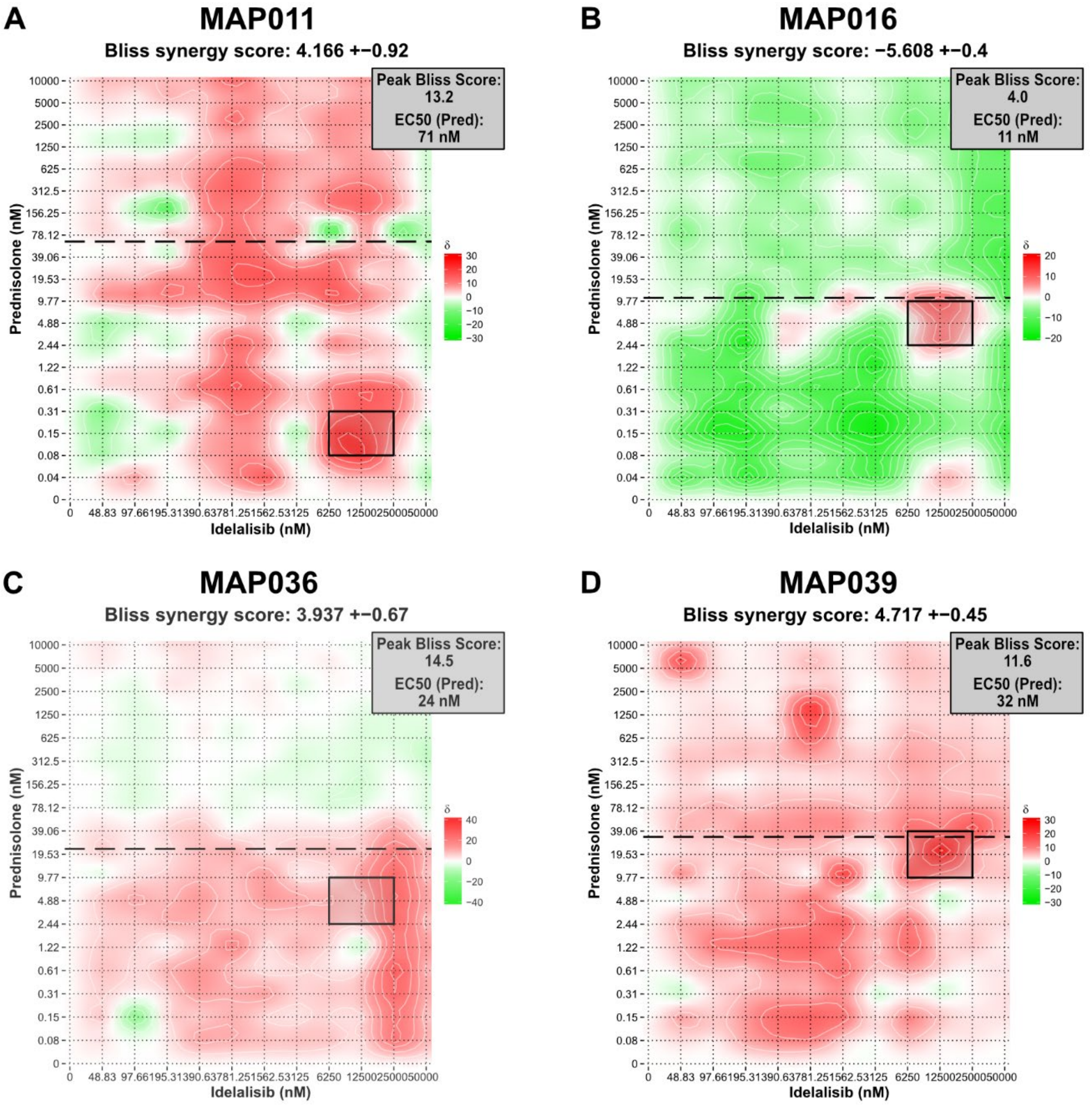

**Supplemental Figure 2A. Bliss synergy plots for standard risk primary specimens treated with prednisolone and idelalisib.** (**A-D**) NCI standard risk specimens with favorable cytogenetic features. The area of peak Bliss score is outlined in black on each plot with the value given in the box at the top right of each plot. EC50 for prednisolone is the average of 2 replicates per specimen and is given in the box at the top right of each plot. The approximate location of the EC50 is indicated on each plot by the long dashed black line.

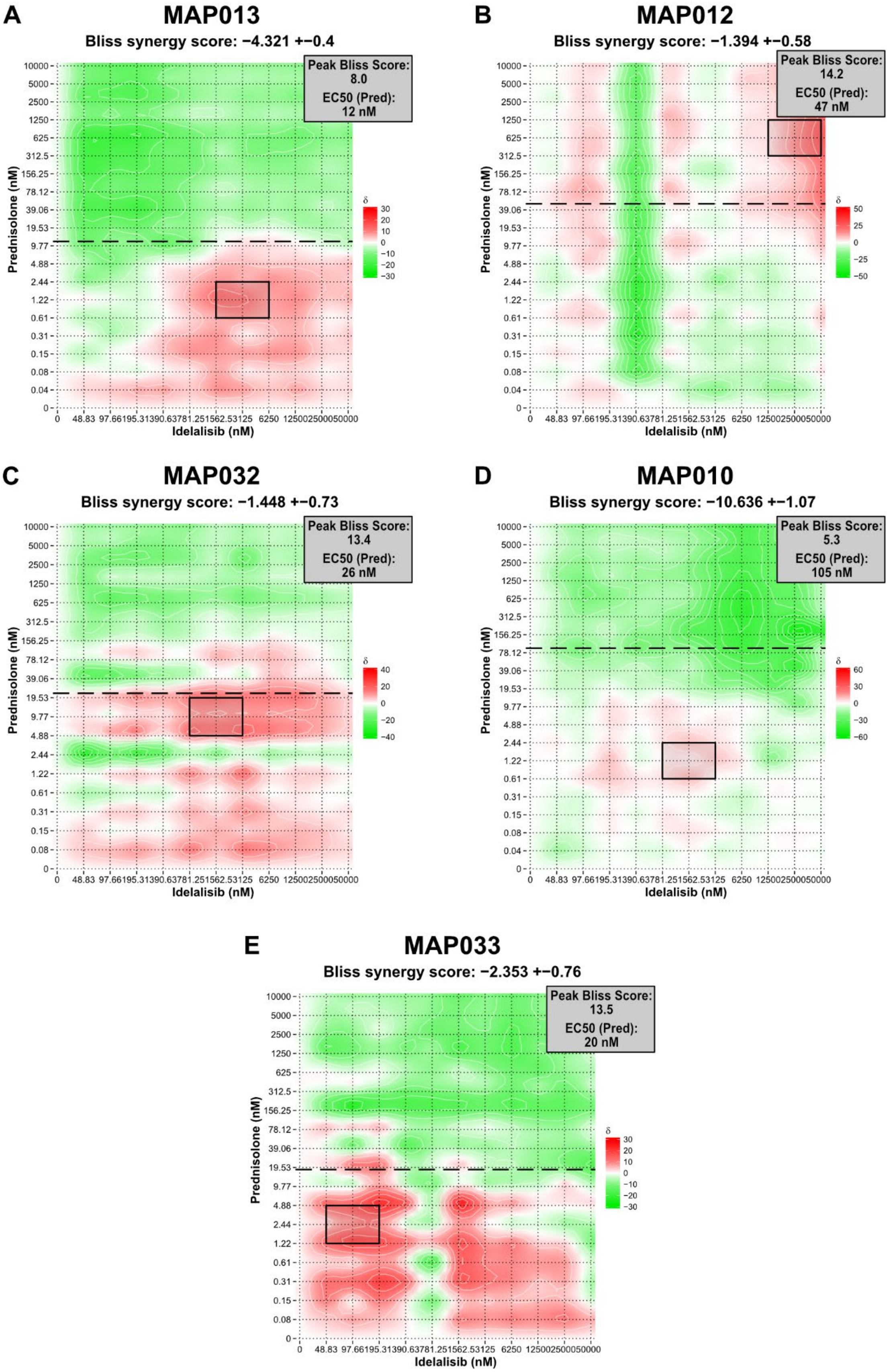
A
MAP013
Bliss synergy score: −4.321 +−0.4
Peak Bliss Score: 8.0
EC50 (Pred): 12 nM
B
MAP012
Bliss synergy score: −1.394 +−0.58
Peak Bliss Score: 14.2
EC50 (Pred): 47 nM
C
MAP032
Bliss synergy score: −1.448 +−0.73
Peak Bliss Score: 13.4
EC50 (Pred): 26 nM
D
MAP010
Bliss synergy score: −10.636 +−1.07
Peak Bliss Score: 5.3
EC50 (Pred): 105 nM
E
MAP033
Bliss synergy score: −2.353 +−0.76
Peak Bliss Score: 13.5
EC50 (Pred): 20 nM
Prednisolone (nM)
Idelalisib (nM)
δ

**Supplemental Figure 2B. Bliss synergy plots for standard risk primary specimens treated with prednisolone and idelalisib.** (**A-C**) NCI standard risk specimens with neutral cytogenetic features, and (**D-E**) NCI standard risk specimens with unfavorable cytogenetic features. The area of peak Bliss score is outlined in black on each plot with the value given in the box at the top right of each plot. EC50 for prednisolone is the average of 2 replicates per specimen and is given in the box at the top right of each plot. The approximate location of the EC50 is indicated on each plot by the long dashed black line.

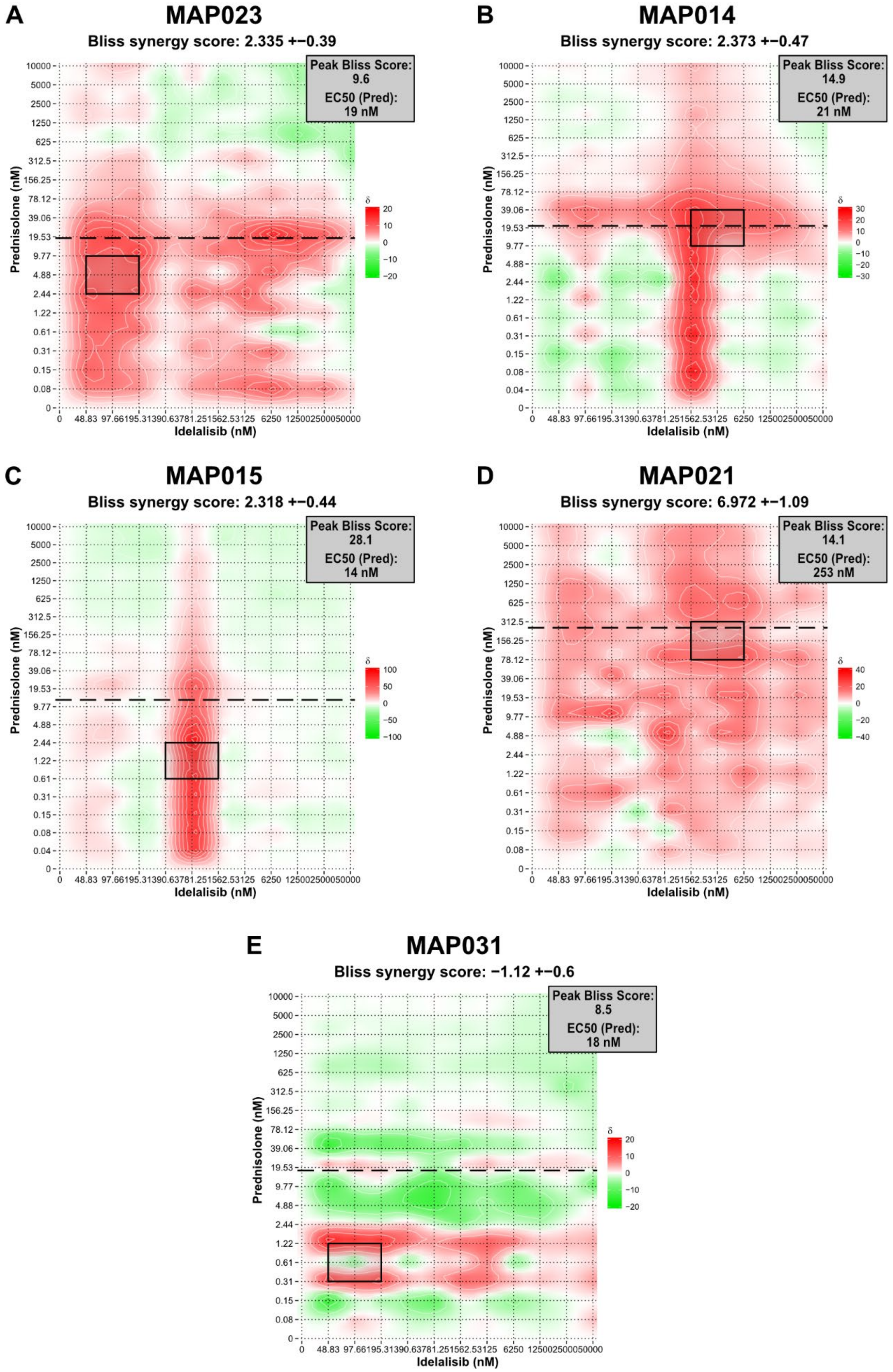
A
MAP023
Bliss synergy score: 2.335 +−0.39
Peak Bliss Score: 9.6
EC50 (Pred): 19 nM
B
MAP014
Bliss synergy score: 2.373 +−0.47
Peak Bliss Score: 14.9
EC50 (Pred): 21 nM
C
MAP015
Bliss synergy score: 2.318 +−0.44
Peak Bliss Score: 28.1
EC50 (Pred): 14 nM
D
MAP021
Bliss synergy score: 6.972 +−1.09
Peak Bliss Score: 14.1
EC50 (Pred): 253 nM
E
MAP031
Bliss synergy score: −1.12 +−0.6
Peak Bliss Score: 8.5
EC50 (Pred): 18 nM
Prednisolone (nM)
Idelalisib (nM)
δ
10000
5000
2500
1250
625
312.5
156.25
78.12
39.06
19.53
9.77
4.88
2.44
1.22
0.61
0.31
0.15
0.08
0.04
0
0 48.83 97.66195.31390.63781.251562.53125 6250 125002500050000

**Supplemental Figure 2C. Bliss synergy plots for high risk primary specimens treated with prednisolone and idelalisib.** (**A**) NCI high risk specimen with favorable cytogenetic features, and (**B-E**) NCI high risk specimens with neutral cytogenetic features. The area of peak Bliss score is outlined in black on each plot with the value given in the box at the top right of each plot. EC50 for prednisolone is the average of 2 replicates per specimen and is given in the box at the top right of each plot. The approximate location of the EC50 is indicated on each plot by the long dashed black line.

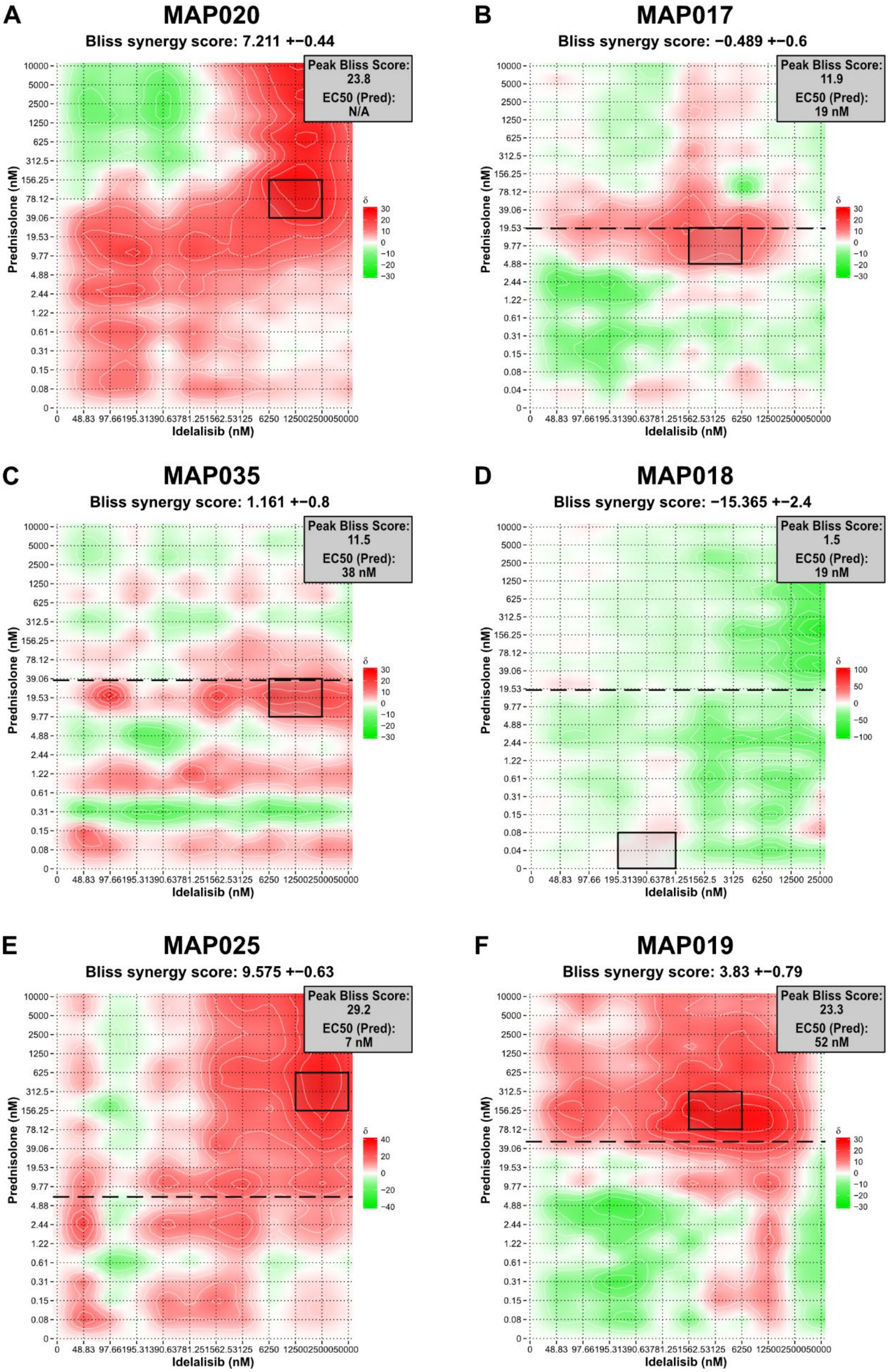
A
MAP020
Bliss synergy score: 7.211 +−0.44
Peak Bliss Score: 23.8
EC50 (Pred): N/A
B
MAP017
Bliss synergy score: −0.489 +−0.6
Peak Bliss Score: 11.9
EC50 (Pred): 19 nM
C
MAP035
Bliss synergy score: 1.161 +−0.8
Peak Bliss Score: 11.5
EC50 (Pred): 38 nM
D
MAP018
Bliss synergy score: −15.365 +−2.4
Peak Bliss Score: 1.5
EC50 (Pred): 19 nM
E
MAP025
Bliss synergy score: 9.575 +−0.63
Peak Bliss Score: 29.2
EC50 (Pred): 7 nM
F
MAP019
Bliss synergy score: 3.83 +−0.79
Peak Bliss Score: 23.3
EC50 (Pred): 52 nM
Prednisolone (nM)
Idelalisib (nM)
δ

**Supplemental Figure 2D. Bliss synergy plots for additional primary specimens treated with prednisolone and idelalisib.** (**A**) NCI high risk specimen with CRLF2 rearrangement, (**B-C**) NCI high risk specimens with unfavorable cytogenetic features, (**D-E**) specimens from infants with B-ALL, and (**F**) specimen from first late marrow relapse of B-ALL. The area of peak Bliss score is outlined in black on each plot with the value given in the box at the top right of each plot. EC50 for prednisolone is the average of 2 replicates per specimen and is given in the box at the top right of each plot, except for MAP020 (**A**) which was insensitive to prednisolone. The approximate location of the EC50 is indicated on each plot by the long dashed black line.

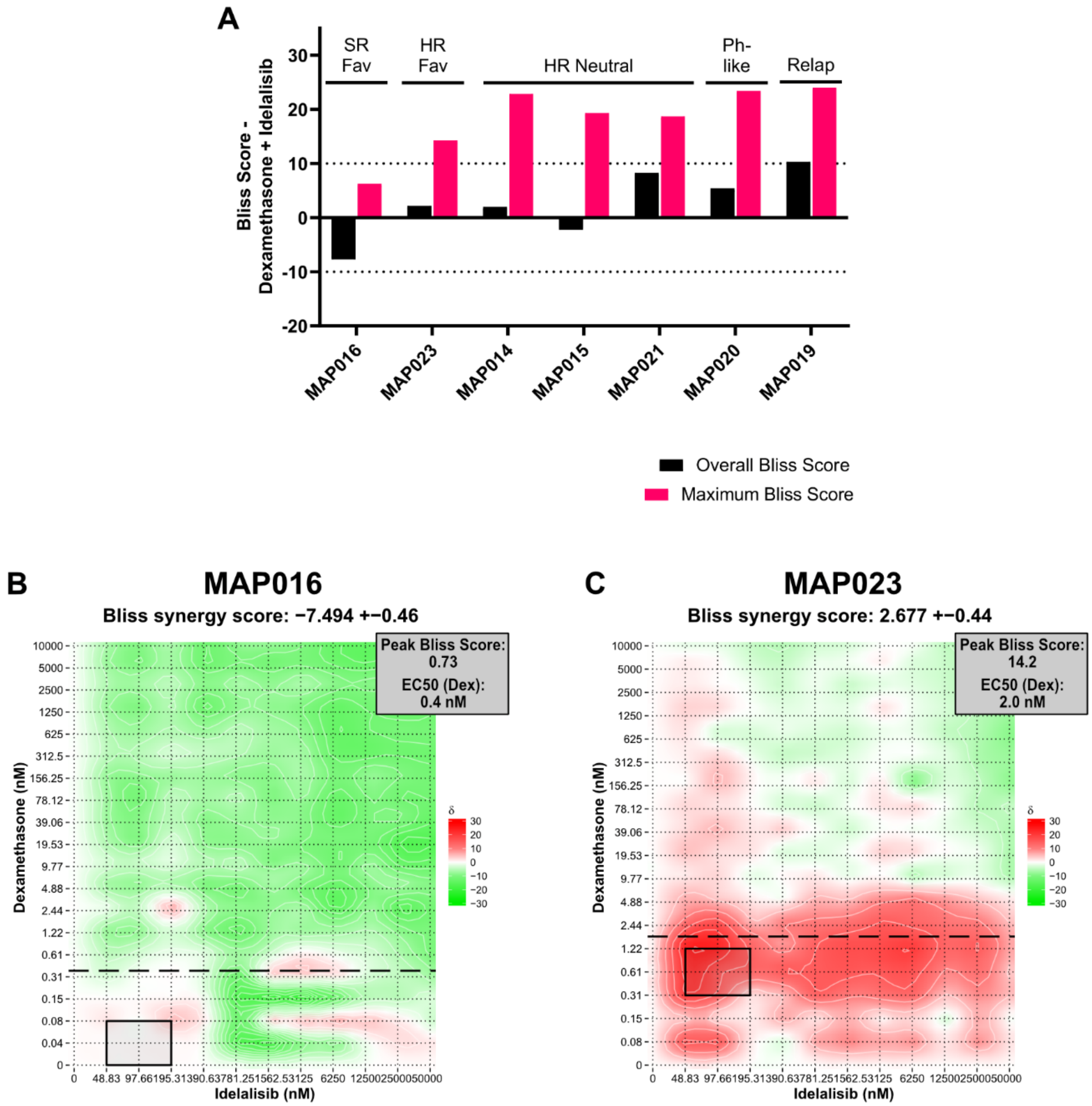

**Supplemental Figure 3A. Combination dexamethasone and idelalisib treatment in primary specimens produces synergy similar to prednisolone and idelalisib.** (**A**) Summary of overall (black) and maximum/peak (pink) Bliss scores for seven primary specimens. SR = NCI Standard Risk. HR = NCI High Risk. Fav = favorable cytogenetics. Relap = relapsed B-ALL. (**B-C**) Bliss synergy plots for primary specimens treated with dexamethasone and idelalisib with favorable cytogenetic features and (**B**) NCI standard risk B-ALL or (**C**) NCI high risk B-ALL. The area of peak Bliss score is outlined in black on each plot with the value given in the box at the top right of each plot. EC50 for dexamethasone is the average of 2 replicates per specimen and is given in the box at the top right of each plot. The approximate location of the EC50 is indicated on each plot by the long dashed black line.

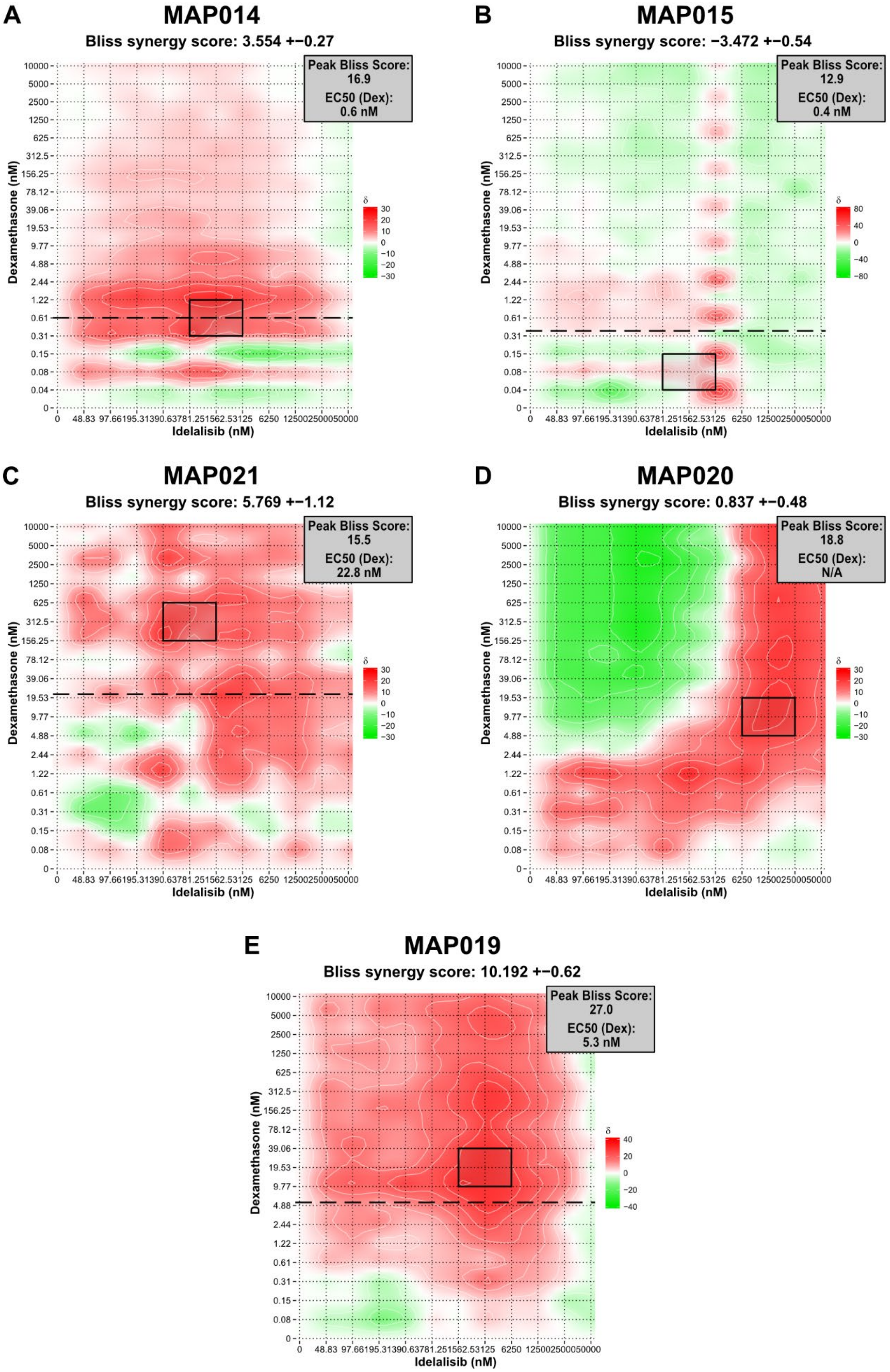
A
MAP014
Bliss synergy score: 3.554 +−0.27
Peak Bliss Score: 16.9
EC50 (Dex): 0.6 nM
B
MAP015
Bliss synergy score: −3.472 +−0.54
Peak Bliss Score: 12.9
EC50 (Dex): 0.4 nM
C
MAP021
Bliss synergy score: 5.769 +−1.12
Peak Bliss Score: 15.5
EC50 (Dex): 22.8 nM
D
MAP020
Bliss synergy score: 0.837 +−0.48
Peak Bliss Score: 18.8
EC50 (Dex): N/A
E
MAP019
Bliss synergy score: 10.192 +−0.62
Peak Bliss Score: 27.0
EC50 (Dex): 5.3 nM
Dexamethasone (nM)
Idelalisib (nM)
δ

**Supplemental Figure 3B. Combination dexamethasone and idelalisib treatment in primary specimens produces synergy similar to prednisolone and idelalisib.** Bliss synergy plots for primary specimens treated with dexamethasone and idelalisib with (**A-C**) NCI high risk B-ALL with neutral cytogenetic features, (**D**) NCI high risk B-ALL with CRLF2 rearrangement, or (**E**) first late marrow relapse of B-ALL. The area of peak Bliss score is outlined in black on each plot with the value given in the box at the top right of each plot. EC50 for dexamethasone is the average of 2 replicates per specimen and is given in the box at the top right of each plot, except for MAP020 (**D**) which was insensitive to dexamethasone. The approximate location of the EC50 is indicated on each plot by the long dashed black line.

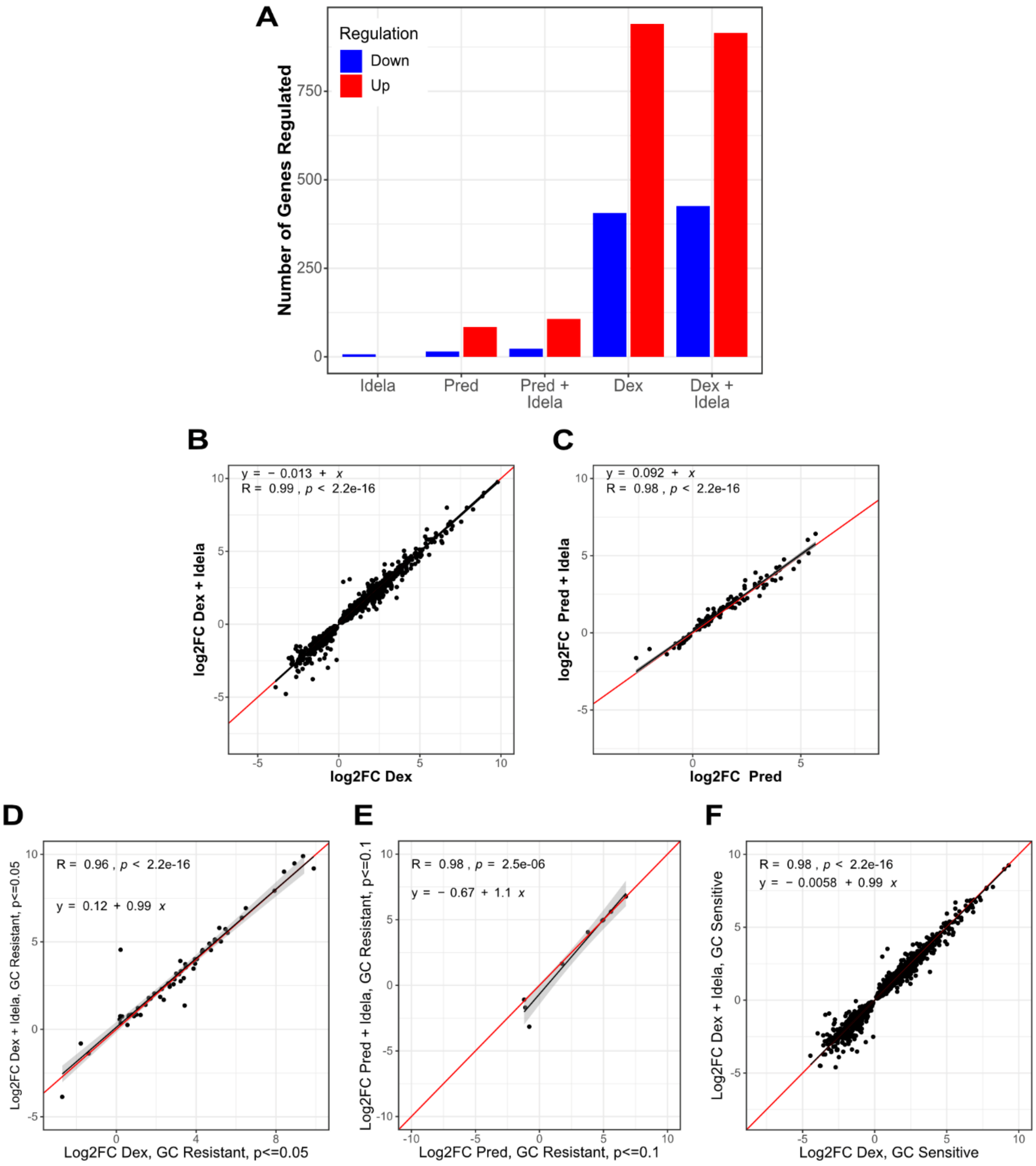

**Supplemental Figure 4. Gene expression in glucocorticoid-resistant and glucocorticoid-sensitive primary patient specimens treated with glucocorticoids with and without idelalisib.** (**A**) Overall number of genes upregulated (red) and downregulated (blue) in all 7 primary patient specimens treated with idelalisib only (idela, 500 nM), prednisolone only (pred, 25-50 nM), prednisolone + idelalisib (pred+idela), dexamethasone only (dex, 25-50 nM), or dexamethasone + idelalisib (dex+idela). (**B-C**) Comparison of gene expression with (**B**) dexamethasone only (x-axis) versus dexamethasone plus idelalisib (y-axis) and (**C**) prednisolone only (x-axis) versus prednisolone plus idelalisib (y-axis) in all 7 primary specimens. (**D-E**) Comparison of gene expression log2 fold change in 2 glucocorticoid-resistant specimens with (**D**) dexamethasone only (x-axis) versus dexamethasone plus idelalisib (y-axis) and (**E**) prednisolone only (x-axis) versus prednisolone plus idelalisib (y-axis). (**F**) Comparison of gene expression log2 fold change in 5 glucocorticoid-sensitive specimens treated with dexamethasone only (x-axis) versus dexamethasone plus idelalisib (y-axis). For all plots, the Pearson correlation and regression equations are reported.

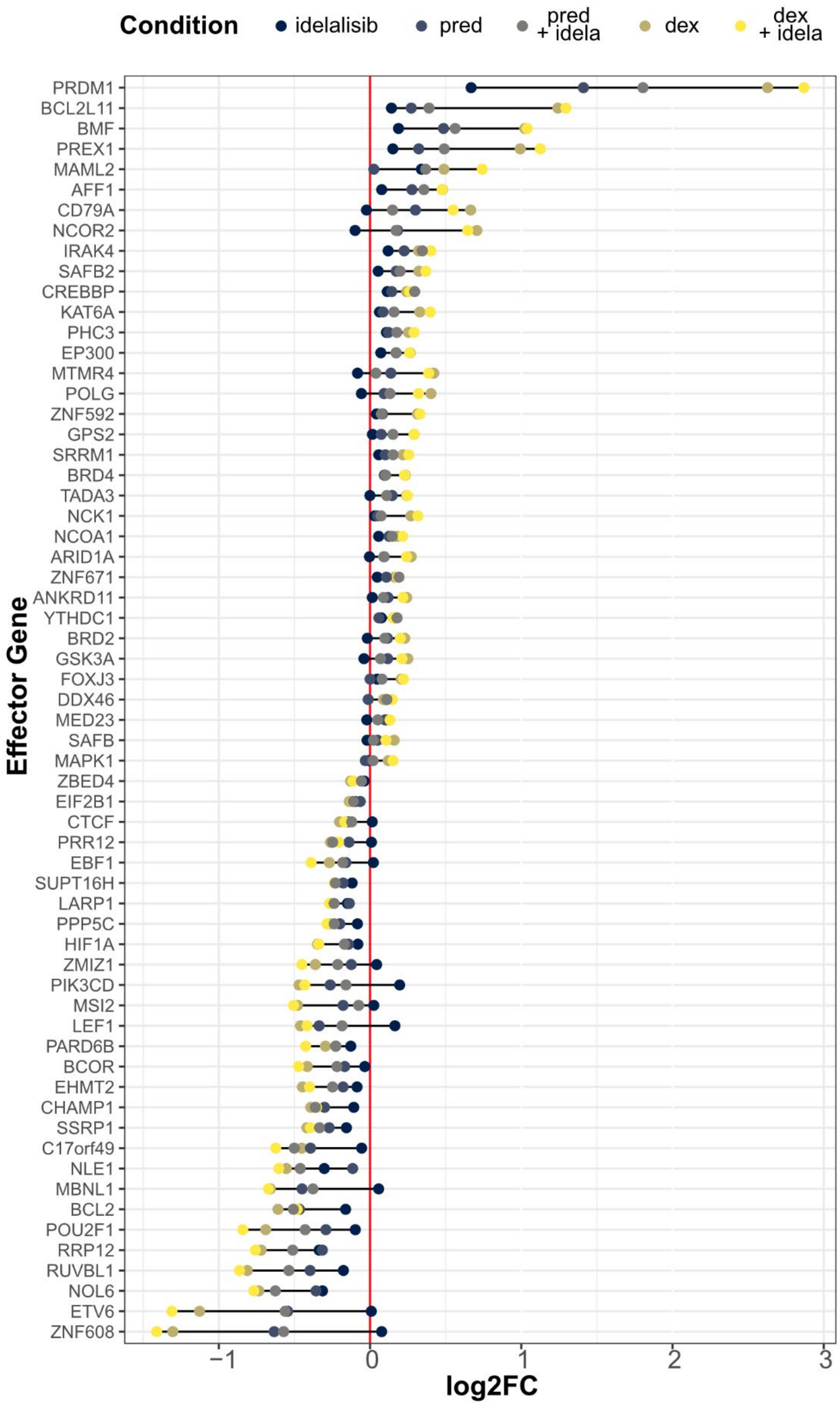

Condition
idelalisib
pred
pred + idela
dex
dex + idela
PRDM1
BCL2L11
BMF
PREX1
MAML2
AFF1
CD79A
NCOR2
IRAK4
SAFB2
CREBBP
KAT6A
PHC3
EP300
MTMR4
POLG
ZNF592
GPS2
SRRM1
BRD4
TADA3
NCK1
NCOA1
ARID1A
ZNF671
ANKRD11
YTHDC1
BRD2
GSK3A
FOXJ3
DDX46
MED23
SAFB
MAPK1
ZBED4
EIF2B1
CTCF
PRR12
EBF1
SUPT16H
LARP1
PPP5C
HIF1A
ZMIZ1
PIK3CD
MSI2
LEF1
PARD6B
BCOR
EHMT2
CHAMP1
SSRP1
C17orf49
NLE1
MBNL1
BCL2
POU2F1
RRP12
RUVBL1
NOL6
ETV6
ZNF608
Effector Gene
−1
0
1
2
3
log2FC

**Supplemental Figure 5A. Glucocorticoid-sensitive primary specimens exhibit similar changes in effector gene expression to NALM6 cells.** The log2 fold change of effector genes in response to combination dexamethasone (dex) and idelalisib (idela) treatment.

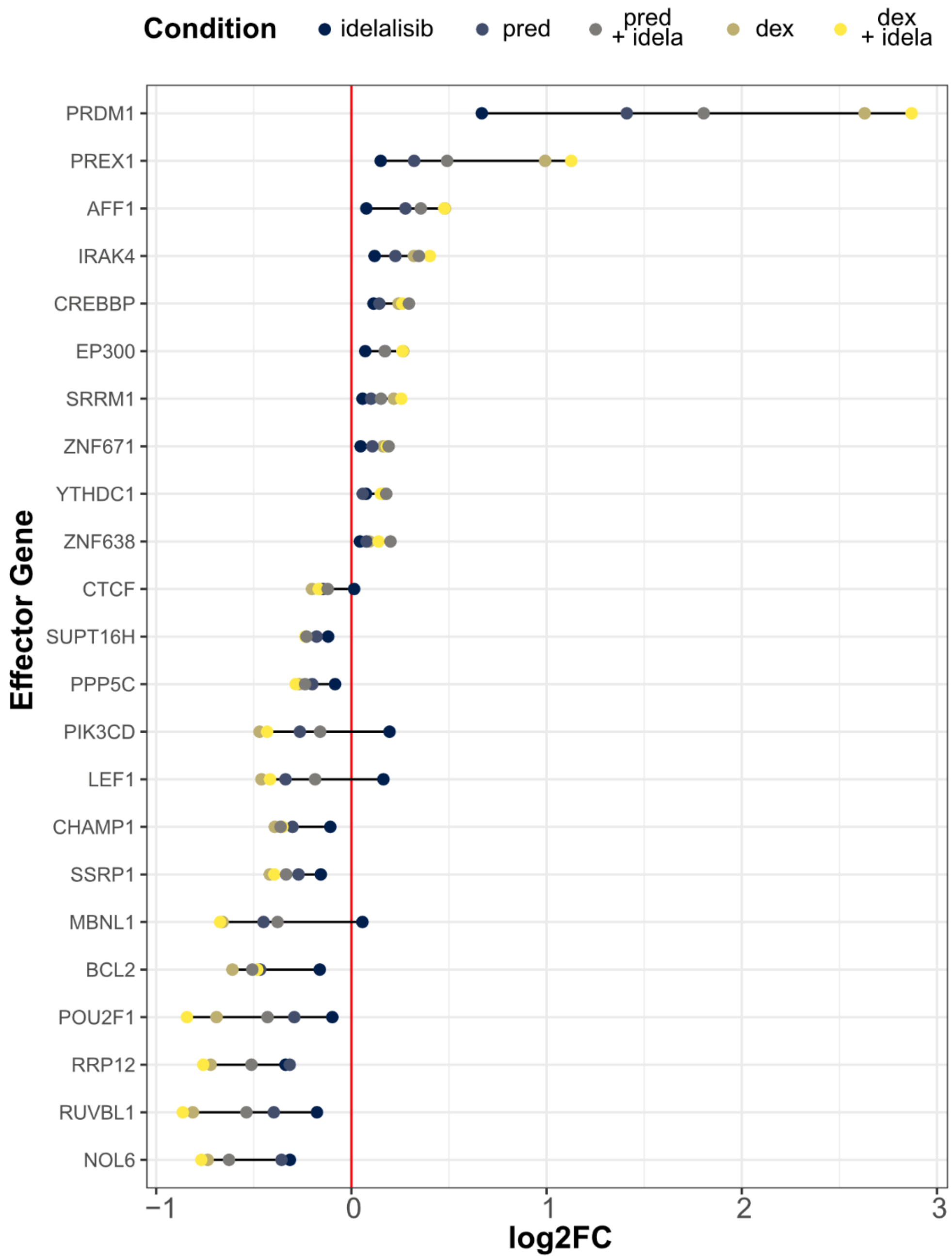

**Supplemental Figure 5B. Glucocorticoid-sensitive primary specimens exhibit similar changes in effector gene expression to NALM6 cells.** The log2 fold change of effector genes in response to combination prednisolone (pred) and idelalisib (idela) treatment.

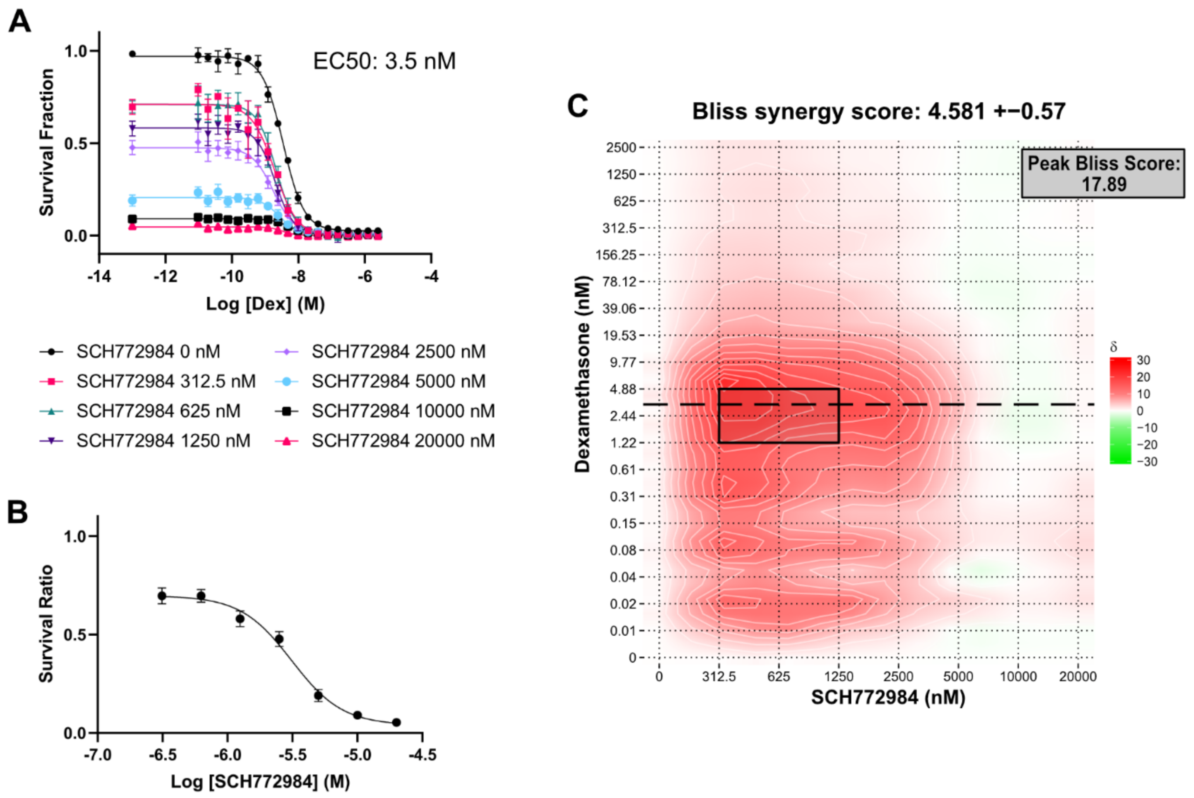

**Supplemental Figure 6. Synergy evaluation of NALM6 cells treated with dexamethasone and the ERK1/2 inhibitor SCH772984.** (A) Dose-response curves for dexamethasone (Dex) at different doses of SCH772984, with 3 replicates per concentration. Dexamethasone alone (SCH772984 0 nM) is shown in the top black line. (B) Dose-response curve for SCH772984 alone, 3 replicates per concentration. (C) Synergy plot of the combination treatment. Bliss score greater than 10 indicates synergy, 10 to -10 indicates additivity, and less than -10 indicates antagonism. Peak Bliss score from the black outlined area is given in the box below the plot. Long dashed black line indicates the approximate EC50 for dexamethasone.

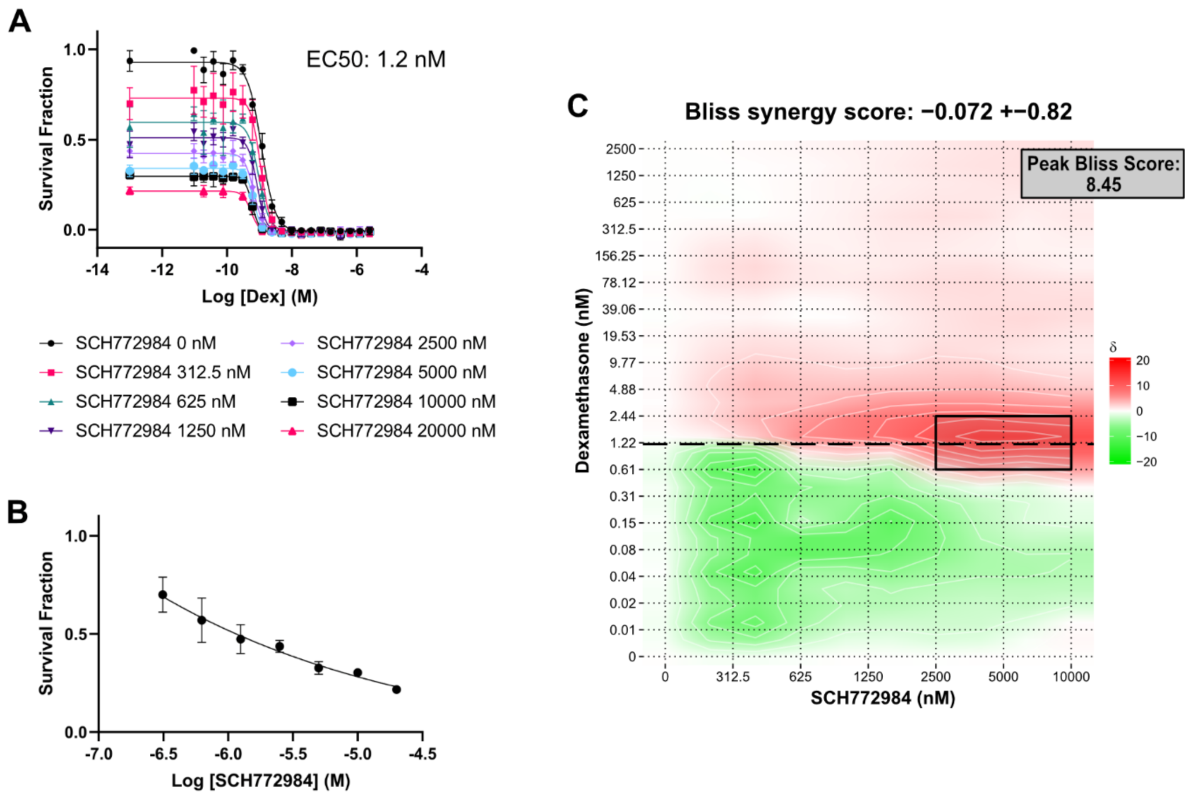

**Supplemental Figure 7. Synergy evaluation of Sup-B15 cells treated with dexamethasone and the ERK1/2 inhibitor SCH772984.** (A) Dose-response curves for dexamethasone (Dex) at different doses of SCH772984, with 3 replicates per concentration. Dexamethasone alone (SCH772984 0 nM) is shown in the top black line. (B) Dose-response curve for SCH772984 alone, 3 replicates per concentration. (C) Synergy plot of the combination treatment. Bliss score greater than 10 indicates synergy, 10 to -10 indicates additivity, and less than -10 indicates antagonism. Peak Bliss score from the black outlined area is given in the box below the plot. Long dashed black line indicates the approximate EC50 for dexamethasone.

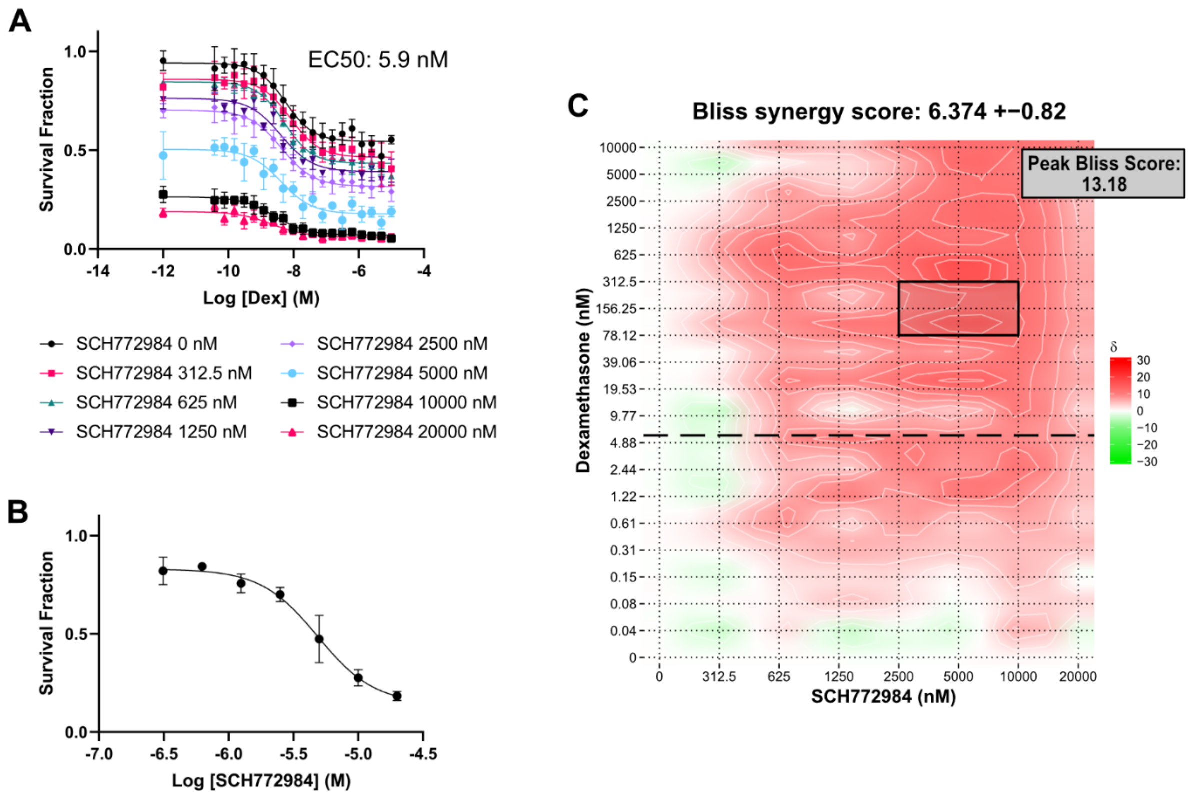

**Supplemental Figure 8. Synergy evaluation of RCH-ACV cells treated with dexamethasone and the ERK1/2 inhibitor SCH772984.** (A) Dose-response curves for dexamethasone (Dex) at different doses of SCH772984, with 3 replicates per concentration. Dexamethasone alone (SCH772984 0 nM) is shown in the top black line. (B) Dose-response curve for SCH772984 alone, 3 replicates per concentration. (C) Synergy plot of the combination treatment. Bliss score greater than 10 indicates synergy, 10 to -10 indicates additivity, and less than -10 indicates antagonism. Peak Bliss score from the black outlined area is given in the box below the plot. Long dashed black line indicates the approximate EC50 for dexamethasone, which is below the peak Bliss score area in these non-glucocorticoid sensitive cells.

### Dex+Idela RNAseq

Miles Pufall

4/3/2023

### Background

This is an R Markdown document. Markdown is a simple formatting syntax for authoring HTML, PDF, and MS Word documents. For more details on using R Markdown see http://rmarkdown.rstudio.com (http://rmarkdown.rstudio.com).

When you click the **Knit** button a document will be generated that includes both content as well as the output of any embedded R code chunks within the document. You can embed an R code chunk like this:

### Read in Screen Results

```
full_rhos <- readxl::read_excel("~/Library/CloudStorage/Dropbox/miles/screen/full_rhos_1
80815.xlsx")
sig_rhos <- dplyr::filter(full_rhos, Rho.P.value < 0.05)
full_gammas <- read_csv("~/Library/CloudStorage/Dropbox/miles/screen/full_gammas_180815.
csv")
sig_gammas <- dplyr::filter(full_gammas, Gamma.P.value < 0.01)

cagek_rhos <- readxl::read_excel("~/Library/CloudStorage/Dropbox/miles/screen/CAGEK_rhos
_1508.xlsx")
c_sig_rhos <- dplyr::filter(cagek_rhos, `Rho P value` < 0.05)
cagek_gammas <- readxl::read_excel("~/Library/CloudStorage/Dropbox/miles/screen/CAGEK_rh
os_1508.xlsx", sheet = 2)
c_sig_gammas <-dplyr::filter(cagek_gammas, `Gamma P value` < 0.01)
```

### Import RNA-seq data

#### Identify count tables

```
cond <- read_csv("cond_idel.csv") %>%
  mutate(sample = replace(sample, sample=="Hb3_quant", "hb3_quant")) %>%
  mutate(dex = as.factor(dex)) %>%
  mutate(idela = as.factor(idela)) %>%
  rename(names = sample)
```

```
## Rows: 18 Columns: 4
## ── Column specification ────────────────────────────────────────────────────────
## Delimiter: ","
## chr (2): sample, condition
## dbl (2): dex, idela
##
## ℹ Use `spec()` to retrieve the full column specification for this data.
## ℹ Specify the column types or set `show_col_types = FALSE` to quiet this message.
```

```
cond$files <- file.path("salm", "quants", cond$names, "quant.sf")
cond <- as_tibble(cond)

file.exists(cond$files)
```

```
##  [1] TRUE TRUE TRUE TRUE TRUE TRUE TRUE TRUE TRUE TRUE TRUE TRUE TRUE TRUE TRUE
## [16] TRUE TRUE TRUE
```

```
se <- tximeta(cond)
```

```
## importing quantifications
## reading in files with read_tsv
## 1 2 3 4 5 6 7 8 9 10 11 12 13 14 15 16 17 18
## found matching transcriptome:
## [ GENCODE - Homo sapiens - release 42 ]
## loading existing TxDb created: 2022-12-30 00:09:26
## loading existing transcript ranges created: 2022-12-30 00:09:27
## fetching genome info for GENCODE
```

```
dim(se)
```

```
## [1] 251550     18
```

### Import into DESeq

```
gse <- summarizeToGene(se)
```

```
## loading existing TxDb created: 2022-12-30 00:09:26
```

```
## obtaining transcript-to-gene mapping from database
```

```
## loading existing gene ranges created: 2022-12-30 00:10:30
```

```
## summarizing abundance
```

```
## summarizing counts
```

```
## summarizing length
```

```
dim(gse)
```

```
## [1] 62262    18
```

```
round( colSums(assay(gse)) / 1e6, 1 )
```

```
## veh1_quant veh2_quant veh3_quant  lb1_quant  lb2_quant  lb3_quant  hb1_quant
##       18.1       23.0       21.4       24.1       20.0       30.9       19.8
##  hb2_quant  hb3_quant ide1_quant ide2_quant ide3_quant  lo1_quant  lo2_quant
##       17.5       28.0       21.8       27.6       23.9       21.4       24.0
##  lo3_quant  hi1_quant  hi2_quant  hi3_quant
##       26.3       18.6       19.7       29.9
```

```
#Export count table
count_table <- round(assays(gse)$counts, 0) %>%
  as_tibble(rownames = "ensembl")
write_csv(count_table, "nalm6_idela_gene_count_table.csv")

dds <- DESeqDataSet(gse, ~dex + idela + dex:idela)
```

```
## using counts and average transcript lengths from tximeta
```

```
dds$group <- factor(paste0(dds$dex, dds$idela))
design(dds) <- ~ group
```

### Pre filter

```
dds <- dds[ rowSums(counts(dds)) > 36, ]
nrow(dds)
```

```
## [1] 21242
```

### Exploratory Data Analysis

#### Sample comparison heatmap

```
vsd <- vst(dds, blind = FALSE) #fast
```

```
## using 'avgTxLength' from assays(dds), correcting for library size
```

```
dds <- estimateSizeFactors(dds)
```

```
## using 'avgTxLength' from assays(dds), correcting for library size
```

```
sampleDists <- dist(t(assay(vsd)))

sampleDistMatrix <- as.matrix( sampleDists )
rownames(sampleDistMatrix) <- vsd$names
colnames(sampleDistMatrix) <- NULL
colors <- colorRampPalette( rev(brewer.pal(9, "Blues")) )(255)
pheatmap(sampleDistMatrix,
         clustering_distance_rows = sampleDists,
         clustering_distance_cols = sampleDists)
```

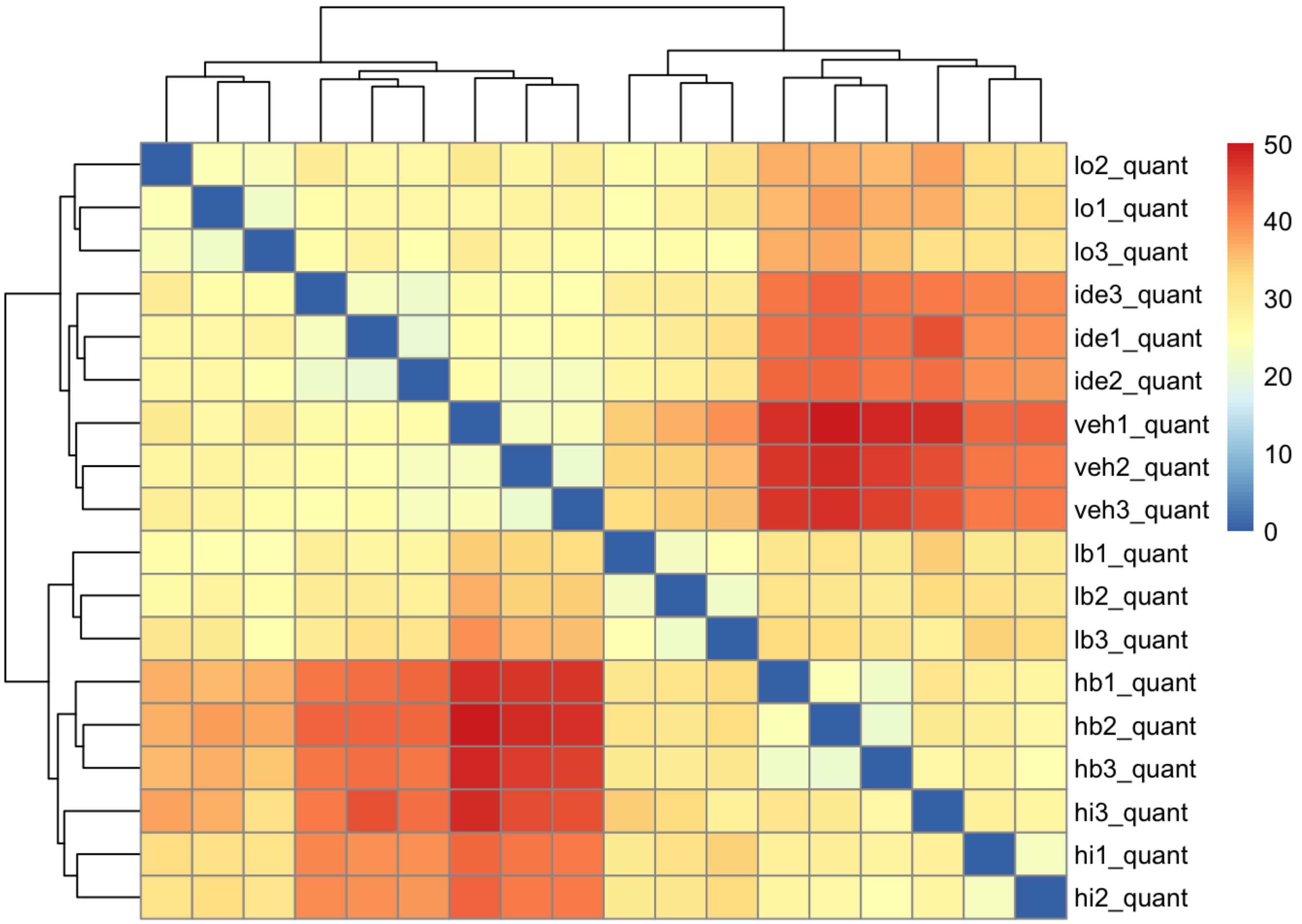

### Poisson-based heatmap

```
poisd <- PoissonDistance(t(counts(dds)))

samplePoisDistMatrix <- as.matrix( poisd$dd )
rownames(samplePoisDistMatrix) <- dds$names
colnames(samplePoisDistMatrix) <- NULL
pheatmap(samplePoisDistMatrix,
         clustering_distance_rows = poisd$dd,
         clustering_distance_cols = poisd$dd)
```

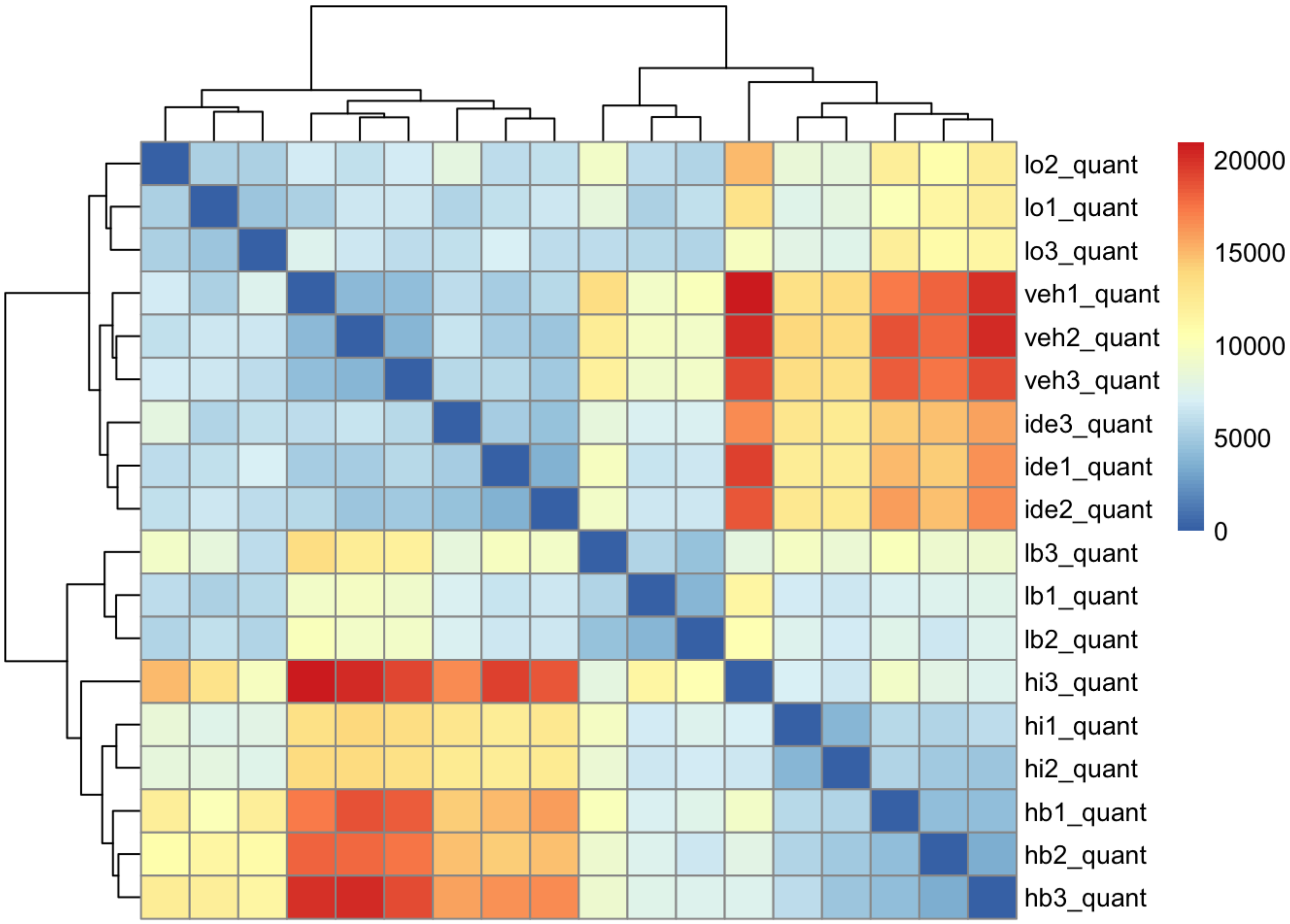

### PCA Plot

```
plotPCA(vsd, intgroup = c("dex", "idela")) +
  theme_bw() +
  theme(axis.title = element_text(face = "bold"))
```

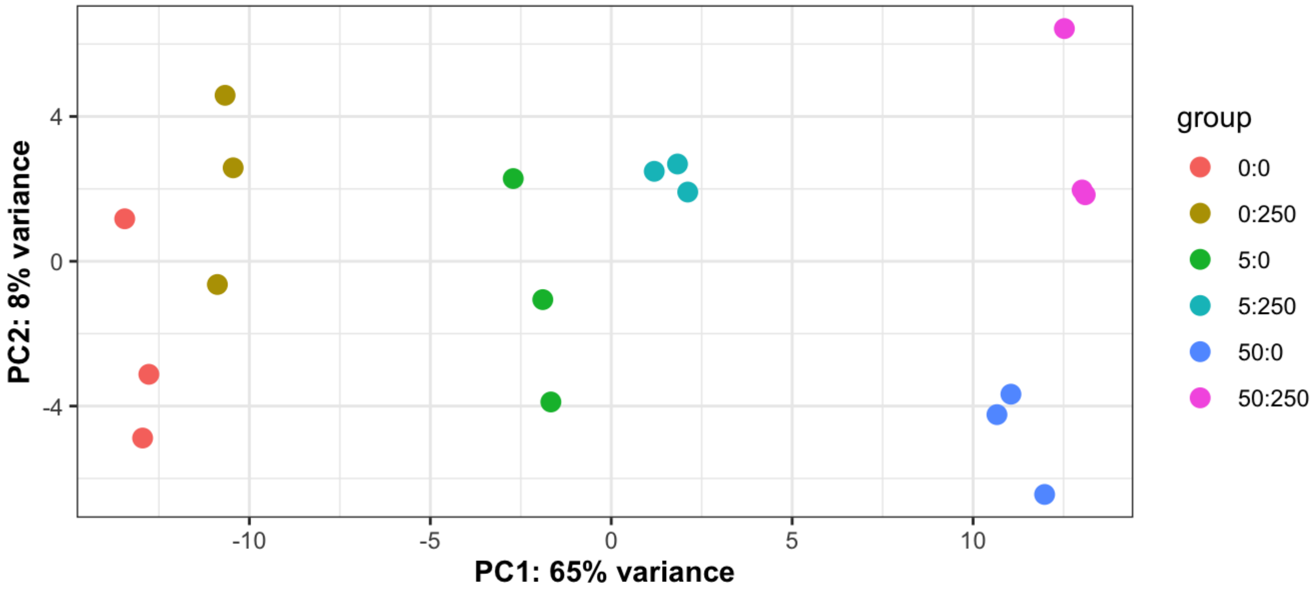

```
ggsave("nalm6_pca.pdf")
```

```
## Saving 7 x 5 in image
```

### MDS Plots

```
mds <- as.data.frame(colData(vsd))  %>%
  cbind(cmdscale(sampleDistMatrix))
ggplot(mds, aes(x = `1`, y = `2`, color = dex, shape = idela)) +
  geom_point(size = 3) +
  coord_fixed() +
  theme_bw()
```

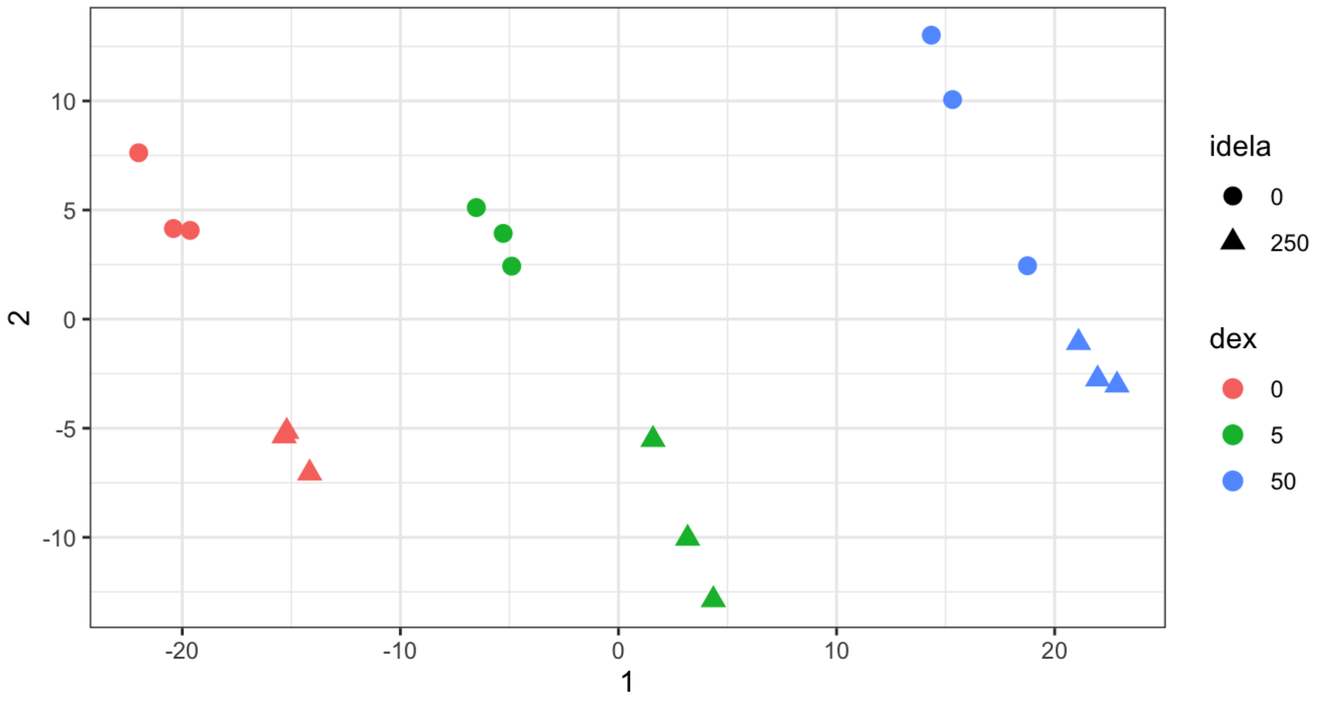

```
ggsave("mds_dex_idela.pdf", width = 4, height = 4)

mdsPois <- as.data.frame(colData(dds)) %>%
  cbind(cmdscale(samplePoisDistMatrix))
ggplot(mdsPois, aes(x = `1`, y = `2`, color = dex, shape = idela)) +
  geom_point(size = 3) +
  coord_fixed() +
  theme_bw()
```

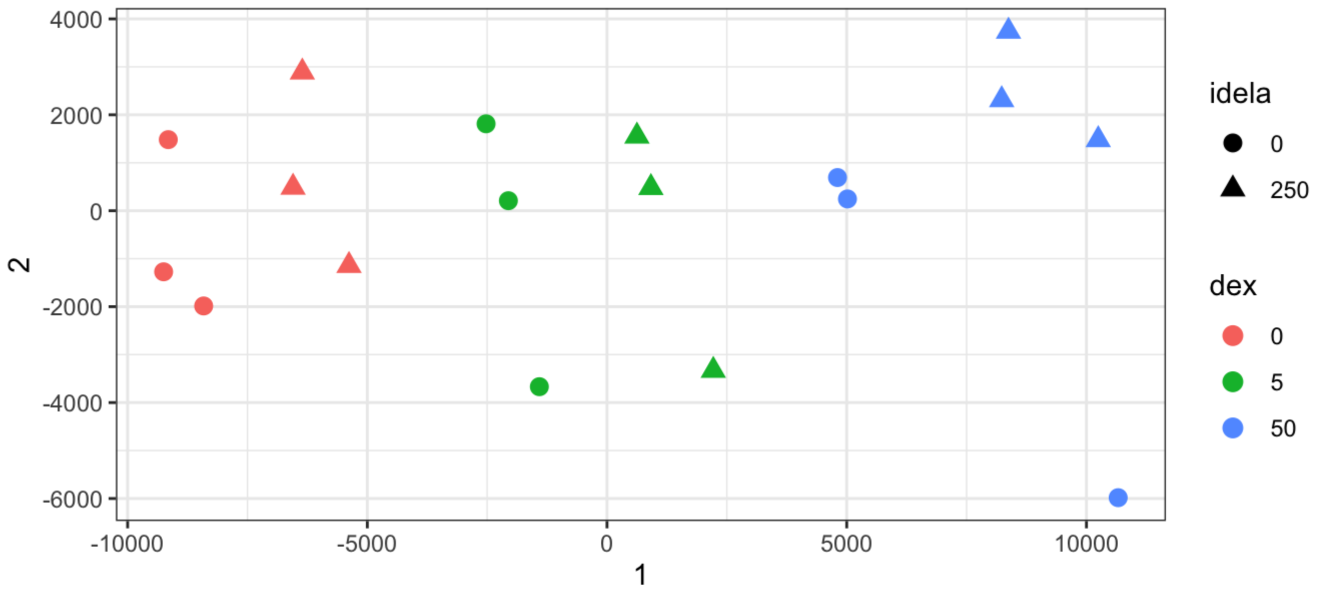

### Calculate differential expression

```
dds <- DESeq(dds)
```

```
## using pre-existing normalization factors
```

```
## estimating dispersions
```

```
## gene-wise dispersion estimates
```

```
## mean-dispersion relationship
```

```
## final dispersion estimates
```

```
## fitting model and testing
```

```
resultsNames(dds)
```

```
## [1] "Intercept"         "group_0250_vs_00"  "group_50_vs_00"
## [4] "group_500_vs_00"   "group_50250_vs_00" "group_5250_vs_00"
```

```
results <- results(dds)
```

### Make results tables for treatments versus control

```
res.hidex <- results(dds, name = "group_500_vs_00")
head(res.hidex[ order(res.hidex$padj, decreasing = FALSE), ])
```

```
## log2 fold change (MLE): group 500 vs 00
## Wald test p-value: group 500 vs 00
## DataFrame with 6 rows and 6 columns
##                      baseMean log2FoldChange     lfcSE      stat       pvalue
##                     <numeric>      <numeric> <numeric> <numeric>    <numeric>
## ENSG00000143119.14  5893.710        2.30880 0.0567756   40.6654  0.00000e+00
## ENSG00000096060.15  7856.732        1.97255 0.0550563   35.8279 4.06130e-281
## ENSG00000265972.6   4263.711        3.07612 0.0911625   33.7433 1.34239e-249
## ENSG00000248302.3    921.545        4.34853 0.1312012   33.1440 6.91000e-241
## ENSG00000159200.18  3711.420        2.39813 0.0734715   32.6403 1.09914e-233
## ENSG00000256235.3   1054.201        3.29998 0.1032100   31.9734 2.55308e-224
##                            padj
##                       <numeric>
## ENSG00000143119.14  0.00000e+00
## ENSG00000096060.15 4.22700e-277
## ENSG00000265972.6  9.31442e-246
## ENSG00000248302.3  3.59597e-237
## ENSG00000159200.18 4.57595e-230
## ENSG00000256235.3  8.85748e-221
```

```
sum(res.hidex$padj < 0.01, na.rm=TRUE)
```

```
## [1] 3778
```

```
summary(res.hidex, alpha = 0.01)
```

```
##
## out of 21242 with nonzero total read count
## adjusted p-value < 0.01
## LFC > 0 (up)       : 1696, 8%
## LFC < 0 (down)     : 2082, 9.8%
## outliers [1]       : 14, 0.066%
## low counts [2]     : 412, 1.9%
## (mean count < 2)
## [1] see 'cooksCutoff' argument of ?results
## [2] see 'independentFiltering' argument of ?results
```

```
res.lodex <- results(dds, name = "group_50_vs_00")
head(res.lodex[ order(res.lodex$padj, decreasing = FALSE), ])
```

```
## log2 fold change (MLE): group 50 vs 00
## Wald test p-value: group 50 vs 00
## DataFrame with 6 rows and 6 columns
##                      baseMean log2FoldChange     lfcSE      stat       pvalue
##                     <numeric>      <numeric> <numeric> <numeric>    <numeric>
## ENSG00000256235.3   1054.201        2.56699 0.1042310   24.6279 6.34421e-134
## ENSG00000159200.18  3711.420        1.71182 0.0739485   23.1488 1.49357e-118
## ENSG00000248302.3    921.545        3.08684 0.1333862   23.1421 1.74456e-118
## ENSG00000109501.15   804.747        3.61453 0.1660013   21.7741 4.08126e-105
## ENSG00000174944.9   3855.746        5.78615 0.2802596   20.6457  1.06759e-94
## ENSG00000143119.14  5893.710        1.14805 0.0573598   20.0149  4.08097e-89
##                            padj
##                       <numeric>
## ENSG00000256235.3  9.29236e-130
## ENSG00000159200.18 8.51751e-115
## ENSG00000248302.3  8.51751e-115
## ENSG00000109501.15 1.49446e-101
## ENSG00000174944.9   3.12741e-91
## ENSG00000143119.14  9.96233e-86
```

```
sum(res.lodex$padj < 0.01, na.rm=TRUE)
```

```
## [1] 649
```

```
summary(res.lodex, alpha = 0.01)
```

```
## 
## out of 21242 with nonzero total read count
## adjusted p-value < 0.01
## LFC > 0 (up)       : 407, 1.9%
## LFC < 0 (down)     : 242, 1.1%
## outliers [1]       : 14, 0.066%
## low counts [2]     : 6581, 31%
## (mean count < 14)
## [1] see 'cooksCutoff' argument of ?results
## [2] see 'independentFiltering' argument of ?results
```

```
res.idel <-  results(dds, name = "group_0250_vs_00")
head(res.idel[ order(res.idel$padj, decreasing = FALSE), ])
```

```
## log2 fold change (MLE): group 0250 vs 00
## Wald test p-value: group 0250 vs 00
## DataFrame with 6 rows and 6 columns
##                      baseMean log2FoldChange     lfcSE      stat       pvalue
##                     <numeric>      <numeric> <numeric> <numeric>    <numeric>
## ENSG00000134853.12 21714.814       1.334182 0.0578140   23.0771 7.85603e-118
## ENSG00000165507.9    621.084       1.959074 0.1356858   14.4383  2.97023e-47
## ENSG00000170365.10  8105.290       0.615688 0.0472780   13.0227  9.08835e-39
## ENSG00000086730.17   839.007      -1.132965 0.0956698  -11.8425  2.35467e-32
## ENSG00000087495.17   965.110       1.176041 0.0999210   11.7697  5.59153e-32
## ENSG00000107537.14  1409.967       1.199796 0.1132524   10.5940  3.17724e-26
##                             padj
##                        <numeric>
## ENSG00000134853.12 1.15067e-113
## ENSG00000165507.9   2.17525e-43
## ENSG00000170365.10  4.43723e-35
## ENSG00000086730.17  8.62223e-29
## ENSG00000087495.17  1.63798e-28
## ENSG00000107537.14  7.75617e-23
```

```
sum(res.idel$padj < 0.01, na.rm=TRUE)
```

```
## [1] 418
```

```
summary(res.idel, alpha = 0.01)
```

```
##
## out of 21242 with nonzero total read count
## adjusted p-value < 0.01
## LFC > 0 (up)       : 208, 0.98%
## LFC < 0 (down)     : 210, 0.99%
## outliers [1]       : 14, 0.066%
## low counts [2]     : 6581, 31%
## (mean count < 14)
## [1] see 'cooksCutoff' argument of ?results
## [2] see 'independentFiltering' argument of ?results
```

```
res.lob <- results(dds, name = "group_5250_vs_00")
head(res.lob[ order(res.lob$padj, decreasing = FALSE), ])
```

```
## log2 fold change (MLE): group 5250 vs 00
## Wald test p-value: group 5250 vs 00
## DataFrame with 6 rows and 6 columns
##                      baseMean log2FoldChange     lfcSE      stat       pvalue
##                     <numeric>      <numeric> <numeric> <numeric>    <numeric>
## ENSG00000159200.18  3711.420        2.31732 0.0733828   31.5785 7.27599e-219
## ENSG00000256235.3   1054.201        2.95346 0.1033185   28.5860 1.00331e-179
## ENSG00000134853.12 21714.814        1.59611 0.0577802   27.6239 5.74750e-168
## ENSG00000248302.3    921.545        3.62203 0.1320445   27.4304 1.19207e-165
## ENSG00000170365.10  8105.290        1.16277 0.0469627   24.7595 2.44999e-135
## ENSG00000174944.9   3855.746        6.46468 0.2801177   23.0784 7.62301e-118
##                            padj
##                       <numeric>
## ENSG00000159200.18 1.21531e-214
## ENSG00000256235.3  8.37914e-176
## ENSG00000134853.12 3.20002e-164
## ENSG00000248302.3  4.97777e-162
## ENSG00000170365.10 8.18444e-132
## ENSG00000174944.9  2.12212e-114
```

```
sum(res.lob$padj < 0.01, na.rm=TRUE)
```

```
## [1] 2398
```

```
summary(res.lob, alpha = 0.01)
```

```
##
## out of 21242 with nonzero total read count
## adjusted p-value < 0.01
## LFC > 0 (up)       : 1053, 5%
## LFC < 0 (down)     : 1345, 6.3%
## outliers [1]       : 14, 0.066%
## low counts [2]     : 4525, 21%
## (mean count < 7)
## [1] see 'cooksCutoff' argument of ?results
## [2] see 'independentFiltering' argument of ?results
```

```
res.hib <- results(dds, name = "group_50250_vs_00")
head(res.hib[ order(res.hib$padj, decreasing = FALSE), ])
```

```
## log2 fold change (MLE): group 50250 vs 00
## Wald test p-value: group 50250 vs 00
## DataFrame with 6 rows and 6 columns
##                     baseMean log2FoldChange     lfcSE      stat       pvalue
##                    <numeric>      <numeric> <numeric> <numeric>    <numeric>
## ENSG00000143119.14  5893.710        2.30042 0.0567700   40.5218  0.00000e+00
## ENSG00000159200.18  3711.420        2.98128 0.0731251   40.7696  0.00000e+00
## ENSG00000248302.3    921.545        4.90967 0.1306716   37.5726  0.00000e+00
## ENSG00000265972.6   4263.711        3.09309 0.0911349   33.9397 1.73045e-252
## ENSG00000101445.10  3816.773        2.58802 0.0783920   33.0138 5.15278e-239
## ENSG00000256235.3   1054.201        3.33627 0.1031139   32.3552 1.17282e-229
##                            padj
##                       <numeric>
## ENSG00000143119.14  0.00000e+00
## ENSG00000159200.18  0.00000e+00
## ENSG00000248302.3   0.00000e+00
## ENSG00000265972.6  8.82705e-249
## ENSG00000101445.10 2.10275e-235
## ENSG00000256235.3  3.98836e-226
```

```
sum(res.hib$padj < 0.01, na.rm=TRUE)
```

```
## [1] 4965
```

```
summary(res.hib, alpha = 0.01)
```

```
##
## out of 21242 with nonzero total read count
## adjusted p-value < 0.01
## LFC > 0 (up)       : 2164, 10%
## LFC < 0 (down)     : 2801, 13%
## outliers [1]       : 14, 0.066%
## low counts [2]     : 824, 3.9%
## (mean count < 3)
## [1] see 'cooksCutoff' argument of ?results
## [2] see 'independentFiltering' argument of ?results
```

#### Annotate results tables

```
results_table <- function(res_name, deseq_obj, new_name) {
  df <- results(deseq_obj, name = res_name)
  df <- as.data.frame(df)
  df <- df[,c(1:3, 5:6)]
  colnames(df) <- c("base_mean", paste0(new_name, "_log2FC"), paste0(new_name,"_lfcse"),
paste0(new_name, "_pval"), paste0(new_name, "_adjp"))
  new_name <- df
  return(new_name)
}
lfc_table <- function(res_name, deseq_obj, new_name) {
  df <- lfcShrink(deseq_obj, coef = res_name)
  df <- as.data.frame(df)
  colnames(df) <- c("base_mean", paste0(new_name, "_log2FC"), paste0(new_name,"_lfcse"),
paste0(new_name, "_pval"), paste0(new_name, "_adjp"))
  new_name <- df
  return(new_name)
}

idel <- results_table("group_0250_vs_00", dds, "idel")
lodex <- results_table("group_50_vs_00", dds, "lodex")
hidex <- results_table("group_500_vs_00", dds, "hidex")
loboth <- results_table("group_5250_vs_00", dds, "loboth")
hiboth <- results_table("group_50250_vs_00", dds, "hiboth")

#idel <- lfc_table("group_0250_vs_00", dds, "idel")
#lodex <- lfc_table("group_50_vs_00", dds, "lodex")
#hidex <- lfc_table("group_500_vs_00", dds, "hidex")
#loboth <- lfc_table("group_5250_vs_00", dds, "loboth")
#hiboth <- lfc_table("group_50250_vs_00", dds, "hiboth")

sum_table <-
  cbind(idel, lodex[, c(2:5)]) %>%
  cbind(.,hidex[, c(2:5)]) %>%
  cbind(.,loboth[, c(2:5)]) %>%
  cbind(.,hiboth[, c(2:5)])

add_geneids <- function(genelist) {
  genelist$symbol <- mapIds(org.Hs.eg.db, keys=str_sub(row.names(genelist), 1, 15), colu
mn="SYMBOL", keytype="ENSEMBL", multiVals="first")
  genelist$entrez <- mapIds(org.Hs.eg.db, keys=str_sub(row.names(genelist), 1, 15), colu
mn="ENTREZID", keytype="ENSEMBL", multiVals="first")
  genelist$genename <- mapIds(org.Hs.eg.db, str_sub(row.names(genelist), 1, 15), column
="GENENAME", keytype="ENSEMBL", multiVals="first")
  #genelist <- genelist %>% drop_na(log2FoldChange)
  return(genelist)
}
sum_table <- add_geneids(sum_table)
```

```
## 'select()' returned 1:many mapping between keys and columns
## 'select()' returned 1:many mapping between keys and columns
## 'select()' returned 1:many mapping between keys and columns
```

```
sum_tbl_2 <- sum_table %>%
  dplyr::select(0,(length(sum_table)-2):length(sum_table), everything()) %>%
  rownames_to_column(var = "ensembl") %>%
  arrange(symbol, ensembl) %>%
  filter(!duplicated(symbol)) %>%
  as_tibble()

write_csv(sum_tbl_2, "idela_genex_220609.csv")

sum_tbl_2
```

```
## # A tibble: 15,705 × 25
##    ensembl     symbol entrez genename base_mean idel_log2FC idel_lfcse idel_pval
##    <chr>       <chr>  <chr>  <chr>        <dbl>       <dbl>      <dbl>     <dbl>
##  1 ENSG000001… A1BG   1      alpha-1…    270.      -0.165       0.310     0.596
##  2 ENSG000002… A1BG-… 503538 A1BG an…    146.       0.255       0.195     0.191
##  3 ENSG000001… A2M    2      alpha-2…     37.9     -1.35        0.597     0.0238
##  4 ENSG000002… A2M-A… 144571 A2M ant…      5.48    -0.174       0.722     0.809
##  5 ENSG000000… AAAS   8086   aladin …    584.      -0.0217      0.0893    0.808
##  6 ENSG000000… AACS   65985  acetoac…    328.       0.100       0.141     0.479
##  7 ENSG000001… AAGAB  79719  alpha a…   1325.      -0.0426      0.131     0.745
##  8 ENSG000001… AAK1   22848  AP2 ass…   1801.      -0.104       0.122     0.397
##  9 ENSG000000… AAMDC  28971  adipoge…    109.      -0.00438     0.199     0.982
## 10 ENSG000001… AAMP   14     angio a…    811.      -0.0667      0.113     0.554
## # ℹ 15,695 more rows
## # ℹ 17 more variables: idel_adjp <dbl>, lodex_log2FC <dbl>, lodex_lfcse <dbl>,
## #   lodex_pval <dbl>, lodex_adjp <dbl>, hidex_log2FC <dbl>, hidex_lfcse <dbl>,
## #   hidex_pval <dbl>, hidex_adjp <dbl>, loboth_log2FC <dbl>,
## #   loboth_lfcse <dbl>, loboth_pval <dbl>, loboth_adjp <dbl>,
## #   hiboth_log2FC <dbl>, hiboth_lfcse <dbl>, hiboth_pval <dbl>,
## #   hiboth_adjp <dbl>
```

### Make a longer summary table

```
sum_lng <- sum_tbl_2 %>%
  pivot_longer(cols = !c(1:5), names_to = c("treat", "stat"), names_sep = "_", values_to
= "value") %>%
  pivot_wider(names_from = "stat", values_from = "value") %>%
  replace_na(list(pval = 1, adjp = 1)) %>%
  mutate(treat = factor(treat, c("idel", "lodex", "loboth", "hidex", "hiboth")))

sum_lng
```

```
## # A tibble: 78,525 × 10
##    ensembl    symbol entrez genename base_mean treat  log2FC lfcse   pval   adjp
##    <chr>      <chr>  <chr>  <chr>        <dbl> <fct>   <dbl> <dbl>  <dbl>  <dbl>
##  1 ENSG00000… A1BG   1      alpha-1…      270. idel  -0.165  0.310 0.596  0.883
##  2 ENSG00000… A1BG   1      alpha-1…      270. lodex  0.223  0.310 0.472  0.799
##  3 ENSG00000… A1BG   1      alpha-1…      270. hidex  0.0919 0.310 0.767  0.888
##  4 ENSG00000… A1BG   1      alpha-1…      270. lobo…  0.0653 0.309 0.833  0.921
##  5 ENSG00000… A1BG   1      alpha-1…      270. hibo…  0.685  0.309 0.0265 0.0764
##  6 ENSG00000… A1BG-… 503538 A1BG an…      146. idel   0.255  0.195 0.191  0.620
##  7 ENSG00000… A1BG-… 503538 A1BG an…      146. lodex  0.121  0.194 0.535  0.833
##  8 ENSG00000… A1BG-… 503538 A1BG an…      146. hidex  0.478  0.193 0.0132 0.0532
##  9 ENSG00000… A1BG-… 503538 A1BG an…      146. lobo…  0.397  0.192 0.0387 0.131
## 10 ENSG00000… A1BG-… 503538 A1BG an…      146. hibo…  0.392  0.194 0.0437 0.115
## # ℹ 78,515 more rows
```

```
sum_rnaseq_tbl <- sum_lng %>%
  group_by(treat) %>%
  summarise(Up = sum(adjp <= 0.01 & log2FC > 0), Down = sum(adjp <= 0.01 & log2FC < 0))
%>%
  mutate(Total = Up + Down)

sum_lng %>%
  group_by(treat) %>%
  summarise(Up = sum(adjp <= 0.01 & log2FC > 0), Down = sum(adjp <= 0.01 & log2FC < 0))
%>%
  pivot_longer(cols = c("Up", "Down"), names_to = "Regulation", values_to = "Number") %
>%
  ggplot(aes(treat, Number, fill = Regulation)) +
  geom_col(width = 0.8, position=position_dodge(0.9)) +
  scale_fill_manual(values=c('blue','red')) +
  scale_x_discrete(breaks=c("idel", "lodex", "loboth", "hidex", "hiboth"), labels=c("Ide
la", "5nM Dex", "5 nM Dex +\nIdela", "50 nM Dex", "50 nM Dex +\nIdela")) +
  theme_bw() +
  ylab("Number of Genes Regulated") +
  theme(axis.text.x = element_text(angle = 45, hjust = 1), axis.title.x=element_blank(),
axis.title.y = element_text(face = "bold"), legend.position = c(0.2, 0.8))
```

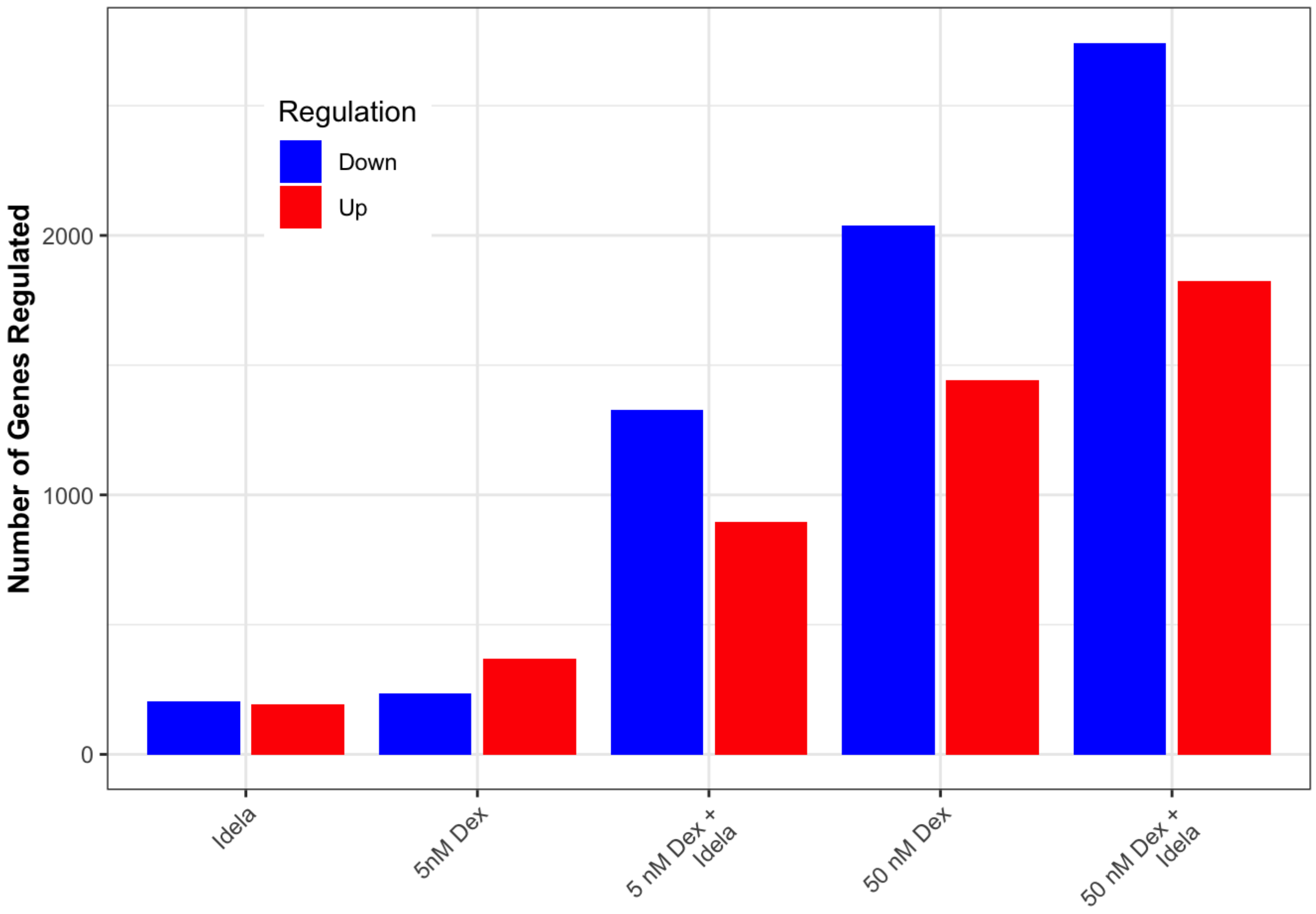

```
ggsave("nalm_idela_up_down_summary.pdf", width = 4, height = 4)
```

### Description of dex+idela vs dex alone

```
lo_and_both_reg <- dplyr::filter(sum_tbl_2, lodex_adjp <= 0.01 | loboth_adjp <= 0.01 | idel_adjp <= 0.01)
cc = character(nrow(lo_and_both_reg)) ## will hold the color designations
cc[lo_and_both_reg$loboth_adjp <= 0.01] = "white"
cc[lo_and_both_reg$idel_adjp <= 0.01 & lo_and_both_reg$lodex_adjp <= 0.01 & lo_and_both_reg$loboth_adjp <= 0.01] = "black"
cc[lo_and_both_reg$idel_adjp <= 0.01 & !lo_and_both_reg$loboth_adjp <= 0.01 & !lo_and_both_reg$lodex_adjp <= 0.01  ] = "red"
cc[lo_and_both_reg$lodex_adjp <= 0.01 & !lo_and_both_reg$idel_adjp <= 0.01 & !lo_and_both_reg$loboth_adjp <= 0.01 ] = "green"
cc[lo_and_both_reg$idel_adjp <= 0.01 & lo_and_both_reg$lodex_adjp <= 0.01 & !lo_and_both_reg$loboth_adjp <= 0.01] = "yellow"
lo_and_both_reg <- cbind(lo_and_both_reg,cc)

lm_lo_and_both <- lm(lo_and_both_reg$lodex_log2FC ~ lo_and_both_reg$loboth_log2FC)
hi_and_both_reg <- dplyr::filter(sum_table, hidex_adjp <= 0.01 | hiboth_adjp <= 0.01)
lm_hi_and_both <- lm(hi_and_both_reg$hidex_log2FC ~ hi_and_both_reg$hiboth_log2FC)

par(mfrow = c(c(1,2)))
plot(lo_and_both_reg$lodex_log2FC, lo_and_both_reg$loboth_log2FC,
     col = ifelse(lo_and_both_reg$lodex_adjp <= 0.01, "red", "black"), xlim = c(-10,10), ylim = c(-10,10), pch = 20,
     xlab = "log2 Fold Change (5nM Dex)", ylab = "log2 Fold Change (5nM Dex + 250nM Idela)", main = "5 nM Dex + Idelalisib")
abline(0,1)
plot(hi_and_both_reg$hidex_log2FC, hi_and_both_reg$hiboth_log2FC,
     col = ifelse(hi_and_both_reg$hidex_adjp <=0.01, "red", "black"), xlim = c(-10,10), ylim = c(-10,10), pch = 20,
     xlab = "log2 Fold Change (50nM Dex)", ylab = "log2 Fold Change (50nM Dex + 250nM Idela)", main = "50 nM Dex + Idelalisib")
abline(0,1)
```

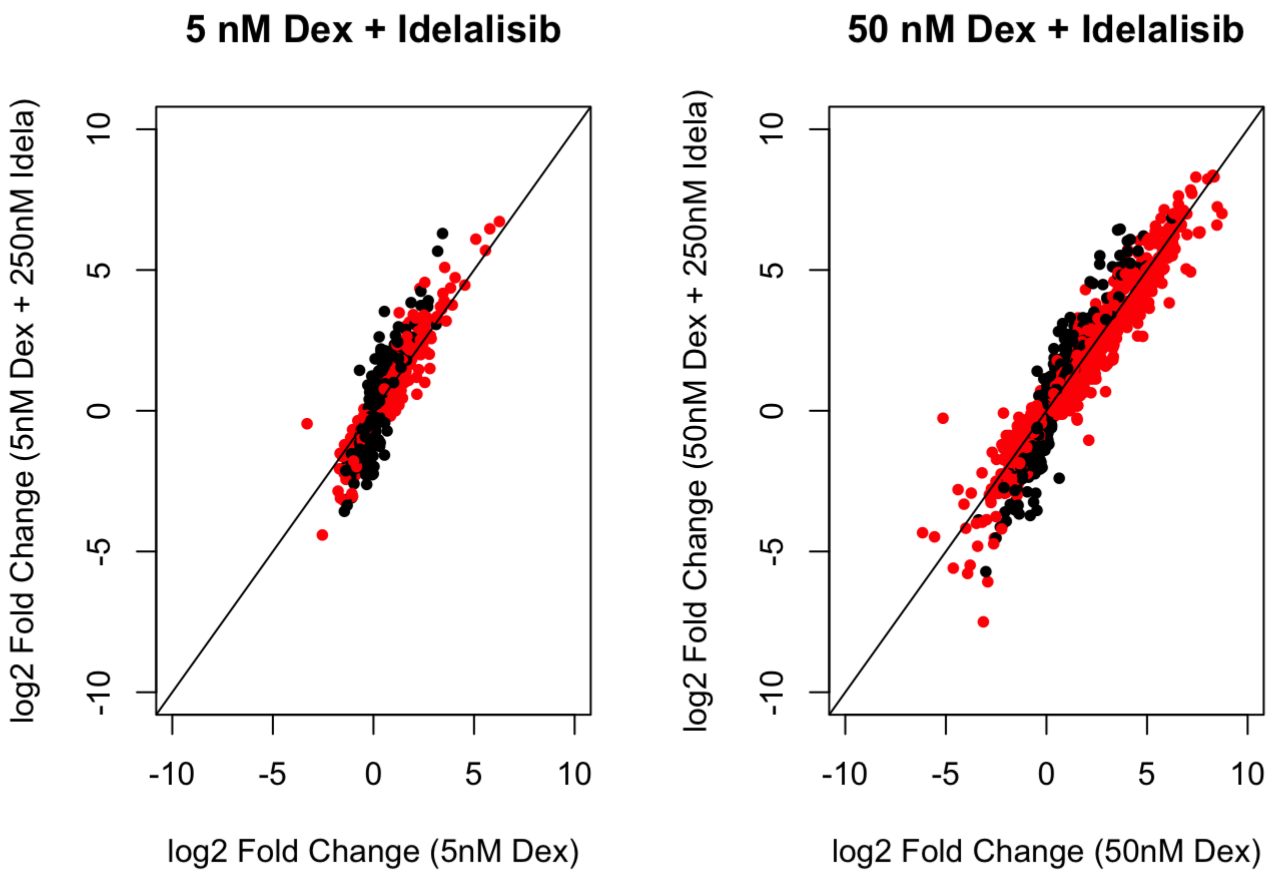
5 nM Dex + Idelalisib
log2 Fold Change (5nM Dex + 250nM Idela)
log2 Fold Change (5nM Dex)
-10
-5
0
5
10
50 nM Dex + Idelalisib
log2 Fold Change (50nM Dex + 250nM Idela)
log2 Fold Change (50nM Dex)

```r
lodex_reg_filt <- dplyr::filter(sum_table, lodex_adjp <= 0.01)
p1 <- ggscatter(lodex_reg_filt, x = "lodex_log2FC", y = "loboth_log2FC",
          add = "reg.line", conf.int = TRUE,
          cor.coef = TRUE, cor.method = "pearson",
          xlab = "log2FC, 5nM Dex", ylab = "log2FC, Dex + Idela") +
  xlim(-6,8) + ylim(-6,8) +
  stat_cor(label.x.npc = "left", label.y.npc = "top") +
  stat_regline_equation(label.y = 8, aes(label = after_stat(eq.label))) +
  geom_abline(intercept = 0, slope = 1, colour = "red") +
  theme_bw()

hidex_reg_filt <- dplyr::filter(sum_tbl_2, hidex_adjp <= 0.01 & abs(hiboth_log2FC) < 15
& abs(hidex_log2FC) < 15)
#hidex_reg_filt <- dplyr::filter(sum_table, hidex_adjp <= 0.01)

p2 <- ggscatter(hidex_reg_filt, x = "hidex_log2FC", y = "hiboth_log2FC",
          add = "reg.line", conf.int = TRUE,
          cor.coef = TRUE, cor.method = "pearson",
          xlab = "log2FC, 50nM Dex", ylab = "log2FC, Dex + Idela") +
  xlim(-6,8) + ylim(-6,8) +
  stat_cor(label.x.npc = "left", label.y.npc = "top") +
  stat_regline_equation(label.y = 8, aes(label = after_stat(eq.label))) +
  geom_abline(intercept = 0, slope = 1, colour = "red") +
  theme_bw()

g <- grid.arrange(p1, p2, nrow = 1)
```

```
## Warning: Removed 2 rows containing non-finite values (`stat_smooth()`).
```

```
## Warning: Removed 2 rows containing non-finite values (`stat_cor()`).
## Removed 2 rows containing non-finite values (`stat_cor()`).
```

```
## Warning: Removed 2 rows containing non-finite values
## (`stat_regline_equation()`).
```

```
## Warning: Removed 2 rows containing missing values (`geom_point()`).
```

```
## Warning: Removed 1 rows containing missing values (`geom_smooth()`).
```

```
## Warning: Removed 6 rows containing non-finite values (`stat_smooth()`).
```

```
## Warning: Removed 6 rows containing non-finite values (`stat_cor()`).
## Removed 6 rows containing non-finite values (`stat_cor()`).
```

```
## Warning: Removed 6 rows containing non-finite values
## (`stat_regline_equation()`).
```

```
## Warning: Removed 6 rows containing missing values (`geom_point()`).
```

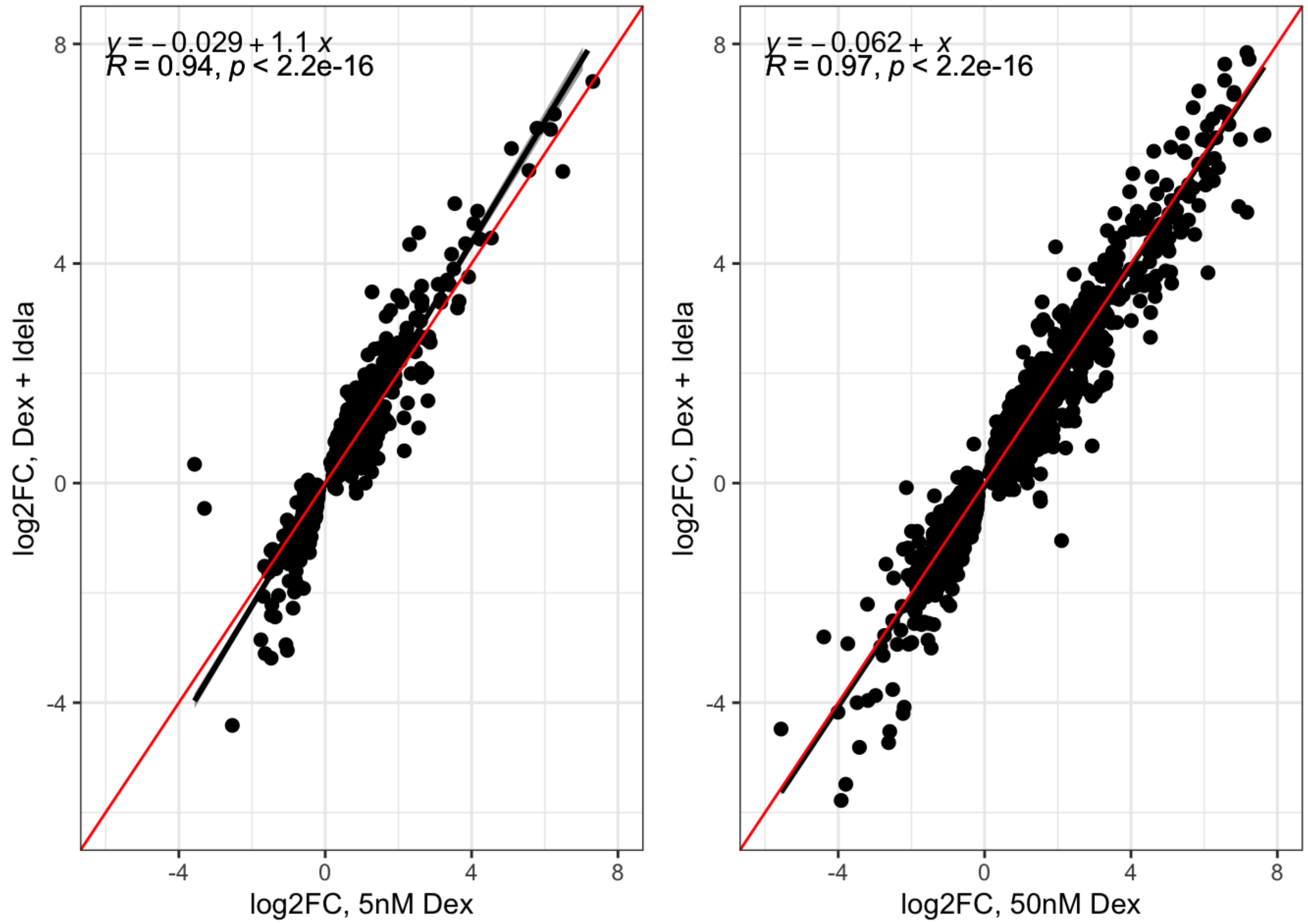

```
ggsave(filename = "nalm6_dex_idel_5_50_scatter.pdf", height = 4, width = 8, g)
```

### Do a correlation plot to determine the general behavior of dex + idela

```
lodex_reg_filt <- dplyr::filter(sum_tbl_2, lodex_adjp <= 0.01 & abs(lodex_log2FC) < 10)

# ttest for enhanced upregulation by idela
lodex_reg_filt %>%
  filter(loboth_log2FC > 0) %>%
  t.test(.$loboth_log2FC, .$lodex_log2FC, data= .)
```

```
##
##  Welch Two Sample t-test
##
## data:  .$loboth_log2FC and .$lodex_log2FC
## t = 2.1342, df = 710.77, p-value = 0.03316
## alternative hypothesis: true difference in means is not equal to 0
## 95 percent confidence interval:
##  0.01263657 0.30292009
## sample estimates:
## mean of x mean of y
##  1.204455  1.046677
```

```
test_up <- lodex_reg_filt %>%
  filter(loboth_log2FC > 0)

b1 <- lodex_reg_filt %>%
  dplyr::select(lodex_log2FC, loboth_log2FC) %>%
  filter(loboth_log2FC > 0) %>%
  pivot_longer(cols = c("lodex_log2FC", "loboth_log2FC"), names_to = "treat", values_to
= "log2FC") %>%
  mutate(treat = factor(treat, levels = c("lodex_log2FC", "loboth_log2FC"))) %>%
  ggplot(aes(treat, log2FC)) +
  ggtitle("Low dex, upregulated genes") + ylab("log2FC") + xlab("") +
  geom_boxplot() +
  theme(axis.text.x = element_text(angle = 45, hjust = 1)) +
  theme_bw()

b_up <- lodex_reg_filt%>%
  dplyr::select(lodex_log2FC, loboth_log2FC) %>%
  filter(loboth_log2FC > 0 | lodex_log2FC > 0) %>%
  pivot_longer(cols = c("lodex_log2FC", "loboth_log2FC"), names_to = "treat", values_to
= "log2FC") %>%
  mutate(treat = factor(treat, levels = c("lodex_log2FC", "loboth_log2FC"))) %>%
  ggplot(aes(treat, log2FC)) +
  ylab("log2FC") + xlab("") +
  geom_boxplot() +
  ylim(-6,8) +
  theme_bw() +
  theme(axis.text.x = element_text(angle = 45, hjust = 1))

b_down <- lodex_reg_filt%>%
  dplyr::select(lodex_log2FC, loboth_log2FC) %>%
  filter(loboth_log2FC < 0 | lodex_log2FC < 0) %>%
  pivot_longer(cols = c("lodex_log2FC", "loboth_log2FC"), names_to = "treat", values_to
= "log2FC") %>%
  mutate(treat = factor(treat, levels = c("lodex_log2FC", "loboth_log2FC"))) %>%
  ggplot(aes(treat, log2FC)) +
  ylab("log2FC") + xlab("") +
  geom_boxplot() +
  ylim(-6,8) +
  theme_bw() +
  theme(axis.text.x = element_text(angle = 45, hjust = 1))

box_lowboth <- grid.arrange(b_up, b_down, nrow = 1)
```

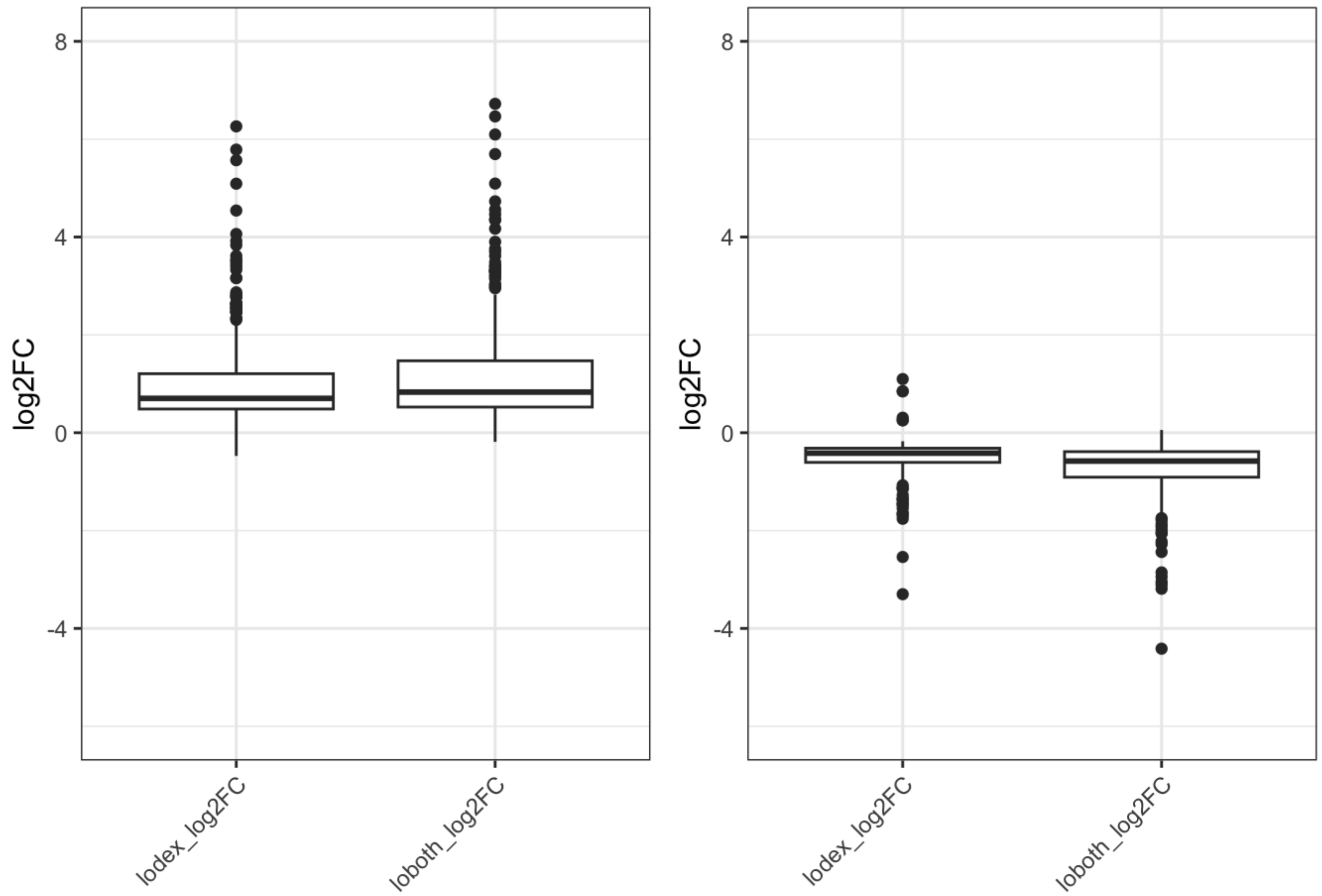

```
ggsave(filename = "boxplot_dex_idel_5.pdf", height = 4, width = 3, box_lowboth)

t.test(test_up$loboth_log2FC, test_up$lodex_log2FC, paired = TRUE)
```

```
##
##  Paired t-test
##
## data:  test_up$loboth_log2FC and test_up$lodex_log2FC
## t = 6.5445, df = 363, p-value = 2.036e-10
## alternative hypothesis: true mean difference is not equal to 0
## 95 percent confidence interval:
##  0.1103686 0.2051880
## sample estimates:
## mean difference
##       0.1577783
```

```
test_down <- lodex_reg_filt %>%
  filter(loboth_log2FC < 0)

t.test(test_down$loboth_log2FC, test_down$lodex_log2FC, paired = TRUE)
```

```
##
##  Paired t-test
##
## data:  test_down$loboth_log2FC and test_down$lodex_log2FC
## t = -8.6088, df = 239, p-value = 1.005e-15
## alternative hypothesis: true mean difference is not equal to 0
## 95 percent confidence interval:
##  -0.2873247 -0.1803154
## sample estimates:
## mean difference
##      -0.2338201
```

```
t.test(abs(lodex_reg_filt$loboth_log2FC), abs(lodex_reg_filt$lodex_log2FC), paired = TRU
E)
```

```
##
##  Paired t-test
##
## data:  abs(lodex_reg_filt$loboth_log2FC) and abs(lodex_reg_filt$lodex_log2FC)
## t = 9.5454, df = 603, p-value < 2.2e-16
## alternative hypothesis: true mean difference is not equal to 0
## 95 percent confidence interval:
##  0.1385816 0.2103782
## sample estimates:
## mean difference
##       0.1744799
```

```
mean(abs(lodex_reg_filt$loboth_log2FC)) - mean(abs(lodex_reg_filt$lodex_log2FC))
```

```
## [1] 0.1744799
```

```
t.test(abs(hidex_reg_filt$hiboth_log2FC), abs(hidex_reg_filt$hidex_log2FC), paired = TRU
E)
```

```
##
##  Paired t-test
##
## data:  abs(hidex_reg_filt$hiboth_log2FC) and abs(hidex_reg_filt$hidex_log2FC)
## t = 9.6827, df = 3480, p-value < 2.2e-16
## alternative hypothesis: true mean difference is not equal to 0
## 95 percent confidence interval:
##  0.04579721 0.06905314
## sample estimates:
## mean difference
##      0.05742518
```

```
mean(abs(hidex_reg_filt$hiboth_log2FC)) - mean(abs(hidex_reg_filt$hidex_log2FC))
```

```
## [1] 0.05742518
```

```
lodex_up_lng <- lodex_reg_filt %>%
  dplyr::select(lodex_log2FC, loboth_log2FC) %>%
  filter(loboth_log2FC > 0) %>%
  pivot_longer(cols = c("lodex_log2FC", "loboth_log2FC"), names_to = "treat", values_to
= "log2FC")

ggplot(lodex_up_lng, aes(treat, log2FC)) +
  geom_boxplot()
```

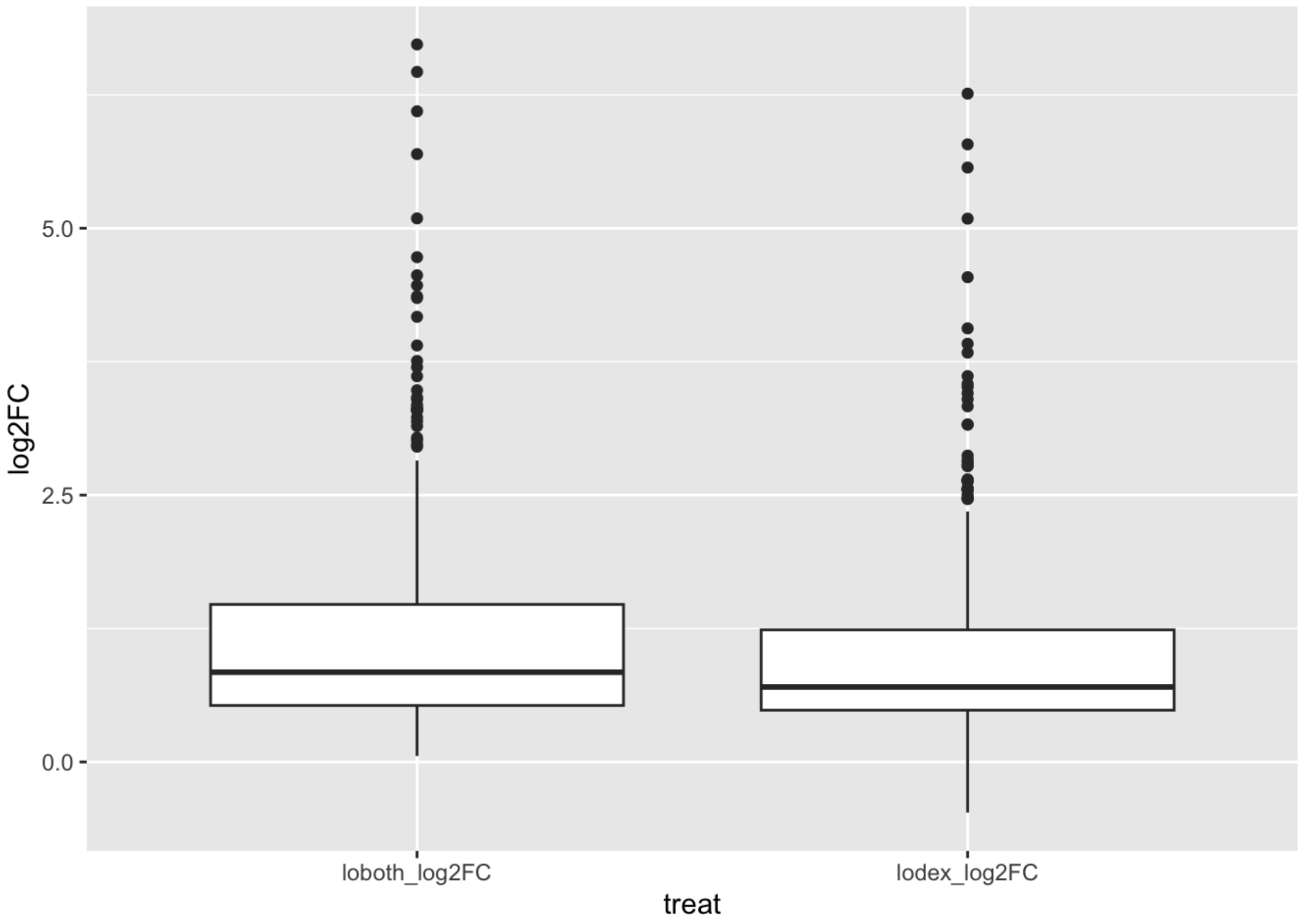

### Other comparisons

```
res.lob_lo <- results(dds, contrast = c("group", "5250", "50"))
head(res.lob_lo[ order(res.lob_lo$padj, decreasing = FALSE), ])
```

```
## log2 fold change (MLE): group 5250 vs 50
## Wald test p-value: group 5250 vs 50
## DataFrame with 6 rows and 6 columns
##                      baseMean log2FoldChange     lfcSE      stat      pvalue
##                     <numeric>      <numeric> <numeric> <numeric>   <numeric>
## ENSG00000134853.12 21714.814       0.903692 0.0575220  15.71037 1.28426e-55
## ENSG00000086730.17   839.007      -1.127217 0.0992807 -11.35383 7.09809e-30
## ENSG00000111371.16 14645.798       0.456088 0.0406732  11.21347 3.50181e-29
## ENSG00000127325.19   161.349       2.203259 0.1964678  11.21435 3.46725e-29
## ENSG00000145358.6    540.945      -1.327342 0.1330047  -9.97966 1.87102e-23
## ENSG00000168209.6   1767.982      -0.719078 0.0813786  -8.83620 9.90276e-19
##                           padj
##                      <numeric>
## ENSG00000134853.12 1.82827e-51
## ENSG00000086730.17 5.05242e-26
## ENSG00000111371.16 1.24629e-25
## ENSG00000127325.19 1.24629e-25
## ENSG00000145358.6  5.32716e-20
## ENSG00000168209.6  2.34960e-15
```

```
sum(res.lob_lo$padj < 0.01, na.rm=TRUE)
```

```
## [1] 572
```

```
summary(res.lob_lo, alpha = 0.01)
```

```
##
## out of 21242 with nonzero total read count
## adjusted p-value < 0.01
## LFC > 0 (up)       : 234, 1.1%
## LFC < 0 (down)     : 338, 1.6%
## outliers [1]       : 14, 0.066%
## low counts [2]     : 6992, 33%
## (mean count < 16)
## [1] see 'cooksCutoff' argument of ?results
## [2] see 'independentFiltering' argument of ?results
```

```
res.hib_hi <- results(dds, contrast = c("group", "50250", "500"))
head(res.hib_hi[ order(res.hib_hi$padj, decreasing = FALSE), ])
```

```
## log2 fold change (MLE): group 50250 vs 500
## Wald test p-value: group 50250 vs 500
## DataFrame with 6 rows and 6 columns
##                     baseMean log2FoldChange     lfcSE      stat      pvalue
##                    <numeric>      <numeric> <numeric> <numeric>   <numeric>
## ENSG00000134853.12 21714.814       0.793701 0.0577642  13.74038 5.81696e-43
## ENSG00000127325.19   161.349       2.365213 0.1891950  12.50146 7.32923e-36
## ENSG00000211672.2   1717.413       1.005801 0.0874044  11.50745 1.21000e-30
## ENSG00000235621.10   232.026       1.371629 0.1301860  10.53592 5.90037e-26
## ENSG00000285578.1    449.134       1.251376 0.1263179   9.90656 3.89837e-23
## ENSG00000065911.13  5270.897      -0.511357 0.0559595  -9.13798 6.36303e-20
##                           padj
##                      <numeric>
## ENSG00000134853.12 7.56263e-39
## ENSG00000127325.19 4.76437e-32
## ENSG00000211672.2  5.24375e-27
## ENSG00000235621.10 1.91777e-22
## ENSG00000285578.1  1.01366e-19
## ENSG00000065911.13 1.37876e-16
```

```
sum(res.hib_hi$padj < 0.01, na.rm=TRUE)
```

```
## [1] 511
```

```
summary(res.hib_hi, alpha = 0.01)
```

```
##
## out of 21242 with nonzero total read count
## adjusted p-value < 0.01
## LFC > 0 (up)       : 201, 0.95%
## LFC < 0 (down)     : 310, 1.5%
## outliers [1]       : 14, 0.066%
## low counts [2]     : 8227, 39%
## (mean count < 26)
## [1] see 'cooksCutoff' argument of ?results
## [2] see 'independentFiltering' argument of ?results
```

```
res.dex_hilo <- results(dds, contrast = c("group", "500", "50"))
head(res.dex_hilo[ order(res.dex_hilo$padj, decreasing = FALSE), ])
```

```
## log2 fold change (MLE): group 500 vs 50
## Wald test p-value: group 500 vs 50
## DataFrame with 6 rows and 6 columns
##                      baseMean log2FoldChange     lfcSE      stat       pvalue
##                     <numeric>      <numeric> <numeric> <numeric>    <numeric>
## ENSG00000265972.6   4263.711       2.100826 0.0891110   23.5754 6.89690e-123
## ENSG00000285417.1    472.855       2.678949 0.1241803   21.5731 3.21663e-103
## ENSG00000143119.14  5893.710       1.160749 0.0549125   21.1382  3.54572e-99
## ENSG00000096060.15  7856.732       0.958836 0.0539498   17.7728  1.14898e-70
## ENSG00000165810.17   568.781       2.128813 0.1223836   17.3946  9.06530e-68
## ENSG00000092820.19  4460.222       0.830240 0.0505469   16.4251  1.26416e-60
##                             padj
##                        <numeric>
## ENSG00000265972.6  1.12364e-118
## ENSG00000285417.1   2.62027e-99
## ENSG00000143119.14  1.92556e-95
## ENSG00000096060.15  4.67981e-67
## ENSG00000165810.17  2.95384e-64
## ENSG00000092820.19  3.43262e-57
```

```
sum(res.dex_hilo$padj < 0.01, na.rm=TRUE)
```

```
## [1] 1686
```

```
summary(res.dex_hilo, alpha = 0.01)
```

```
##
## out of 21242 with nonzero total read count
## adjusted p-value < 0.01
## LFC > 0 (up)       : 774, 3.6%
## LFC < 0 (down)     : 912, 4.3%
## outliers [1]       : 14, 0.066%
## low counts [2]     : 4936, 23%
## (mean count < 8)
## [1] see 'cooksCutoff' argument of ?results
## [2] see 'independentFiltering' argument of ?results
```

```
res.lob_idel <- results(dds, contrast = c("group", "5250", "0250"))
head(res.lob_idel[ order(res.lob_idel$padj, decreasing = FALSE), ])
```

```
## log2 fold change (MLE): group 5250 vs 0250
## Wald test p-value: group 5250 vs 0250
## DataFrame with 6 rows and 6 columns
##                      baseMean log2FoldChange     lfcSE      stat       pvalue
##                     <numeric>      <numeric> <numeric> <numeric>    <numeric>
## ENSG00000256235.3    1054.201        2.65312 0.0966499   27.4509 6.78403e-166
## ENSG00000248302.3     921.545        2.91832 0.1120499   26.0448 1.54093e-149
## ENSG00000159200.18   3711.420        1.66459 0.0708415   23.4973 4.34136e-122
## ENSG00000143119.14   5893.710        1.32565 0.0571890   23.1801 7.22828e-119
## ENSG00000164938.14   2806.620        4.20289 0.1873109   22.4380 1.67497e-111
## ENSG00000174944.9    3855.746        5.96869 0.2686013   22.2214 2.13455e-109
##                             padj
##                        <numeric>
## ENSG00000256235.3  1.04956e-161
## ENSG00000248302.3  1.19199e-145
## ENSG00000159200.18 2.23884e-118
## ENSG00000143119.14 2.79572e-115
## ENSG00000164938.14 5.18269e-108
## ENSG00000174944.9  5.50395e-106
```

```
sum(res.lob_idel$padj < 0.01, na.rm=TRUE)
```

```
## [1] 921
```

```
summary(res.lob_idel, alpha = 0.01)
```

```
##
## out of 21242 with nonzero total read count
## adjusted p-value < 0.01
## LFC > 0 (up)       : 480, 2.3%
## LFC < 0 (down)     : 441, 2.1%
## outliers [1]       : 14, 0.066%
## low counts [2]     : 5757, 27%
## (mean count < 11)
## [1] see 'cooksCutoff' argument of ?results
## [2] see 'independentFiltering' argument of ?results
```

```
res.hib_idel <- results(dds, contrast = c("group", "50250", "0250"))
head(res.hib_idel[ order(res.hib_idel$padj, decreasing = FALSE), ])
```

```
## log2 fold change (MLE): group 50250 vs 0250
## Wald test p-value: group 50250 vs 0250
## DataFrame with 6 rows and 6 columns
##                     baseMean log2FoldChange     lfcSE      stat       pvalue
##                    <numeric>      <numeric> <numeric> <numeric>    <numeric>
## ENSG00000143119.14  5893.710        2.44158 0.0566614   43.0906  0.00000e+00
## ENSG00000248302.3    921.545        4.20596 0.1104286   38.0876  0.00000e+00
## ENSG00000096060.15  7856.732        1.82493 0.0549998   33.1807 2.04391e-241
## ENSG00000159200.18  3711.420        2.32855 0.0705746   32.9942 9.83844e-239
## ENSG00000101445.10  3816.773        2.56407 0.0777833   32.9643 2.63702e-238
## ENSG00000265972.6   4263.711        2.95921 0.0902500   32.7890 8.44555e-236
##                            padj
##                       <numeric>
## ENSG00000143119.14  0.00000e+00
## ENSG00000248302.3   0.00000e+00
## ENSG00000096060.15 1.41820e-237
## ENSG00000159200.18 5.11992e-235
## ENSG00000101445.10 1.09784e-234
## ENSG00000265972.6  2.93004e-232
```

```
sum(res.hib_idel$padj < 0.01, na.rm=TRUE)
```

```
## [1] 4001
```

```
summary(res.hib_idel, alpha = 0.01)
```

```
##
## out of 21242 with nonzero total read count
## adjusted p-value < 0.01
## LFC > 0 (up)       : 1734, 8.2%
## LFC < 0 (down)     : 2267, 11%
## outliers [1]       : 14, 0.066%
## low counts [2]     : 412, 1.9%
## (mean count < 2)
## [1] see 'cooksCutoff' argument of ?results
## [2] see 'independentFiltering' argument of ?results
```

##Genes with an interaction between idela and dex

```
dds_int <- DESeqDataSet(gse, ~dex + idela + dex:idela)
```

```
## using counts and average transcript lengths from tximeta
```

```
#dds_int <- DESeqDataSetFromTximport(txi, cond, ~ idela + dex + idela:dex)
  #Filter
dds_int <- dds_int[ rowSums(counts(dds_int)) > 36, ]
  #Diff regulation
#Then actually look for differences between samples
dds_int <- DESeq(dds_int)
```

```
## estimating size factors
```

```
## using 'avgTxLength' from assays(dds), correcting for library size
```

```
## estimating dispersions
```

```
## gene-wise dispersion estimates
```

```
## mean-dispersion relationship
```

```
## final dispersion estimates
```

```
## fitting model and testing
```

```
resultsNames(dds_int)
```

```
## [1] "Intercept"      "dex_5_vs_0"     "dex_50_vs_0"    "idela_250_vs_0"
## [5] "dex5.idela250"  "dex50.idela250"
```

```
res_int <- results(dds_int)
```

### Generate results tables for coomparisons

These results should pretty much match the results from above

```
res_int.lodex <- results(dds_int, name = "dex_5_vs_0")
head(res_int.lodex[ order(res_int.lodex$padj, decreasing = FALSE), ])
```

```
## log2 fold change (MLE): dex 5 vs 0
## Wald test p-value: dex 5 vs 0
## DataFrame with 6 rows and 6 columns
##                     baseMean log2FoldChange     lfcSE      stat       pvalue
##                    <numeric>      <numeric> <numeric> <numeric>    <numeric>
## ENSG00000256235.3   1054.201        2.56699 0.1042310   24.6279 6.34414e-134
## ENSG00000159200.18  3711.420        1.71182 0.0739485   23.1488 1.49356e-118
## ENSG00000248302.3    921.545        3.08684 0.1333862   23.1421 1.74454e-118
## ENSG00000109501.15   804.747        3.61453 0.1660013   21.7741 4.08120e-105
## ENSG00000174944.9   3855.746        5.78615 0.2802596   20.6457  1.06754e-94
## ENSG00000143119.14  5893.710        1.14805 0.0573598   20.0149  4.08096e-89
##                            padj
##                       <numeric>
## ENSG00000256235.3  9.29226e-130
## ENSG00000159200.18 8.51742e-115
## ENSG00000248302.3  8.51742e-115
## ENSG00000109501.15 1.49443e-101
## ENSG00000174944.9   3.12724e-91
## ENSG00000143119.14  9.96230e-86
```

```
sum(res_int.lodex$padj < 0.01, na.rm=TRUE)
```

```
## [1] 649
```

```
summary(res_int.lodex, alpha = 0.01)
```

```
##
## out of 21242 with nonzero total read count
## adjusted p-value < 0.01
## LFC > 0 (up)       : 407, 1.9%
## LFC < 0 (down)     : 242, 1.1%
## outliers [1]       : 14, 0.066%
## low counts [2]     : 6581, 31%
## (mean count < 14)
## [1] see 'cooksCutoff' argument of ?results
## [2] see 'independentFiltering' argument of ?results
```

```
sh_res_int.lodex <- lfcShrink(dds_int, coef="dex_5_vs_0", type="apeglm")
```

```
## using 'apeglm' for LFC shrinkage. If used in published research, please cite:
##     Zhu, A., Ibrahim, J.G., Love, M.I. (2018) Heavy-tailed prior distributions for
##     sequence count data: removing the noise and preserving large differences.
##     Bioinformatics. https://doi.org/10.1093/bioinformatics/bty895
```

```
head(sh_res_int.lodex[ order(sh_res_int.lodex$padj, decreasing = FALSE), ])
```

```
## log2 fold change (MAP): dex 5 vs 0
## Wald test p-value: dex 5 vs 0
## DataFrame with 6 rows and 5 columns
##                     baseMean log2FoldChange     lfcSE       pvalue         padj
##                    <numeric>      <numeric> <numeric>    <numeric>    <numeric>
## ENSG00000256235.3   1054.201        2.55429 0.1043825 6.34414e-134 9.29226e-130
## ENSG00000159200.18  3711.420        1.70325 0.0740799 1.49356e-118 8.51742e-115
## ENSG00000248302.3    921.545        3.07536 0.1336594 1.74454e-118 8.51742e-115
## ENSG00000109501.15   804.747        3.59773 0.1662127 4.08120e-105 1.49443e-101
## ENSG00000174944.9   3855.746        5.75886 0.2808174  1.06754e-94  3.12724e-91
## ENSG00000143119.14  5893.710        1.13795 0.0574369  4.08096e-89  9.96230e-86
```

```
res_int.hidex <- results(dds_int, name = "dex_50_vs_0")
head(res_int.hidex[ order(res_int.hidex$padj, decreasing = FALSE), ])
```

```
## log2 fold change (MLE): dex 50 vs 0
## Wald test p-value: dex 50 vs 0
## DataFrame with 6 rows and 6 columns
##                     baseMean log2FoldChange     lfcSE      stat       pvalue
##                    <numeric>      <numeric> <numeric> <numeric>    <numeric>
## ENSG00000143119.14  5893.710        2.30880 0.0567756   40.6654  0.00000e+00
## ENSG00000096060.15  7856.732        1.97255 0.0550563   35.8279 4.06128e-281
## ENSG00000265972.6   4263.711        3.07612 0.0911625   33.7433 1.34237e-249
## ENSG00000248302.3    921.545        4.34853 0.1312012   33.1440 6.90993e-241
## ENSG00000159200.18  3711.420        2.39813 0.0734715   32.6403 1.09913e-233
## ENSG00000256235.3   1054.201        3.29998 0.1032100   31.9734 2.55304e-224
##                            padj
##                       <numeric>
## ENSG00000143119.14  0.00000e+00
## ENSG00000096060.15 4.22698e-277
## ENSG00000265972.6  9.31429e-246
## ENSG00000248302.3  3.59593e-237
## ENSG00000159200.18 4.57590e-230
## ENSG00000256235.3  8.85736e-221
```

```
sum(res_int.hidex$padj < 0.01, na.rm=TRUE)
```

```
## [1] 3779
```

```
summary(res_int.hidex, alpha = 0.01)
```

```
##
## out of 21242 with nonzero total read count
## adjusted p-value < 0.01
## LFC > 0 (up)       : 1696, 8%
## LFC < 0 (down)     : 2083, 9.8%
## outliers [1]       : 14, 0.066%
## low counts [2]     : 412, 1.9%
## (mean count < 2)
## [1] see 'cooksCutoff' argument of ?results
## [2] see 'independentFiltering' argument of ?results
```

```
res_int.idel <- results(dds_int, name = "idela_250_vs_0")
head(res_int.idel[ order(res_int.idel$padj, decreasing = FALSE), ])
```

```
## log2 fold change (MLE): idela 250 vs 0
## Wald test p-value: idela 250 vs 0
## DataFrame with 6 rows and 6 columns
##                     baseMean log2FoldChange     lfcSE      stat       pvalue
##                    <numeric>      <numeric> <numeric> <numeric>    <numeric>
## ENSG00000134853.12 21714.814       1.334182 0.0578140   23.0771 7.85601e-118
## ENSG00000165507.9    621.084       1.959074 0.1356858   14.4383  2.97021e-47
## ENSG00000170365.10  8105.290       0.615688 0.0472780   13.0227  9.08834e-39
## ENSG00000086730.17   839.007      -1.132965 0.0956698  -11.8425  2.35466e-32
## ENSG00000087495.17   965.110       1.176041 0.0999210   11.7697  5.59152e-32
## ENSG00000107537.14  1409.967       1.199796 0.1132524   10.5940  3.17723e-26
##                            padj
##                       <numeric>
## ENSG00000134853.12 1.15067e-113
## ENSG00000165507.9   2.17524e-43
## ENSG00000170365.10  4.43723e-35
## ENSG00000086730.17  8.62219e-29
## ENSG00000087495.17  1.63798e-28
## ENSG00000107537.14  7.75614e-23
```

```
sum(res_int.idel$padj < 0.01, na.rm=TRUE)
```

```
## [1] 418
```

```
summary(res_int.idel, alpha = 0.01)
```

```
##
## out of 21242 with nonzero total read count
## adjusted p-value < 0.01
## LFC > 0 (up)       : 208, 0.98%
## LFC < 0 (down)     : 210, 0.99%
## outliers [1]       : 14, 0.066%
## low counts [2]     : 6581, 31%
## (mean count < 14)
## [1] see 'cooksCutoff' argument of ?results
## [2] see 'independentFiltering' argument of ?results
```

```
test2 <- results(dds_int, contrast = list( c("dex_50_vs_0", "idela_250_vs_0", "dex50.ide
la250")))
summary(test2, alpha = 0.01)
```

```
##
## out of 21242 with nonzero total read count
## adjusted p-value < 0.01
## LFC > 0 (up)       : 2164, 10%
## LFC < 0 (down)     : 2802, 13%
## outliers [1]       : 14, 0.066%
## low counts [2]     : 824, 3.9%
## (mean count < 3)
## [1] see 'cooksCutoff' argument of ?results
## [2] see 'independentFiltering' argument of ?results
```

These are the interaction results.

The interaction term, answering: is the condition effect *different* across genotypes? In this case, idela is the genotype - so does dex do something different in the presence of idela? results(dds, name="idela250.dex…")

```
res_int.5int <- results(dds_int, name = "dex5.idela250")
head(res_int.5int[ order(res_int.5int$padj, decreasing = FALSE), ])
```

```
## log2 fold change (MLE): dex5.idela250
## Wald test p-value: dex5.idela250
## DataFrame with 6 rows and 6 columns
##                      baseMean log2FoldChange     lfcSE      stat      pvalue
##                     <numeric>      <numeric> <numeric> <numeric>   <numeric>
## ENSG00000100280.17   2462.281       0.653120 0.1177848   5.54503 2.93910e-08
## ENSG00000165507.9     621.084      -0.983224 0.1808470  -5.43677 5.42545e-08
## ENSG00000111371.16  14645.798       0.308036 0.0578936   5.32073 1.03351e-07
## ENSG00000134853.12  21714.814      -0.430490 0.0815552  -5.27851 1.30238e-07
## ENSG00000088305.19    562.270       0.888653 0.1700857   5.22474 1.74401e-07
## ENSG00000066923.18    865.979       0.828551 0.1675293   4.94571 7.58675e-07
##                            padj
##                       <numeric>
## ENSG00000100280.17 0.000564681
## ENSG00000165507.9  0.000564681
## ENSG00000111371.16 0.000677758
## ENSG00000134853.12 0.000677758
## ENSG00000088305.19 0.000726065
## ENSG00000066923.18 0.002632095
```

```
sum(res_int.5int$padj < 0.05, na.rm=TRUE)
```

```
## [1] 18
```

```
summary(res_int.5int, alpha = 0.05)
```

```
##
## out of 21242 with nonzero total read count
## adjusted p-value < 0.05
## LFC > 0 (up)       : 13, 0.061%
## LFC < 0 (down)     : 5, 0.024%
## outliers [1]       : 14, 0.066%
## low counts [2]     : 412, 1.9%
## (mean count < 2)
## [1] see 'cooksCutoff' argument of ?results
## [2] see 'independentFiltering' argument of ?results
```

```
res_int.50int <- results(dds_int, name = "dex50.idela250")
head(res_int.50int[ order(res_int.50int$padj, decreasing = FALSE), ])
```

```
## log2 fold change (MLE): dex50.idela250
## Wald test p-value: dex50.idela250
## DataFrame with 6 rows and 6 columns
##                      baseMean log2FoldChange     lfcSE      stat      pvalue
##                     <numeric>      <numeric> <numeric> <numeric>   <numeric>
## ENSG00000165507.9    621.084      -1.361687 0.1790091  -7.60680 2.80962e-14
## ENSG00000134853.12 21714.814      -0.540481 0.0817261  -6.61332 3.75801e-11
## ENSG00000103995.14  4195.779      -0.620638 0.1034555  -5.99908 1.98444e-09
## ENSG00000170365.10  8105.290      -0.392716 0.0658787  -5.96120 2.50397e-09
## ENSG00000120833.14  5901.363      -0.632532 0.1101617  -5.74185 9.36483e-09
## ENSG00000157557.13  2168.807      -0.494089 0.0926364  -5.33364 9.62654e-08
##                           padj
##                      <numeric>
## ENSG00000165507.9  4.80838e-10
## ENSG00000134853.12 3.21573e-07
## ENSG00000103995.14 1.07132e-05
## ENSG00000170365.10 1.07132e-05
## ENSG00000120833.14 3.20539e-05
## ENSG00000157557.13 2.74581e-04
```

```
sum(res_int.50int$padj < 0.05, na.rm=TRUE)
```

```
## [1] 72
```

```
summary(res_int.50int, alpha = 0.05)
```

```
##
## out of 21242 with nonzero total read count
## adjusted p-value < 0.05
## LFC > 0 (up)       : 30, 0.14%
## LFC < 0 (down)     : 42, 0.2%
## outliers [1]       : 14, 0.066%
## low counts [2]     : 4114, 19%
## (mean count < 6)
## [1] see 'cooksCutoff' argument of ?results
## [2] see 'independentFiltering' argument of ?results
```

### Make a list of genes with strong interaction terms

#### Function to add Gene symbols

```
res.5int <- add_geneids(res_int.5int)
```

```
## 'select()' returned 1:many mapping between keys and columns
## 'select()' returned 1:many mapping between keys and columns
## 'select()' returned 1:many mapping between keys and columns
```

```
res.5int_sig <- res.5int %>%
  as.data.frame() %>%
  rownames_to_column('ensembl') %>%
  dplyr::filter(padj <= 0.05, abs(log2FoldChange) <= 10)

res.50int <- add_geneids(res_int.50int)
```

```
## 'select()' returned 1:many mapping between keys and columns
## 'select()' returned 1:many mapping between keys and columns
## 'select()' returned 1:many mapping between keys and columns
```

```
res.50int_sig <- res.50int %>%
  as.data.frame() %>%
  rownames_to_column('ensembl') %>%
  dplyr::filter(padj <= 0.05, abs(log2FoldChange) <= 10)

#Are any of these common between them?

common_int <- res.5int_sig %>%
  inner_join(res.50int_sig, by = "ensembl")
```

### Bar charts for key genes

Join results for lowdex into table

New function to add gene IDs to a tibble

```
add_geneids_tbl <- function(genelist) {
  genelist$symbol <- mapIds(org.Hs.eg.db, keys=str_sub(genelist$ensembl, 1, 15), column
="SYMBOL", keytype="ENSEMBL", multiVals="first")
  genelist$entrez <- mapIds(org.Hs.eg.db, keys=str_sub(genelist$ensembl, 1, 15), column
="ENTREZID", keytype="ENSEMBL", multiVals="first")
  genelist$genename <- mapIds(org.Hs.eg.db, keys=str_sub(genelist$ensembl, 1, 15), colum
n="GENENAME", keytype="ENSEMBL", multiVals="first")
#  genelist <- genelist %>% drop_na(log2FoldChange)
  return(genelist)
}
```

```
idela_drg <- res.idel %>%
  as.data.frame()
names(idela_drg) <- paste0("idela.", names(idela_drg))
idela_drg <- idela_drg %>%
  rownames_to_column(var = "ensembl")

lodex_drg <- res.lodex %>%
  as.data.frame()
names(lodex_drg) <- paste0("lodex.", names(lodex_drg))
lodex_drg <- lodex_drg %>%
  rownames_to_column(var = "ensembl")

lob_drg <- res.lob %>%
  as.data.frame()
names(lob_drg) <- paste0("loboth.", names(lob_drg))
lob_drg <- lob_drg %>%
  rownames_to_column(var = "ensembl")

lodex_merge <- idela_drg %>%
  left_join(lodex_drg, by = "ensembl") %>%
  left_join(lob_drg, by = "ensembl") %>%
  add_geneids_tbl() %>%
  arrange(symbol, ensembl) %>%
  filter(!duplicated(symbol))
```

```
## 'select()' returned 1:many mapping between keys and columns
## 'select()' returned 1:many mapping between keys and columns
## 'select()' returned 1:many mapping between keys and columns
```

```
lodex_merge_longer <- lodex_merge %>%
  pivot_longer(idela.baseMean:loboth.padj, names_to = c("treatment",".value"), names_pat
tern = "(^\\w+).(\\w+$)") %>%
  mutate(treatment = factor(treatment, levels = c("idela", "lodex","loboth")))
```

### Plot normally regulated genes

Start with a set that we know

```
reg_genes <- c("RCAN1", "KCNJ2", "FKBP5", "P2RY14", "SOCS2", "IL7R")
lodex_merge_longer %>%
  dplyr::filter(symbol %in% reg_genes) %>%
  ggplot(aes(treatment, log2FoldChange, fill = treatment)) +
  labs(x = "", y = "log2FoldChange") +
  geom_col(position = "dodge") +
  scale_fill_viridis(discrete = T, option = "E") +
  facet_wrap(~symbol) +theme_bw() +
  theme(legend.position="none", axis.text.x = element_text(angle = 45, hjust = 1)) +
  geom_errorbar(aes(ymin=log2FoldChange-lfcSE, ymax=log2FoldChange+lfcSE), position = po
sition_dodge(width = 0.9), width=0.5, colour="black", size = 0.5)
```

```
## Warning: Using `size` aesthetic for lines was deprecated in ggplot2 3.4.0.
## ℹ Please use `linewidth` instead.
## This warning is displayed once every 8 hours.
## Call `lifecycle::last_lifecycle_warnings()` to see where this warning was
## generated.
```

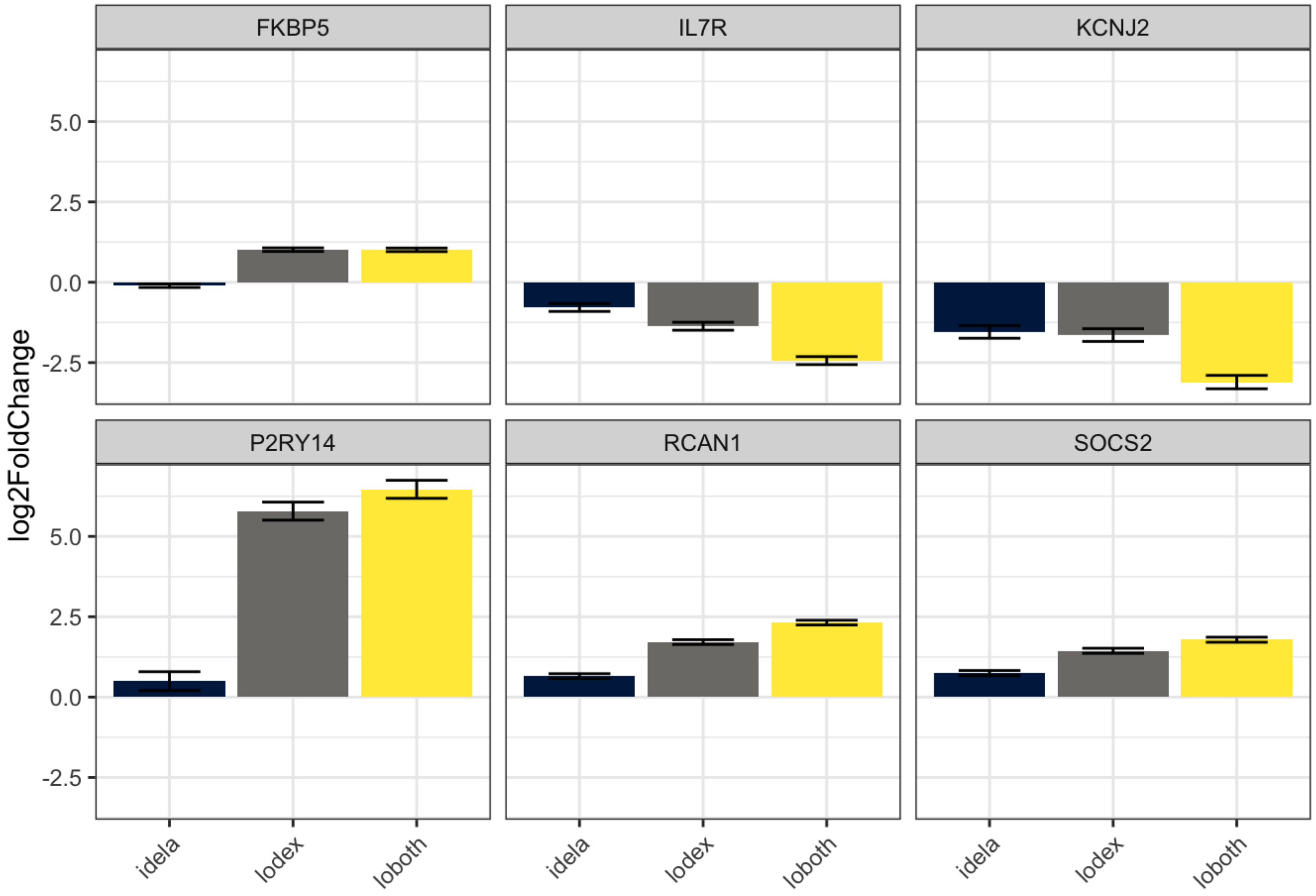

```
reg_genes <- c("BCL2L11", "BCL2", "IL7R", "TXNIP", "MBNL1", "BMF")
lodex_merge_longer %>%
  dplyr::filter(symbol %in% reg_genes) %>%
  ggplot(aes(treatment, log2FoldChange, fill = treatment)) +
  labs(x = "", y = "log2FoldChange") +
  geom_col(position = "dodge") +
  scale_fill_viridis(discrete = T, option = "E") +
  facet_wrap(~symbol) +theme_bw() +
  theme(legend.position="none", axis.text.x = element_text(angle = 45, hjust = 1)) +
  geom_errorbar(aes(ymin=log2FoldChange-lfcSE, ymax=log2FoldChange+lfcSE), position = po
sition_dodge(width = 0.9), width=0.5, colour="black", size = 0.5)
```

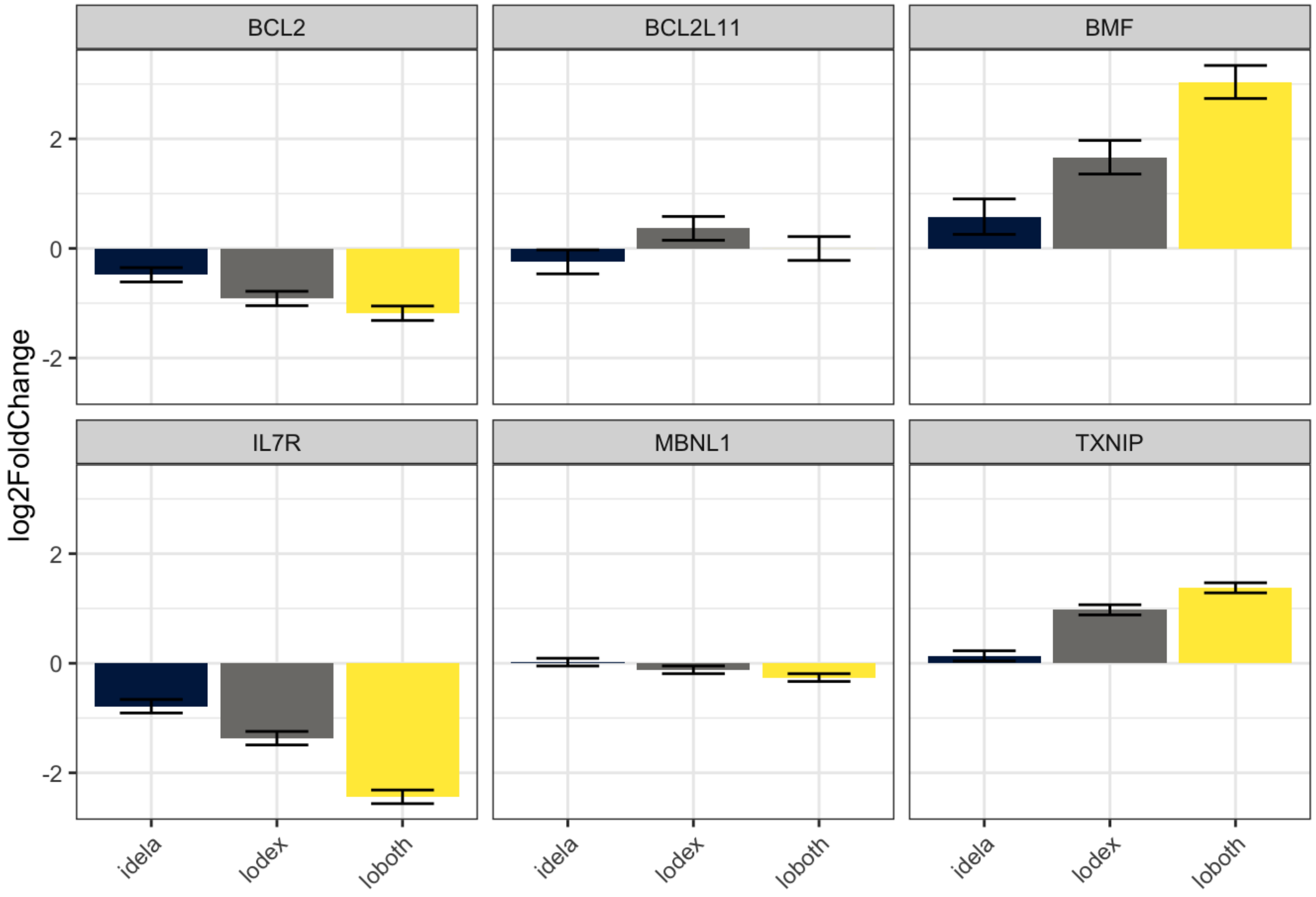

### Plot interaction genes

```
int_genes <- res.5int_sig %>%
  filter(abs(log2FoldChange) < 10) %>%
  na.omit() %>%
  pull(symbol)

lodex_merge_longer %>%
  dplyr::filter(symbol %in% int_genes) %>%
  ggplot(aes(treatment, log2FoldChange, fill = treatment)) +
  labs(x = "", y = "log2FoldChange") +
  geom_col(position = "dodge") +
  scale_fill_viridis(discrete = T, option = "E") +
  facet_wrap(~symbol) + theme_bw() +
  theme(legend.position="none", axis.text.x = element_text(angle = 45, hjust = 1)) +
  geom_errorbar(aes(ymin=log2FoldChange-lfcSE, ymax=log2FoldChange+lfcSE), position = po
sition_dodge(width = 0.9), width=0.5, colour="black", size = 0.5)
```

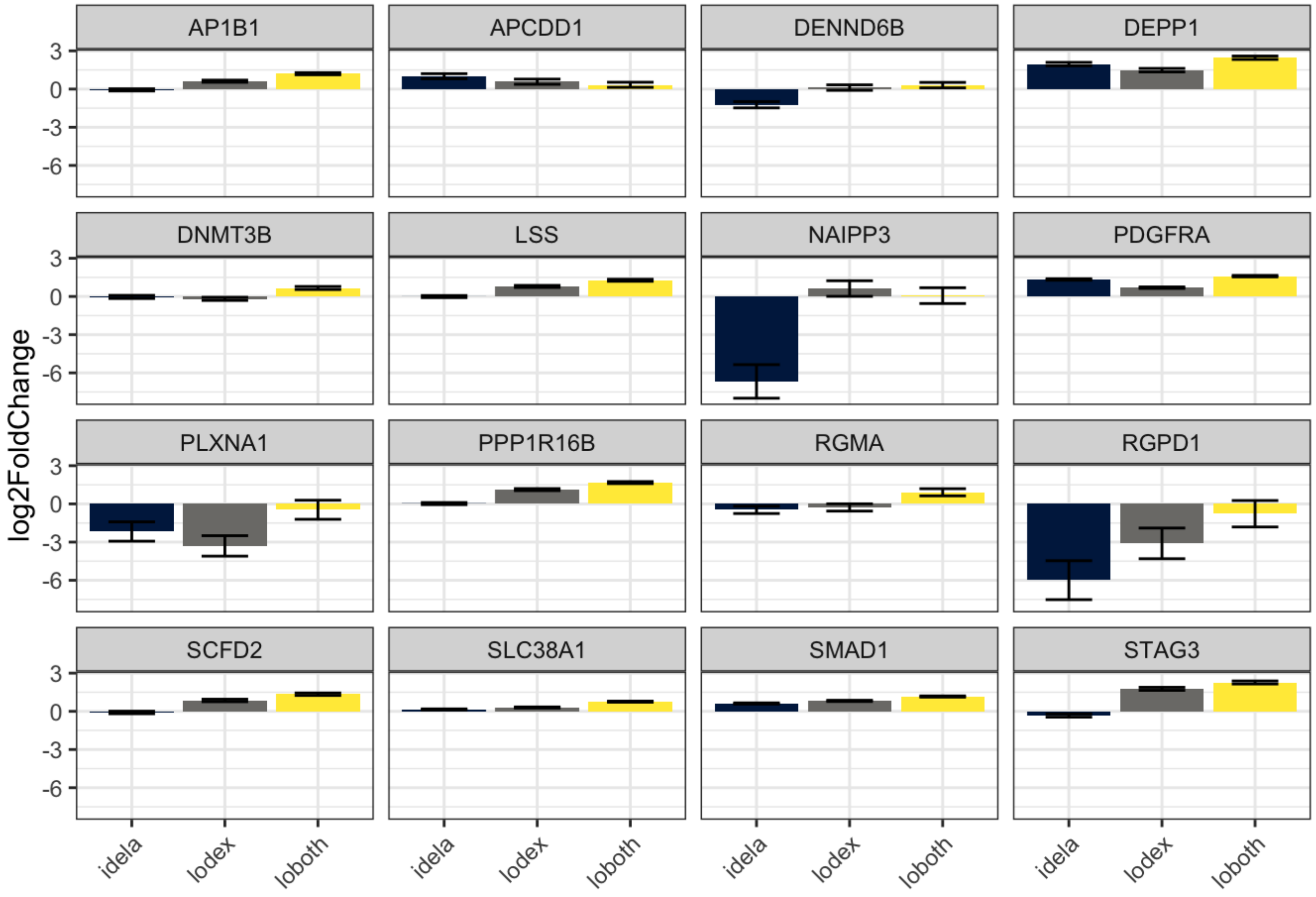

### Identifying Effector Genes

Effector genes are those genes regulated by chemotherapeutic drugs that contribute to B-ALL cell death. We have four sets of regulated genes:

1. Regulated by low dex (5 nM)
2. Regulated by high dex (50 nM)
3. Regulated by low dex + idela
4. Regulated by high dex + idela

The question is, which regulated genes contribute to cell death? We have two measures of contribution to cell death - shRNA screen measuring the effect of each gene on *growth* and *sensitivity* to dex. We did this screen in two ways, in still flasks for 6,000 genes (CAGEK) and in stir flasks for all protein coding genes (full).

Using these screens we can identify genes as:

Positive Effectors - when upregulated they contribute to cell death. This is evidenced as upregulation and more resistant when knocked down Negative Effectors - when downregulated they contribute to cell death. This is evidenced as downregulation and more sensitive when knocked down Buffering genes - when regulation opposes cell death. This is evidenced by upregulation, but more sensitive when knocked down, or vice versa.

This is best visualized by a scatter plot with the screen phenotype on the x-axis and the fold regulation on the y-axis.

```
#Merge screen and regulation data
names(full_rhos) <- c("number", "entrez", "symbol", "genename", "shRNAs", "Remaining_shR
NAs", "Rho_phenotype", "Rho_pvalue", "Rho_qvalue")
full_rhos_min <- full_rhos[,c(3,7,8)]
names(cagek_rhos) <- c("entrez", "symbol", "genename", "shRNAs", "Remaining_shRNAs", "CA
GEK_Rho_phenotype", "CAGEK_Rho_pvalue")
```

```
## Warning: The `value` argument of `names<-` must have the same length as `x` as of tib
ble
## 3.0.0.
## This warning is displayed once every 8 hours.
## Call `lifecycle::last_lifecycle_warnings()` to see where this warning was
## generated.
```

```
## Warning: The `value` argument of `names<-` can't be empty as of tibble 3.0.0.
## This warning is displayed once every 8 hours.
## Call `lifecycle::last_lifecycle_warnings()` to see where this warning was
## generated.
```

```
cagek_rhos_min <- cagek_rhos[,c(1,6,7)] %>%
  mutate(entrez = as.character(entrez))
names(full_gammas) <- c("number", "entrez", "symbol", "genename", "shRNAs", "Remaining_s
hRNAs", "Gamma_phenotype", "Gamma_pvalue", "Gamma_qvalue")
full_gammas_min <- full_gammas[,c(3,7,8)]
names(cagek_gammas) <- c("entrez", "symbol", "genename", "shRNAs", "Remaining_shRNAs",
"CAGEK_Gamma_phenotype", "CAGEK_Gamma_pvalue")
cagek_gammas_min <- cagek_gammas[,c(1,6,7)] %>%
  mutate(entrez = as.character(entrez))

# Just makes tables with both expression and screen phenotype data

full_effector_tbl <- sum_tbl_2 %>%
  inner_join(full_rhos_min, by = "symbol") %>%
  inner_join(full_gammas_min, by = "symbol")

cagek_effector_tbl <- sum_tbl_2 %>%
  inner_join(cagek_rhos_min, by = "entrez") %>%
  inner_join(cagek_gammas_min, by = "entrez")
```

### What are the effector genes for High Dex?

1. Positive Effectors

```r
pos_full_effectors <- full_effector_tbl %>%
  filter(Rho_pvalue < 0.05 & hidex_adjp < 0.05 & Rho_phenotype > 0 & hidex_log2FC > 0)

write.csv(pos_full_effectors, "pos_nalm6_full_effectors.csv")

pos_cagek_effectors <- cagek_effector_tbl %>%
  filter(CAGEK_Rho_pvalue < 0.05 & hidex_adjp < 0.05 & CAGEK_Rho_phenotype > 0 & hidex_l
og2FC > 0)

all_pos_eff <- pos_full_effectors %>%
  full_join(pos_cagek_effectors, by = "symbol")

common_pos_eff <- pos_full_effectors %>%
  inner_join(pos_cagek_effectors, by = "symbol")
```

```r
ggplot(common_pos_eff, aes(hidex_log2FC.x, hiboth_log2FC.x, label = symbol)) +
  labs(title = "Highest confidence positive effectors", y = "50 nM Dex + Idela, log2FC",
x = "50 nM Dex, log2FC") +
  geom_point() +
  geom_text_repel(size = 3, max.overlaps = 12, max.iter = 1000000, label.padding = 0.1)
+
  geom_abline(slope = 1, intercept = 0, color = "red", size = 1) +
  xlim(0,5.1) + ylim(0, 5.1) +
  theme_bw() +
  theme(axis.title = element_text(face="bold"), plot.title = element_text(face = "bol
d")) +
  annotate("rect", xmin = 0, xmax = 5.1, ymin = 0, ymax = 5.1, fill= "purple", alpha =
0.1)
```

```
## Warning in geom_text_repel(size = 3, max.overlaps = 12, max.iter = 1e+06, :
## Ignoring unknown parameters: `label.padding`
```

```
## Warning: ggrepel: 9 unlabeled data points (too many overlaps). Consider
## increasing max.overlaps
```

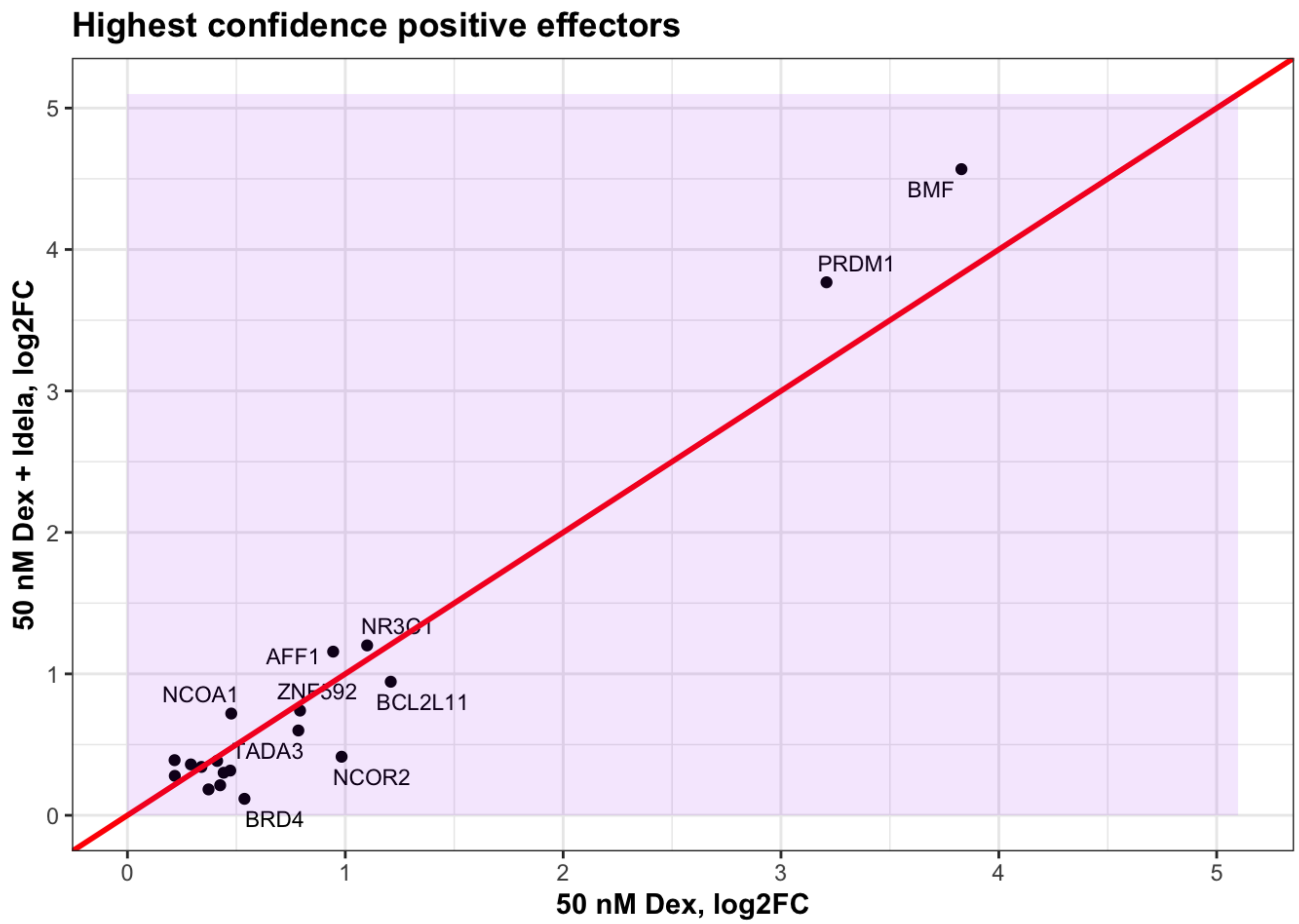

```
ggsave("common_pos_effectors.pdf", width = 5, height= 5)
```

```
## Warning: ggrepel: 9 unlabeled data points (too many overlaps). Consider
## increasing max.overlaps
```

Pos effectors at low dex

```
ggplot(common_pos_eff, aes(lodex_log2FC.x,loboth_log2FC.x, label = symbol)) +
  labs(title = "Highest confidence positive effectors", y = "5 nM Dex + Idela, log2FC",
x = "5 nM Dex, log2FC") +
  geom_point() +
  geom_text_repel(size = 3, max.overlaps = 12, max.iter = 1000000, label.padding = 0.1)
+
  geom_abline(slope = 1, intercept = 0, color = "red", size = 1) +
  xlim(0, 3.5) + ylim(0, 3.5) +
  theme_bw() +
  theme(axis.title = element_text(face="bold"), plot.title = element_text(face = "bol
d")) +
  annotate("rect", xmin = 0, xmax = 3.5, ymin = 0, ymax = 3.5, fill= "purple", alpha =
0.1)
```

```
## Warning in geom_text_repel(size = 3, max.overlaps = 12, max.iter = 1e+06, :
## Ignoring unknown parameters: `label.padding`
```

```
## Warning: Removed 3 rows containing missing values (`geom_point()`).
```

```
## Warning: Removed 3 rows containing missing values (`geom_text_repel()`).
```

```
## Warning: ggrepel: 8 unlabeled data points (too many overlaps). Consider
## increasing max.overlaps
```

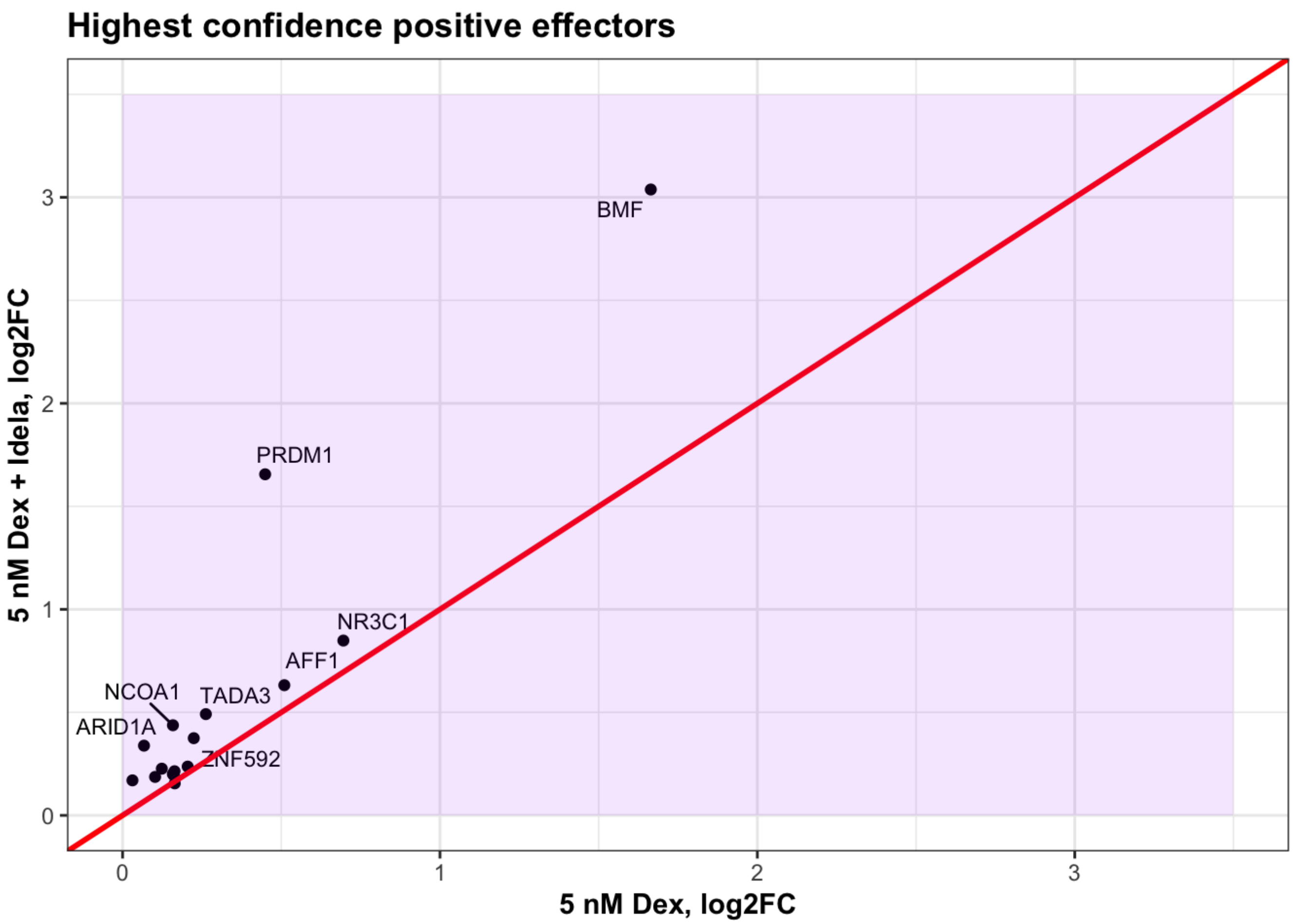

Does addition of idela cause regulation of common positive effectors to look like hidex?

```
ggplot(common_pos_eff) +
  geom_point(aes(lodex_log2FC.x, loboth_log2FC.x), color = "black") +
  geom_point(aes(lodex_log2FC.x, hidex_log2FC.x, label = symbol), color = "red") +
  labs(y = "50 nM Dex, red, 5 nM Dex + Idela, black", x = "5 nM Dex, log2FC") +
  #geom_text_repel(size = 3, max.overlaps = 12, max.iter = 1000000, label.padding = 0.1)
+
  geom_abline(slope = 1, intercept = 0, color = "red", size = 1) +
  # xlim(0, 3.5) + ylim(0, 3.5) +
  theme_bw() +
  theme(axis.title = element_text(face="bold"), plot.title = element_text(face = "bol
d")) +
  annotate("rect", xmin = 0, xmax = 3.5, ymin = 0, ymax = 3.5, fill= "purple", alpha =
0.1)
```

```
## Warning in geom_point(aes(lodex_log2FC.x, hidex_log2FC.x, label = symbol), :
## Ignoring unknown aesthetics: label
```

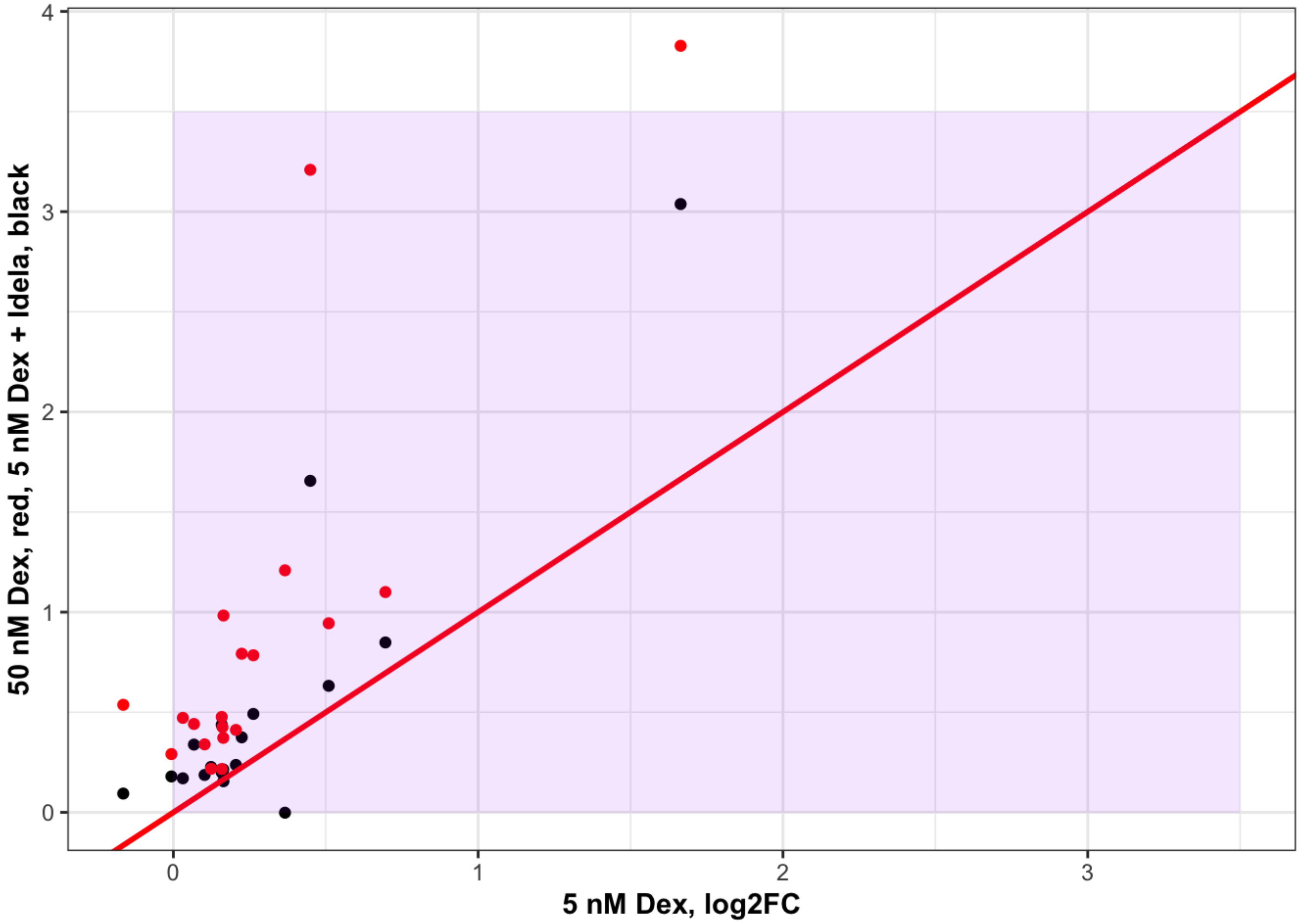

```
cpe_lng <- common_pos_eff %>%
  select(symbol, lodex_log2FC.x, loboth_log2FC.x, hidex_log2FC.x, hiboth_log2FC.x) %>%
  pivot_longer(cols = c("lodex_log2FC.x", "loboth_log2FC.x", "hidex_log2FC.x", "hiboth_l
og2FC.x"), values_to = "log2FC", names_to = "Condition")
ggplot(cpe_lng, aes(x= log2FC, y= reorder(symbol, log2FC))) +
    geom_line() +
    geom_point(aes(color=Condition), size=2) +
    scale_color_viridis(discrete = T, option = "E") +
    theme_bw() + ylab("Effector Gene")+
    theme(legend.position="top")
```

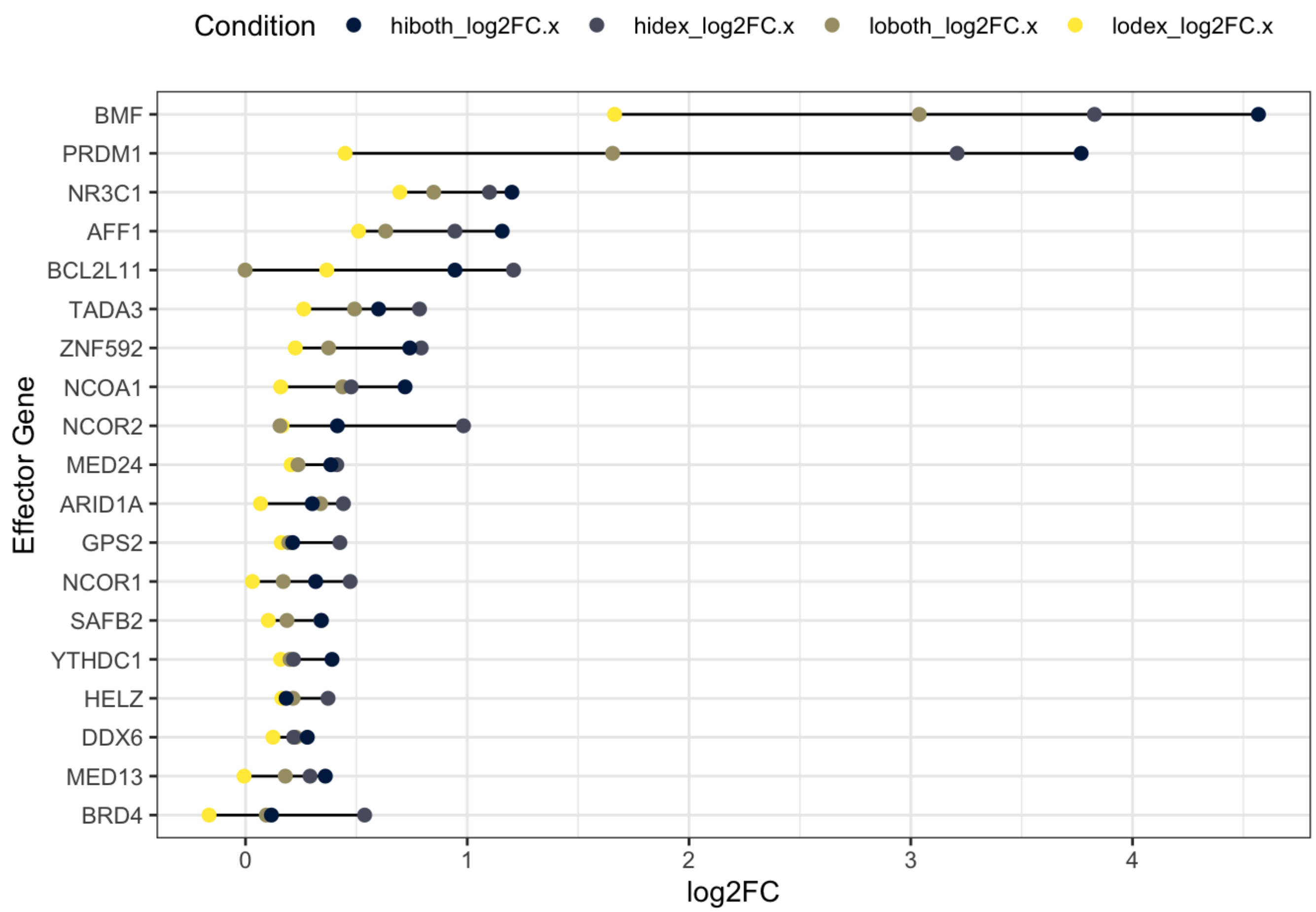

```
ggsave("pos_eff_reg_cond.pdf", width = 8, height = 4)
```

2. Negative Effectors

```
neg_full_effectors <- full_effector_tbl %>%
  filter(Rho_pvalue < 0.05 & hidex_adjp < 0.05 & Rho_phenotype < 0 & hidex_log2FC < 0)

write.csv(neg_full_effectors, "neg_nalm6_full_effectors.csv")

neg_cagek_effectors <- cagek_effector_tbl %>%
  filter(CAGEK_Rho_pvalue < 0.05 & hidex_adjp < 0.05 & CAGEK_Rho_phenotype < 0 & hidex_l
og2FC < 0)

all_neg_eff <- neg_full_effectors %>%
  full_join(neg_cagek_effectors, by = "symbol")

common_neg_eff <- neg_full_effectors %>%
  inner_join(neg_cagek_effectors, by = "symbol")

ggplot(common_neg_eff, aes(hidex_log2FC.x, hiboth_log2FC.x, label = symbol)) +
  labs(title = "Highest confidence negative effectors", y = "50 nM Dex + Idela, log2FC",
x = "50 nM Dex, log2FC") +
  geom_point() +
  geom_text_repel(size = 3, max.overlaps = 12, max.iter = 1000000) +
  geom_abline(slope = 1, intercept = 0, color = "red", size = 1) +
  xlim(-1.6, 0.1) + ylim(-1.6, 0.1) +
  theme_bw() +
  theme(axis.title = element_text(face="bold"), plot.title = element_text(face = "bol
d")) +
  annotate("rect", xmin = -1.6, xmax = 0.1, ymin = -1.6, ymax = 0.1 , fill= "green", alp
ha = 0.1)
```

```
## Warning: Removed 1 rows containing missing values (`geom_point()`).
```

```
## Warning: Removed 1 rows containing missing values (`geom_text_repel()`).
```

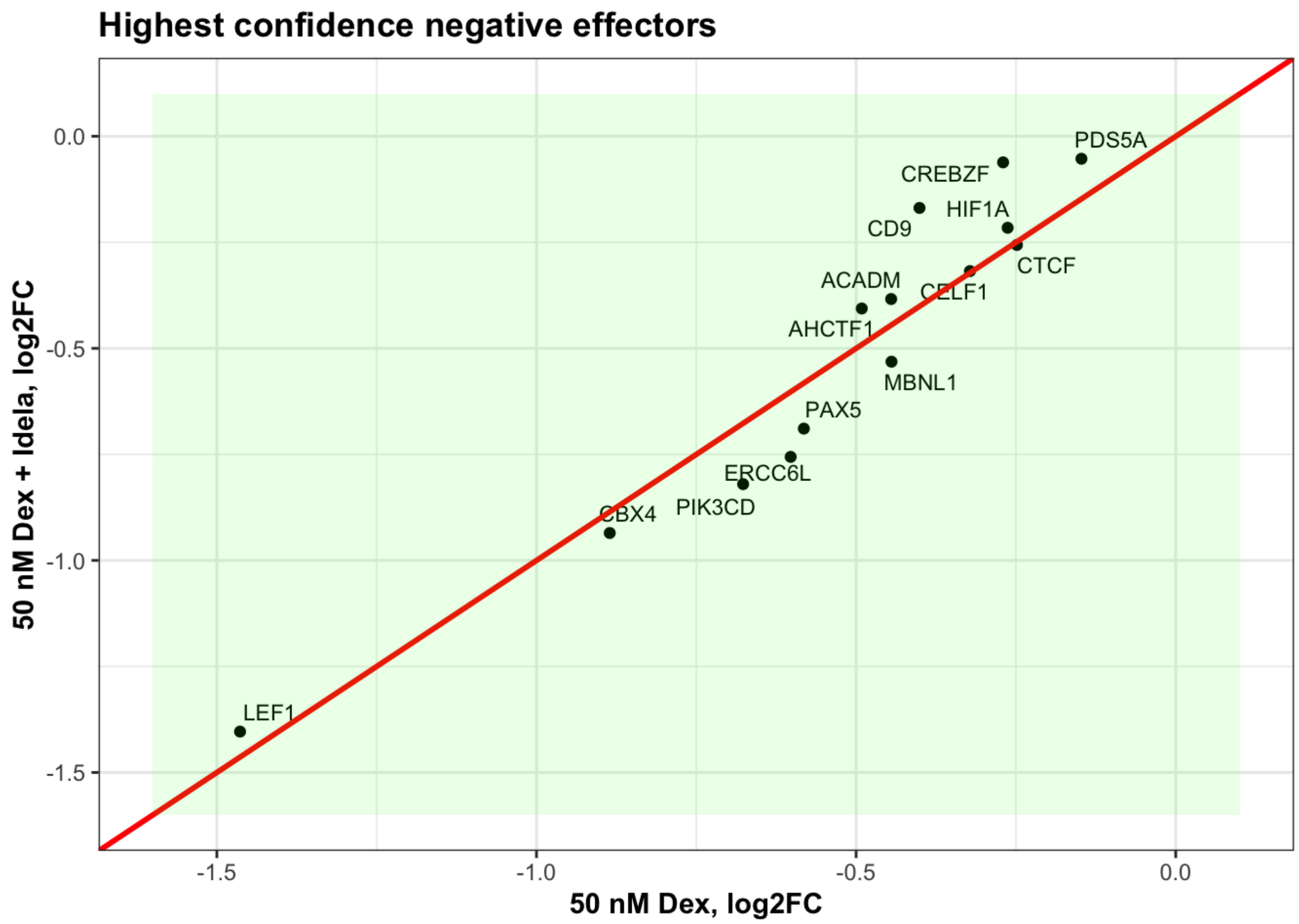

```
ggsave("common_neg_effectors.pdf", width = 5, height= 5)
```

```
## Warning: Removed 1 rows containing missing values (`geom_point()`).
## Removed 1 rows containing missing values (`geom_text_repel()`).
```

```
cne_lng <- common_neg_eff %>%
  select(symbol, lodex_log2FC.x, loboth_log2FC.x, hidex_log2FC.x, hiboth_log2FC.x) %>%
  pivot_longer(cols = c("lodex_log2FC.x", "loboth_log2FC.x", "hidex_log2FC.x", "hiboth_l
og2FC.x"), values_to = "log2FC", names_to = "Condition")
ggplot(cne_lng, aes(x= log2FC, y= reorder(symbol, log2FC))) +
    geom_line() +
    geom_point(aes(color=Condition), size=2) +
    scale_color_viridis(discrete = T, option = "E") +
    theme_bw() + ylab("Effector Gene")+
    theme(legend.position="top")
```

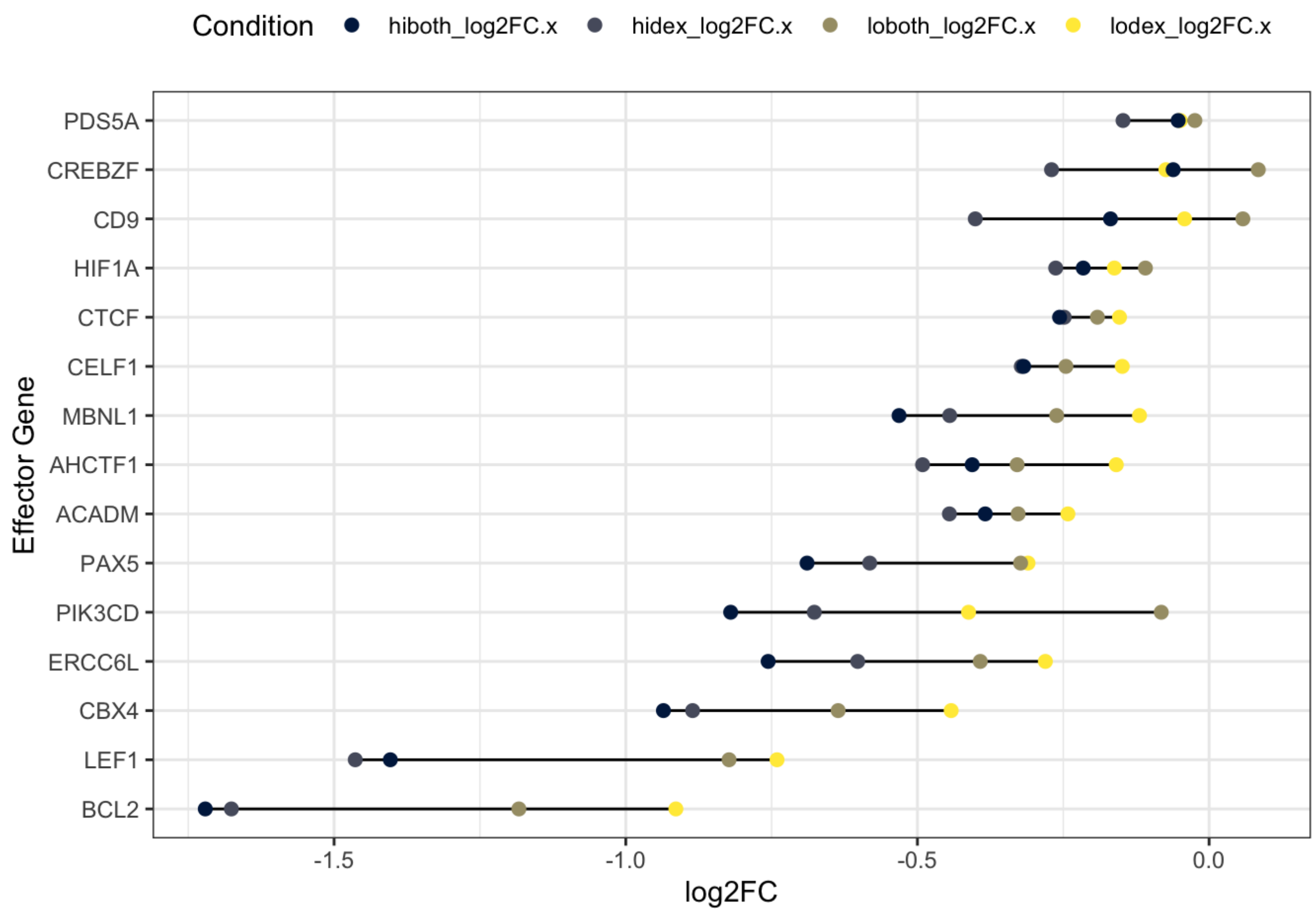

```
ggsave("neg_eff_reg_cond.pdf", width = 8, height = 4)

ce_lng <- bind_rows(cpe_lng, cne_lng)

ggplot(ce_lng, aes(x= log2FC, y= reorder(symbol, log2FC))) +
    geom_line() +
    geom_point(aes(color=Condition), size=2) +
    scale_color_viridis(discrete = T, option = "E") +
    theme_bw() + ylab("Effector Gene")+
    theme(legend.position="top")
```

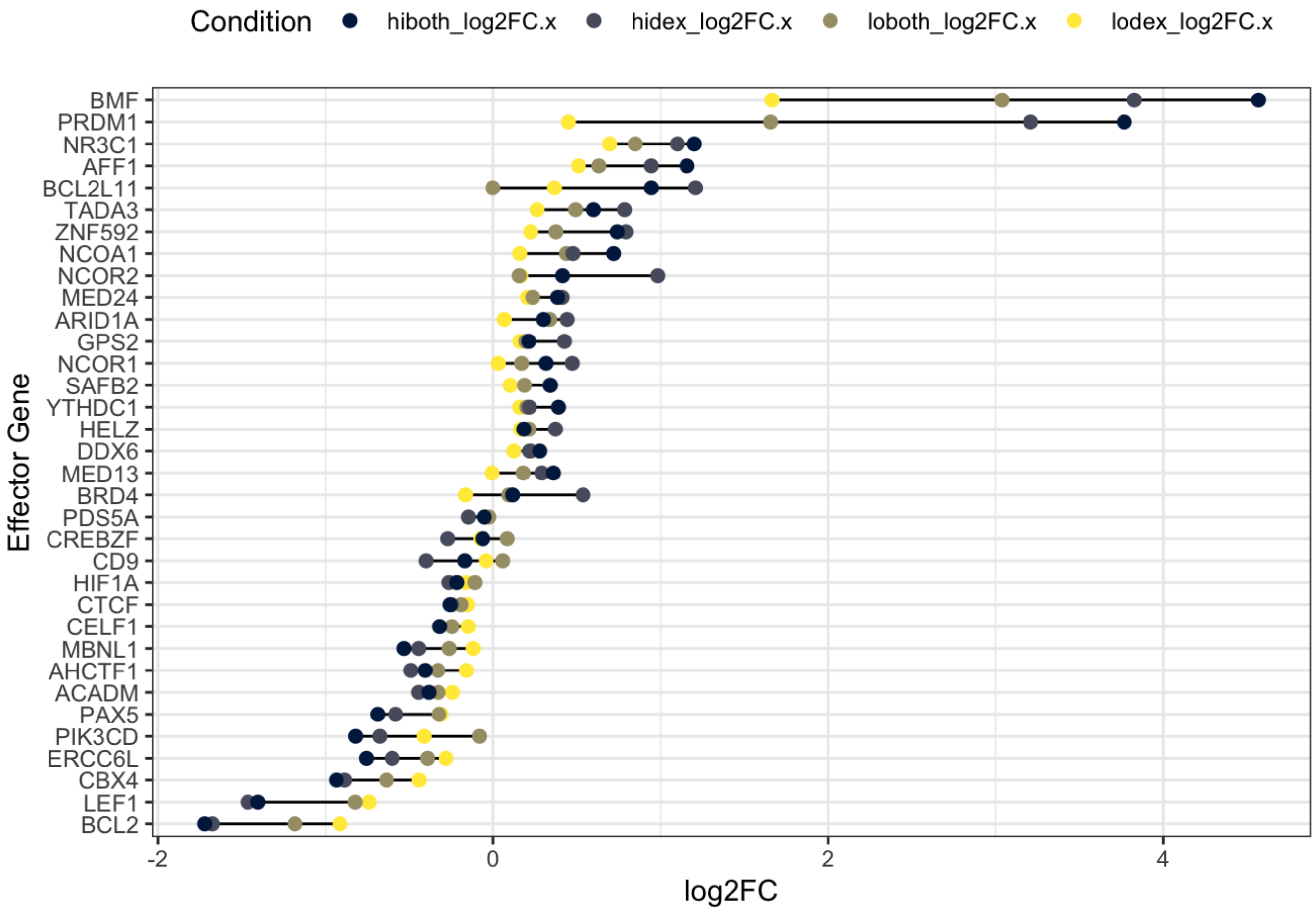

```
ggsave("eff_reg_cond.pdf", width = 7, height = 6)
```

for lodex

### Are effector genes more strongly regulated in response to idela

#### than other genes?

```
# ttest for enhanced upregulation by idela

test_up <- lodex_reg_filt %>%
  filter(loboth_log2FC > 0)

eff_1 <- ce_lng %>%
  dplyr::filter(Condition %in% c("lodex_log2FC.x", "loboth_log2FC.x")) %>%
  mutate(Condition = factor(Condition, levels = c("lodex_log2FC.x", "loboth_log2FC.x")))
%>%
  ggplot(aes(Condition, log2FC)) +
  ggtitle("Low dex, upregulated genes") + ylab("log2FC") + xlab("") +
  geom_boxplot() +
  theme(axis.text.x = element_text(angle = 45, hjust = 1)) +
  theme_bw()

eff_up <- ce_lng %>%
  dplyr::filter(Condition %in% c("lodex_log2FC.x", "loboth_log2FC.x") & log2FC > 0) %>%
  mutate(Condition = factor(Condition, levels = c("lodex_log2FC.x", "loboth_log2FC.x")))
%>%
  ggplot(aes(Condition, log2FC)) +
  ylab("log2FC") + xlab("") +
  geom_boxplot() +
  ylim(-1.5, 3.5) +
  theme_bw() +
  theme(axis.text.x = element_text(angle = 45, hjust = 1))

eff_down <- ce_lng %>%
  dplyr::filter(Condition %in% c("lodex_log2FC.x", "loboth_log2FC.x") & log2FC < 0) %>%
  mutate(Condition = factor(Condition, levels = c("lodex_log2FC.x", "loboth_log2FC.x")))
%>%
  ggplot(aes(Condition, log2FC)) +
  ylab("log2FC") + xlab("") +
  geom_boxplot() +
  ylim(-1.5, 3.5) +
  theme_bw() +
  theme(axis.text.x = element_text(angle = 45, hjust = 1))

box_eff_lowboth <- grid.arrange(eff_up, eff_down, nrow = 1)
```

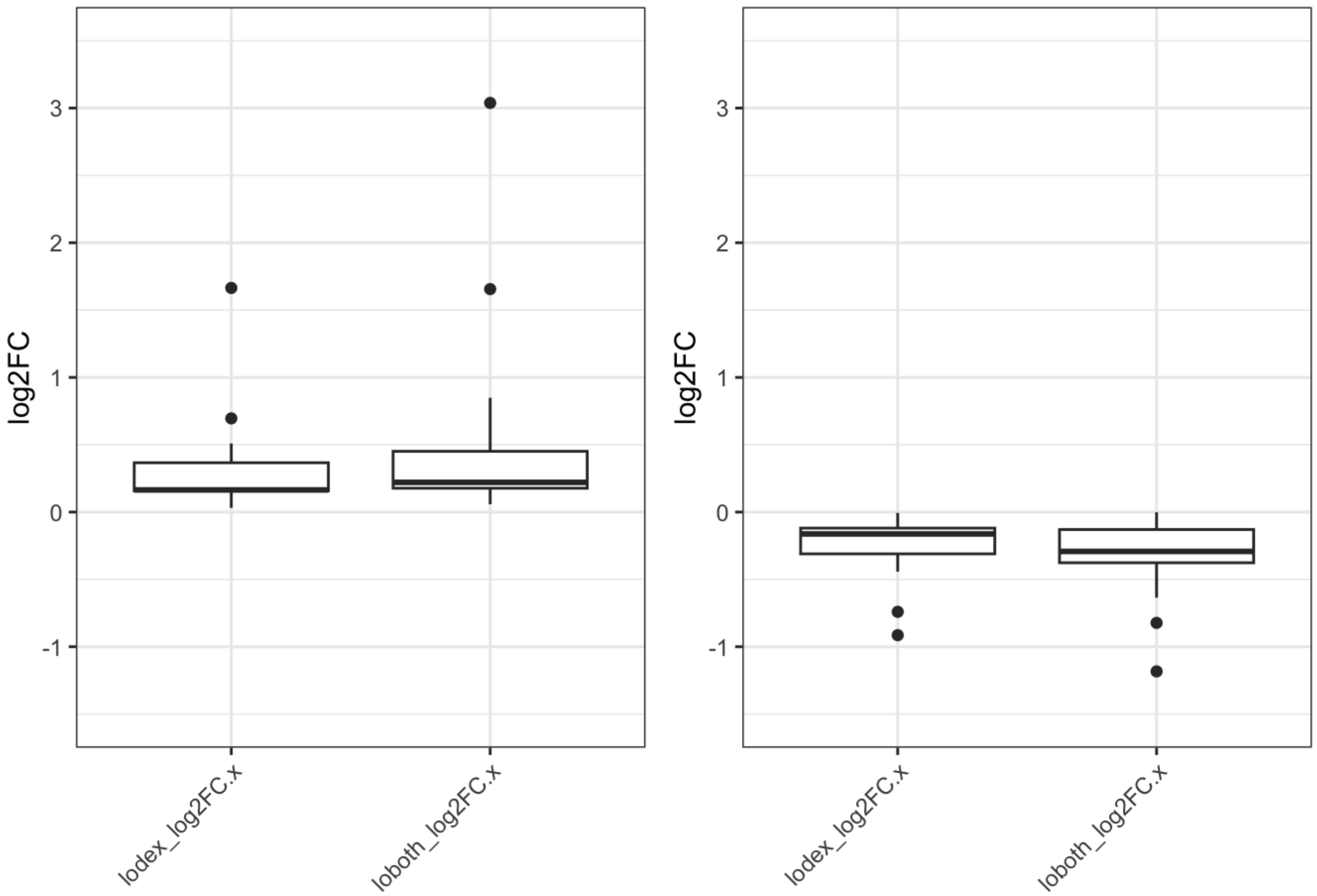

```
ggsave(filename = "boxplot_eff_dex_dex_lo.pdf", height = 4, width = 3, box_eff_lowboth)

sum_eff <- sum_table %>%
  dplyr::filter(symbol %in% ce_lng$symbol)

eff_up_table <- sum_eff %>%
  dplyr::filter(loboth_log2FC > 0 | lodex_log2FC > 0)

t.test(eff_up_table$loboth_log2FC, eff_up_table$lodex_log2FC, paired = TRUE)
```

```
##
##  Paired t-test
##
## data:  eff_up_table$loboth_log2FC and eff_up_table$lodex_log2FC
## t = 2.6259, df = 20, p-value = 0.01619
## alternative hypothesis: true mean difference is not equal to 0
## 95 percent confidence interval:
##  0.04500288 0.39270091
## sample estimates:
## mean difference
##       0.2188519
```

```r
eff_down_table <- sum_eff %>%
  dplyr::filter(loboth_log2FC < 0 | lodex_log2FC < 0)

t.test(eff_down_table$loboth_log2FC, eff_down_table$lodex_log2FC, paired = TRUE)
```

```
##
##  Paired t-test
##
## data:  eff_down_table$loboth_log2FC and eff_down_table$lodex_log2FC
## t = -0.58989, df = 17, p-value = 0.563
## alternative hypothesis: true mean difference is not equal to 0
## 95 percent confidence interval:
##  -0.11642160  0.06554521
## sample estimates:
## mean difference
##      -0.0254382
```

```r
t.test(abs(sum_eff$loboth_log2FC), abs(sum_eff$lodex_log2FC), paired = TRUE)
```

```
##
##  Paired t-test
##
## data:  abs(sum_eff$loboth_log2FC) and abs(sum_eff$lodex_log2FC)
## t = 2.5568, df = 33, p-value = 0.01535
## alternative hypothesis: true mean difference is not equal to 0
## 95 percent confidence interval:
##  0.02894599 0.25444377
## sample estimates:
## mean difference
##       0.1416949
```

```r
sum_eff %>%
  dplyr::filter(hiboth_log2FC > 0 | hidex_log2FC > 0) %>%
  t.test(.$hiboth_log2FC, .$hidex_log2FC, data = ., paired = TRUE)
```

```
##
##  Paired t-test
##
## data:  .$hiboth_log2FC and .$hidex_log2FC
## t = -0.037824, df = 18, p-value = 0.9702
## alternative hypothesis: true mean difference is not equal to 0
## 95 percent confidence interval:
##  -0.1527794  0.1473756
## sample estimates:
## mean difference
##    -0.002701932
```

```
sum_eff %>%
  dplyr::filter(hiboth_log2FC < 0 | hidex_log2FC < 0) %>%
  t.test(.$hiboth_log2FC, .$hidex_log2FC, data = ., paired = TRUE)
```

```
##
##  Paired t-test
##
## data:  .$hiboth_log2FC and .$hidex_log2FC
## t = 0.44287, df = 14, p-value = 0.6646
## alternative hypothesis: true mean difference is not equal to 0
## 95 percent confidence interval:
##  -0.05094853  0.07746414
## sample estimates:
## mean difference
##      0.01325781
```

```
t.test(abs(sum_eff$hiboth_log2FC), abs(sum_eff$hidex_log2FC), paired = TRUE)
```

```
##
##  Paired t-test
##
## data:  abs(sum_eff$hiboth_log2FC) and abs(sum_eff$hidex_log2FC)
## t = -0.17724, df = 33, p-value = 0.8604
## alternative hypothesis: true mean difference is not equal to 0
## 95 percent confidence interval:
##  -0.09183354  0.07711567
## sample estimates:
## mean difference
##    -0.007358935
```

```
ggplot(common_neg_eff, aes(lodex_log2FC.x, loboth_log2FC.x, label = symbol)) +
  labs(title = "Highest confidence negative effectors", y = "5 nM Dex + Idela, log2FC",
x = "5 nM Dex, log2FC") +
  geom_point() +
  geom_text_repel(size = 3, max.overlaps = 12, max.iter = 1000000) +
  geom_abline(slope = 1, intercept = 0, color = "red", size = 1) +
  xlim(-1.6, 0.1) + ylim(-1.6, 0.1) +
  theme_bw() +
  theme(axis.title = element_text(face="bold"), plot.title = element_text(face = "bol
d")) +
  annotate("rect", xmin = -1.6, xmax = 0.1, ymin = -1.6, ymax = 0.1 , fill= "green", alp
ha = 0.1)
```

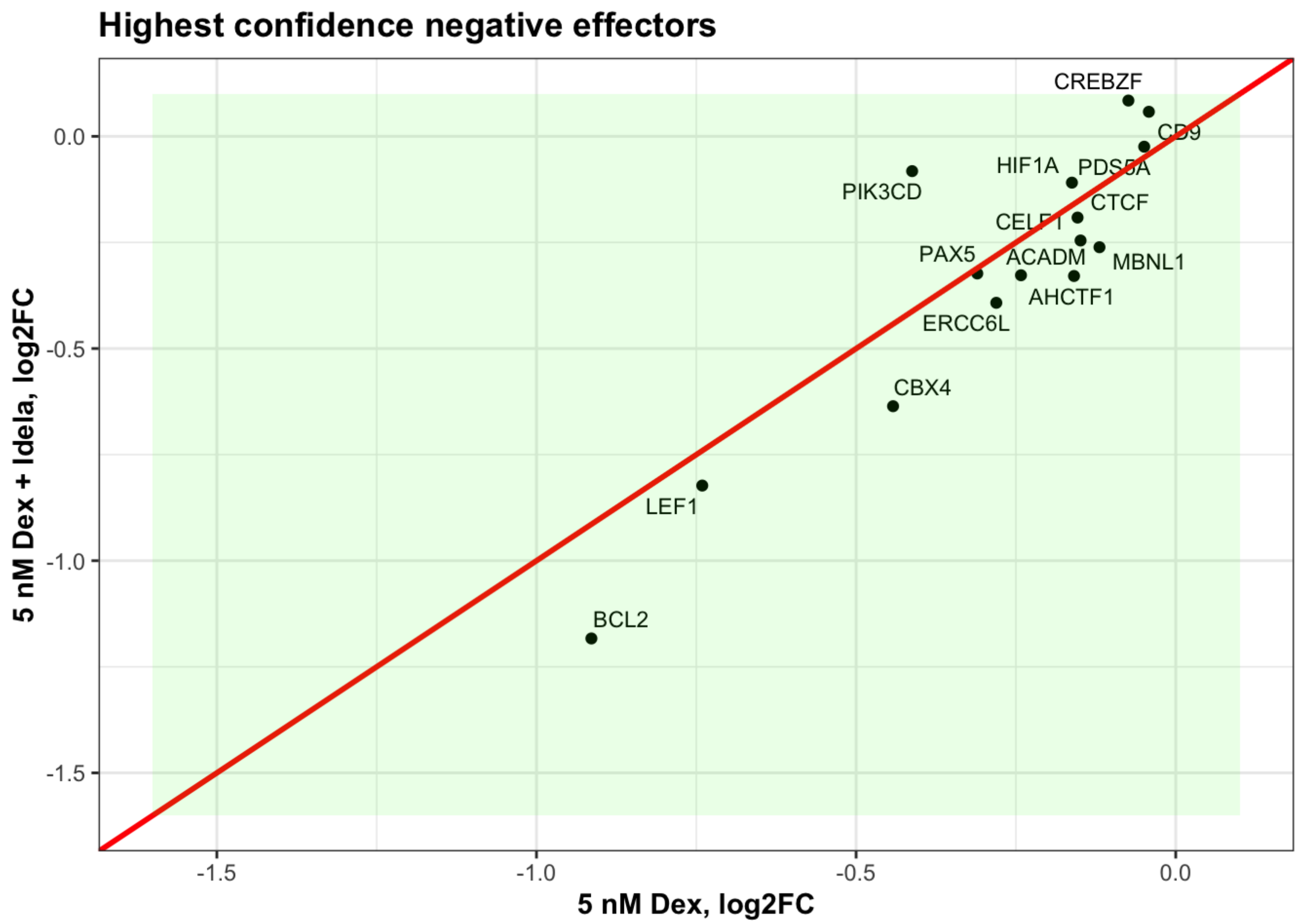

### Are any interaction genes effectors?

```
intersect(pos_cagek_effectors$symbol, res.50int_sig$symbol)
```

```
## [1] "GPS2" "SON"
```

```
intersect(pos_cagek_effectors$symbol, res.5int_sig$symbol)
```

```
## character(0)
```

```
intersect(pos_full_effectors$symbol, res.50int_sig$symbol)
```

```
## [1] "GPS2"   "MED13L"
```

```
intersect(pos_full_effectors$symbol, res.5int_sig$symbol)
```

```
## character(0)
```

```
intersect(neg_cagek_effectors$symbol, res.50int_sig$symbol)
```

```
## character(0)
```

```
intersect(neg_cagek_effectors$symbol, res.5int_sig$symbol)
```

```
## character(0)
```

```
intersect(neg_full_effectors$symbol, res.50int_sig$symbol)
```

```
## [1] "AKAP1" "MME"
```

```
intersect(neg_full_effectors$symbol, res.5int_sig$symbol)
```

```
## character(0)
```

3. Buffering Genes

```
full_buff <- full_effector_tbl %>%
  filter(Rho_pvalue < 0.05 & hidex_adjp < 0.05 & Rho_phenotype < 0 & hidex_log2FC > 0 |
Rho_pvalue < 0.05 & hidex_adjp < 0.05 & Rho_phenotype > 0 & hidex_log2FC < 0)

cagek_buff <- cagek_effector_tbl %>%
  filter(CAGEK_Rho_pvalue < 0.05 & hidex_adjp < 0.05 & CAGEK_Rho_phenotype < 0 & hidex_l
og2FC > 0 | CAGEK_Rho_pvalue < 0.05 & hidex_adjp < 0.05 & CAGEK_Rho_phenotype > 0 & hide
x_log2FC < 0)

all_buff <- full_buff %>%
  full_join(cagek_buff, by = "symbol")

common_buff <- full_buff %>%
  inner_join(cagek_buff, by = "symbol")

ggplot(common_buff, aes(hidex_log2FC.x, hiboth_log2FC.x, label = symbol)) +
  ggtitle("Highest confidence buffering genes") + ylab("50 nM Dex + Idela, log2FC") + xl
ab("50 nM Dex, log2FC") +
  geom_point() +
  geom_text_repel(size = 3, max.overlaps = 12, max.iter = 1000000) +
  geom_abline(slope = 1, intercept = 0, color = "red", size = 1) +
  xlim(-1, 1) + ylim(-1, 1) +
  geom_hline(yintercept = 0) + geom_vline(xintercept = 0) +
  theme_bw() +
  theme(axis.title = element_text(face="bold"), plot.title = element_text(face = "bol
d")) +
  annotate("rect", xmin = -Inf, xmax = 0, ymin = -Inf, ymax = 0, fill= "yellow", alpha =
0.1) +
  annotate("rect", xmin = 0, xmax = Inf, ymin = 0, ymax = Inf, fill= "yellow", alpha =
0.1)
```

```
## Warning: Removed 1 rows containing missing values (`geom_point()`).
```

```
## Warning: Removed 1 rows containing missing values (`geom_text_repel()`).
```

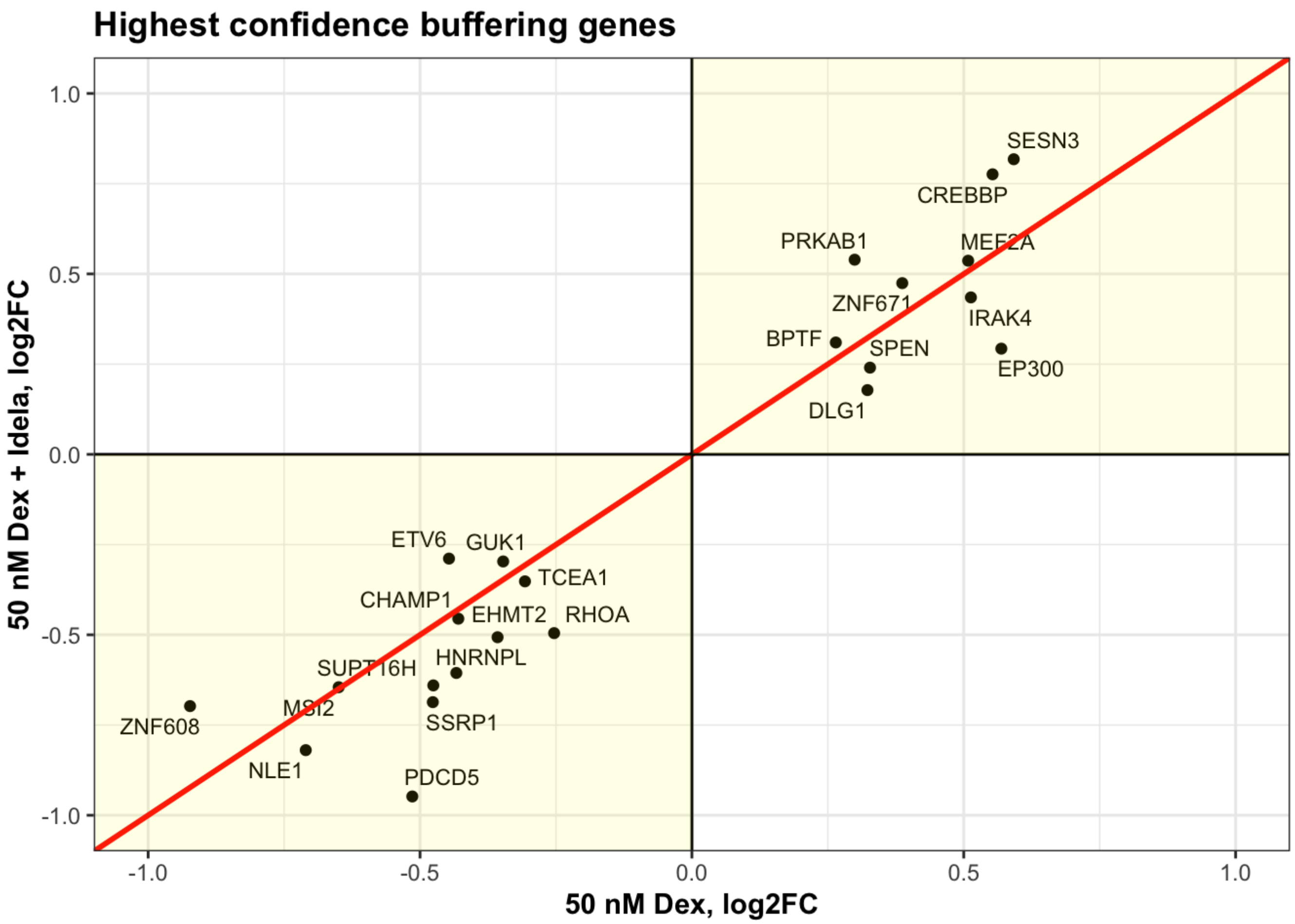

```
ggsave("common_buff_genes.pdf", width = 6, height= 6)
```

```
## Warning: Removed 1 rows containing missing values (`geom_point()`).
## Removed 1 rows containing missing values (`geom_text_repel()`).
```

### What are the dex + idela effectors

1. Positive effectors

```
#Genes that are upregulated with 50 nM Dex and Idela that are significant contributors t
o dex-induced cell death in both screen versions

di_pos_full_effectors <- full_effector_tbl %>%
  filter(Rho_pvalue < 0.05 & hiboth_adjp < 0.05 & Rho_phenotype > 0 & hiboth_log2FC > 0)

di_pos_cagek_effectors <- cagek_effector_tbl %>%
  filter(CAGEK_Rho_pvalue < 0.05 & hiboth_adjp < 0.05 & CAGEK_Rho_phenotype > 0 & hiboth
_log2FC > 0)

# The positive effectors in either of the screens

di_all_pos_eff <- di_pos_full_effectors %>%
  full_join(di_pos_cagek_effectors, by = "symbol")

# Positive effectors that were positive in both screens

di_common_pos_eff <- di_pos_full_effectors %>%
  inner_join(di_pos_cagek_effectors, by = "symbol")

intersect(di_common_pos_eff$symbol, common_pos_eff$symbol)
```

```
##  [1] "AFF1"    "ARID1A"  "BCL2L11" "BMF"     "DDX6"    "MED13"   "MED24"
##  [8] "NCOA1"   "NCOR1"   "NCOR2"   "NR3C1"   "PRDM1"   "SAFB2"   "TADA3"
## [15] "YTHDC1"  "ZNF592"
```

```
di_specific_effectors <- setdiff(di_common_pos_eff$symbol, common_pos_eff$symbol)
dex_specific_effectors <- setdiff(common_pos_eff$symbol, di_common_pos_eff$symbol)

## Which are gained w/ addition of idela?

#di_specific <- setdiff(di_all_pos_eff$symbol, di_common)
#dex_specific <- setdiff(all_pos_eff$symbol, di_common)

full_effector_tbl %>%
  filter(symbol %in% di_specific_effectors) %>%
  ggplot(aes(hidex_log2FC, hiboth_log2FC, label = symbol)) +
  labs(title = "Expression of Idela enhanced hidex effector genes") +
  geom_point() +
  geom_text_repel(size = 3, max.overlaps = 12, max.iter = 1000000, label.padding = 0.1)
+
  geom_abline(slope = 1, intercept = 0, color = "red", size = 1) +
  xlim(-0.1,1) + ylim(-0.1, 1) +
  theme_bw()
```

```
## Warning in geom_text_repel(size = 3, max.overlaps = 12, max.iter = 1e+06, :
## Ignoring unknown parameters: `label.padding`
```

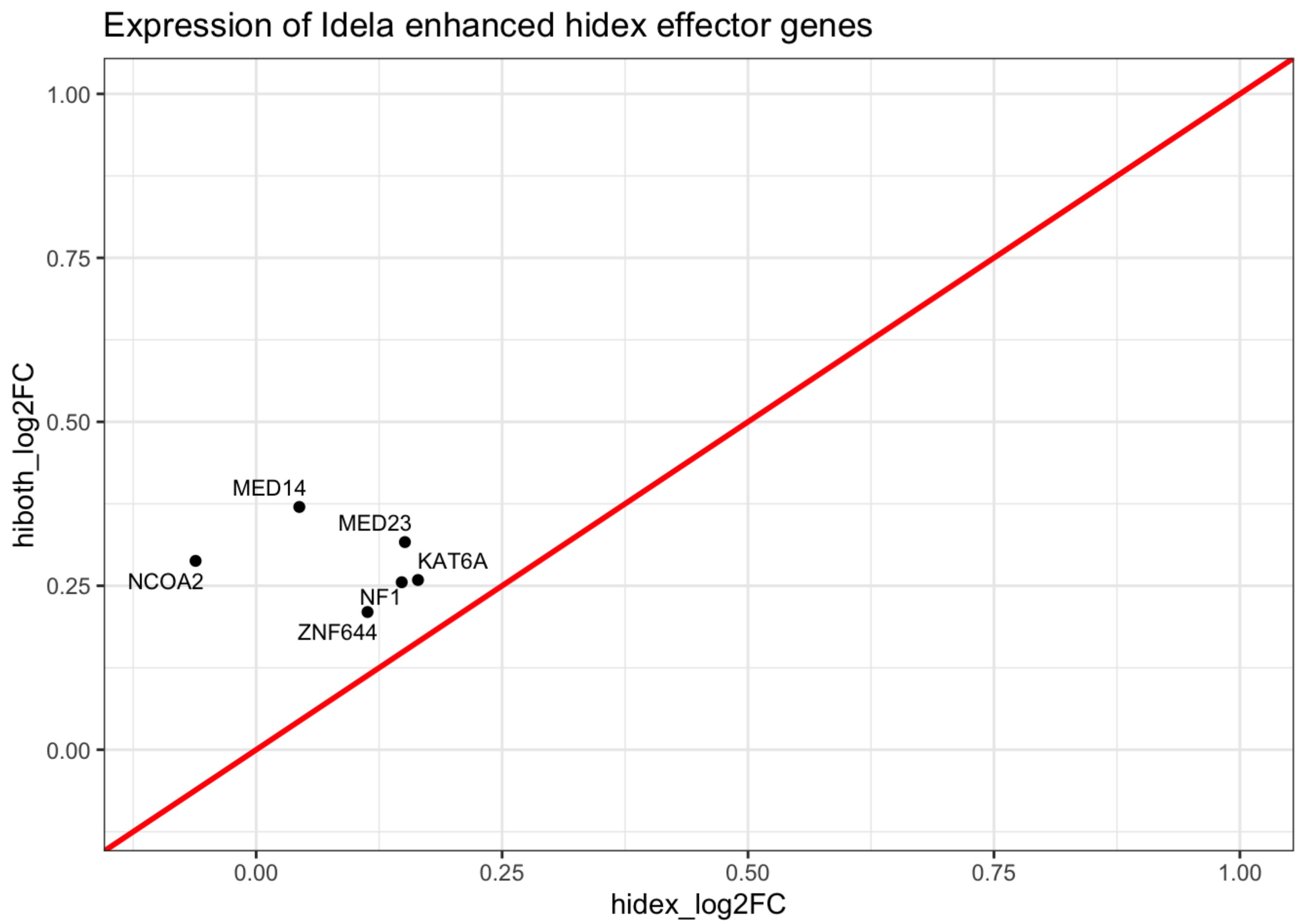

```
#dex_specific_data <- all_pos_eff %>%
#  filter(symbol %in% dex_specific)

full_effector_tbl %>%
  filter(symbol %in% dex_specific_effectors) %>%
  ggplot(aes(hidex_log2FC, hiboth_log2FC, label = symbol)) +
  labs(title = "Expression of Idela diminished hidex effector genes") +
  geom_point() +
  geom_text_repel(size = 3, max.overlaps = 12, max.iter = 1000000, label.padding = 0.1)
+
  geom_abline(slope = 1, intercept = 0, color = "red", size = 1) +
  xlim(-0.1,1) + ylim(-0.1, 1) +
  theme_bw()
```

```
## Warning in geom_text_repel(size = 3, max.overlaps = 12, max.iter = 1e+06, :
## Ignoring unknown parameters: `label.padding`
```

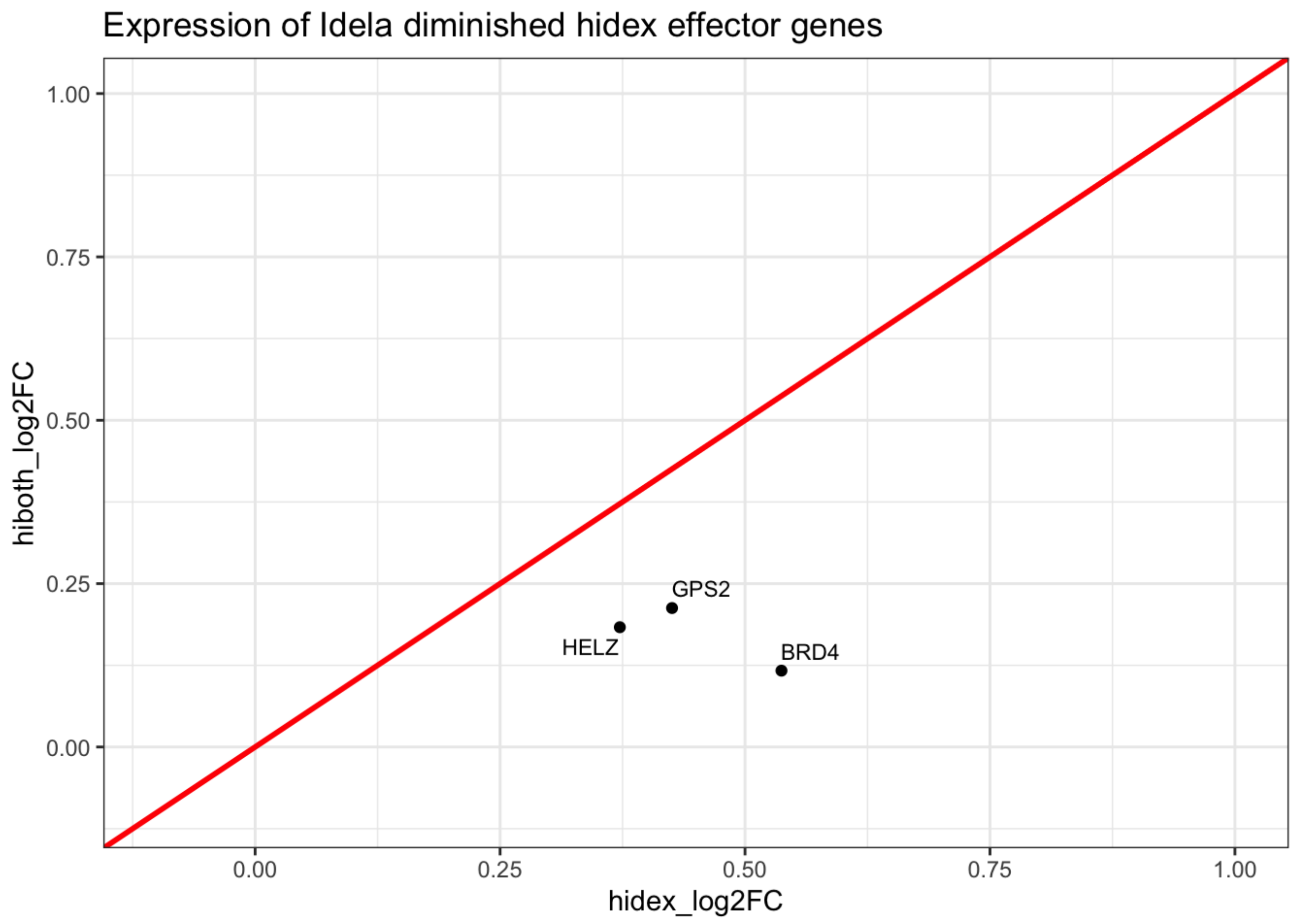

```
#full_effector_tbl %>%
#  filter(symbol %in% di_common) %>%
#  ggplot(aes(hidex_log2FC, hiboth_log2FC, label = symbol)) +
  # labs(title = "Effector genes with HiDex +/- Idela") +
  # geom_point() +
  # geom_text_repel(size = 3, max.overlaps = 12, max.iter = 1000000, label.padding = 0.
1) +
  # geom_abline(slope = 1, intercept = 0, color = "red", size = 1) +
  # xlim(-0.25,6) + ylim(-0.25, 6) +
  # theme_bw()
```

```
#Filter for significant genes

hidex_full_eff_loose <- full_effector_tbl %>%
  filter(Rho_pvalue < 0.05 & hidex_adjp < 0.05 & abs(hidex_log2FC) < 10) %>%
  mutate(quadrant = case_when(hidex_log2FC > 0 & Rho_phenotype > 0 ~ "darkorchid4",
                              hidex_log2FC < 0 & Rho_phenotype > 0 ~ "darkorange2",
                              hidex_log2FC < 0 & Rho_phenotype < 0 ~ "forestgreen",
                              hidex_log2FC > 0 & Rho_phenotype < 0 ~ "darkorange2"))
## positive effectors
hidex_full_eff_loose %>%
  filter(hidex_log2FC > 0 & Rho_phenotype > 0) %>%
  count()
```

```
## # A tibble: 1 × 1
##       n
##   <int>
## 1    85
```

```
## negative effectors
hidex_full_eff_loose %>%
  filter(hidex_log2FC < 0 & Rho_phenotype < 0) %>%
  count()
```

```
## # A tibble: 1 × 1
##       n
##   <int>
## 1   183
```

```
#Buffering genes

hidex_full_eff_loose %>%
  filter(hidex_log2FC > 0 & Rho_phenotype < 0 | hidex_log2FC < 0 & Rho_phenotype > 0) %
>%
  count()
```

```
## # A tibble: 1 × 1
##       n
##   <int>
## 1   229
```

```r
hidex_full_eff <- full_effector_tbl %>%
  filter(Rho_pvalue < 0.01 & hidex_adjp < 0.01 & abs(hidex_log2FC) < 10) %>%
  mutate(quadrant = case_when(hidex_log2FC > 0 & Rho_phenotype > 0 ~ "darkorchid4",
                              hidex_log2FC < 0 & Rho_phenotype > 0 ~ "darkorange2",
                              hidex_log2FC < 0 & Rho_phenotype < 0 ~ "forestgreen",
                              hidex_log2FC > 0 & Rho_phenotype < 0 ~ "darkorange2"))

ggplot(hidex_full_eff, aes(Rho_phenotype, hidex_log2FC, label = symbol)) +
  geom_point(colour = hidex_full_eff$quadrant) +
  geom_hline(yintercept = 0) + geom_vline(xintercept = 0) +
  geom_text_repel(size = 3, max.overlaps = 12, force = 2) +
  labs(title = "50 nM Dex Effectors, Full Screen", y = "50 nM Dex, log2FC", x = "Contrib
ution to Dex Sensitivity") +
  theme_bw() +
  theme(axis.title = element_text(face="bold")) +
  annotate("rect", xmin = Inf, xmax = 0, ymin = Inf, ymax = 0, fill= "purple", alpha =
0.1)  +
  annotate("rect", xmin = -Inf, xmax = 0, ymin = -Inf, ymax = 0 , fill= "green", alpha =
0.1) +
  annotate("rect", xmin = 0, xmax = Inf, ymin = 0, ymax = -Inf, fill= "yellow", alpha =
0.1) +
  annotate("rect", xmin = 0, xmax = -Inf, ymin = Inf, ymax = 0, fill= "yellow", alpha =
0.1)
```

```
## Warning: ggrepel: 120 unlabeled data points (too many overlaps). Consider
## increasing max.overlaps
```

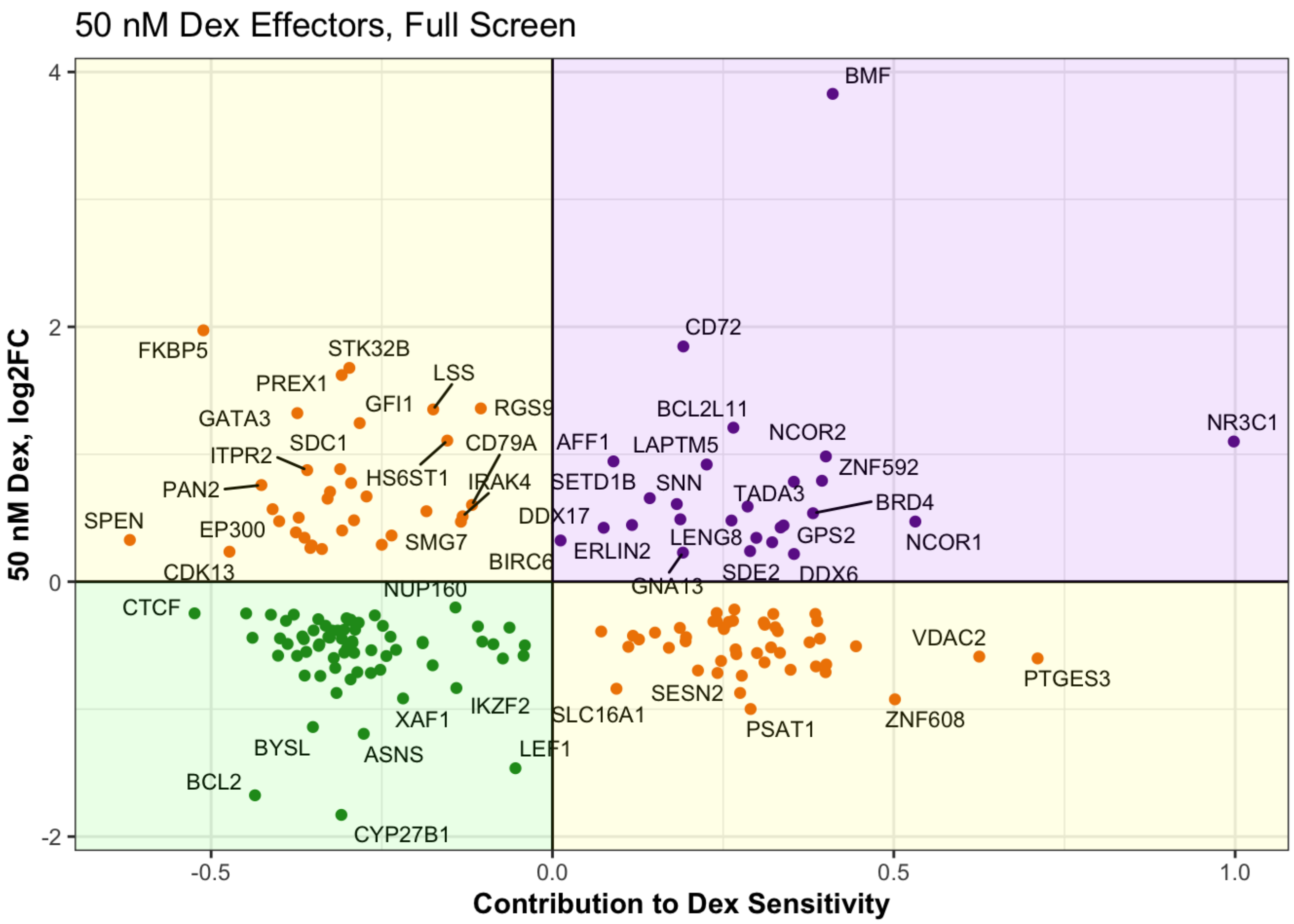

```
ggsave("hidex_full_effector_plot.pdf", width = 8, height = 8)
```

```
## Warning: ggrepel: 83 unlabeled data points (too many overlaps). Consider
## increasing max.overlaps
```

```
ggplot(hidex_full_eff, aes(Rho_phenotype, hidex_log2FC, label = symbol)) +
  geom_point(colour = hidex_full_eff$quadrant) +
  geom_hline(yintercept = 0) + geom_vline(xintercept = 0) +
  xlim(-0.55, 0) + ylim(-1.9,0) +
  geom_text_repel(size = 3, max.overlaps = 12, force = 2) +
  labs(title = "50 nM Negative Dex Effectors, Full Screen", y = "50 nM Dex, log2FC", x =
"Contribution to Dex Sensitivity") +
  theme_bw() +
  theme(axis.title = element_text(face="bold")) +
  annotate("rect", xmin = Inf, xmax = 0, ymin = Inf, ymax = 0, fill= "purple", alpha =
0.1)  +
  annotate("rect", xmin = -Inf, xmax = 0, ymin = -Inf, ymax = 0 , fill= "green", alpha =
0.1) +
  annotate("rect", xmin = 0, xmax = Inf, ymin = 0, ymax = -Inf, fill= "yellow", alpha =
0.1) +
  annotate("rect", xmin = 0, xmax = -Inf, ymin = Inf, ymax = 0, fill= "yellow", alpha =
0.1)
```

```
## Warning: Removed 106 rows containing missing values (`geom_point()`).
```

```
## Warning: Removed 106 rows containing missing values (`geom_text_repel()`).
```

```
## Warning: ggrepel: 18 unlabeled data points (too many overlaps). Consider
## increasing max.overlaps
```

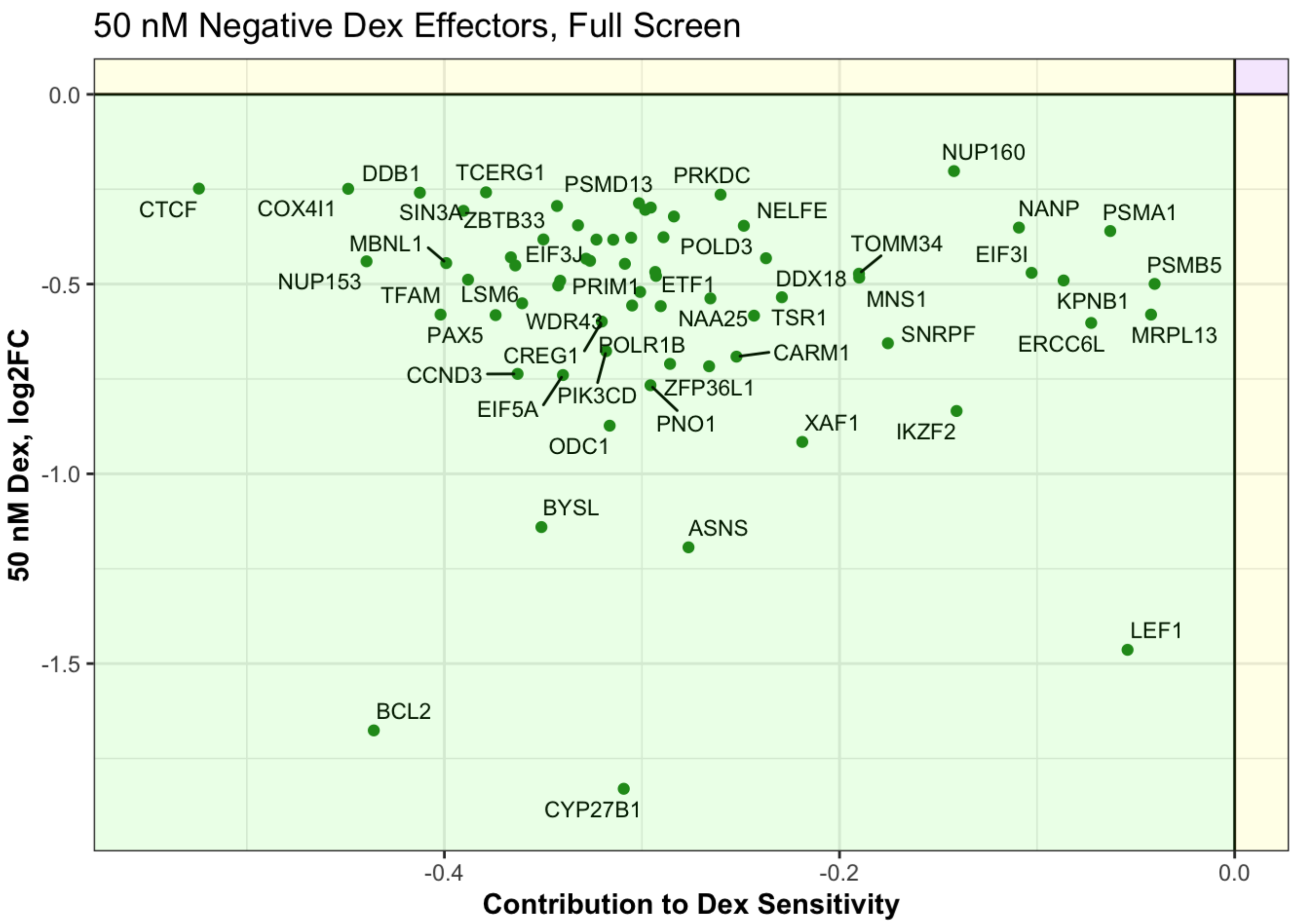

```r
ggsave("hidex_full_negeffector_plot.pdf", width = 8, height = 8)
```

```
## Warning: Removed 106 rows containing missing values (`geom_point()`).
```

```
## Warning: Removed 106 rows containing missing values (`geom_text_repel()`).
```

```
ggplot(hidex_full_eff, aes(Rho_phenotype, hidex_log2FC, label = symbol)) +
  geom_point(colour = hidex_full_eff$quadrant) +
  geom_hline(yintercept = 0) + geom_vline(xintercept = 0) +
  xlim(0, 0.6) + ylim(0,2) +
  geom_text_repel(size = 3, max.overlaps = 12, max.iter = 1000000, label.padding = 0.1)
+
  labs(title = "50 nM Positive Dex Effectors, Full Screen", y = "50 nM Dex, log2FC", x =
"Contribution to Dex Sensitivity") +
  theme_bw() +
  theme(axis.title = element_text(face="bold")) +
  annotate("rect", xmin = Inf, xmax = 0, ymin = Inf, ymax = 0, fill= "purple", alpha =
0.1)  +
  annotate("rect", xmin = -Inf, xmax = 0, ymin = -Inf, ymax = 0 , fill= "green", alpha =
0.1) +
  annotate("rect", xmin = 0, xmax = Inf, ymin = 0, ymax = -Inf, fill= "yellow", alpha =
0.1) +
  annotate("rect", xmin = 0, xmax = -Inf, ymin = Inf, ymax = 0, fill= "yellow", alpha =
0.1)
```

```
## Warning in geom_text_repel(size = 3, max.overlaps = 12, max.iter = 1e+06, :
## Ignoring unknown parameters: `label.padding`
```

```
## Warning: Removed 149 rows containing missing values (`geom_point()`).
```

```
## Warning: Removed 149 rows containing missing values (`geom_text_repel()`).
```

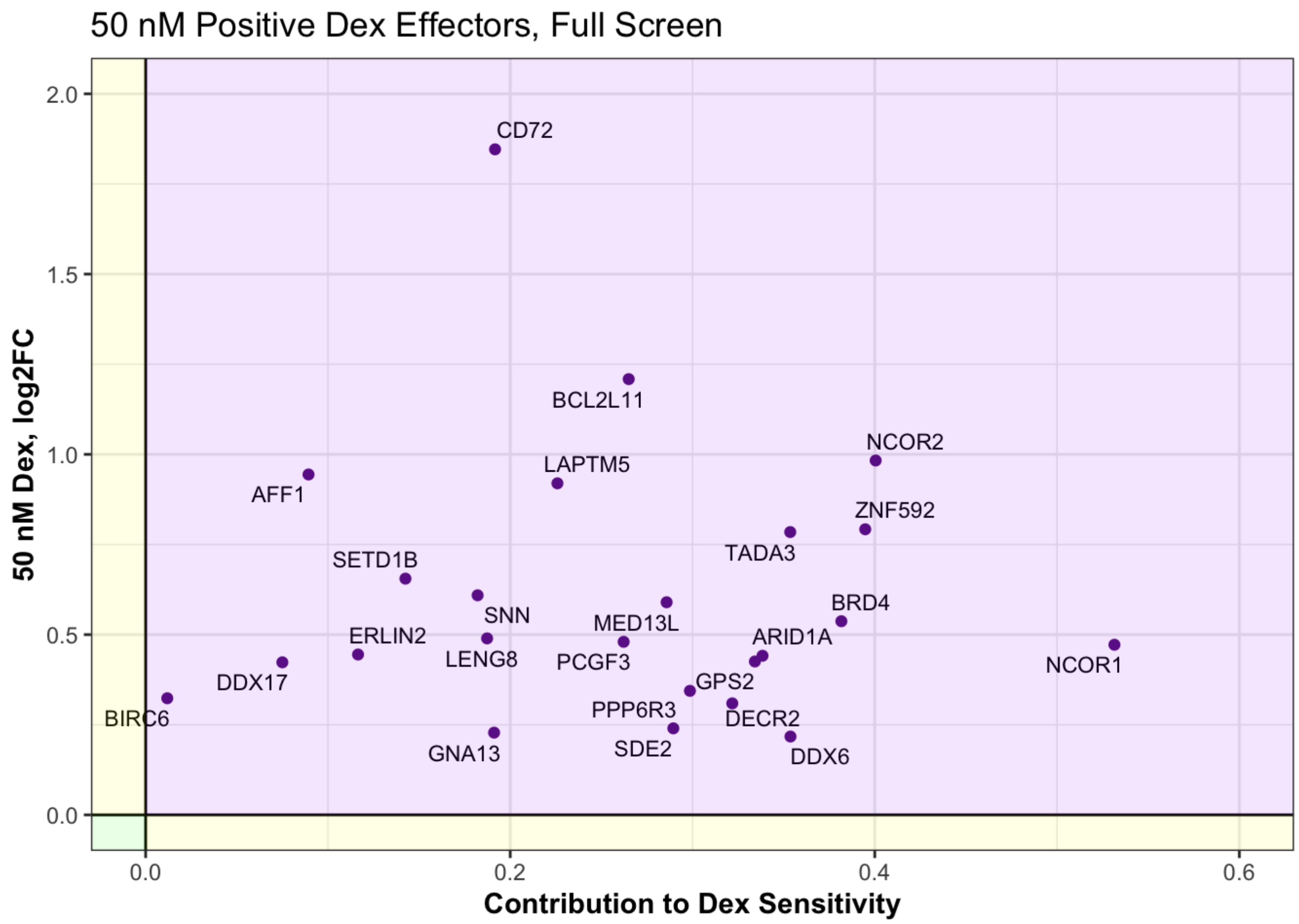

```
ggsave("hidex_full_poseffector_plot.pdf", width = 8, height = 8)
```

```
## Warning: Removed 149 rows containing missing values (`geom_point()`).
## Removed 149 rows containing missing values (`geom_text_repel()`).
```

```
hidex_cagek_eff <- cagek_effector_tbl %>%
  filter(CAGEK_Rho_pvalue < 0.01 & hidex_adjp < 0.01 & abs(hidex_log2FC) < 10) %>%
  mutate(quadrant = case_when(hidex_log2FC > 0 & CAGEK_Rho_phenotype > 0 ~ "darkorchid
4",
                              hidex_log2FC < 0 & CAGEK_Rho_phenotype > 0 ~ "darkorange
2",
                              hidex_log2FC < 0 & CAGEK_Rho_phenotype < 0 ~ "forestgree
n",
                              hidex_log2FC > 0 & CAGEK_Rho_phenotype < 0 ~ "darkorange
2"))

ggplot(hidex_cagek_eff, aes(CAGEK_Rho_phenotype, hidex_log2FC, label = symbol)) +
  geom_point(colour = hidex_cagek_eff$quadrant) +
  geom_hline(yintercept = 0) + geom_vline(xintercept = 0) +
  geom_text_repel(size = 3, max.overlaps = 12, max.iter = 1000000, label.padding = 0.1)
+
  labs(title = "50 nM Dex Effectors, CAGEK", y = "50 nM Dex, log2FC", x = "Contribution
to Dex Sensitivity") +
  theme_bw() +
  theme(axis.title = element_text(face="bold")) +
  annotate("rect", xmin = Inf, xmax = 0, ymin = Inf, ymax = 0, fill= "purple", alpha =
0.1)  +
  annotate("rect", xmin = -Inf, xmax = 0, ymin = -Inf, ymax = 0 , fill= "green", alpha =
0.1) +
  annotate("rect", xmin = 0, xmax = Inf, ymin = 0, ymax = -Inf, fill= "yellow", alpha =
0.1) +
  annotate("rect", xmin = 0, xmax = -Inf, ymin = Inf, ymax = 0, fill= "yellow", alpha =
0.1)
```

```
## Warning in geom_text_repel(size = 3, max.overlaps = 12, max.iter = 1e+06, :
## Ignoring unknown parameters: `label.padding`
```

```
## Warning: ggrepel: 59 unlabeled data points (too many overlaps). Consider
## increasing max.overlaps
```

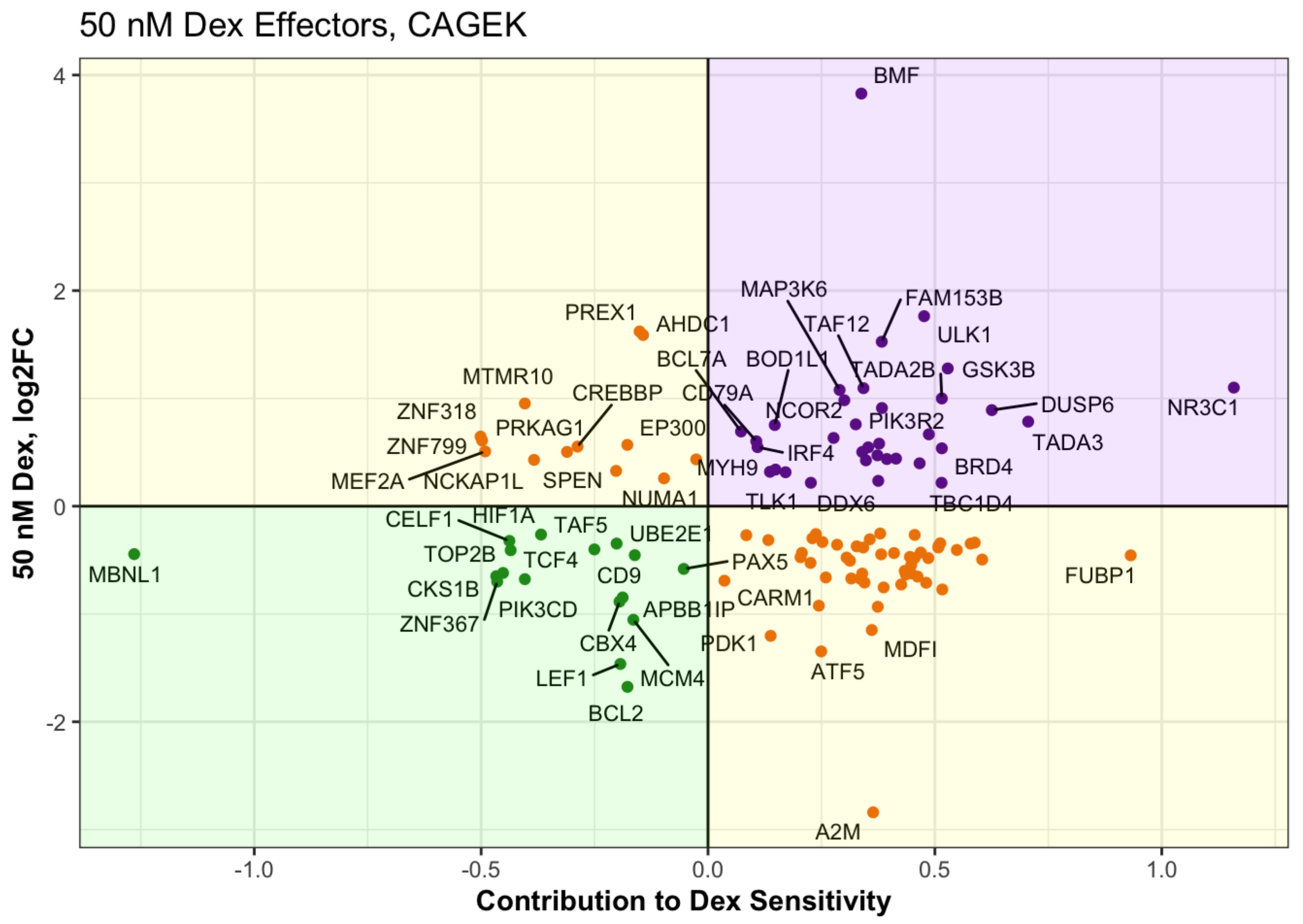

```
ggsave("hidex_cagek_effector_plot.pdf", width = 10, height = 10)
```

```
## Warning: ggrepel: 36 unlabeled data points (too many overlaps). Consider
## increasing max.overlaps
```

```
ggplot(hidex_cagek_eff, aes(CAGEK_Rho_phenotype, hidex_log2FC, label = symbol)) +
  geom_point(colour = hidex_cagek_eff$quadrant) +
  geom_hline(yintercept = 0) + geom_vline(xintercept = 0) +
  xlim(-1.3, 0) + ylim(-1.6,0) +
  geom_text_repel(size = 3, max.overlaps = 12, max.iter = 1000000, label.padding = 0.1)
+
  labs(title = "50 nM Dex Effectors, CAGEK", y = "50 nM Dex, log2FC", x = "Contribution
to Dex Sensitivity") +
  theme_bw() +
  theme(axis.title = element_text(face="bold")) +
  annotate("rect", xmin = Inf, xmax = 0, ymin = Inf, ymax = 0, fill= "purple", alpha =
0.1)  +
  annotate("rect", xmin = -Inf, xmax = 0, ymin = -Inf, ymax = 0 , fill= "green", alpha =
0.1) +
  annotate("rect", xmin = 0, xmax = Inf, ymin = 0, ymax = -Inf, fill= "yellow", alpha =
0.1) +
  annotate("rect", xmin = 0, xmax = -Inf, ymin = Inf, ymax = 0, fill= "yellow", alpha =
0.1)
```

```
## Warning in geom_text_repel(size = 3, max.overlaps = 12, max.iter = 1e+06, :
## Ignoring unknown parameters: `label.padding`
```

```
## Warning: Removed 99 rows containing missing values (`geom_point()`).
```

```
## Warning: Removed 99 rows containing missing values (`geom_text_repel()`).
```

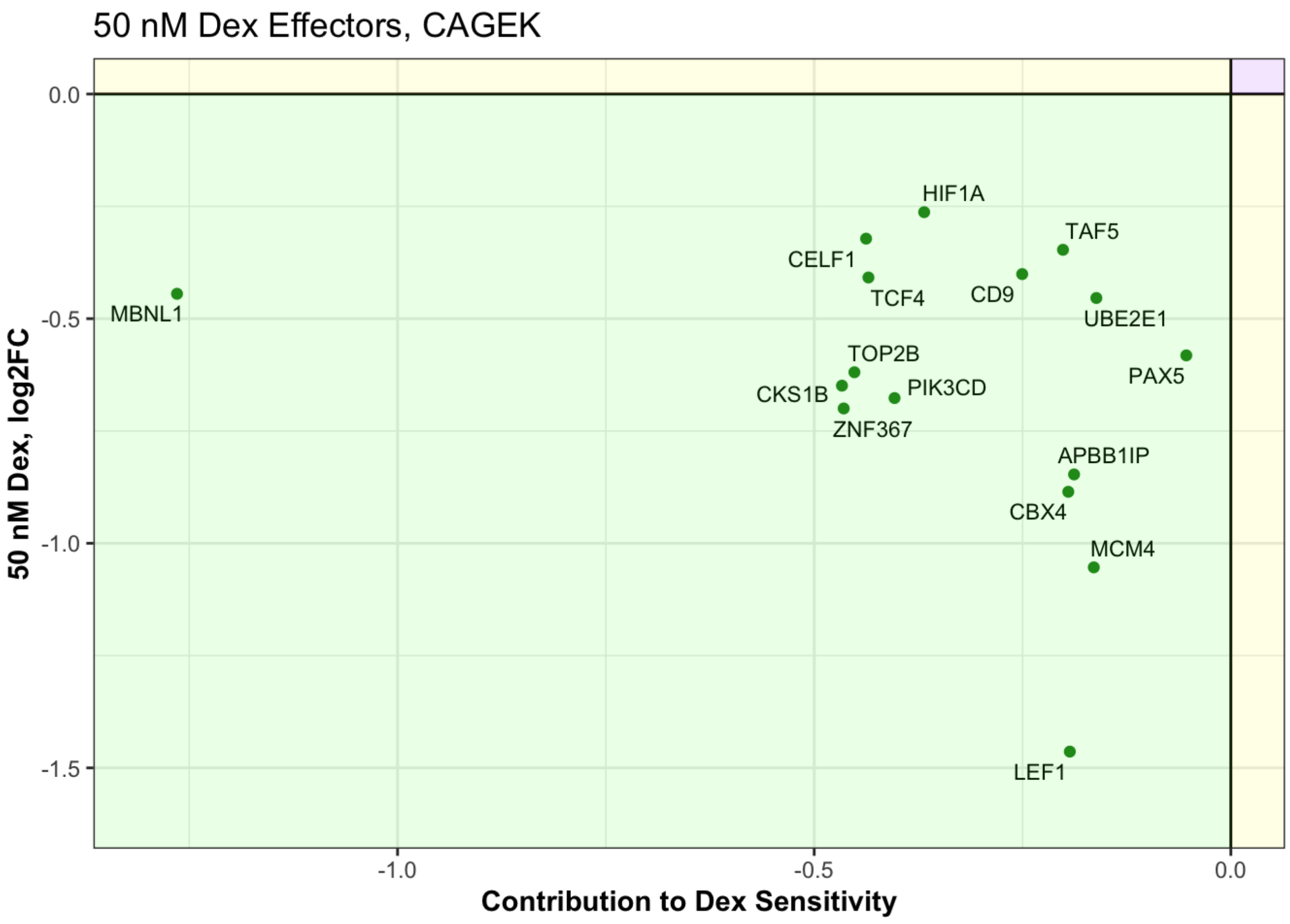

```
ggsave("hidex_cagek_negeffector_plot.pdf", width = 10, height = 10)
```

```
## Warning: Removed 99 rows containing missing values (`geom_point()`).
## Removed 99 rows containing missing values (`geom_text_repel()`).
```

```
ggplot(hidex_cagek_eff, aes(CAGEK_Rho_phenotype, hidex_log2FC, label = symbol)) +
  geom_point(colour = hidex_cagek_eff$quadrant) +
  geom_hline(yintercept = 0) + geom_vline(xintercept = 0) +
  xlim(0, 0.8) + ylim(0,2.5) +
  geom_text_repel(size = 3, max.overlaps = 12, max.iter = 1000000, label.padding = 0.1)
+
  labs(title = "50 nM Dex Effectors, CAGEK", y = "50 nM Dex, log2FC", x = "Contribution
to Dex Sensitivity") +
  theme_bw() +
  theme(axis.title = element_text(face="bold")) +
  annotate("rect", xmin = Inf, xmax = 0, ymin = Inf, ymax = 0, fill= "purple", alpha =
0.1)  +
  annotate("rect", xmin = -Inf, xmax = 0, ymin = -Inf, ymax = 0 , fill= "green", alpha =
0.1) +
  annotate("rect", xmin = 0, xmax = Inf, ymin = 0, ymax = -Inf, fill= "yellow", alpha =
0.1) +
  annotate("rect", xmin = 0, xmax = -Inf, ymin = Inf, ymax = 0, fill= "yellow", alpha =
0.1)
```

```
## Warning in geom_text_repel(size = 3, max.overlaps = 12, max.iter = 1e+06, :
## Ignoring unknown parameters: `label.padding`
```

```
## Warning: Removed 83 rows containing missing values (`geom_point()`).
```

```
## Warning: Removed 83 rows containing missing values (`geom_text_repel()`).
```

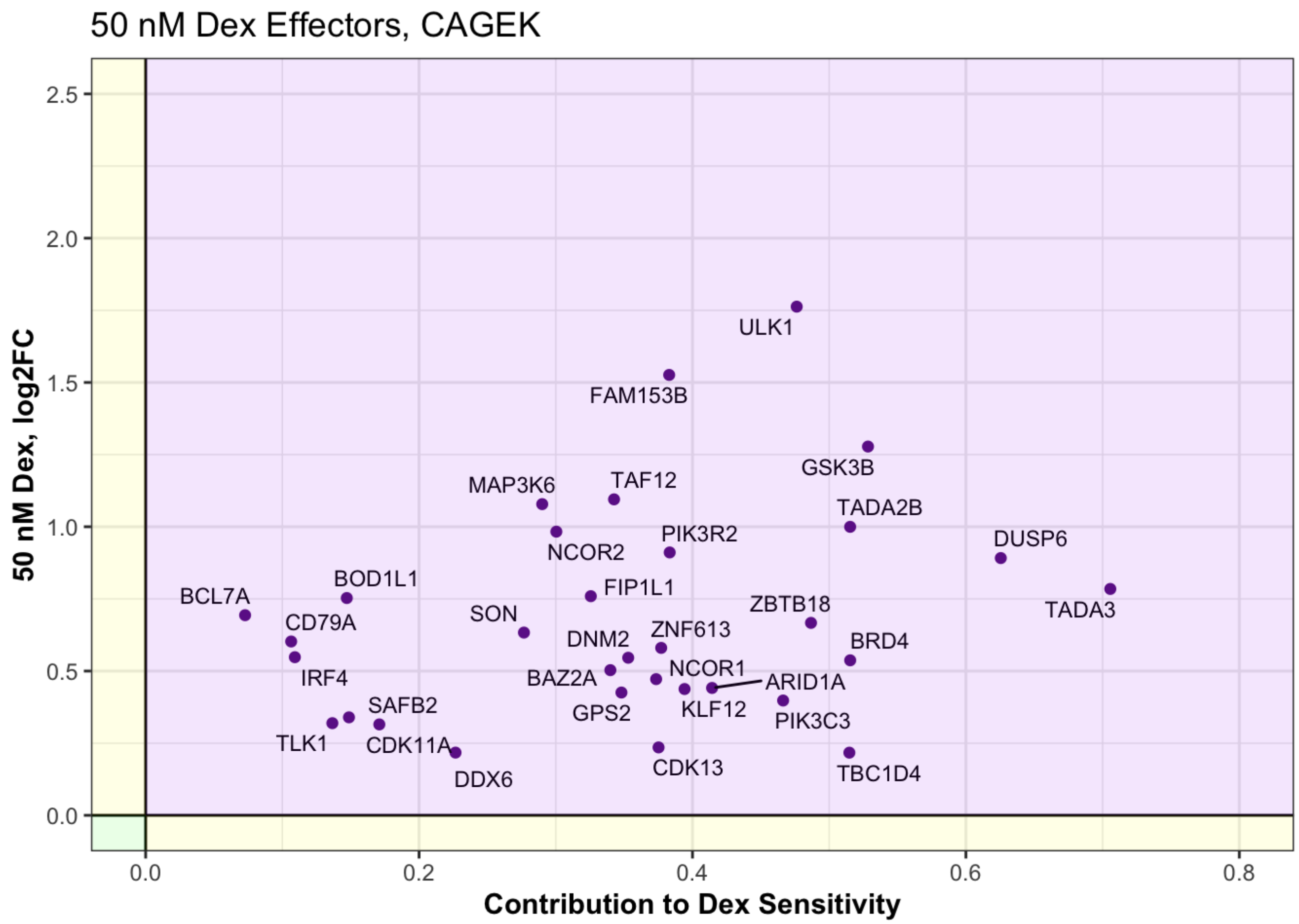

```
ggsave("hidex_cagek_poseffector_plot.pdf", width = 10, height = 10)
```

```
## Warning: Removed 83 rows containing missing values (`geom_point()`).
## Removed 83 rows containing missing values (`geom_text_repel()`).
```

### But are any of these affecting growth or GC senstivity?

Sensitivity from full screen

```
lo_int_full_rhos <- full_rhos %>%
  filter(symbol %in% int_genes) %>%
  filter(Rho_pvalue <= 0.05)
nrow(lo_int_full_rhos)
```

```
## [1] 1
```

Growth from full screen

```
lo_int_full_gammas <- full_gammas %>%
  filter(symbol %in% int_genes) %>%
  filter(Gamma_qvalue <= 0.05)
nrow(lo_int_full_gammas)
```

```
## [1] 0
```

Sensitivity from CAGEK screen

```
cagek_rhos <- cagek_rhos[, c(1:7)]

lo_int_cagek_rhos <- cagek_rhos %>%
  filter(symbol %in% int_genes) %>%
  filter(CAGEK_Rho_pvalue <= 0.05)
nrow(lo_int_cagek_rhos)
```

```
## [1] 0
```

Growth from CAGEK screen

```
lo_int_cagek_gammas <- cagek_gammas %>%
  filter(symbol %in% int_genes) %>%
  filter(CAGEK_Gamma_pvalue <= 0.05)
nrow(lo_int_cagek_gammas)
```

```
## [1] 0
```

Conclusion: Enhanced cell death is from additive regulation of effector genes, not from a few very differently regulated genes.

### But what about genes showing an interaction at 50 nM Dex?

```
hi_int_genes <- res.50int_sig %>%
  filter(abs(log2FoldChange) < 10) %>%
  na.omit() %>%
  pull(symbol)
```

Sensitivity from full screen

```
hi_int_full_rhos <- full_rhos %>%
  filter(symbol %in% hi_int_genes) %>%
  filter(Rho_pvalue <= 0.05)
nrow(hi_int_full_rhos)
```

```
## [1] 9
```

Growth from full screen

```
hi_int_full_gammas <- full_gammas %>%
  filter(symbol %in% hi_int_genes) %>%
  filter(Gamma_qvalue <= 0.05)
nrow(hi_int_full_gammas)
```

```
## [1] 3
```

Sensitivity from CAGEK screen

```
hi_int_cagek_rhos <- cagek_rhos %>%
  filter(symbol %in% hi_int_genes) %>%
  filter(CAGEK_Rho_pvalue <= 0.05)
nrow(hi_int_cagek_rhos)
```

```
## [1] 3
```

Growth from CAGEK screen

```
hi_int_cagek_gammas <- cagek_gammas %>%
  filter(symbol %in% hi_int_genes) %>%
  filter(CAGEK_Gamma_pvalue <= 0.05)
nrow(hi_int_cagek_gammas)
```

```
## [1] 7
```

A few genes that show an interaction between 50 nM Dex and Idela show a phenotype in the screens, suggesting that these may be effector genes of idela synergy.

Let’s look at the nature of the interaction to see if it might contribute to synergy.

First, make a table for regulated genes under all conditions:

```
hi_and_both_reg_filt <- hi_and_both_reg %>%
  rownames_to_column('ensembl') %>%
  arrange(symbol, ensembl) %>%
  filter(!duplicated(symbol))

hi_and_both_reg_longer <- hi_and_both_reg_filt %>%
  pivot_longer(idel_log2FC:hiboth_adjp, names_to = c("treatment","stat"), names_pattern
= "(^\\w+)_(\\w+$)") %>%
  pivot_wider(names_from = stat, values_from = value) %>%
  mutate(treatment = factor(treatment, levels = c("idel", "lodex","loboth","hidex","hibo
th")))

hi_and_both_reg_longer %>%
  dplyr::filter(symbol %in% hi_int_full_rhos$symbol) %>%
  ggplot(aes(treatment, log2FC, fill = treatment)) +
  labs(x = "", y = "log2FoldChange") +
  geom_col(position = "dodge") +
  scale_fill_viridis(discrete = T, option = "E") +
  facet_wrap(~symbol, scales = "free") + theme_bw() +
  theme(legend.position="none", axis.text.x = element_text(angle = 45, hjust = 1)) +
  geom_errorbar(aes(ymin=log2FC-lfcse, ymax=log2FC+lfcse), position = position_dodge(wid
th = 0.9), width=0.5, colour="black", size = 0.5)
```

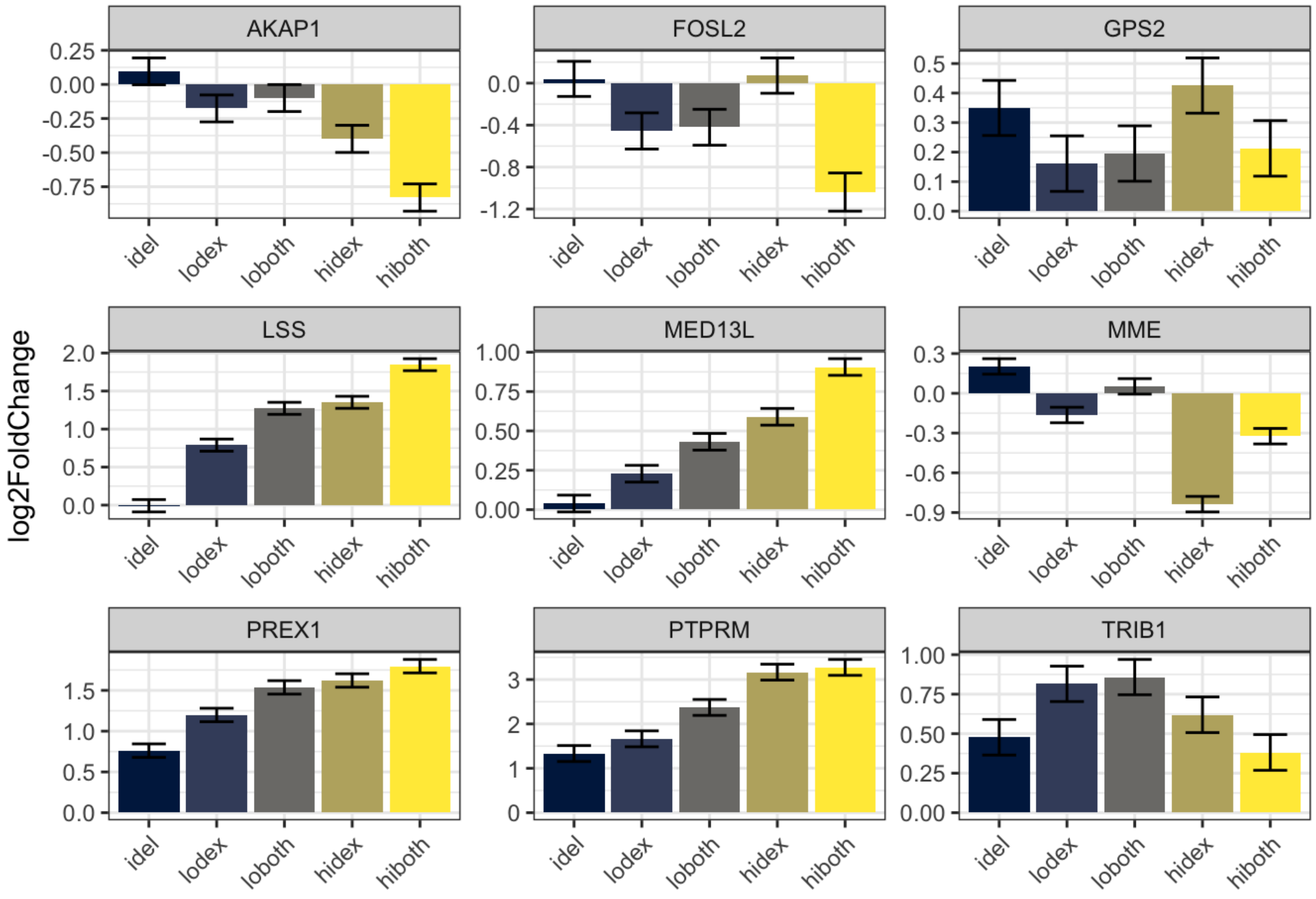

#### Synergistic high dex interaction genes

```
hi_int_genes <- res.50int_sig %>%
  filter(abs(log2FoldChange) < 10) %>%
  filter(log2FoldChange > 0) %>%
  na.omit() %>%
  pull(symbol)

hi_and_both_reg_longer %>%
  dplyr::filter(symbol %in% hi_int_genes) %>%
  ggplot(aes(treatment, log2FC, fill = treatment)) +
  labs(x = "", y = "log2FoldChange") +
  geom_col(position = "dodge") +
  scale_fill_viridis(discrete = T, option = "E") +
  facet_wrap(~symbol) +theme_bw() +
  theme(legend.position="none", axis.text.x = element_text(angle = 45, hjust = 1)) +
  geom_errorbar(aes(ymin=log2FC-lfcse, ymax=log2FC+lfcse), position = position_dodge(wid
th = 0.9), width=0.5, colour="black", size = 0.5)
```

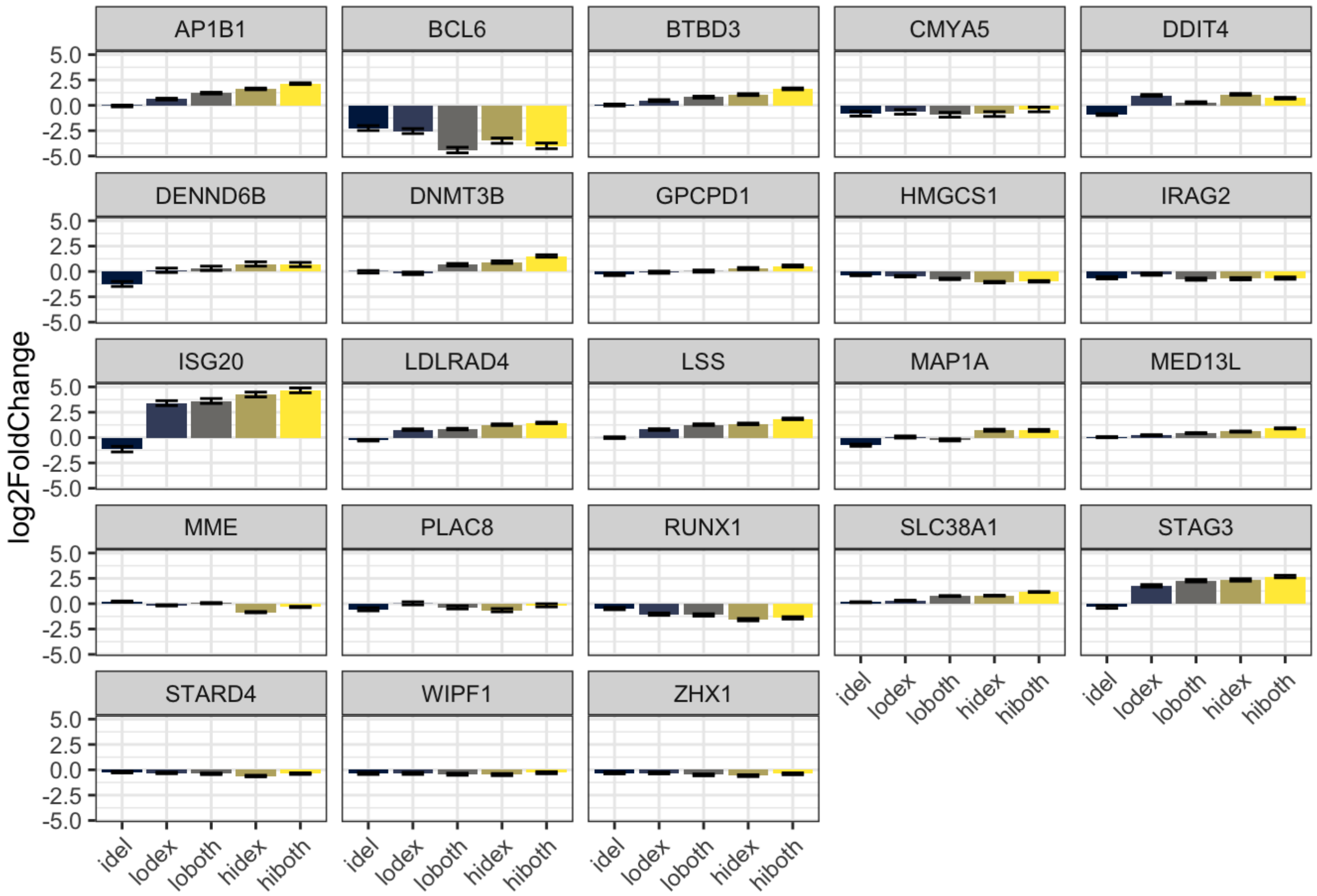

### Common interaction genes b/w 5 nM Dex and 50 nM Dex

#### Syngergistic

```
common_int_syn <- common_int %>%
  dplyr::filter(log2FoldChange.x > 0) %>%
  na.omit()

hi_and_both_reg_longer %>%
  dplyr::filter(symbol %in% common_int_syn$symbol.y) %>%
  ggplot(aes(treatment, log2FC, fill = treatment)) +
  labs(x = "", y = "log2FoldChange") +
  geom_col(position = "dodge") +
  scale_fill_viridis(discrete = T, option = "E") +
  facet_wrap(~symbol) +theme_bw() +
  theme(legend.position="none", axis.text.x = element_text(angle = 45, hjust = 1)) +
  geom_errorbar(aes(ymin=log2FC-lfcse, ymax=log2FC+lfcse), position = position_dodge(wid
th = 0.9), width=0.5, colour="black", size = 0.5)
```

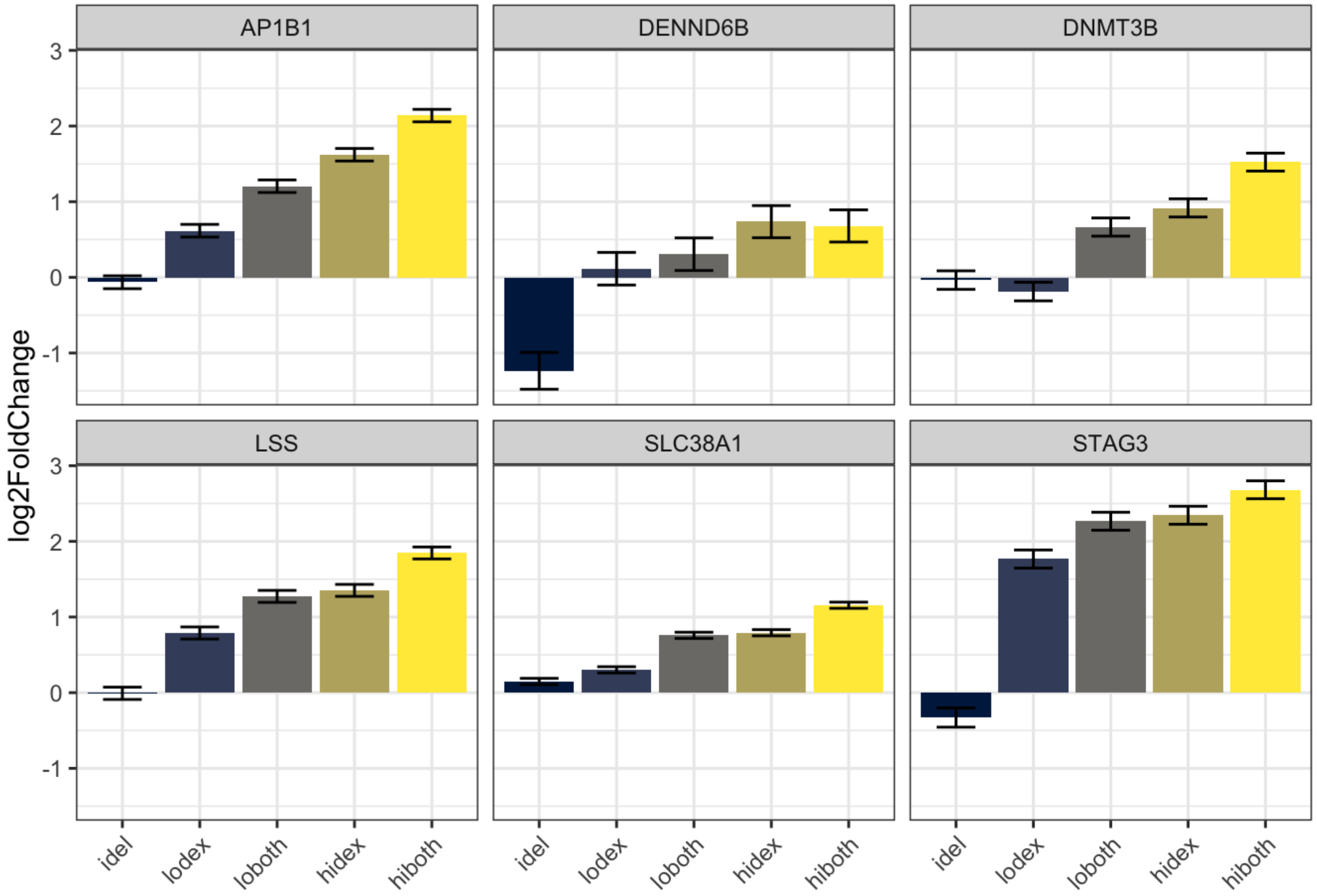

#### Buffering

```
common_int_buff <- common_int %>%
  dplyr::filter(log2FoldChange.x < 0) %>%
  na.omit()

hi_and_both_reg_longer %>%
  dplyr::filter(symbol %in% common_int_buff$symbol.y) %>%
  ggplot(aes(treatment, log2FC, fill = treatment)) +
  labs(x = "", y = "log2FoldChange") +
  geom_col(position = "dodge") +
  scale_fill_viridis(discrete = T, option = "E") +
  facet_wrap(~symbol) +theme_bw() +
  theme(legend.position="none", axis.text.x = element_text(angle = 45, hjust = 1)) +
  geom_errorbar(aes(ymin=log2FC-lfcse, ymax=log2FC+lfcse), position = position_dodge(wid
th = 0.9), width=0.5, colour="black", size = 0.5)
```

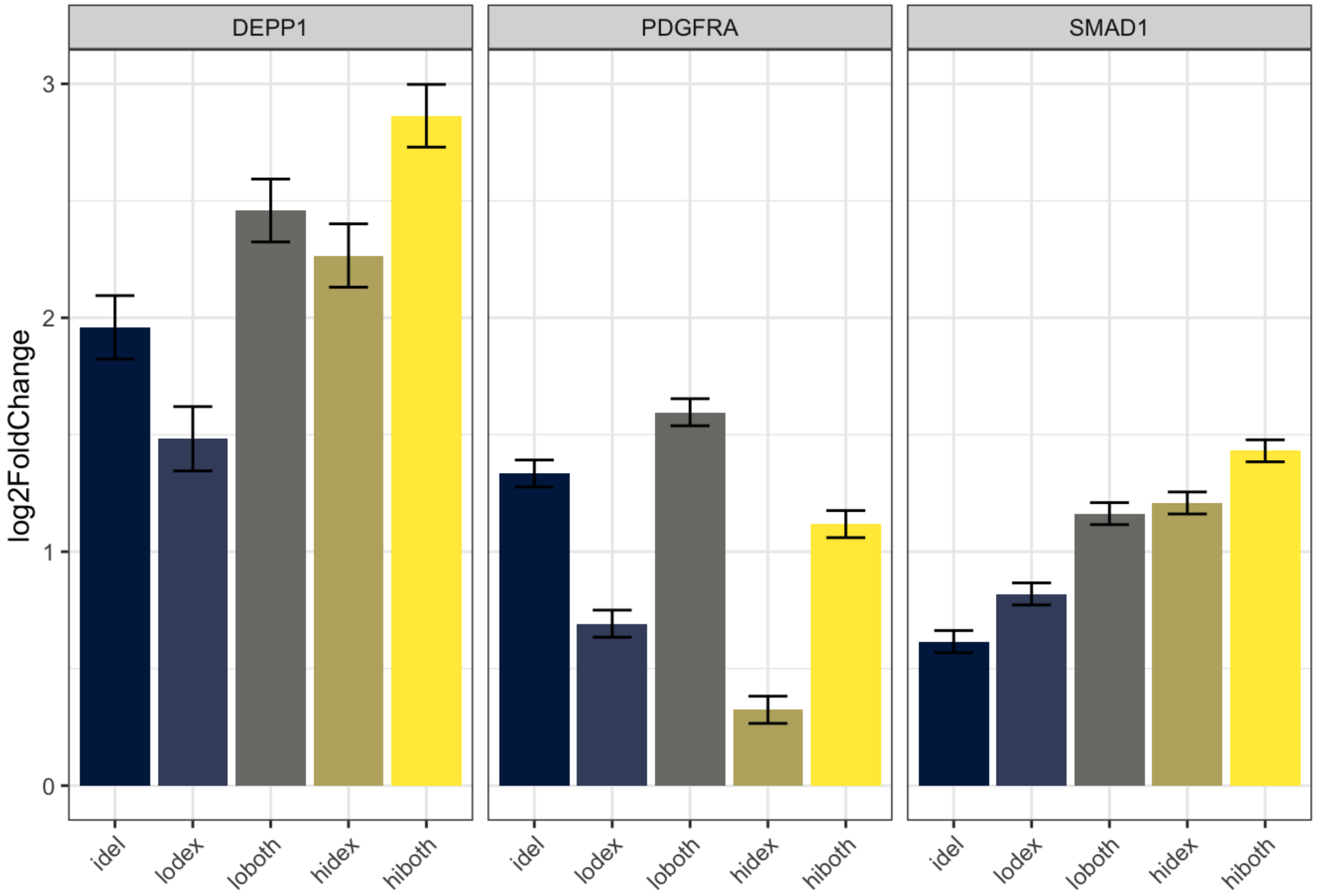

#### How many are greater than additive?

```
hi_int <- hi_and_both_reg_filt %>%
  dplyr::filter(symbol %in% hi_int_genes)

hi_syn_genes <- dplyr::filter(hi_int, abs(hiboth_log2FC) > abs(idel_log2FC + hidex_log2F
C))
```

So 22 genes are regulated more that additively by hidex and idela

#### How many are effectors?

```
hi_syn_full_rhos <- full_rhos %>%
  filter(symbol %in% hi_syn_genes$symbol) %>%
  filter(Rho_pvalue <= 0.05)
nrow(hi_syn_full_rhos)
```

```
## [1] 2
```

One gene - *DDX39A is a positive effector gene* that is synergistically regulated by dex & idela.

```
hi_syn_cagek_rhos <- cagek_rhos %>%
  filter(symbol %in% hi_syn_genes$symbol) %>%
  filter(CAGEK_Rho_pvalue <= 0.05)
nrow(hi_syn_cagek_rhos)
```

```
## [1] 0
```

One gene, *MCM7, is a buffering gene*, meaning it is contributes to dex-induced cell death, but is synergistically downregulated by dex & idela.

```
hi_syn_genes %>%
  pivot_longer(idel_log2FC:hiboth_adjp, names_to = c("treatment","stat"), names_pattern
= "(^\\w+)_(\\w+$)") %>%
  pivot_wider(names_from = stat, values_from = value) %>%
  mutate(treatment = factor(treatment, levels = c("idel", "lodex","loboth","hidex","hibo
th"))) %>%
  ggplot(aes(treatment, log2FC, fill = treatment)) +
  labs(x = "", y = "log2FoldChange") +
  geom_col(position = "dodge") +
  scale_fill_viridis(discrete = T, option = "E") +
  facet_wrap(~symbol) +theme_bw() +
  theme(legend.position="none", axis.text.x = element_text(angle = 45, hjust = 1), pane
l.grid.major = element_line("lightgray",0.25),
        panel.grid.minor = element_line("lightgray",0.1)) +
  geom_errorbar(aes(ymin=log2FC-lfcse, ymax=log2FC+lfcse), position = position_dodge(wid
th = 0.9), width=0.4, colour="black", size = 0.4)
```

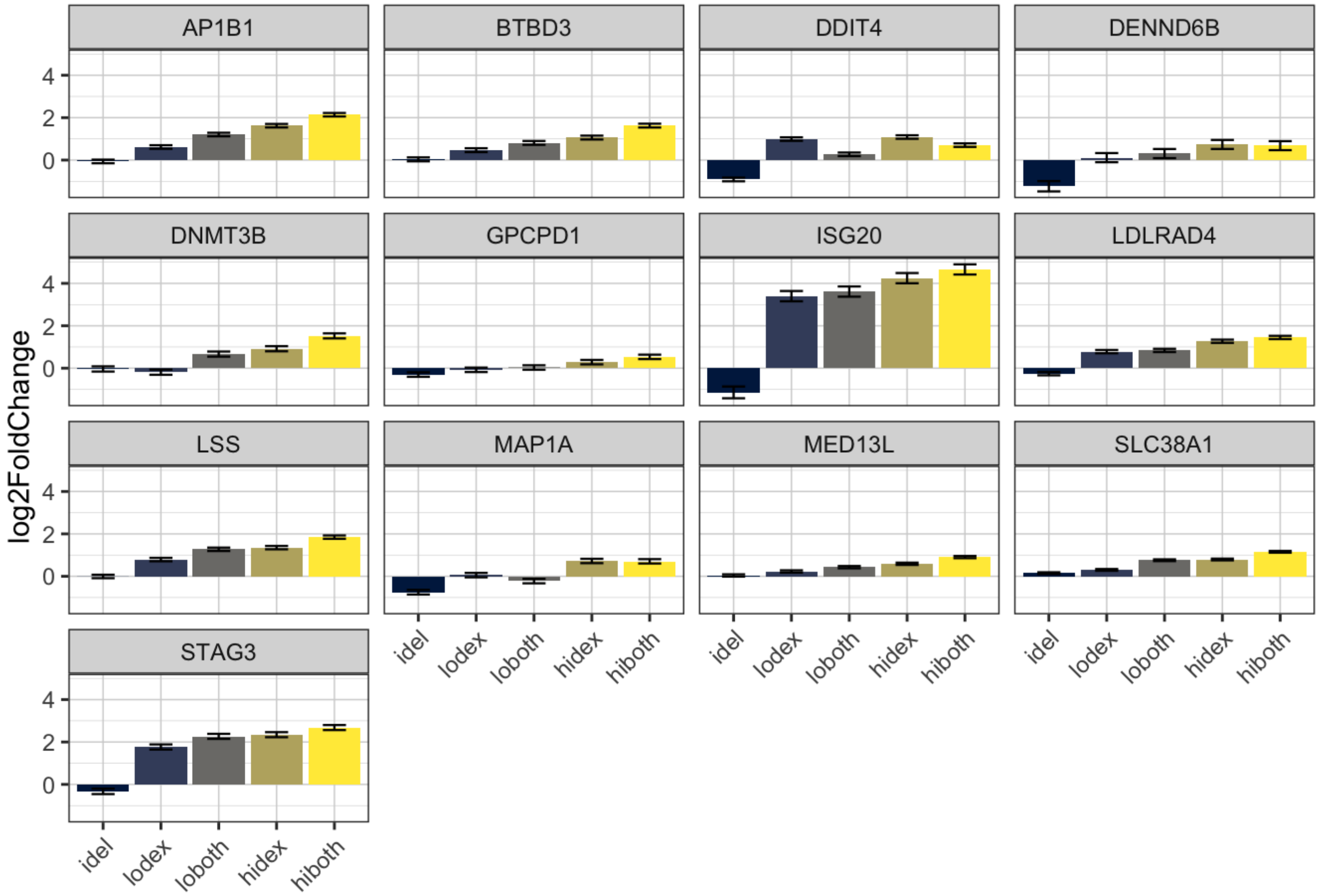

```
ggsave("hidex_syn_interacting_genes.pdf", width = 8, height = 6)
```

### Which effector genes are most strongly regulated?

The way to do this would be to take the significant rho genes, and overlap them with the significantly regulated genes. For those, we then want to look at those with the biggest fold change difference between dex and dex + idela. So let's do it:

```
#let's combine the tables of significant phenotype and significant regulation
olap_rhos_reg <- sig_rhos %>%
  inner_join(lo_and_both_reg, by = c("Symbol" = "symbol")) %>%
  mutate(diff_lo_idel = lodex_log2FC - loboth_log2FC)
n_distinct(olap_rhos_reg$Symbol)
```

```
## [1] 256
```

```
# Same w/ CAGEK screen
c_olap_rhos_reg <- c_sig_rhos %>%
  inner_join(lo_and_both_reg, by = c("Symbol" = "symbol")) %>%
  mutate(diff_lo_idel = lodex_log2FC - loboth_log2FC)
n_distinct(c_olap_rhos_reg)
```

```
## [1] 142
```

```
doub_imp_genes <- intersect(olap_rhos_reg$Symbol, c_olap_rhos_reg$Symbol)
```

Genes that consistently give a phenotype in both screens are our most confident hits. Of these, there are 37 that are significantly regulated under some condition. Perhaps the best thing to do is to make graphs for each of these and pick the ones that are most interesting to test.

```
lo_and_both_reg_longer <- lo_and_both_reg %>%
  pivot_longer(idel_log2FC:hiboth_adjp, names_to = c("treatment","stat"), names_pattern
= "(^\\w+)_(\\w+$)") %>%
  pivot_wider(names_from = stat, values_from = value) %>%
  mutate(treatment = factor(treatment, levels = c("idel", "lodex","loboth","hidex","hibo
th")))

lo_and_both_reg_longer %>%
  dplyr::filter(symbol %in% doub_imp_genes) %>%
  ggplot(aes(treatment, log2FC, fill = treatment)) +
  labs(x = "", y = "log2FoldChange") +
  geom_col(position = "dodge") +
  scale_fill_viridis(discrete = T, option = "E") +
  facet_wrap(~symbol, scales = "free") +theme_bw() +
  theme(legend.position="none", axis.text.x = element_text(angle = 45, hjust = 1)) +
  geom_errorbar(aes(ymin=log2FC-lfcse, ymax=log2FC+lfcse), position = position_dodge(wid
th = 0.9), width=0.5, colour="black", size = 0.5)
```

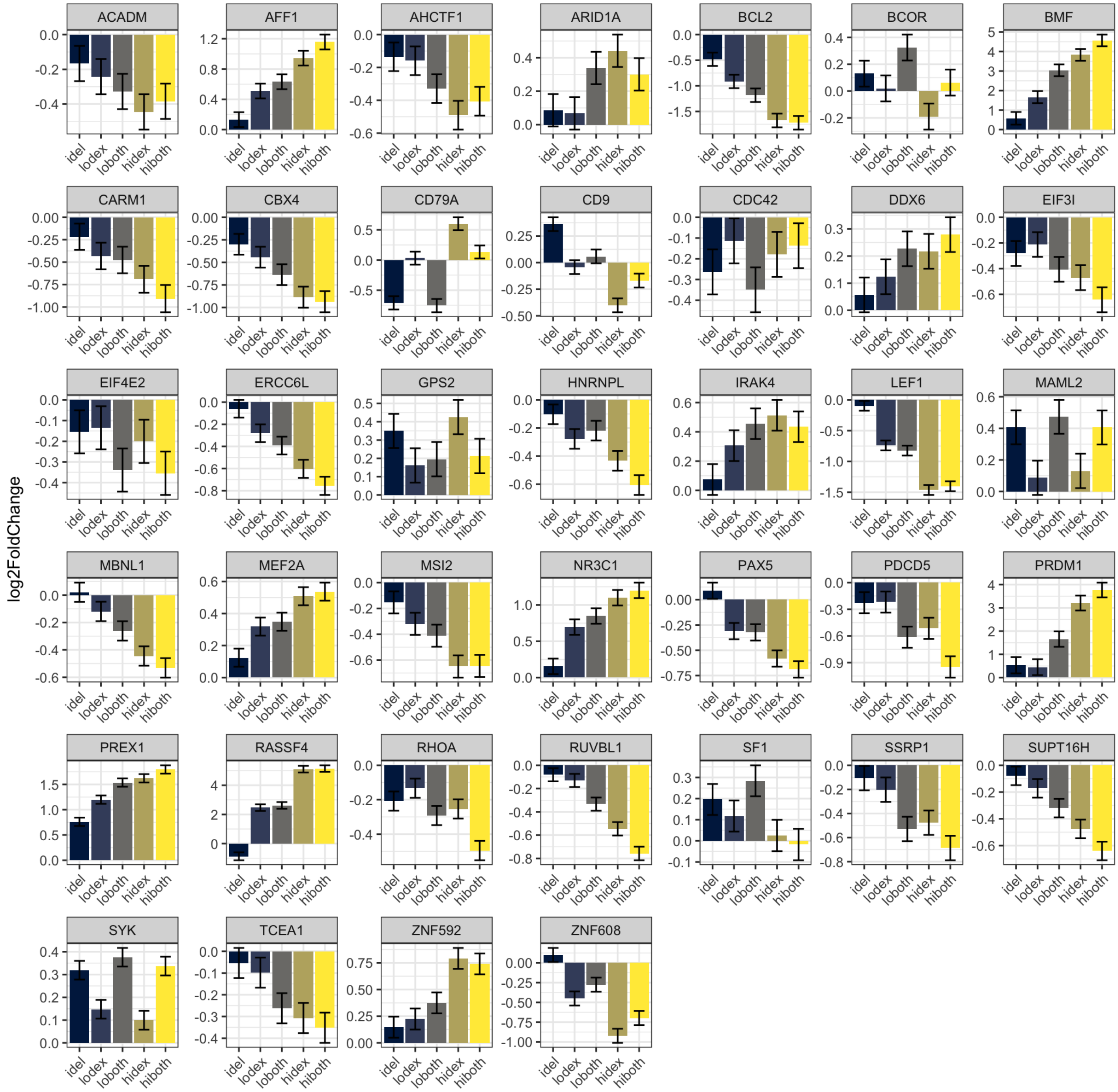

```
lo_and_both_reg_longer %>%
  dplyr::filter(symbol %in% reg_genes) %>%
  ggplot(aes(treatment, log2FC, fill = treatment)) +
  labs(x = "", y = "log2FoldChange") +
  geom_col(position = "dodge") +
  scale_fill_viridis(discrete = T, option = "E") +
  facet_wrap(~symbol, scales = "free") +theme_bw() +
  theme(legend.position="none", axis.text.x = element_text(angle = 45, hjust = 1)) +
  geom_errorbar(aes(ymin=log2FC-lfcse, ymax=log2FC+lfcse), position = position_dodge(wid
th = 0.9), width=0.5, colour="black", size = 0.5)
```

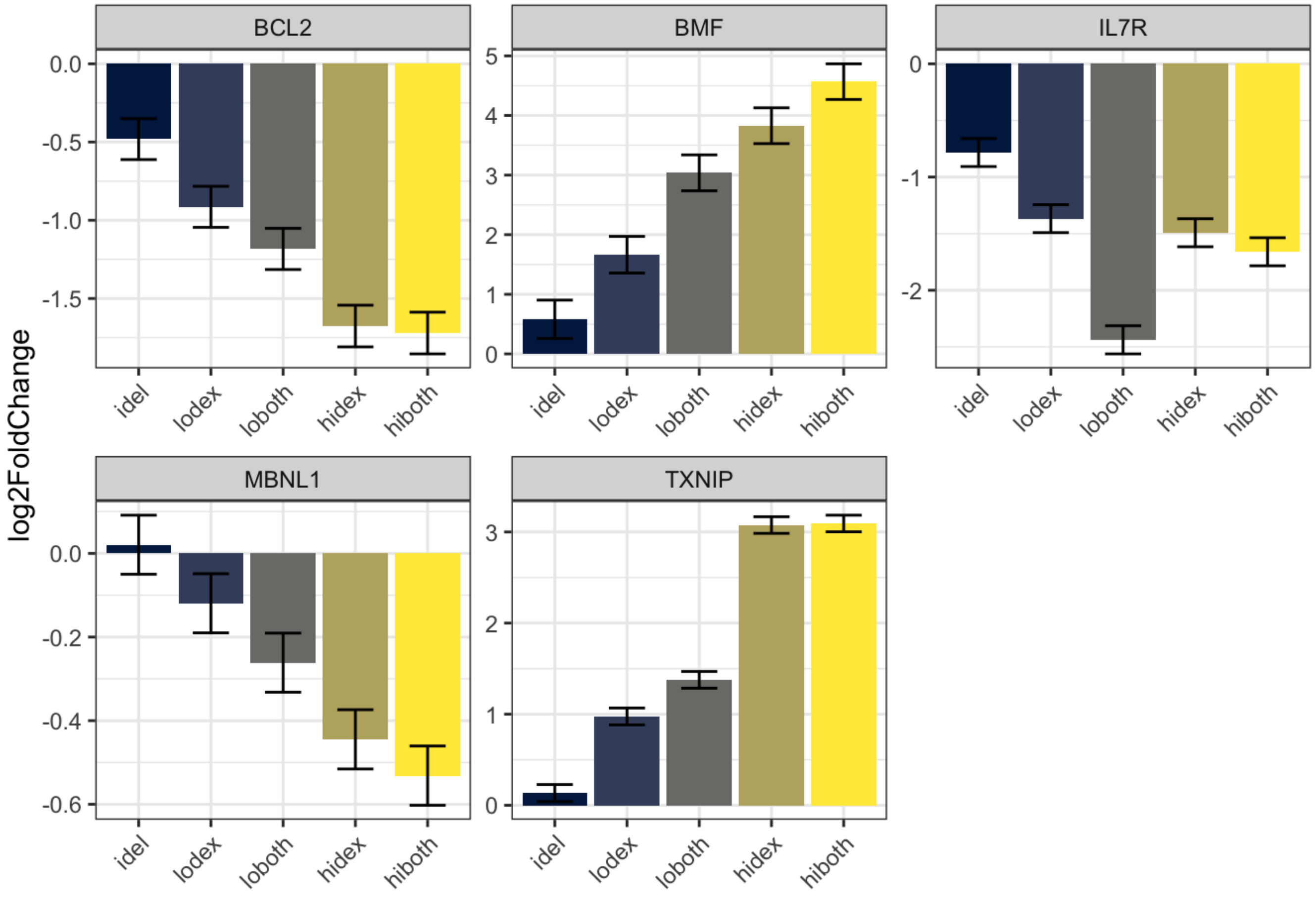

```
sum_longer <- sum_tbl_2 %>%
  pivot_longer(idel_log2FC:hiboth_adjp, names_to = c("treatment","stat"), names_pattern
= "(^\\w+)_(\\w+$)") %>%
  pivot_wider(names_from = stat, values_from = value) %>%
  mutate(treatment = factor(treatment, levels = c("idel", "lodex","loboth","hidex","hibo
th")))

jess_genes <- c("BCL2", "BCL2L11", "IL7R", "TXNIP")

sum_longer %>%
  dplyr::filter(symbol %in% jess_genes) %>%
  ggplot(aes(treatment, log2FC, fill = treatment)) +
  labs(x = "", y = "log2FoldChange") +
  geom_col(position = "dodge") +
  scale_fill_viridis(discrete = T, option = "E") +
  facet_wrap(~symbol) +theme_bw() +
  theme(legend.position="none", axis.text.x = element_text(angle = 45, hjust = 1)) +
  geom_errorbar(aes(ymin=log2FC-lfcse, ymax=log2FC+lfcse), position = position_dodge(wid
th = 0.9), width=0.5, colour="black", size = 0.5)
```

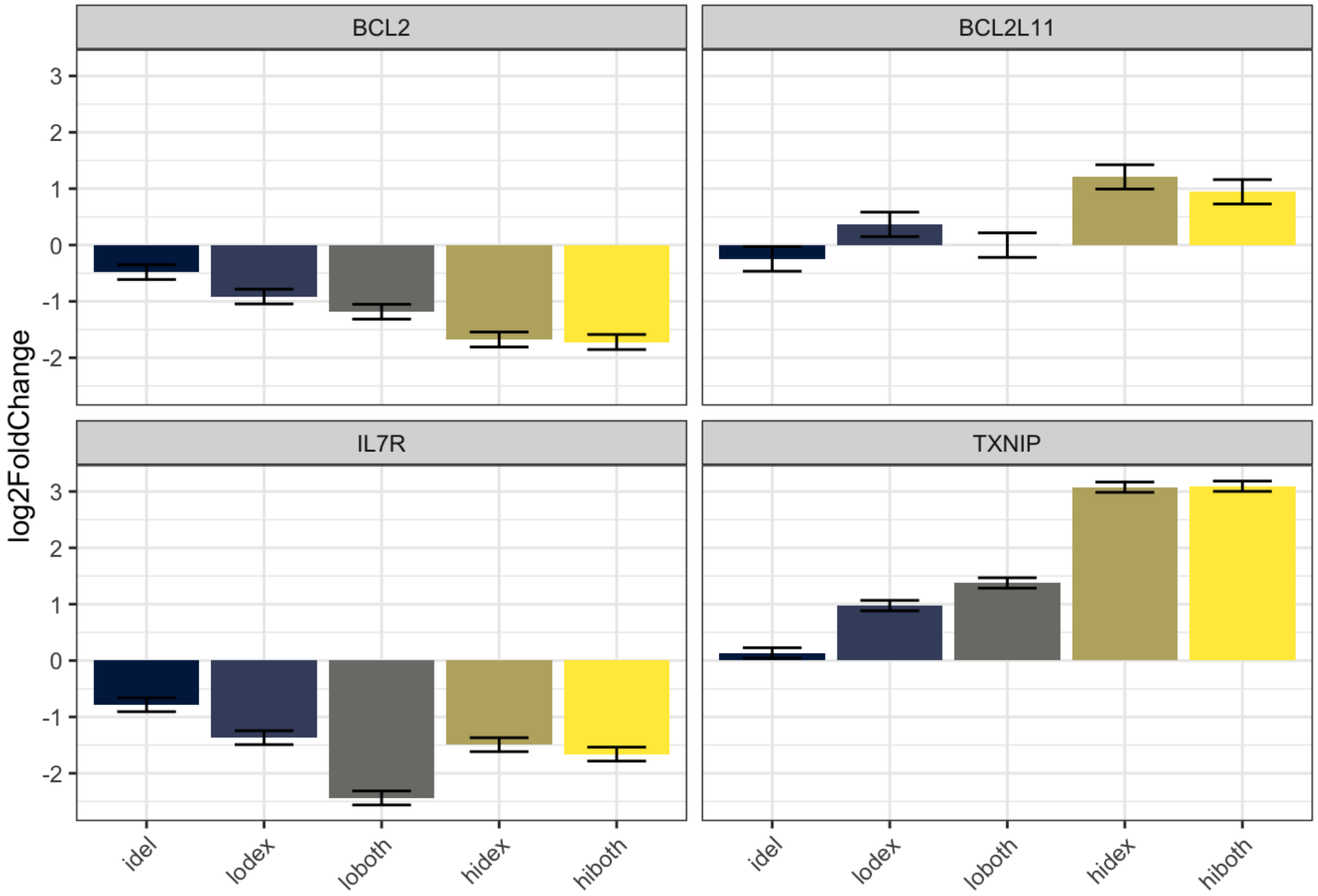

```
ggsave(filename = "eff_genes_barplot.pdf", height = 4, width = 4)
```

```
x <- c("CHD4
MTA2
RBBP4
RBBP7
GATAD2A
GATAD2B
HDAC1
HDAC2
SMARCC2
SMARCA4
SMARCD2
SMARCC1
SMARCE1
SMARCB1
SMARCA5
SMARCD1
SMARCA2
ARID1A
ARID1B
ARID2
NCOR1
NCOR2
Sin3A
SUDS3
FAM60A
PHF12
KDM1A
KDM2B
KDM3A
KDM3B
KDM5A
KDM6A
KAT5
KMT2D
WDR5
ASH2L
MED14
MED15
MED17
MED22
MED25
MED8
TRIM24
TRIM28
PELP1
PAXIP1
RCOR1
RCOR3
TCF20
NCOA3
NCOA6
TADA2A
```

```
ZBTB20
ZBTB9
ZBED3
HCFC1
PPM1G
PPP4R1
MCM4
MCM5
ZNF512")
cofactors <- read_table(x, col_names = FALSE)

full_rhos_min %>%
  filter(Rho_pvalue <= 0.05 & symbol %in% cofactors$X1)
```

```
## # A tibble: 14 × 3
##    symbol  Rho_phenotype Rho_pvalue
##    <chr>           <dbl>      <dbl>
##  1 ARID1A          0.338   1.48e- 4
##  2 CHD4            0.392   7.50e- 9
##  3 HDAC2          -0.326   9.07e- 3
##  4 KAT5           -0.288   1.73e- 2
##  5 KDM1A          -0.281   1.65e- 2
##  6 KDM2B           0.112   2.71e- 3
##  7 KDM5A          -0.250   1.31e- 3
##  8 KMT2D          -0.400   4.50e- 9
##  9 MED14           0.325   2.59e- 5
## 10 NCOR1           0.531   4.57e- 7
## 11 NCOR2           0.400   2.10e-11
## 12 PAXIP1          0.132   3.82e- 2
## 13 SMARCD1         0.196   1.30e- 3
## 14 TADA2A         -0.298   3.51e- 2
```

### Analyzing Patient Samples with RUV and DESeq2

zimmermanjo

11/28/2022

#### This markdown analyzes 7 primary patient B-ALL specimens to determine differences in gene regulation with glucocorticoids +/- idelalisib for all samples and also subsetting into glucocorticoid sensitive vs. glucocorticoid resistant specimens. The approach incorporates RUVSeq into the DESeq workflow to identify a set of empirical control genes and use these as controls in our model design. This will follow the DESeq2 workflow for section 2.3 through 3.1 (reading in data), then section 8.2 (RUV), then section 5 on (running differential expression analysis) - workflow here (https://bioconductor.org/packages/release/workflows/vignettes/rnaseqGene/inst/doc/rnaseqGen ruv-with-deseq2).

Import the sample data and select for columns to make a conditions tables - section 2.3

Display the conditions table when you are done. Ensure that the treatment, time, and cell type are all complete.

```
sample_table <- list.files("quants/") %>%
  as_tibble() %>%
  separate(col = "value", into = c("number", "txnum", "replicate", "treatment", "patient")) %>%
  mutate(gc = str_extract(treatment, pattern = "Dex|Pred")) %>%
  mutate(idela = str_extract(treatment, pattern = "Idela")) %>%
  replace_na(list(gc = "Veh", idela = "Veh")) %>%
  mutate(treament = as.factor(treatment), patient = as.factor(patient), gc = as.factor(gc), idela = as.factor(idela))
```

According to the JSON files:

What are the average number of reads per sample and what is the average mapping percentage?

```
dir <- "quants"

Sample <- list.files("quants/")

test <- sapply(list.files(dir), function(x) rjson::fromJSON(file = paste0("quants/", x, "/aux_info/meta_info.json")))

#Output is a list

table <- as.data.frame(t(test))

## There are a few variables with multiple values per observation.  They might be interesting, but we'll select them out
table_filt <- table %>%
  dplyr::select(-quant_errors, -eq_class_properties, -length_classes)

table_filt <- add_column(table_filt, Sample, .before = TRUE)
tidy_tbl <- map_df(table_filt, unlist)

tidy_tbl <- tidy_tbl %>%
  separate(col = Sample, into = c("number", "txnum", "replicate", "treatment", "patient"), remove = FALSE)
```

Average number of reads:

```
ave_reads <- tidy_tbl %>%
  pull(num_processed) %>%
  mean() %>%
  round(0)

ave_reads
```

```
## [1] 40324996
```

```
ggplot(tidy_tbl, aes(Sample, num_mapped, fill = patient)) +
  geom_col() +
  geom_hline(yintercept = mean(tidy_tbl$num_mapped)) +
  theme(axis.text.x = element_text(angle = 45, hjust=1))
```

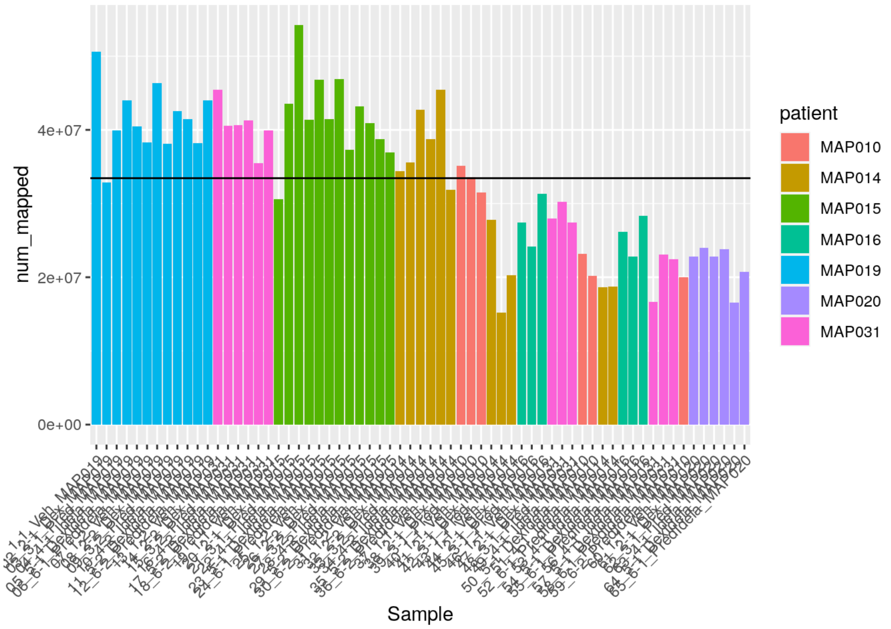

```
ave_mapped <- tidy_tbl %>%
  pull(percent_mapped) %>%
  mean() %>%
  round(0)

ave_mapped
```

```
## [1] 83
```

```
ggplot(tidy_tbl, aes(Sample, percent_mapped, fill = patient)) +
  geom_col() +
  geom_hline(yintercept = mean(tidy_tbl$percent_mapped)) +
  theme(axis.text.x = element_text(angle = 45, hjust=1))
```

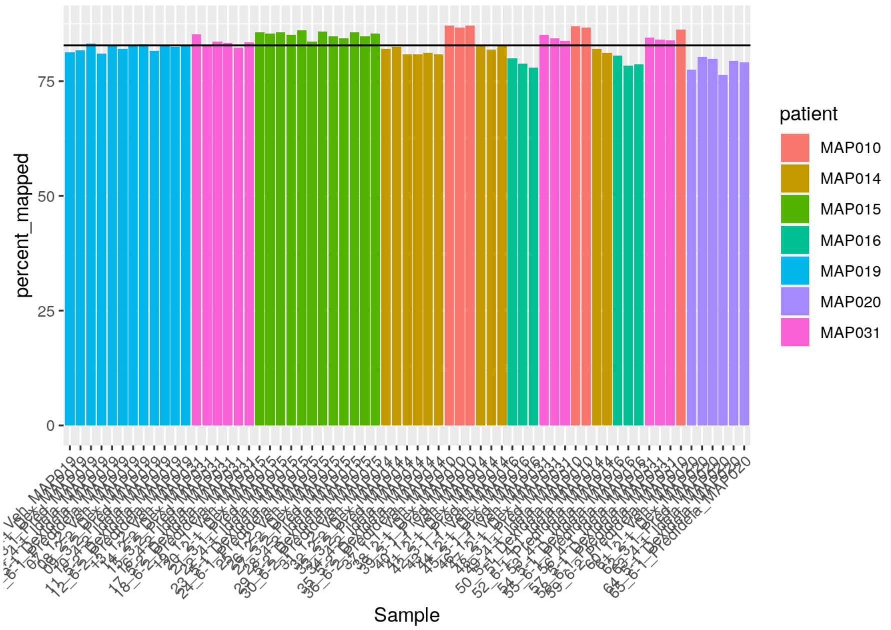

The average number of reads is 4.0324996^{7}

The average percent mapped is 83%

#### Import count tables into R

```
test_sample <- list.files(dir)
all.equal(test_sample, tidy_tbl$Sample)
```

```
## [1] TRUE
```

```
sample_table$names <- test_sample
sample_table$files <- file.path(dir, sample_table$names, "quant.sf")
file.exists(sample_table$files)
```

```
##  [1] TRUE TRUE TRUE TRUE TRUE TRUE TRUE TRUE TRUE TRUE TRUE TRUE TRUE TRUE TRUE
## [16] TRUE TRUE TRUE TRUE TRUE TRUE TRUE TRUE TRUE TRUE TRUE TRUE TRUE TRUE TRUE
## [31] TRUE TRUE TRUE TRUE TRUE TRUE TRUE TRUE TRUE TRUE TRUE TRUE TRUE TRUE TRUE
## [46] TRUE TRUE TRUE TRUE TRUE TRUE TRUE TRUE TRUE TRUE TRUE TRUE TRUE TRUE TRUE
## [61] TRUE TRUE TRUE TRUE TRUE
```

```
# Note that the order of factors had to be changed to put Vehicle first, since it is the control condition

sample_table <- sample_table %>%
  dplyr::select(-treament) %>%
  mutate(treatment = as_factor(treatment)) %>%
  mutate(gc = factor(gc, levels = c("Veh", "Dex", "Pred"))) %>%
  mutate(idela = factor(idela, levels = c("Veh", "Idela")))

se <- tximeta(sample_table)
```

```
## importing quantifications
```

```
## reading in files with read_tsv
```

```
## 1 2 3 4 5 6 7 8 9 10 11 12 13 14 15 16 17 18 19 20 21 22 23 24 25 26 27 28 29 30 31 32 33 34 35 36 37 38 39 40 41 42 43 4
4 45 46 47 48 49 50 51 52 53 54 55 56 57 58 59 60 61 62 63 64 65
## found matching transcriptome:
## [ GENCODE - Homo sapiens - release 38 ]
## loading existing TxDb created: 2021-10-11 19:29:18
## loading existing transcript ranges created: 2021-10-11 19:29:20
## fetching genome info for GENCODE
```

```
## Error in .order_seqlevels(chrom_sizes[, "chrom"]) :
##   !anyNA(m32) is not TRUE
```

##### Summarize to gene for gene-level analysis

```
dim(se)
```

```
## [1] 236186     65
```

```
gse <- summarizeToGene(se)
```

```
## loading existing TxDb created: 2021-10-11 19:29:18
```

```
## obtaining transcript-to-gene mapping from database
```

```
## loading existing gene ranges created: 2021-10-11 19:29:47
```

```
## summarizing abundance
```

```
## summarizing counts
```

```
## summarizing length
```

```
dim(gse)
```

```
## [1] 60230    65
```

##### Export count table

```
count_table <- round(assays(gse)$counts, 0) %>%
  as_tibble(rownames = "ensembl")
 # write_csv(count_table, "pt_samples_GCidela_count_table.csv")
```

##### Specify the model (formula) into DESeq and assign to object “dds”

Per DESeq2 guide section 8.2, we need to run DESeq and results first without any batch effect to obtain p-values for the analysis.

```
dds <- DESeqDataSet(gse, design = ~ idela + gc)
```

```
## using counts and average transcript lengths from tximeta
```

```
nrow(dds)
```

```
## [1] 60230
```

#### Plot PCA of samples

```
vsd <- vst(dds, blind = FALSE)

plotPCA(vsd, intgroup = c("gc", "idela"))
```

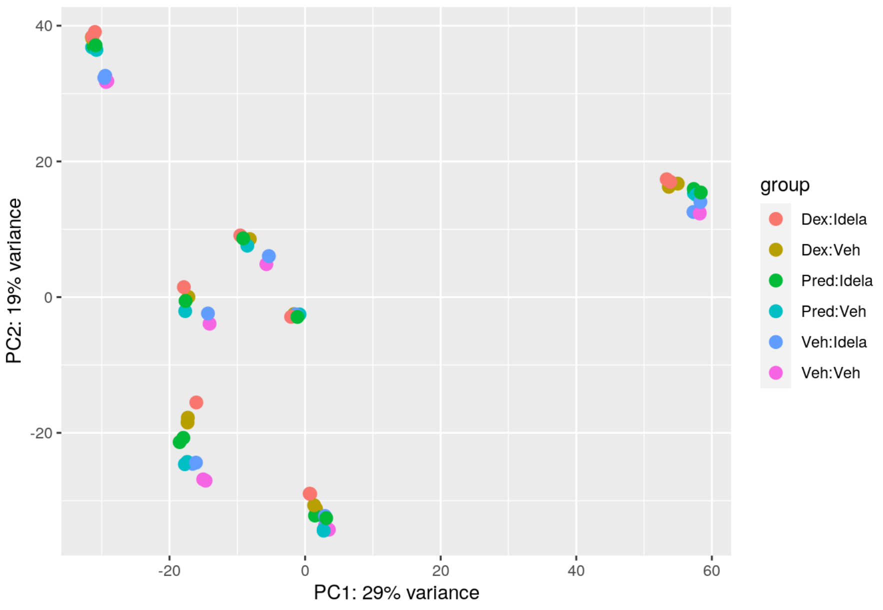

```
pcaData <- plotPCA(vsd, intgroup = c( "gc", "idela", "patient"), returnData = TRUE)

percentVar <- round(100 * attr(pcaData, "percentVar"))

ggplot(pcaData, aes(x = PC1, y = PC2, color = gc, shape = idela)) +
  geom_point(size =3) +
  xlab(paste0("PC1: ", percentVar[1], "% variance")) +
  ylab(paste0("PC2: ", percentVar[2], "% variance")) +
  coord_fixed() +
  ggtitle("PCA with VST data")
```

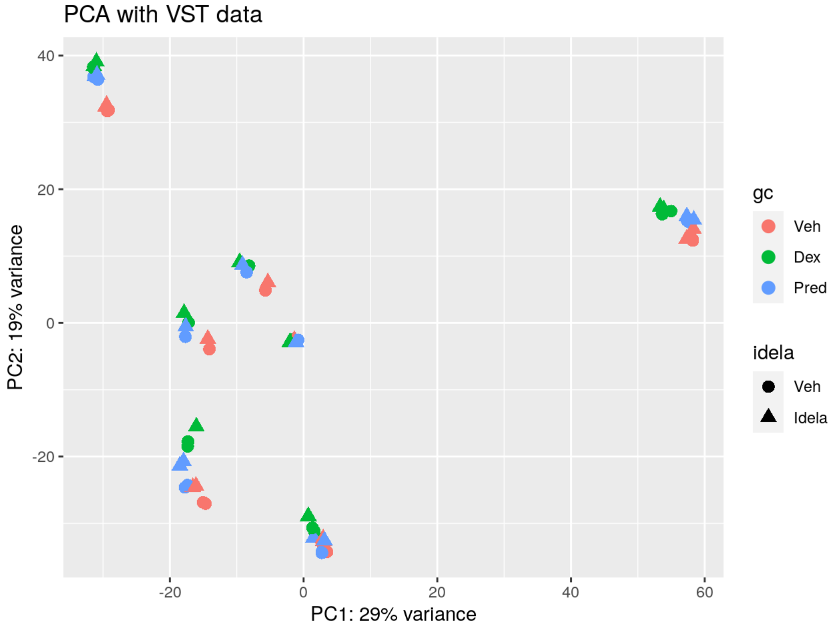

The data likely group best by sample. After that there seems to be a typical progression in PC2 from Veh ==> Pred ==> Dex with grades between that may be due to idela for most of the clusters.

##### Perform differential gene expression testing in order to use RUV:

```
dds <- DESeq(dds)
```

```
## estimating size factors
```

```
## using 'avgTxLength' from assays(dds), correcting for library size
```

```
## estimating dispersions
```

```
## gene-wise dispersion estimates
```

```
## mean-dispersion relationship
```

```
## final dispersion estimates
```

```
## fitting model and testing
```

```
## -- replacing outliers and refitting for 743 genes
## -- DESeq argument 'minReplicatesForReplace' = 7
## -- original counts are preserved in counts(dds)
```

```
## estimating dispersions
```

```
## fitting model and testing
```

```
resultsNames(dds)
```

```
## [1] "Intercept"          "idela_Idela_vs_Veh" "gc_Dex_vs_Veh"
## [4] "gc_Pred_vs_Veh"
```

##### Creating a results table named “res” to continue with using RUV per section 8.2 in the DESeq2 workflow

```
res <- results(dds)
```

##### Pulling out empirical control genes:

```
set <- newSeqExpressionSet(counts(dds))
idx  <- rowSums(counts(set) > 5) >= 2
set  <- set[idx, ]
set <- betweenLaneNormalization(set, which="upper")
not.sig <- rownames(res)[which(res$pvalue > .1)]
empirical <- rownames(set)[ rownames(set) %in% not.sig ]
set <- RUVg(set, empirical, k=2)
pData(set)
```

```
##                                    W_1         W_2
## 01_1-1_Veh_MAP019       -0.1570925289  0.14231896
## 02_2-1_Dex_MAP019       -0.1403925511  0.16901237
## 03_3-1_Pred_MAP019      -0.1410906668  0.15670503
## 04_4-1_Idela_MAP019     -0.1593219045  0.15000677
## 05_5-1_DexIdela_MAP019  -0.1357574912  0.17454907
## 06_6-1_PredIdela_MAP019 -0.1457858846  0.16619738
## 07_1-2_Veh_MAP019       -0.1519952449  0.14176847
## 08_2-2_Dex_MAP019       -0.1345479360  0.16554538
## 09_3-2_Pred_MAP019      -0.1435080022  0.16068488
## 10_4-2_Idela_MAP019     -0.1561746962  0.14493180
## 11_5-2_DexIdela_MAP019  -0.1384382329  0.17294157
## 12_6-2_PredIdela_MAP019 -0.1433285505  0.16094102
## 13_1-2_Veh_MAP031        0.2418623013  0.06098237
## 14_2-2_Dex_MAP031        0.2151264048  0.09551423
## 15_3-2_Pred_MAP031       0.2306562541  0.08533669
## 16_4-2_Idela_MAP031      0.2337462364  0.07171820
## 17_5-2_DexIdela_MAP031   0.2132941673  0.10391073
## 18_6-2_PredIdela_MAP031  0.2299313494  0.09232810
## 19_1-1_Veh_MAP015        0.0058478318 -0.18984994
## 20_2-1_Dex_MAP015        0.0033730192 -0.15183915
## 21_3-1_Pred_MAP015       0.0009658334 -0.16855994
## 22_4-1_Idela_MAP015     -0.0018835534 -0.16514001
## 23_5-1_DexIdela_MAP015   0.0029599804 -0.14928873
## 24_6-1_PredIdela_MAP015 -0.0087207666 -0.15861903
## 25_1-2_Veh_MAP015        0.0047588461 -0.17495536
## 26_2-2_Dex_MAP015        0.0016809477 -0.15319929
## 27_3-2_Pred_MAP015       0.0003779556 -0.16602859
## 28_4-2_Idela_MAP015     -0.0002723644 -0.16745141
## 29_5-2_DexIdela_MAP015   0.0007426903 -0.14297827
## 30_6-2_PredIdela_MAP015  0.0012814861 -0.16042630
## 31_1-2_Veh_MAP014       -0.0443161394 -0.13836320
## 32_2-2_Dex_MAP014       -0.0376411321 -0.09838826
## 33_3-2_Pred_MAP014      -0.0471367751 -0.12366609
## 34_4-2_Idela_MAP014     -0.0554887447 -0.11873862
## 35_5-2_DexIdela_MAP014  -0.0396406502 -0.08925541
## 36_6-2_PredIdela_MAP014 -0.0534785509 -0.11198082
## 37_1-1_Veh_MAP010        0.0029601286 -0.08484722
## 38_2-1_Dex_MAP010        0.0056717579 -0.08350589
## 39_3-1_Pred_MAP010       0.0047123508 -0.08436380
## 40_1-1_Veh_MAP014       -0.0365505469 -0.14025923
## 41_2-1_Dex_MAP014       -0.0373123851 -0.10545953
## 42_3-1_Pred_MAP014      -0.0367077207 -0.12826168
## 43_1-1_Veh_MAP016       -0.0976943107  0.00219403
## 44_2-1_Dex_MAP016       -0.0964185611  0.02546402
## 45_3-1_Pred_MAP016      -0.1071751676  0.01890825
## 46_1-1_Veh_MAP031        0.2489326445  0.06418422
## 47_2-1_Dex_MAP031        0.2314771498  0.09620950
## 48_3-1_Pred_MAP031       0.2400327857  0.08464642
## 49_4-1_Idela_MAP010      0.0030157818 -0.08784798
## 50_5-1_DexIdela_MAP010   0.0004730731 -0.08954625
## 51_4-1_Idela_MAP014     -0.0431129078 -0.12898647
## 52_6-1_PredIdela_MAP014 -0.0456913610 -0.11659781
## 53_4-1_Idela_MAP016     -0.0971200219  0.01160844
## 54_5-1_DexIdela_MAP016  -0.0984152137  0.03977316
## 55_6-1_PredIdela_MAP016 -0.1073027881  0.02738673
## 56_4-1_Idela_MAP031      0.2535928428  0.07361277
## 57_5-1_DexIdela_MAP031   0.2294073326  0.09934085
## 58_6-1_PredIdela_MAP031  0.2473632430  0.09285890
## 59_6-2_PredIdela_MAP010  0.0010542946 -0.08515617
## 60_1-1_Veh_MAP020       -0.0053662093  0.10665495
## 61_2-1_Dex_MAP020        0.0001281042  0.12181926
## 62_3-1_Pred_MAP020       0.0020090928  0.11441358
## 63_4-1_Idela_MAP020     -0.0062768296  0.11595460
## 64_5-1_DexIdela_MAP020  -0.0079997586  0.12918846
## 65_6-1_PredIdela_MAP020  0.0017202627  0.12394929
```

Plotting the factors estimated by RUV:

```
par(mfrow = c(2, 1), mar = c(3,5,3,1))
for (i in 1:2) {
  stripchart(pData(set)[, i] ~ dds$patient, vertical = TRUE, main = paste0("W", i))
  abline(h = 0)
 }
```

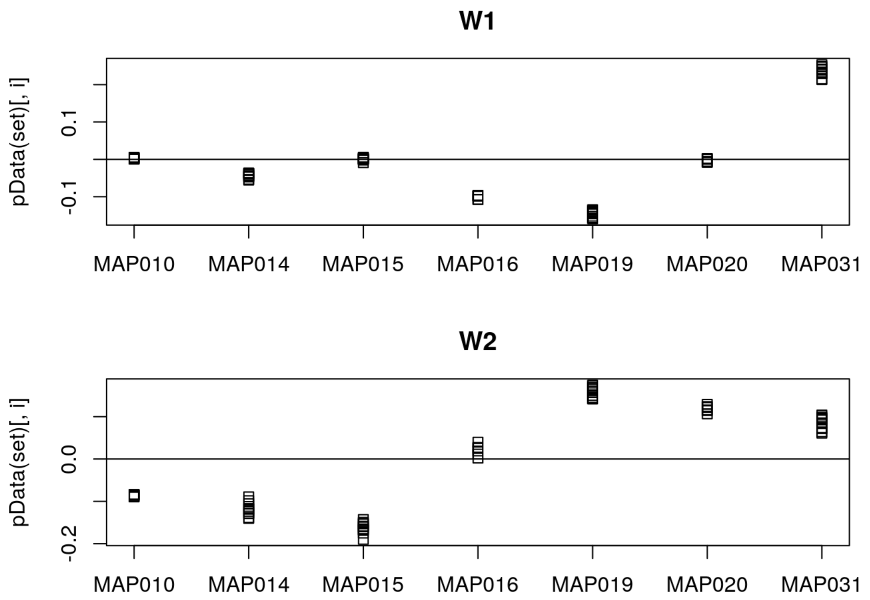

Adding the unwanted variation to the model design

Also adding in grouping variable at this stage

```
ddsruv <- dds

ddsruv$W1 <- set$W_1
ddsruv$W2 <- set$W_2

# design(ddsruv) <- ~ W1 + W2 + idela + gc + idela:gc

ddsruv$group <- factor(paste0(ddsruv$gc, ddsruv$idela))
ddsruv$group <- relevel(ddsruv$group, "VehVeh")

design(ddsruv) <- ~ W1 + W2 + group
```

Re-run DESeq with this new design to re-estimate parameters and results.

```
ddsruv <- DESeq(ddsruv)
```

```
## using pre-existing normalization factors
```

```
## estimating dispersions
```

```
## found already estimated dispersions, replacing these
```

```
## gene-wise dispersion estimates
```

```
## mean-dispersion relationship
```

```
## final dispersion estimates
```

```
## fitting model and testing
```

```
resultsNames(ddsruv)
```

```
## [1] "Intercept"                 "W1"
## [3] "W2"                        "group_DexIdela_vs_VehVeh"
## [5] "group_DexVeh_vs_VehVeh"    "group_PredIdela_vs_VehVeh"
## [7] "group_PredVeh_vs_VehVeh"   "group_VehIdela_vs_VehVeh"
```

##### Now I will filter out genes with < 2 reads on average per sample.

if there are 65 samples, that'd be 2 * 65 or 130

```
ddsruv <- ddsruv[ rowSums(counts(ddsruv)) > 130, ]
nrow(ddsruv)
```

```
## [1] 25100
```

#### Now attempting to create results tables

##### Dex Alone

```
dex_res <- results(ddsruv, name = "group_DexVeh_vs_VehVeh", independentFiltering = TRUE, alpha = 0.01)

summary(dex_res)
```

```
##
## out of 25100 with nonzero total read count
## adjusted p-value < 0.01
## LFC > 0 (up)       : 940, 3.7%
## LFC < 0 (down)     : 406, 1.6%
## outliers [1]       : 0, 0%
## low counts [2]     : 0, 0%
## (mean count < 2)
## [1] see 'cooksCutoff' argument of ?results
## [2] see 'independentFiltering' argument of ?results
```

##### Pred Alone

```
pred_res <- results(ddsruv, name = "group_PredVeh_vs_VehVeh", independentFiltering = TRUE, alpha = 0.01)

summary(pred_res)
```

```
##
## out of 25100 with nonzero total read count
## adjusted p-value < 0.01
## LFC > 0 (up)       : 84, 0.33%
## LFC < 0 (down)     : 15, 0.06%
## outliers [1]       : 0, 0%
## low counts [2]     : 0, 0%
## (mean count < 2)
## [1] see 'cooksCutoff' argument of ?results
## [2] see 'independentFiltering' argument of ?results
```

##### Idela Alone

```
idela_res <- results(ddsruv, name = "group_VehIdela_vs_VehVeh", independentFiltering = TRUE, alpha = 0.01)

summary(idela_res)
```

```
##
## out of 25100 with nonzero total read count
## adjusted p-value < 0.01
## LFC > 0 (up)       : 0, 0%
## LFC < 0 (down)     : 7, 0.028%
## outliers [1]       : 0, 0%
## low counts [2]     : 0, 0%
## (mean count < 2)
## [1] see 'cooksCutoff' argument of ?results
## [2] see 'independentFiltering' argument of ?results
```

##### Dex + Idela

```
di_res <- results(ddsruv, name = "group_DexIdela_vs_VehVeh", independentFiltering = TRUE, alpha = 0.01)

summary(di_res)
```

```
##
## out of 25100 with nonzero total read count
## adjusted p-value < 0.01
## LFC > 0 (up)       : 915, 3.6%
## LFC < 0 (down)     : 426, 1.7%
## outliers [1]       : 0, 0%
## low counts [2]     : 0, 0%
## (mean count < 2)
## [1] see 'cooksCutoff' argument of ?results
## [2] see 'independentFiltering' argument of ?results
```

Pred + Idela

```
pi_res <- results(ddsruv, name = "group_PredIdela_vs_VehVeh", independentFiltering = TRUE, alpha = 0.01)

summary(pi_res)
```

```
##
## out of 25100 with nonzero total read count
## adjusted p-value < 0.01
## LFC > 0 (up)       : 120, 0.48%
## LFC < 0 (down)     : 27, 0.11%
## outliers [1]       : 0, 0%
## low counts [2]     : 3893, 16%
## (mean count < 7)
## [1] see 'cooksCutoff' argument of ?results
## [2] see 'independentFiltering' argument of ?results
```

#### Continue with making tables and plots to compare gene regulation between conditions

##### Prep and merge tables - wrote a function to help with this

```
results_table <- function(res_name, deseq_obj, new_name) {
  df <- results(deseq_obj, name = res_name)
  df <- as.data.frame(df)
  df <- df[,c(1:3, 5:6)]
  colnames(df) <- c("base_mean", paste0(new_name, "_log2FC"), paste0(new_name,"_lfcse"), paste0(new_name, "_pval"), paste0(n
ew_name, "_adjp"))
  new_name <- df
  return(new_name)
}

idela <- results_table("group_VehIdela_vs_VehVeh", ddsruv, "idela")
dex_only <- results_table("group_DexVeh_vs_VehVeh", ddsruv, "dex")
pred_only <- results_table("group_PredVeh_vs_VehVeh", ddsruv, "pred")
dex_idela <- results_table("group_DexIdela_vs_VehVeh", ddsruv, "di")
pred_idela <- results_table("group_PredIdela_vs_VehVeh", ddsruv, "pi")

sum_table <-
  cbind(idela, dex_only[, c(2:5)]) %>%
  cbind(., pred_only[, c(2:5)]) %>%
  cbind(., dex_idela[, c(2:5)]) %>%
  cbind(., pred_idela[, c(2:5)])

add_geneids <- function(genelist) {
  genelist$symbol <- mapIds(org.Hs.eg.db, keys=substr(row.names(genelist), 1, 15), column="SYMBOL", keytype="ENSEMBL", multi
Vals="first")
  genelist$entrez <- mapIds(org.Hs.eg.db, keys=substr(row.names(genelist), 1, 15), column="ENTREZID", keytype="ENSEMBL", mul
tiVals="first")
  genelist$genename <- mapIds(org.Hs.eg.db, keys=substr(row.names(genelist), 1, 15), column="GENENAME", keytype="ENSEMBL", m
ultiVals="first")
  #genelist <- genelist %>% drop_na(log2FoldChange)
  return(genelist)
}

sum_table <- add_geneids(sum_table)
```

```
## 'select()' returned 1:many mapping between keys and columns
## 'select()' returned 1:many mapping between keys and columns
## 'select()' returned 1:many mapping between keys and columns
```

```
sum_tbl <- sum_table %>%
  dplyr::select(0,(length(sum_table)-2):length(sum_table), everything()) %>%
  rownames_to_column(var = "Ensembl_geneid") %>%
  as_tibble()
```

Making bar charts of the number of genes which are regulated in each treatment condition for visualization of results

```
sum_lng <- sum_tbl %>%
  pivot_longer(cols = !c(1:5), names_to = c("treat", "stat"), names_sep = "_", values_to = "value") %>%
  pivot_wider(names_from = "stat", values_from = "value") %>%
  replace_na(list(pval = 1, adjp = 1)) %>%
  mutate(treat = factor(treat, c("idela", "pred", "pi", "dex", "di")))

sum_lng %>%
  group_by(treat) %>%
  summarise(Up = sum(adjp <= 0.01 & log2FC > 0), Down = sum(adjp <= 0.01 & log2FC < 0)) %>%
  pivot_longer(cols = c("Up", "Down"), names_to = "Regulation", values_to = "Number") %>%
  ggplot(aes(treat, Number, fill = Regulation)) +
  geom_col(width = 0.8, position=position_dodge(0.9)) +
  scale_fill_manual(values=c('blue','red')) +
  scale_x_discrete(breaks=c("idela", "pred", "pi", "dex", "di"), labels=c("Idela", "Pred", "Pred +\nIdela", "Dex", "Dex +\nI
dela")) +
  theme_bw() +
  ylab("Number of Genes Regulated") +
  theme(axis.title.x=element_blank(), axis.title.y = element_text(face = "bold"), legend.position = c(0.12, 0.85))
```

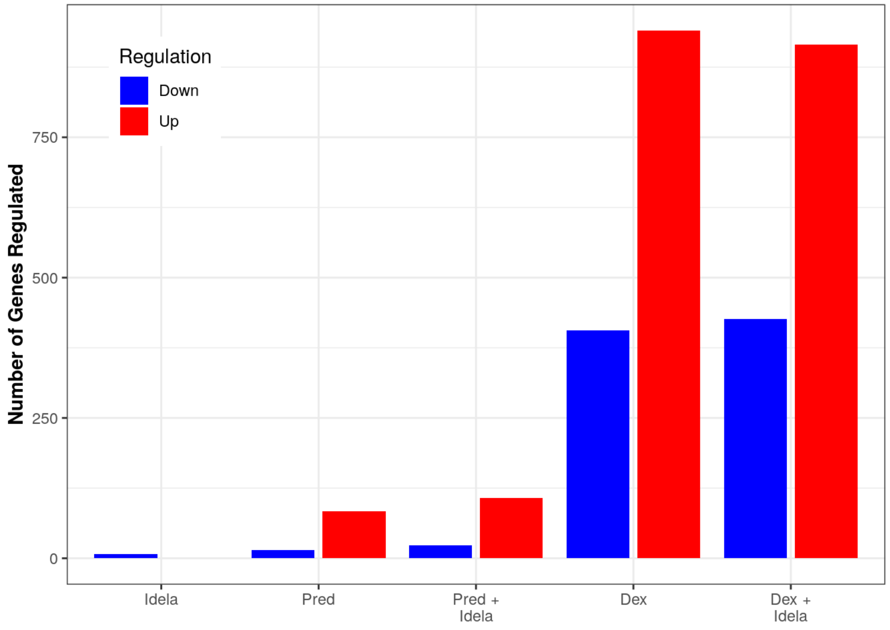

```
  # ggsave("pt_samples_up_down_summary_20221123.pdf", width = 5, height = 4)
  # ggsave("pt_samples_up_down_summary_20221123.png", width = 5, height = 4)
  # ggsave("pt_samples_up_down_summary_20221123.svg", width = 5, height = 4)
```

Plotting Dex vs. Dex + Idela

```
dex_vs_di_all <- sum_tbl %>%
  dplyr::filter(dex_adjp <= 0.01 | di_adjp <= 0.01) %>%
  dplyr::filter(abs(dex_log2FC) < 10 & abs(di_log2FC) < 10) %>%
  ggplot(aes(x = dex_log2FC, y = di_log2FC)) +
  geom_point() +
  xlim(-6, 10) + ylim(-8, 10) +
  geom_abline(slope = 1, intercept = 0, color = "red") +
  geom_smooth(method = "lm", color = "black") +
  stat_cor(label.x.npc = "left", label.y.npc = "top") +
  stat_regline_equation(label.y = 8, aes(label = ..eq.label..)) +
  xlab("Dex Log2FC") +
  ylab("Dex+Idela Log2FC") +
  theme_bw()

dex_vs_di_all
```

```
## Warning: The dot-dot notation (`..eq.label..`) was deprecated in ggplot2 3.4.0.
## ℹ Please use `after_stat(eq.label)` instead.
```

```
## `geom_smooth()` using formula = 'y ~ x'
```

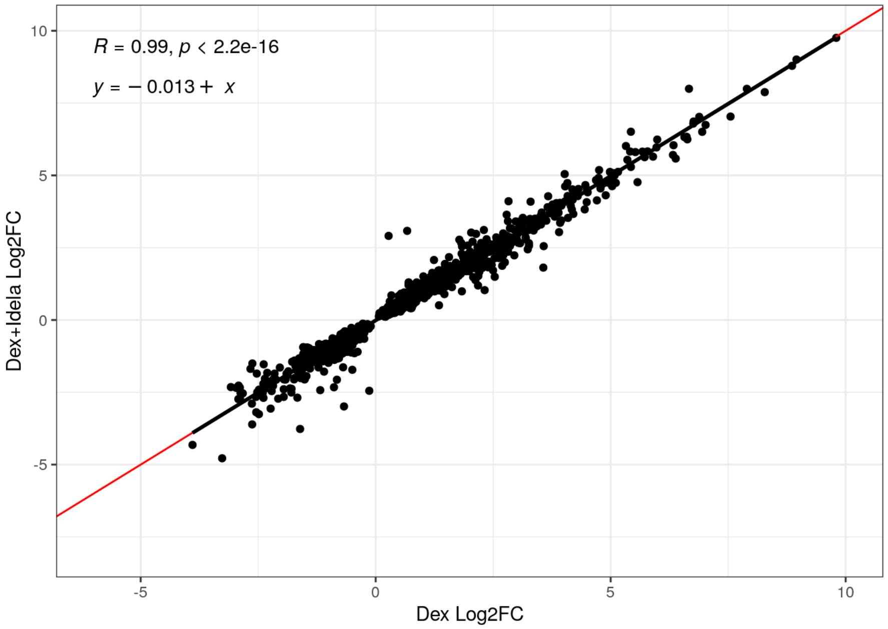

Plotting Pred vs Pred + Idela

```
pred_vs_pi_all <- sum_tbl %>%
  dplyr::filter(pred_adjp <= 0.01 | pi_adjp <= 0.01) %>%
  dplyr::filter(abs(pred_log2FC) < 10 & abs(pi_log2FC) < 10) %>%
  ggplot(aes(x = pred_log2FC, y = pi_log2FC)) +
  geom_point() +
  xlim(-4, 8) + ylim(-7, 10) +
  geom_abline(slope = 1, intercept = 0, color = "red") +
  geom_smooth(method = "lm", color = "black") +
  stat_cor(label.x.npc = "left", label.y.npc = "top") +
  stat_regline_equation(label.y = 8, aes(label = ..eq.label..)) +
  xlab("Pred Log2FC") +
  ylab("Pred+Idela Log2FC") +
  theme_bw()

pred_vs_pi_all
```

```
## `geom_smooth()` using formula = 'y ~ x'
```

```
## Warning: Removed 2 rows containing non-finite values (`stat_smooth()`).
```

```
## Warning: Removed 2 rows containing non-finite values (`stat_cor()`).
```

```
## Warning: Removed 2 rows containing non-finite values
## (`stat_regline_equation()`).
```

```
## Warning: Removed 2 rows containing missing values (`geom_point()`).
```

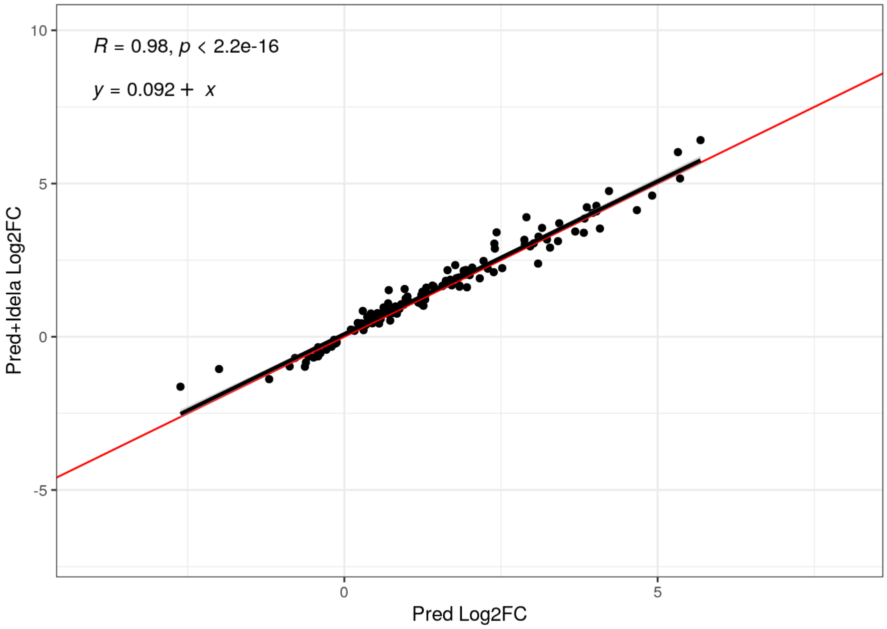

no difference with idela added to dex or pred

#### Repeat the analysis trying to compare sensitive vs. resistant samples

##### Add in a new variable (GCsenivitity) for resistant vs. sensitive

```
test_sample <- list.files(dir)
all.equal(test_sample, tidy_tbl$Sample)
```

```
## [1] TRUE
```

```
sample_table$names <- test_sample

# add a variable for sensitivity
sample_table$GCsensitivity <- sample_table$patient
sample_table <- sample_table %>%
  mutate(GCsensitivity = fct_collapse(GCsensitivity,
                                      sensitive = c("MAP014", "MAP015" , "MAP016", "MAP019", "MAP031"),
                                      resistant = c("MAP010", "MAP020")))

sample_table$files <- file.path(dir, sample_table$names, "quant.sf")
file.exists(sample_table$files)
```

```
##  [1] TRUE TRUE TRUE TRUE TRUE TRUE TRUE TRUE TRUE TRUE TRUE TRUE TRUE TRUE TRUE
## [16] TRUE TRUE TRUE TRUE TRUE TRUE TRUE TRUE TRUE TRUE TRUE TRUE TRUE TRUE TRUE
## [31] TRUE TRUE TRUE TRUE TRUE TRUE TRUE TRUE TRUE TRUE TRUE TRUE TRUE TRUE TRUE
## [46] TRUE TRUE TRUE TRUE TRUE TRUE TRUE TRUE TRUE TRUE TRUE TRUE TRUE TRUE TRUE
## [61] TRUE TRUE TRUE TRUE TRUE
```

```
# Note that the order of factors had to be changed to put Vehicle first, since it is the control condition

sample_table <- sample_table %>%
  mutate(treatment = as_factor(treatment)) %>%
  mutate(gc = factor(gc, levels = c("Veh", "Dex", "Pred"))) %>%
  mutate(idela = factor(idela, levels = c("Veh", "Idela")))

se <- tximeta(sample_table)
```

```
## importing quantifications
```

```
## reading in files with read_tsv
```

```
## 1 2 3 4 5 6 7 8 9 10 11 12 13 14 15 16 17 18 19 20 21 22 23 24 25 26 27 28 29 30 31 32 33 34 35 36 37 38 39 40 41 42 43 4
4 45 46 47 48 49 50 51 52 53 54 55 56 57 58 59 60 61 62 63 64 65
## found matching transcriptome:
## [ GENCODE - Homo sapiens - release 38 ]
## loading existing TxDb created: 2021-10-11 19:29:18
## loading existing transcript ranges created: 2021-10-11 19:29:20
## fetching genome info for GENCODE
```

```
## Error in .order_seqlevels(chrom_sizes[, "chrom"]) :
##   !anyNA(m32) is not TRUE
```

##### Summarize to gene for gene-level analysis

```
dim(se)
```

```
## [1] 236186     65
```

```
gse <- summarizeToGene(se)
```

```
## loading existing TxDb created: 2021-10-11 19:29:18
```

```
## obtaining transcript-to-gene mapping from database
```

```
## loading existing gene ranges created: 2021-10-11 19:29:47
```

```
## summarizing abundance
```

```
## summarizing counts
```

```
## summarizing length
```

```
dim(gse)
```

```
## [1] 60230    65
```

##### Fit to model with group and RUV, but also taking into account GC sensitivity

Did this similarly to above except for adding in the interaction of GCsensitivity with group in the final design.

```
dds <- DESeqDataSet(gse, design = ~ idela + gc)
```

```
## using counts and average transcript lengths from tximeta
```

```
ddsruv <- dds

ddsruv$W1 <- set$W_1
ddsruv$W2 <- set$W_2

# design(ddsruv) <- ~ W1 + W2 + idela + gc + idela:gc

ddsruv$group <- factor(paste0(ddsruv$gc, ddsruv$idela))
ddsruv$group <- relevel(ddsruv$group, "VehVeh")

ddsruv$GCsensitivity <- relevel(ddsruv$GCsensitivity, "sensitive")

design(ddsruv) <- ~ W1 + W2 + GCsensitivity*group
```

##### Now filter out genes with < 2 reads on average per sample.

if there are 65 samples, that'd be 2 * 65 or 130

```
ddsruv <- ddsruv[ rowSums(counts(ddsruv)) > 130, ]
nrow(ddsruv)
```

```
## [1] 25100
```

#### Perform differential gene expression testing using the new model, taking into account GC sensitivity

```
ddsruv <- DESeq(ddsruv)
```

```
## estimating size factors
```

```
## using 'avgTxLength' from assays(dds), correcting for library size
```

```
## estimating dispersions
```

```
## gene-wise dispersion estimates
```

```
## mean-dispersion relationship
```

```
## final dispersion estimates
```

```
## fitting model and testing
```

```
## 6 rows did not converge in beta, labelled in mcols(object)$betaConv. Use larger maxit argument with nbinomWaldTest
```

```
resultsNames(ddsruv)
```

```
##  [1] "Intercept"
##  [2] "W1"
##  [3] "W2"
##  [4] "GCsensitivity_resistant_vs_sensitive"
##  [5] "group_DexIdela_vs_VehVeh"
##  [6] "group_DexVeh_vs_VehVeh"
##  [7] "group_PredIdela_vs_VehVeh"
##  [8] "group_PredVeh_vs_VehVeh"
##  [9] "group_VehIdela_vs_VehVeh"
## [10] "GCsensitivityresistant.groupDexIdela"
## [11] "GCsensitivityresistant.groupDexVeh"
## [12] "GCsensitivityresistant.groupPredIdela"
## [13] "GCsensitivityresistant.groupPredVeh"
## [14] "GCsensitivityresistant.groupVehIdela"
```

#### Looking at then numbers of genes regulated in each condition, comparing sensitive vs. resistant specimens

##### Dex Alone

```
# this one shows the effect of dex alone on the sensitive samples
ddsruv %>%
  results(contrast = list(c("group_DexVeh_vs_VehVeh")), alpha = 0.01) %>%
  summary()
```

```
##
## out of 25100 with nonzero total read count
## adjusted p-value < 0.01
## LFC > 0 (up)       : 1155, 4.6%
## LFC < 0 (down)     : 796, 3.2%
## outliers [1]       : 0, 0%
## low counts [2]     : 974, 3.9%
## (mean count < 3)
## [1] see 'cooksCutoff' argument of ?results
## [2] see 'independentFiltering' argument of ?results
```

```
dex_sens_res <- ddsruv %>%
  results(contrast = list(c("group_DexVeh_vs_VehVeh")), alpha = 0.01)

# this one shows the effect of dex alone plus the GC sensitivity (aka resistant samples)
ddsruv %>%
  results(contrast = list(c("group_DexVeh_vs_VehVeh", "GCsensitivityresistant.groupDexVeh")), alpha = 0.01) %>%
  summary()
```

```
##
## out of 25100 with nonzero total read count
## adjusted p-value < 0.01
## LFC > 0 (up)       : 26, 0.1%
## LFC < 0 (down)     : 15, 0.06%
## outliers [1]       : 0, 0%
## low counts [2]     : 2920, 12%
## (mean count < 5)
## [1] see 'cooksCutoff' argument of ?results
## [2] see 'independentFiltering' argument of ?results
```

```
dex_resist_res <- ddsruv %>%
  results(contrast = list(c("group_DexVeh_vs_VehVeh", "GCsensitivityresistant.groupDexVeh")), alpha = 0.01)

# this one shows just the interaction genes which are different between the sensitive and resistant samples for dex
ddsruv %>%
  results(contrast = list(c( "GCsensitivityresistant.groupDexVeh")), alpha = 0.01) %>%
  summary()
```

```
##
## out of 25100 with nonzero total read count
## adjusted p-value < 0.01
## LFC > 0 (up)       : 4, 0.016%
## LFC < 0 (down)     : 7, 0.028%
## outliers [1]       : 0, 0%
## low counts [2]     : 0, 0%
## (mean count < 2)
## [1] see 'cooksCutoff' argument of ?results
## [2] see 'independentFiltering' argument of ?results
```

Continuing on with pred:

```
# this one shows the effect of pred alone on the sensitive samples
ddsruv %>%
  results(contrast = list(c("group_PredVeh_vs_VehVeh")), alpha = 0.01) %>%
  summary()
```

```
##
## out of 25100 with nonzero total read count
## adjusted p-value < 0.01
## LFC > 0 (up)       : 103, 0.41%
## LFC < 0 (down)     : 43, 0.17%
## outliers [1]       : 0, 0%
## low counts [2]     : 974, 3.9%
## (mean count < 3)
## [1] see 'cooksCutoff' argument of ?results
## [2] see 'independentFiltering' argument of ?results
```

```
pred_sens_res <- ddsruv %>%
  results(contrast = list(c("group_PredVeh_vs_VehVeh")), alpha = 0.01)

# this one shows the effect of pred alone plus the GC sensitivity (aka resistant samples)
ddsruv %>%
  results(contrast = list(c("group_PredVeh_vs_VehVeh", "GCsensitivityresistant.groupPredVeh")), alpha = 0.01) %>%
  summary()
```

```
##
## out of 25100 with nonzero total read count
## adjusted p-value < 0.01
## LFC > 0 (up)       : 5, 0.02%
## LFC < 0 (down)     : 3, 0.012%
## outliers [1]       : 0, 0%
## low counts [2]     : 0, 0%
## (mean count < 2)
## [1] see 'cooksCutoff' argument of ?results
## [2] see 'independentFiltering' argument of ?results
```

```
pred_resist_res <- ddsruv %>%
  results(contrast = list(c("group_PredVeh_vs_VehVeh", "GCsensitivityresistant.groupPredVeh")), alpha = 0.01)

# this one shows just the interaction genes which are different between the sensitive and resistant samples for pred
ddsruv %>%
  results(contrast = list(c( "GCsensitivityresistant.groupPredVeh")), alpha = 0.01) %>%
  summary()
```

```
##
## out of 25100 with nonzero total read count
## adjusted p-value < 0.01
## LFC > 0 (up)       : 1, 0.004%
## LFC < 0 (down)     : 1, 0.004%
## outliers [1]       : 0, 0%
## low counts [2]     : 0, 0%
## (mean count < 2)
## [1] see 'cooksCutoff' argument of ?results
## [2] see 'independentFiltering' argument of ?results
```

##### Idela alone

```
# this one shows the effect of idela alone on the sensitive samples
ddsruv %>%
  results(contrast = list(c("group_VehIdela_vs_VehVeh")), alpha = 0.01) %>%
  summary()
```

```
##
## out of 25100 with nonzero total read count
## adjusted p-value < 0.01
## LFC > 0 (up)       : 2, 0.008%
## LFC < 0 (down)     : 4, 0.016%
## outliers [1]       : 0, 0%
## low counts [2]     : 0, 0%
## (mean count < 2)
## [1] see 'cooksCutoff' argument of ?results
## [2] see 'independentFiltering' argument of ?results
```

```
idela_sens_res <- ddsruv %>%
  results(contrast = list(c("group_VehIdela_vs_VehVeh")), alpha = 0.01)

# this one shows the effect of idela alone plus the GC sensitivity (aka resistant samples)
ddsruv %>%
  results(contrast = list(c("group_VehIdela_vs_VehVeh", "GCsensitivityresistant.groupVehIdela")), alpha = 0.01) %>%
  summary()
```

```
##
## out of 25100 with nonzero total read count
## adjusted p-value < 0.01
## LFC > 0 (up)       : 3, 0.012%
## LFC < 0 (down)     : 12, 0.048%
## outliers [1]       : 0, 0%
## low counts [2]     : 1947, 7.8%
## (mean count < 4)
## [1] see 'cooksCutoff' argument of ?results
## [2] see 'independentFiltering' argument of ?results
```

```
idela_resist_res <- ddsruv %>%
  results(contrast = list(c("group_VehIdela_vs_VehVeh", "GCsensitivityresistant.groupVehIdela")), alpha = 0.01)

# this one shows just the interaction genes which are different between the sensitive and resistant samples for pred
ddsruv %>%
  results(contrast = list(c( "GCsensitivityresistant.groupVehIdela")), alpha = 0.01) %>%
  summary()
```

```
##
## out of 25100 with nonzero total read count
## adjusted p-value < 0.01
## LFC > 0 (up)       : 0, 0%
## LFC < 0 (down)     : 6, 0.024%
## outliers [1]       : 0, 0%
## low counts [2]     : 0, 0%
## (mean count < 2)
## [1] see 'cooksCutoff' argument of ?results
## [2] see 'independentFiltering' argument of ?results
```

Dex+Idela

```
# this one shows the effect of dex+idela on the sensitive samples
ddsruv %>%
  results(contrast = list(c("group_DexIdela_vs_VehVeh")), alpha = 0.01) %>%
  summary()
```

```
##
## out of 25100 with nonzero total read count
## adjusted p-value < 0.01
## LFC > 0 (up)       : 1185, 4.7%
## LFC < 0 (down)     : 842, 3.4%
## outliers [1]       : 0, 0%
## low counts [2]     : 974, 3.9%
## (mean count < 3)
## [1] see 'cooksCutoff' argument of ?results
## [2] see 'independentFiltering' argument of ?results
```

```
di_sens_res <- ddsruv %>%
  results(contrast = list(c("group_DexIdela_vs_VehVeh")), alpha = 0.01)

# this one shows the effect of dex+idela plus the GC sensitivity (aka resistant samples)
ddsruv %>%
  results(contrast = list(c("group_DexIdela_vs_VehVeh", "GCsensitivityresistant.groupDexIdela")), alpha = 0.01) %>%
  summary()
```

```
##
## out of 25100 with nonzero total read count
## adjusted p-value < 0.01
## LFC > 0 (up)       : 23, 0.092%
## LFC < 0 (down)     : 18, 0.072%
## outliers [1]       : 0, 0%
## low counts [2]     : 0, 0%
## (mean count < 2)
## [1] see 'cooksCutoff' argument of ?results
## [2] see 'independentFiltering' argument of ?results
```

```
di_resist_res <- ddsruv %>%
  results(contrast = list(c("group_DexIdela_vs_VehVeh", "GCsensitivityresistant.groupDexIdela")), alpha = 0.01)

# this one shows just the interaction genes which are different between the sensitive and resistant samples for dex+idela
ddsruv %>%
  results(contrast = list(c( "GCsensitivityresistant.groupDexIdela")), alpha = 0.01) %>%
  summary()
```

```
##
## out of 25100 with nonzero total read count
## adjusted p-value < 0.01
## LFC > 0 (up)       : 3, 0.012%
## LFC < 0 (down)     : 16, 0.064%
## outliers [1]       : 0, 0%
## low counts [2]     : 0, 0%
## (mean count < 2)
## [1] see 'cooksCutoff' argument of ?results
## [2] see 'independentFiltering' argument of ?results
```

Pred+Idela

```
# this one shows the effect of pred+idela on the sensitive samples
ddsruv %>%
  results(contrast = list(c("group_PredIdela_vs_VehVeh")), alpha = 0.01) %>%
  summary()
```

```
##
## out of 25100 with nonzero total read count
## adjusted p-value < 0.01
## LFC > 0 (up)       : 216, 0.86%
## LFC < 0 (down)     : 174, 0.69%
## outliers [1]       : 0, 0%
## low counts [2]     : 3893, 16%
## (mean count < 7)
## [1] see 'cooksCutoff' argument of ?results
## [2] see 'independentFiltering' argument of ?results
```

```
pi_sens_res <- ddsruv %>%
  results(contrast = list(c("group_PredIdela_vs_VehVeh")), alpha = 0.01)

# this one shows the effect of pred+idela plus the GC sensitivity (aka resistant samples)
ddsruv %>%
  results(contrast = list(c("group_PredIdela_vs_VehVeh", "GCsensitivityresistant.groupPredIdela")), alpha = 0.01) %>%
  summary()
```

```
##
## out of 25100 with nonzero total read count
## adjusted p-value < 0.01
## LFC > 0 (up)       : 6, 0.024%
## LFC < 0 (down)     : 10, 0.04%
## outliers [1]       : 0, 0%
## low counts [2]     : 0, 0%
## (mean count < 2)
## [1] see 'cooksCutoff' argument of ?results
## [2] see 'independentFiltering' argument of ?results
```

```
pi_resist_res <- ddsruv %>%
  results(contrast = list(c("group_PredIdela_vs_VehVeh", "GCsensitivityresistant.groupPredIdela")), alpha = 0.01)

# this one shows just the interaction genes which are different between the sensitive and resistant samples for pred+idela
ddsruv %>%
  results(contrast = list(c( "GCsensitivityresistant.groupPredIdela")), alpha = 0.01) %>%
  summary()
```

```
##
## out of 25100 with nonzero total read count
## adjusted p-value < 0.01
## LFC > 0 (up)       : 0, 0%
## LFC < 0 (down)     : 5, 0.02%
## outliers [1]       : 0, 0%
## low counts [2]     : 0, 0%
## (mean count < 2)
## [1] see 'cooksCutoff' argument of ?results
## [2] see 'independentFiltering' argument of ?results
```

##### Make tables of the results for plotting to confirm that these are also the same

Prep and merge tables using the same function used for the first analysis, plus a slightly modified version of the function which takes into account the extra effect of GC sensitivity

```
results_table <- function(res_name, deseq_obj, new_name) {
  df <- results(deseq_obj, name = res_name)
  df <- as.data.frame(df)
  df <- df[,c(1:3, 5:6)]
  colnames(df) <- c("base_mean", paste0(new_name, "_log2FC"), paste0(new_name,"_lfcse"), paste0(new_name, "_pval"), paste0(n
ew_name, "_adjp"))
  new_name <- df
  return(new_name)
}

idela_sens <- results_table("group_VehIdela_vs_VehVeh", ddsruv, "idela_sens")
dex_only_sens <- results_table("group_DexVeh_vs_VehVeh", ddsruv, "dex_sens")
pred_only_sens <- results_table("group_PredVeh_vs_VehVeh", ddsruv, "pred_sens")
dex_idela_sens <- results_table("group_DexIdela_vs_VehVeh", ddsruv, "di_sens")
pred_idela_sens <- results_table("group_PredIdela_vs_VehVeh", ddsruv, "pi_sens")

# different function that can take into account the extra condition effect
results_table_resist <- function(res_name, deseq_obj, new_name) {
  df <- results(deseq_obj, res_name)
  df <- as.data.frame(df)
  df <- df[,c(1:3, 5:6)]
  colnames(df) <- c("base_mean", paste0(new_name, "_log2FC"), paste0(new_name,"_lfcse"), paste0(new_name, "_pval"), paste0(n
ew_name, "_adjp"))
  new_name <- df
  return(new_name)
}

dex_only_resist <- results_table_resist(list(c("group_DexVeh_vs_VehVeh", "GCsensitivityresistant.groupDexVeh")), ddsruv, "de
x_resist")
idela_resist <- results_table_resist(list(c("group_VehIdela_vs_VehVeh", "GCsensitivityresistant.groupVehIdela")), ddsruv, "i
dela_resist")
pred_only_resist <- results_table_resist(list(c("group_PredVeh_vs_VehVeh", "GCsensitivityresistant.groupPredVeh")), ddsruv,
"pred_resist")
dex_idela_resist <- results_table_resist(list(c("group_DexIdela_vs_VehVeh", "GCsensitivityresistant.groupDexIdela")), ddsru
v, "di_resist")
pred_idela_resist <- results_table_resist(list(c("group_PredIdela_vs_VehVeh", "GCsensitivityresistant.groupPredIdela")), dds
ruv, "pi_resist")

sum_table_sens <-
  cbind(idela_sens, dex_only_sens[, c(2:5)]) %>%
  cbind(., pred_only_sens[, c(2:5)]) %>%
  cbind(., dex_idela_sens[, c(2:5)]) %>%
  cbind(., pred_idela_sens[, c(2:5)])

# make a separate sum table for resistant samples
sum_table_resist <-
  cbind(idela_resist, dex_only_resist[, c(2:5)]) %>%
  cbind(., pred_only_resist[, c(2:5)]) %>%
  cbind(., dex_idela_resist[, c(2:5)]) %>%
  cbind(., pred_idela_resist[, c(2:5)])

# make a table with both sensitive and resistant samples together to graph sensitive vs. resistant treatments against each o
ther
sum_table_all <-
  cbind(idela_sens, dex_only_sens[, c(2:5)]) %>%
  cbind(., pred_only_sens[, c(2:5)]) %>%
  cbind(., dex_idela_sens[, c(2:5)]) %>%
  cbind(., pred_idela_sens[, c(2:5)]) %>%
  cbind(., idela_resist[, c(2:5)]) %>%
  cbind(., dex_only_resist[, c(2:5)]) %>%
  cbind(., pred_only_resist[, c(2:5)]) %>%
  cbind(., dex_idela_resist[, c(2:5)]) %>%
  cbind(., pred_idela_resist[, c(2:5)])

add_geneids <- function(genelist) {
  genelist$symbol <- mapIds(org.Hs.eg.db, keys=substr(row.names(genelist), 1, 15), column="SYMBOL", keytype="ENSEMBL", multi
Vals="first")
  genelist$entrez <- mapIds(org.Hs.eg.db, keys=substr(row.names(genelist), 1, 15), column="ENTREZID", keytype="ENSEMBL", mul
tiVals="first")
  genelist$genename <- mapIds(org.Hs.eg.db, keys=substr(row.names(genelist), 1, 15), column="GENENAME", keytype="ENSEMBL", m
ultiVals="first")
  #genelist <- genelist %>% drop_na(log2FoldChange)
  return(genelist)
}
```

```
sum_table_sens <- add_geneids(sum_table_sens)
```

```
## 'select()' returned 1:many mapping between keys and columns
## 'select()' returned 1:many mapping between keys and columns
## 'select()' returned 1:many mapping between keys and columns
```

```
sum_tbl_sens <- sum_table_sens %>%
  dplyr::select(0,(length(sum_table_sens)-2):length(sum_table_sens), everything()) %>%
  rownames_to_column(var = "Ensembl_geneid") %>%
  as_tibble()

sum_table_resist <- add_geneids(sum_table_resist)
```

```
## 'select()' returned 1:many mapping between keys and columns
## 'select()' returned 1:many mapping between keys and columns
## 'select()' returned 1:many mapping between keys and columns
```

```
sum_tbl_resist <- sum_table_resist %>%
  dplyr::select(0,(length(sum_table_resist)-2):length(sum_table_resist), everything()) %>%
  rownames_to_column(var = "Ensembl_geneid") %>%
  as_tibble()

sum_table_all <- add_geneids(sum_table_all)
```

```
## 'select()' returned 1:many mapping between keys and columns
## 'select()' returned 1:many mapping between keys and columns
## 'select()' returned 1:many mapping between keys and columns
```

```
sum_tbl_all <- sum_table_all %>%
  dplyr::select(0,(length(sum_table_all)-2):length(sum_table_all), everything()) %>%
  rownames_to_column(var = "Ensembl_geneid") %>%
  as_tibble()
```

Make bar plots with total number of genes regulated in each treatment condition for sensitive samples

```
sum_lng_sens <- sum_tbl_sens %>%
  pivot_longer(cols = !c(1:5), names_to = c("treat", "GCresp", "stat"), names_sep = "_", values_to = "value") %>%
  pivot_wider(names_from = "stat", values_from = "value") %>%
  replace_na(list(pval = 1, adjp = 1)) %>%
  mutate(treat = factor(treat, c("idela", "pred", "pi", "dex", "di")))

sum_lng_sens %>%
  group_by(treat) %>%
  summarise(Up = sum(adjp <= 0.01 & log2FC > 0), Down = sum(adjp <= 0.01 & log2FC < 0)) %>%
  pivot_longer(cols = c("Up", "Down"), names_to = "Regulation", values_to = "Number") %>%
  ggplot(aes(treat, Number, fill = Regulation)) +
  geom_col(width = 0.8, position=position_dodge(0.9)) +
  scale_fill_manual(values=c('blue','red')) +
  scale_x_discrete(breaks=c("idela", "pred", "pi", "dex", "di"), labels=c("Idela", "Pred", "Pred +\nIdela", "Dex", "Dex +\nI
dela")) +
  theme_bw() +
  ylab("Number of Genes Regulated") +
  theme(axis.title.x=element_blank(), axis.title.y = element_text(face = "bold"), legend.position = c(0.12, 0.85))
```

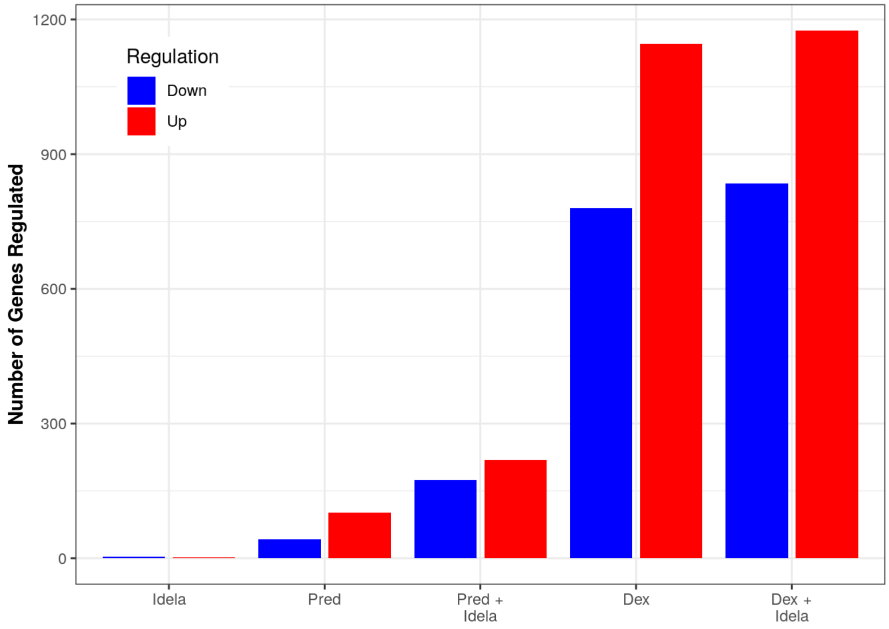

```
ggsave("pts_sens_up_down_summary_20221123.pdf", width = 5, height = 4)
ggsave("pts_sens_up_down_summary_20221123.png", width = 5, height = 4)
ggsave("pts_sens_up_down_summary_20221123.svg", width = 5, height = 4)
```

Make the same type of bar plot for resistant samples

```
sum_lng_resist <- sum_tbl_resist %>%
  pivot_longer(cols = !c(1:5), names_to = c("treat", "GCresp", "stat"), names_sep = "_", values_to = "value") %>%
  pivot_wider(names_from = "stat", values_from = "value") %>%
  replace_na(list(pval = 1, adjp = 1)) %>%
  mutate(treat = factor(treat, c("idela", "pred", "pi", "dex", "di")))

sum_lng_resist %>%
  group_by(treat) %>%
  summarise(Up = sum(adjp <= 0.01 & log2FC > 0), Down = sum(adjp <= 0.01 & log2FC < 0)) %>%
  pivot_longer(cols = c("Up", "Down"), names_to = "Regulation", values_to = "Number") %>%
  ggplot(aes(treat, Number, fill = Regulation)) +
  geom_col(width = 0.8, position=position_dodge(0.9)) +
  scale_fill_manual(values=c('blue','red')) +
  scale_x_discrete(breaks=c("idela", "pred", "pi", "dex", "di"), labels=c("Idela", "Pred", "Pred +\nIdela", "Dex", "Dex +\nI
dela")) +
  theme_bw() +
  ylab("Number of Genes Regulated") +
  theme(axis.title.x=element_blank(), axis.title.y = element_text(face = "bold"), legend.position = c(0.12, 0.85))
```

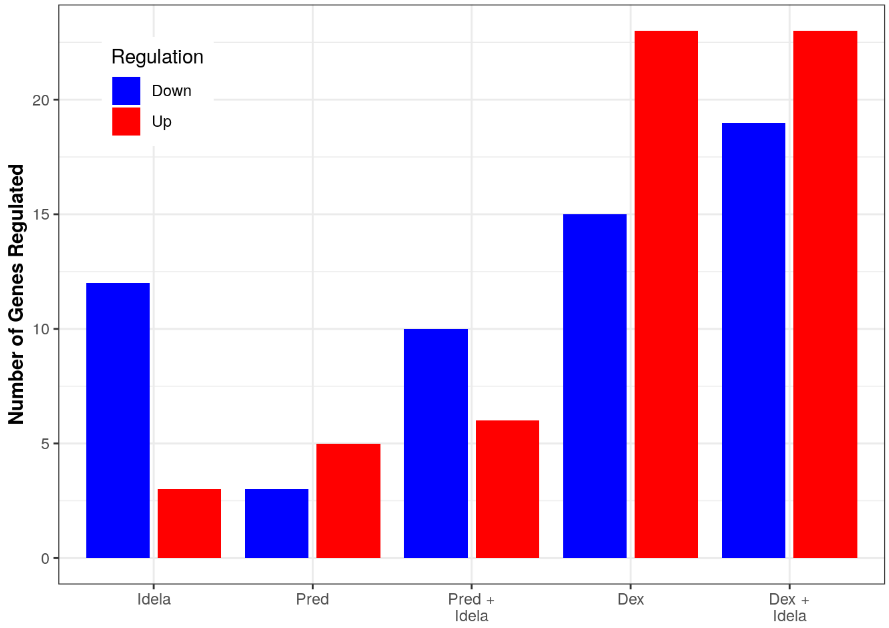

```
ggsave("pts_resist_up_down_summary_20221123.pdf", width = 5, height = 4)
ggsave("pts_resist_up_down_summary_20221123.png", width = 5, height = 4)
ggsave("pts_resist_up_down_summary_20221123.svg", width = 5, height = 4)
```

#### Making plots comparing glucocorticoids +/- idelalisib for GC sensitive and GC resistant specimens

##### Dex vs. Dex+idela for GC sensitive samples

```
sens_dex_idela_plot <- sum_tbl_all %>%
  dplyr::filter(dex_sens_adjp <= 0.01 | di_sens_adjp <= 0.01) %>%
  dplyr::filter(abs(dex_sens_log2FC) < 10 & abs(di_sens_log2FC) < 10) %>%
  ggplot(aes(x = dex_sens_log2FC, y = di_sens_log2FC)) +
  geom_point() +
  xlim(-8, 10) + ylim(-8, 10) +
  geom_abline(slope = 1, intercept = 0, color = "red") +
  geom_smooth(method = "lm", color = "black") +
  stat_cor(label.x.npc = "left", label.y.npc = "top") +
  stat_regline_equation(label.y = 8, aes(label = ..eq.label..)) +
  xlab("Dex Log2FC GC Sensitive") +
  ylab("Dex+Idela Log2FC GC Sensitive") +
  theme_bw()

sens_dex_idela_plot
```

```
## `geom_smooth()` using formula = 'y ~ x'
```

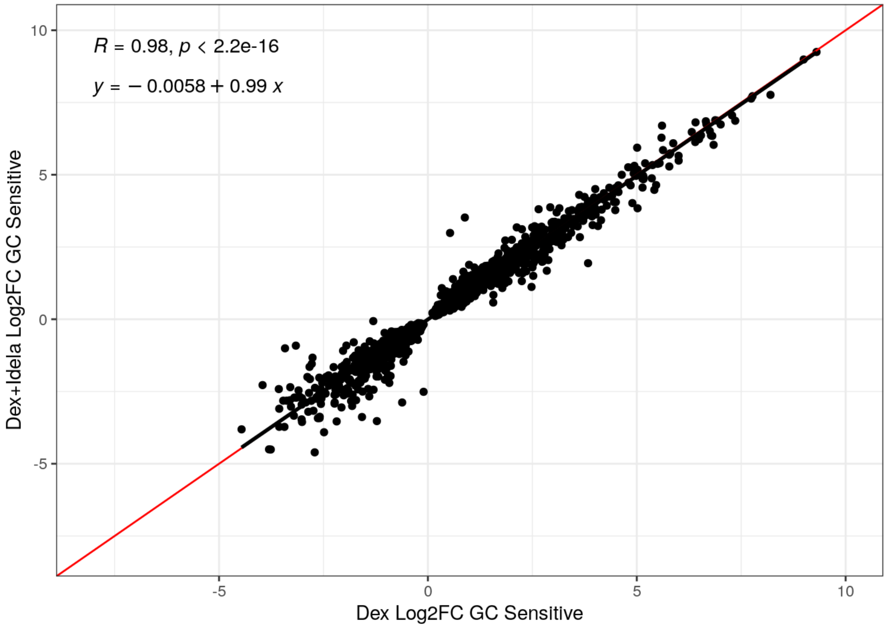

##### Dex vs. Dex+Idela for GC resistant samples

```
resist_dex_idela_plot <- sum_tbl_all %>%
  dplyr::filter(dex_resist_adjp <= 0.05 | di_resist_adjp <= 0.05) %>%
  dplyr::filter(abs(dex_resist_log2FC) < 10 & abs(di_resist_log2FC) < 10) %>%
  ggplot(aes(x = dex_resist_log2FC, y = di_resist_log2FC)) +
  geom_point() +
  xlim(-3, 10) + ylim(-5, 10) +
  geom_abline(slope = 1, intercept = 0, color = "red") +
  geom_smooth(method = "lm", color = "black") +
  stat_cor(label.x.npc = "left", label.y.npc = "top") +
  stat_regline_equation(label.y = 7, aes(label = ..eq.label..)) +
  xlab("Dex Log2FC GC Resistant, p<=0.05") +
  ylab("Dex+Idela Log2FC GC Resistant, p<=0.05") +
  theme_bw()

resist_dex_idela_plot
```

```
## `geom_smooth()` using formula = 'y ~ x'
```

```
## Warning: Removed 3 rows containing non-finite values (`stat_smooth()`).
```

```
## Warning: Removed 3 rows containing non-finite values (`stat_cor()`).
```

```
## Warning: Removed 3 rows containing non-finite values
## (`stat_regline_equation()`).
```

```
## Warning: Removed 3 rows containing missing values (`geom_point()`).
```

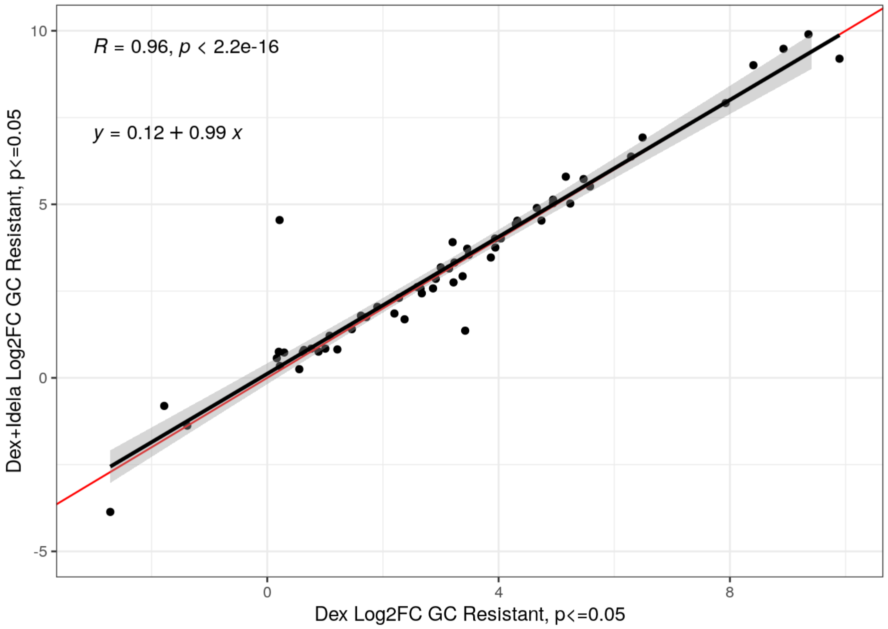

Pred vs. Pred+idela for GC sensitive samples

```
sens_pred_idela_plot <- sum_tbl_all %>%
  dplyr::filter(pred_sens_adjp <= 0.01 | pi_sens_adjp <= 0.01) %>%
  dplyr::filter(abs(pred_sens_log2FC) < 10 & abs(pi_sens_log2FC) < 10) %>%
  ggplot(aes(x = pred_sens_log2FC, y = pi_sens_log2FC)) +
  geom_point() +
  xlim(-5, 10) + ylim(-8, 10) +
  geom_abline(slope = 1, intercept = 0, color = "red") +
  geom_smooth(method = "lm", color = "black") +
  stat_cor(label.x.npc = "left", label.y.npc = "top") +
  stat_regline_equation(label.y = 8, aes(label = ..eq.label..)) +
  xlab("Pred Log2FC GC Sensitive") +
  ylab("Pred+Idela Log2FC GC Sensitive") +
  theme_bw()

sens_pred_idela_plot
```

```
## `geom_smooth()` using formula = 'y ~ x'
```

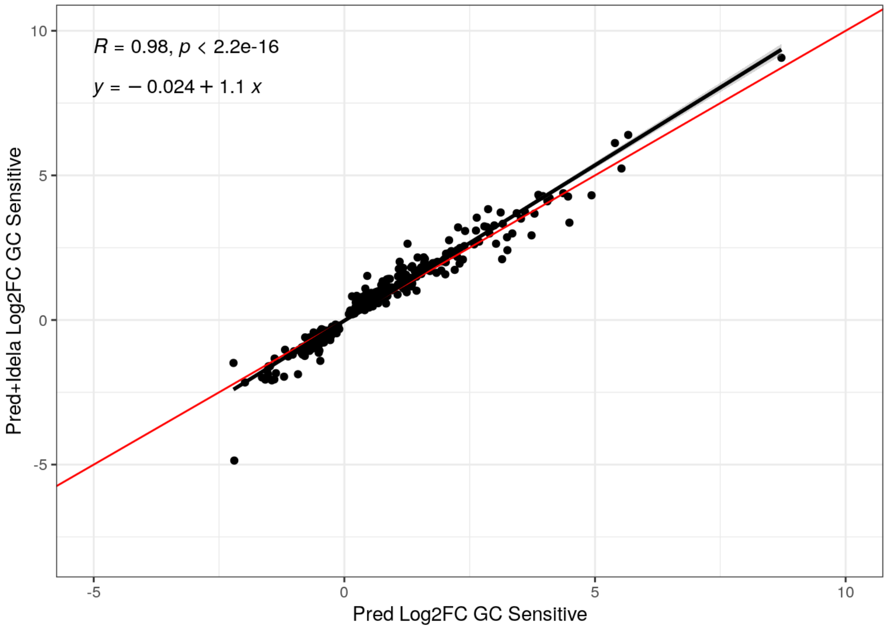

Pred vs. Pred+idela for GC resistant samples

```
resist_pred_idela_plot <- sum_tbl_all %>%
  dplyr::filter(pred_resist_adjp <= 0.1 | pi_resist_adjp <= 0.1) %>%
  dplyr::filter(abs(pred_resist_log2FC) < 10 & abs(pi_resist_log2FC) < 10) %>%
  ggplot(aes(x = pred_resist_log2FC, y = pi_resist_log2FC)) +
  geom_point() +
  xlim(-10, 10) + ylim(-10, 10) +
  geom_abline(slope = 1, intercept = 0, color = "red") +
  geom_smooth(method = "lm", color = "black") +
  stat_cor(label.x.npc = "left", label.y.npc = "top") +
  stat_regline_equation(label.y = 7, aes(label = ..eq.label..)) +
  xlab("Pred Log2FC GC Resistant, p<=0.1") +
  ylab("Pred+Idela Log2FC GC Resistant, p<=0.1") +
  theme_bw()

resist_pred_idela_plot
```

```
## `geom_smooth()` using formula = 'y ~ x'
```

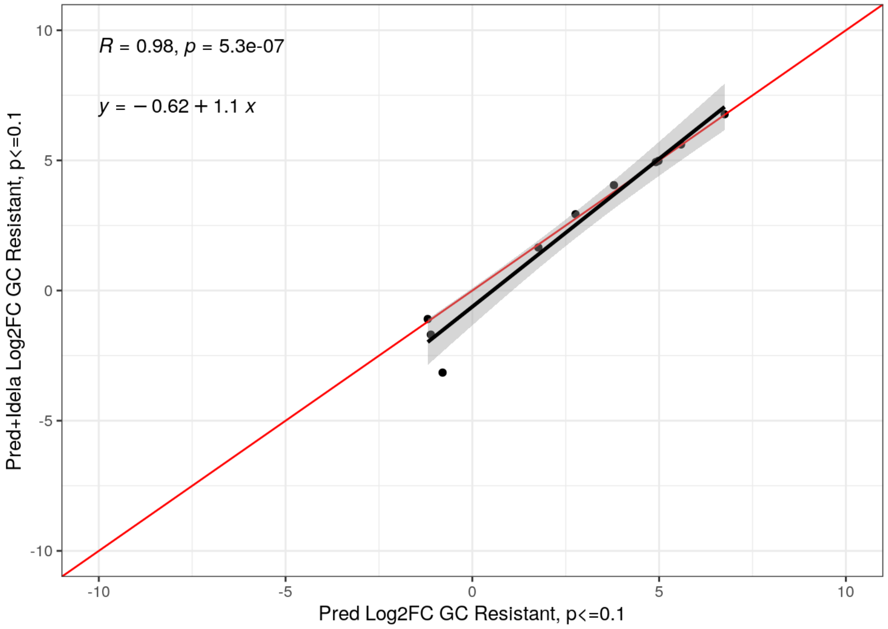

With the new models, it now appears that pred is slightly enhanced by idela in both sensitive and resistant specimens (with the caveat that the resistant specimens have many fewer genes to go off of so take it with a grain of salt), but dex is not.

Saving scatter plots:

```
#ggsave("di_allpts_scatter_bw_20221123.png", plot = dex_vs_di_all, width = 4, height = 4, units = "in")
#ggsave("di_allpts_scatter_bw_20221123.svg", plot = dex_vs_di_all, width = 4, height = 4, units = "in")

#ggsave("pi_allpts_scatter_bw_20221123.png", plot = pred_vs_pi_all, width = 4, height = 4, units = "in")
#ggsave("pi_allpts_scatter_bw_20221123.svg", plot = pred_vs_pi_all, width = 4, height = 4, units = "in")

#ggsave("di_sens_scatter_bw_20221123.png", plot = sens_dex_idela_plot, width = 4, height = 4, units = "in")
#ggsave("di_sens_scatter_bw_20221123.svg", plot = sens_dex_idela_plot, width = 4, height = 4, units = "in")

#ggsave("pi_sens_scatter_bw_20221123.png", plot = sens_pred_idela_plot, width = 4, height = 4, units = "in")
#ggsave("pi_sens_scatter_bw_20221123.svg", plot = sens_pred_idela_plot, width = 4, height = 4, units = "in")

#ggsave("di_resist_scatter_bw_20221123.png", plot = resist_dex_idela_plot, width = 4, height = 4, units = "in")
#ggsave("di_resist_scatter_bw_20221123.svg", plot = resist_dex_idela_plot, width = 4, height = 4, units = "in")

#ggsave("pi_resist_scatter_bw_20221123.png", plot = resist_pred_idela_plot, width = 4, height = 4, units = "in")
#ggsave("pi_resist_scatter_bw_20221123.svg", plot = resist_pred_idela_plot, width = 4, height = 4, units = "in")
```

Session information:

```
sessionInfo()
```

```
## R version 4.1.1 (2021-08-10)
## Platform: x86_64-pc-linux-gnu (64-bit)
## Running under: Ubuntu 18.04.6 LTS
##
## Matrix products: default
## BLAS/LAPACK: /opt/OpenBLAS/lib/libopenblasp-r0.3.3.so
##
## locale:
##  [1] LC_CTYPE=en_US.UTF-8       LC_NUMERIC=C
##  [3] LC_TIME=en_US.UTF-8        LC_COLLATE=en_US.UTF-8
##  [5] LC_MONETARY=en_US.UTF-8    LC_MESSAGES=en_US.UTF-8
##  [7] LC_PAPER=en_US.UTF-8       LC_NAME=C
##  [9] LC_ADDRESS=C               LC_TELEPHONE=C
## [11] LC_MEASUREMENT=en_US.UTF-8 LC_IDENTIFICATION=C
##
## attached base packages:
## [1] parallel  stats4    stats     graphics  grDevices utils     datasets
## [8] methods   base
##
## other attached packages:
##  [1] RUVSeq_1.26.0                edgeR_3.34.1
##  [3] limma_3.48.3                 EDASeq_2.26.1
##  [5] ShortRead_1.50.0             GenomicAlignments_1.28.0
##  [7] Rsamtools_2.8.0              Biostrings_2.60.2
##  [9] XVector_0.32.0               BiocParallel_1.26.2
## [11] ggpubr_0.4.0                 qvalue_2.24.0
## [13] viridis_0.6.2                viridisLite_0.4.1
## [15] forcats_0.5.2                stringr_1.4.1
## [17] dplyr_1.0.10                 purrr_0.3.5
## [19] readr_2.1.3                  tidyr_1.2.1
## [21] tibble_3.1.8                 ggplot2_3.4.0
## [23] tidyverse_1.3.2              ReportingTools_2.32.1
## [25] knitr_1.40                   org.Hs.eg.db_3.13.0
## [27] genefilter_1.74.1            apeglm_1.14.0
## [29] PoiClaClu_1.0.2.1            RColorBrewer_1.1-3
## [31] pheatmap_1.0.12              vsn_3.60.0
## [33] ensembldb_2.16.4             AnnotationFilter_1.16.0
## [35] GenomicFeatures_1.44.2       AnnotationDbi_1.54.1
## [37] rhdf5_2.36.0                 DESeq2_1.32.0
## [39] SummarizedExperiment_1.22.0 Biobase_2.52.0
## [41] MatrixGenerics_1.4.3         matrixStats_0.62.0
## [43] GenomicRanges_1.44.0         GenomeInfoDb_1.28.4
## [45] IRanges_2.26.0               S4Vectors_0.30.2
## [47] BiocGenerics_0.38.0          tximeta_1.10.0
##
## loaded via a namespace (and not attached):
##   [1] rappdirs_0.3.3               rtracklayer_1.52.1
##   [3] AnnotationForge_1.34.1       GGally_2.1.2
##   [5] R.methodsS3_1.8.1            coda_0.19-4
##   [7] ragg_1.2.4                   bit64_4.0.5
##   [9] aroma.light_3.22.0           DelayedArray_0.18.0
##  [11] R.utils_2.11.0               PFAM.db_3.13.0
##  [13] data.table_1.14.4            rpart_4.1-15
##  [15] hwriter_1.3.2.1              KEGGREST_1.32.0
##  [17] RCurl_1.98-1.6               generics_0.1.3
##  [19] preprocessCore_1.54.0        RSQLite_2.2.18
##  [21] bit_4.0.4                    tzdb_0.3.0
##  [23] xml2_1.3.3                   lubridate_1.8.0
##  [25] httpuv_1.6.6                 assertthat_0.2.1
##  [27] gargle_1.2.1                 xfun_0.34
##  [29] tximport_1.20.0              hms_1.1.2
##  [31] jquerylib_0.1.4              evaluate_0.17
##  [33] promises_1.2.0.1             fansi_1.0.3
##  [35] restfulr_0.0.14              progress_1.2.2
##  [37] dbplyr_2.2.1                 readxl_1.4.1
##  [39] Rgraphviz_2.36.0             DBI_1.1.3
##  [41] geneplotter_1.70.0           htmlwidgets_1.5.4
##  [43] reshape_0.8.9                googledrive_2.0.0
##  [45] ellipsis_0.3.2               backports_1.4.1
##  [47] annotate_1.70.0              biomaRt_2.48.3
##  [49] vctrs_0.5.0                  abind_1.4-5
##  [51] cachem_1.0.6                 withr_2.5.0
##  [53] BSgenome_1.60.0              vroom_1.6.0
##  [55] bdsmatrix_1.3-6              checkmate_2.1.0
##  [57] prettyunits_1.1.1            svglite_2.1.0
##  [59] cluster_2.1.2                lazyeval_0.2.2
##  [61] crayon_1.5.2                 labeling_0.4.2
##  [63] pkgconfig_2.0.3              nlme_3.1-152
```

```
##  [65] ProtGenerics_1.24.0          nnet_7.3-16
##  [67] rlang_1.0.6                  lifecycle_1.0.3
##  [69] filelock_1.0.2               affyio_1.62.0
##  [71] BiocFileCache_2.0.0          GOstats_2.58.0
##  [73] modelr_0.1.9                 AnnotationHub_3.0.2
##  [75] dichromat_2.0-0.1            cellranger_1.1.0
##  [77] graph_1.70.0                 Matrix_1.3-4
##  [79] carData_3.0-5                Rhdf5lib_1.14.2
##  [81] reprex_2.0.2                 base64enc_0.1-3
##  [83] googlesheets4_1.0.1          png_0.1-7
##  [85] rjson_0.2.21                 bitops_1.0-7
##  [87] R.oo_1.24.0                  rhdf5filters_1.4.0
##  [89] blob_1.2.3                   jpeg_0.1-9
##  [91] rstatix_0.7.0                ggsignif_0.6.3
##  [93] scales_1.2.1                 memoise_2.0.1
##  [95] GSEABase_1.54.0              magrittr_2.0.3
##  [97] plyr_1.8.7                   zlibbioc_1.38.0
##  [99] compiler_4.1.1               BiocIO_1.2.0
## [101] bbmle_1.0.25                 cli_3.4.1
## [103] affy_1.70.0                  Category_2.58.0
## [105] htmlTable_2.4.0              Formula_1.2-4
## [107] mgcv_1.8-36                  MASS_7.3-54
## [109] tidyselect_1.2.0             stringi_1.7.8
## [111] textshaping_0.3.6            highr_0.9
## [113] emdbook_1.3.12               yaml_2.3.6
## [115] locfit_1.5-9.5               latticeExtra_0.6-29
## [117] grid_4.1.1                   sass_0.4.2
## [119] VariantAnnotation_1.38.0     polynom_1.4-1
## [121] tools_4.1.1                  rstudioapi_0.14
## [123] foreign_0.8-81               gridExtra_2.3
## [125] farver_2.1.1                 digest_0.6.30
## [127] BiocManager_1.30.18          shiny_1.7.3
## [129] Rcpp_1.0.9                   car_3.0-13
## [131] broom_1.0.1                  BiocVersion_3.13.1
## [133] later_1.3.0                  OrganismDbi_1.34.0
## [135] httr_1.4.4                   ggbio_1.40.0
## [137] biovizBase_1.40.0            colorspace_2.0-3
## [139] rvest_1.0.3                  XML_3.99-0.9
## [141] fs_1.5.2                     splines_4.1.1
## [143] RBGL_1.68.0                  systemfonts_1.0.4
## [145] xtable_1.8-4                 jsonlite_1.8.3
## [147] R6_2.5.1                     Hmisc_4.7-0
## [149] pillar_1.8.1                 htmltools_0.5.3
## [151] mime_0.12                    glue_1.6.2
## [153] fastmap_1.1.0                interactiveDisplayBase_1.30.0
## [155] mvtnorm_1.1-3                utf8_1.2.2
## [157] lattice_0.20-44              bslib_0.4.1
## [159] numDeriv_2016.8-1.1          curl_4.3.3
## [161] GO.db_3.13.0                 survival_3.2-11
## [163] rmarkdown_2.17               munsell_0.5.0
## [165] GenomeInfoDbData_1.2.6       haven_2.5.1
## [167] reshape2_1.4.4               gtable_0.3.1
```

### Analysis of synergistic vs. additive samples with RUV, including effector gene analysis

zimmermanjo

1/19/2023

#### This markdown analyzes 2 additive specimens (MAP014 and MAP031) and 2 synergistic specimens (MAP015 and MAP019) to determine differences in gene regulation between these groups with glucocorticoids +/- idelalisib. Each specimen in this analysis included 2 biological replicates, except for MAP014 dexamethasone + idelalisib which had 1 of the replicates fail library preparation. The approach incorporates RUVSeq into the DESeq workflow to identify a set of empirical control genes and use these as controls in our model design. This will follow the DESeq2 workflow for section 2.3 through 3.1 (reading in data), then section 8.2 (RUV), then section 5 on (running differential expression analysis) - workflow here (https://bioconductor.org/packages/release/workflows/vignettes/rnaseqGene/inst/doc/rnaseqGe ruv-with-deseq2).

Import the sample data and select for columns to make a conditions tables - section 2.3

Display the conditions table when you are done. Ensure that the treatment, time, and cell type are all complete.

```
sample_table <- list.files("quants/") %>%
  as_tibble() %>%
  separate(col = "value", into = c("number", "txnum", "replicate", "treatment", "patient")) %>%
  mutate(gc = str_extract(treatment, pattern = "Dex|Pred")) %>%
  mutate(idela = str_extract(treatment, pattern = "Idela")) %>%
  replace_na(list(gc = "Veh", idela = "Veh")) %>%
  mutate(treament = as.factor(treatment), patient = as.factor(patient), gc = as.factor(gc), idela = as.factor(idela))
```

According to the JSON files:

What are the average number of reads per sample and what is the average mapping percentage?

```
dir <- "quants"

Sample <- list.files("quants/")

test <- sapply(list.files(dir), function(x) rjson::fromJSON(file = paste0("quants/", x, "/aux_info/meta_info.json")))

#Output is a list

table <- as.data.frame(t(test))

## There are a few variables with multiple values per observation.  They might be interesting, but we'll select them out
table_filt <- table %>%
  dplyr::select(-quant_errors, -eq_class_properties, -length_classes)

table_filt <- add_column(table_filt, Sample, .before = TRUE)
tidy_tbl <- map_df(table_filt, unlist)

tidy_tbl <- tidy_tbl %>%
  separate(col = Sample, into = c("number", "txnum", "replicate", "treatment", "patient"), remove = FALSE)
```

Average number of reads:

```
ave_reads <- tidy_tbl %>%
  pull(num_processed) %>%
  mean() %>%
  round(0)

ave_reads
```

```
## [1] 43911959
```

```
ggplot(tidy_tbl, aes(Sample, num_mapped, fill = patient)) +
  geom_col() +
  geom_hline(yintercept = mean(tidy_tbl$num_mapped)) +
  theme(axis.text.x = element_text(angle = 45, hjust=1))
```

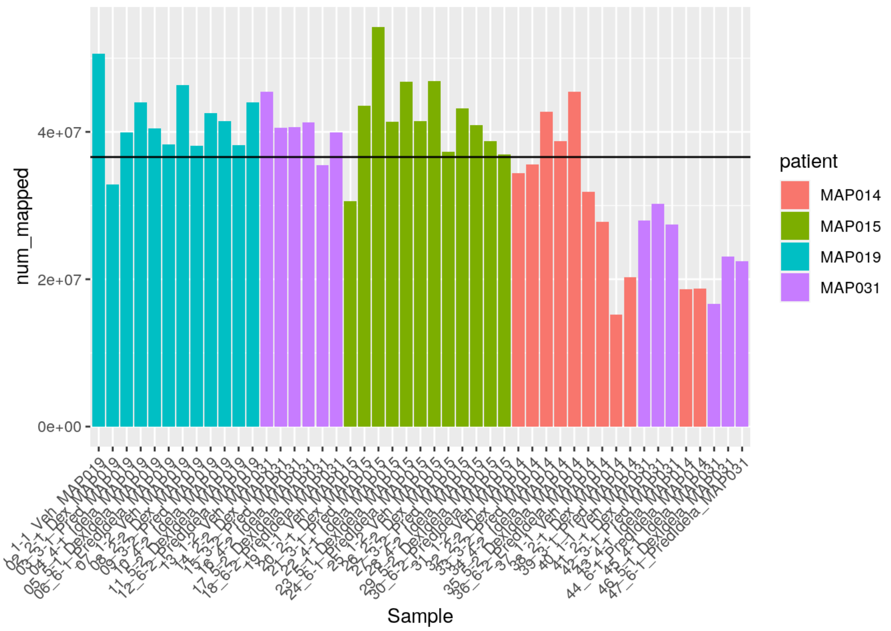

```
ave_mapped <- tidy_tbl %>%
  pull(percent_mapped) %>%
  mean() %>%
  round(0)

ave_mapped
```

```
## [1] 83
```

```
ggplot(tidy_tbl, aes(Sample, percent_mapped, fill = patient)) +
  geom_col() +
  geom_hline(yintercept = mean(tidy_tbl$percent_mapped)) +
  theme(axis.text.x = element_text(angle = 45, hjust=1))
```

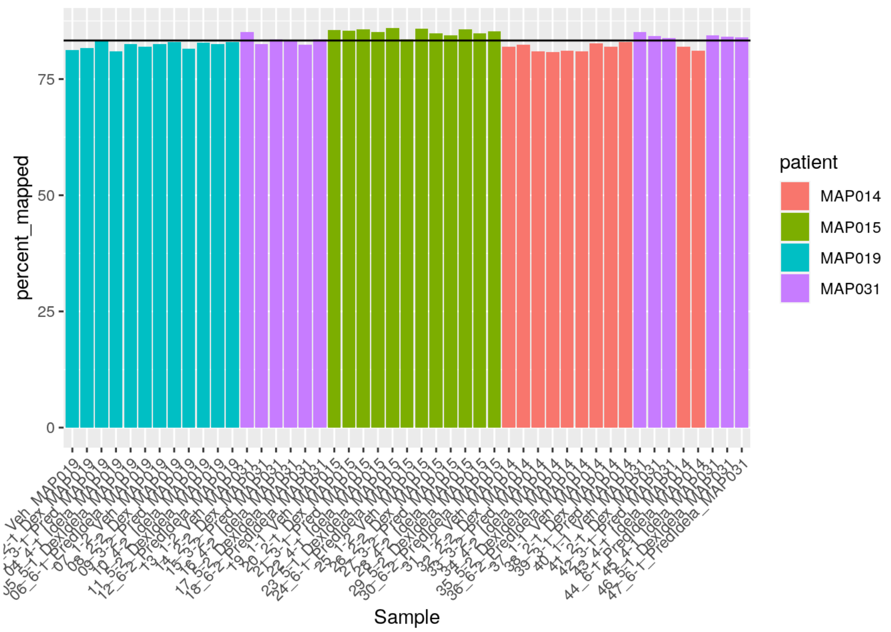

The average number of reads is 4.3911959^{7}

The average percent mapped is 83%

#### Import count tables into R

```
test_sample <- list.files(dir)
all.equal(test_sample, tidy_tbl$Sample)
```

```
## [1] TRUE
```

```
sample_table$names <- test_sample

# add a variable for interaction
sample_table$interaction <- sample_table$patient
sample_table <- sample_table %>%
  mutate(interaction = fct_collapse(interaction,
                                    synergistic = c("MAP015" , "MAP019"),
                                    additive = c("MAP014", "MAP031")))

sample_table$files <- file.path(dir, sample_table$names, "quant.sf")
file.exists(sample_table$files)
```

```
##  [1] TRUE TRUE TRUE TRUE TRUE TRUE TRUE TRUE TRUE TRUE TRUE TRUE TRUE TRUE TRUE
## [16] TRUE TRUE TRUE TRUE TRUE TRUE TRUE TRUE TRUE TRUE TRUE TRUE TRUE TRUE TRUE
## [31] TRUE TRUE TRUE TRUE TRUE TRUE TRUE TRUE TRUE TRUE TRUE TRUE TRUE TRUE TRUE
## [46] TRUE TRUE
```

```
# Note that the order of factors had to be changed to put Vehicle first, since it is the control condition

sample_table <- sample_table %>%
  dplyr::select(-treament) %>%
  mutate(treatment = as_factor(treatment)) %>%
  mutate(gc = factor(gc, levels = c("Veh", "Dex", "Pred"))) %>%
  mutate(idela = factor(idela, levels = c("Veh", "Idela")))

se <- tximeta(sample_table)
```

```
## importing quantifications
```

```
## reading in files with read_tsv
```

```
## 1 2 3 4 5 6 7 8 9 10 11 12 13 14 15 16 17 18 19 20 21 22 23 24 25 26 27 28 29 30 31 32 33 34 35 36 37 38 39 40 41 42 43 4
4 45 46 47
## found matching transcriptome:
## [ GENCODE - Homo sapiens - release 38 ]
## loading existing TxDb created: 2021-10-11 19:29:18
## loading existing transcript ranges created: 2021-10-11 19:29:20
## fetching genome info for GENCODE
```

```
## Error in .order_seqlevels(chrom_sizes[, "chrom"]) :
##   !anyNA(m32) is not TRUE
```

#### Summarize to gene for gene-level analysis

```
dim(se)
```

```
## [1] 236186     47
```

```
gse <- summarizeToGene(se)
```

```
## loading existing TxDb created: 2021-10-11 19:29:18
```

```
## obtaining transcript-to-gene mapping from database
```

```
## loading existing gene ranges created: 2021-10-11 19:29:47
```

```
## summarizing abundance
```

```
## summarizing counts
```

```
## summarizing length
```

```
dim(gse)
```

```
## [1] 60230    47
```

##### Specify the model (formula) into DESeq and assign to object “dds”

Per DESeq2 guide section 8.2, we need to run DESeq and results first without any batch effect to obtain p-values for the analysis.

```
dds <- DESeqDataSet(gse, design = ~ idela + gc)
```

```
## using counts and average transcript lengths from tximeta
```

```
nrow(dds)
```

```
## [1] 60230
```

##### Plot PCA of samples

```
vsd <- vst(dds, blind = FALSE)

plotPCA(vsd, intgroup = c("gc", "idela"))
```

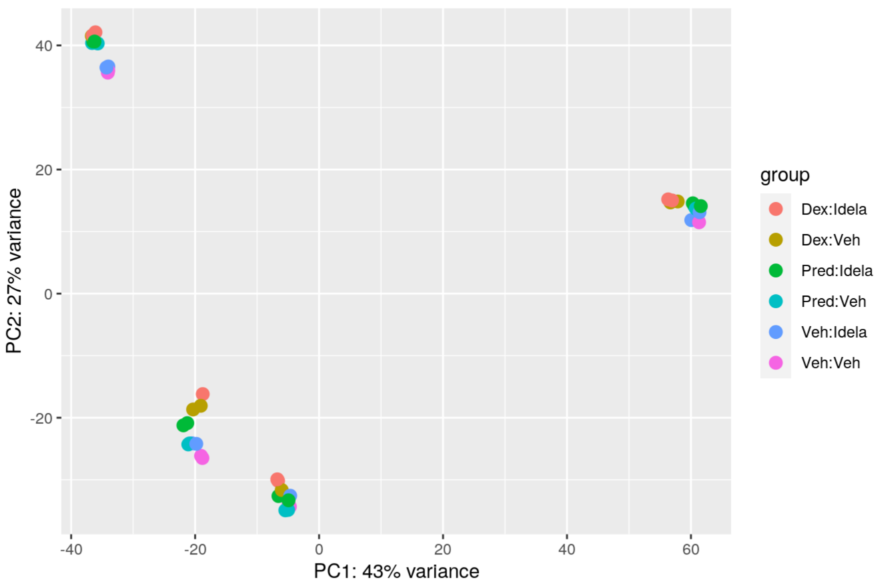

```
pcaData <- plotPCA(vsd, intgroup = c( "gc", "idela", "patient"), returnData = TRUE)

percentVar <- round(100 * attr(pcaData, "percentVar"))

ggplot(pcaData, aes(x = PC1, y = PC2, color = gc, shape = idela)) +
  geom_point(size =3) +
  xlab(paste0("PC1: ", percentVar[1], "% variance")) +
  ylab(paste0("PC2: ", percentVar[2], "% variance")) +
  coord_fixed() +
  ggtitle("PCA with VST data")
```

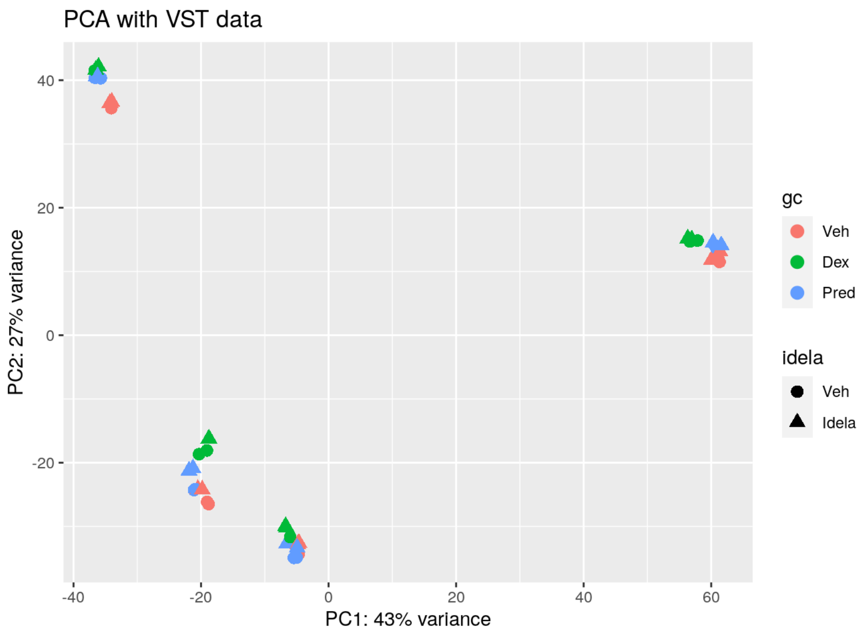

The data likely group best by sample. After that there seems to be a typical progression in PC2 from Veh ==> Pred ==> Dex with grades between that may be due to idela for most of the clusters.

##### Perform differential gene expression testing in order to use RUV:

```
dds <- DESeq(dds)
```

```
## estimating size factors
```

```
## using 'avgTxLength' from assays(dds), correcting for library size
```

```
## estimating dispersions
```

```
## gene-wise dispersion estimates
```

```
## mean-dispersion relationship
```

```
## final dispersion estimates
```

```
## fitting model and testing
```

```
## -- replacing outliers and refitting for 166 genes
## -- DESeq argument 'minReplicatesForReplace' = 7
## -- original counts are preserved in counts(dds)
```

```
## estimating dispersions
```

```
## fitting model and testing
```

```
resultsNames(dds)
```

```
## [1] "Intercept"          "idela_Idela_vs_Veh" "gc_Dex_vs_Veh"
## [4] "gc_Pred_vs_Veh"
```

Creating a results table named “res” to continue with using RUV per section 8.2 in the DESeq2 workflow

```
res <- results(dds)
```

Pulling out empirical control genes:

```
set <- newSeqExpressionSet(counts(dds))
idx  <- rowSums(counts(set) > 5) >= 2
set  <- set[idx, ]
set <- betweenLaneNormalization(set, which="upper")
not.sig <- rownames(res)[which(res$pvalue > .1)]
empirical <- rownames(set)[ rownames(set) %in% not.sig ]
set <- RUVg(set, empirical, k=2)
pData(set)
```

```
##                                  W_1         W_2
## 01_1-1_Veh_MAP019       -0.16298449  0.17551672
## 02_2-1_Dex_MAP019       -0.15181002  0.20039107
## 03_3-1_Pred_MAP019      -0.15254678  0.19118379
## 04_4-1_Idela_MAP019     -0.16505313  0.18216528
## 05_5-1_DexIdela_MAP019  -0.14586269  0.20311504
## 06_6-1_PredIdela_MAP019 -0.15518317  0.19890324
## 07_1-2_Veh_MAP019       -0.16019848  0.17606177
## 08_2-2_Dex_MAP019       -0.14685183  0.19705087
## 09_3-2_Pred_MAP019      -0.15285024  0.19320415
## 10_4-2_Idela_MAP019     -0.16419270  0.18021430
## 11_5-2_DexIdela_MAP019  -0.14939143  0.20323272
## 12_6-2_PredIdela_MAP019 -0.15362373  0.19536332
## 13_1-2_Veh_MAP031        0.23748345  0.06007026
## 14_2-2_Dex_MAP031        0.21762410  0.08929372
## 15_3-2_Pred_MAP031       0.23068603  0.08201865
## 16_4-2_Idela_MAP031      0.23377744  0.06833913
## 17_5-2_DexIdela_MAP031   0.21734459  0.09714503
## 18_6-2_PredIdela_MAP031  0.23172639  0.08808188
## 19_1-1_Veh_MAP015       -0.02641582 -0.18819986
## 20_2-1_Dex_MAP015       -0.02575883 -0.15601721
## 21_3-1_Pred_MAP015      -0.02891894 -0.17036853
## 22_4-1_Idela_MAP015     -0.02692656 -0.16632869
## 23_5-1_DexIdela_MAP015  -0.02506049 -0.15196973
## 24_6-1_PredIdela_MAP015 -0.03500706 -0.16027843
## 25_1-2_Veh_MAP015       -0.02491886 -0.17451070
## 26_2-2_Dex_MAP015       -0.02707737 -0.15739301
## 27_3-2_Pred_MAP015      -0.02776900 -0.16820333
## 28_4-2_Idela_MAP015     -0.02581923 -0.16829321
## 29_5-2_DexIdela_MAP015  -0.02633560 -0.14693439
## 30_6-2_PredIdela_MAP015 -0.02563436 -0.16294303
## 31_1-2_Veh_MAP014       -0.06191984 -0.13493925
## 32_2-2_Dex_MAP014       -0.05243147 -0.09992935
## 33_3-2_Pred_MAP014      -0.06267250 -0.12218992
## 34_4-2_Idela_MAP014     -0.06701611 -0.11801009
## 35_5-2_DexIdela_MAP014  -0.05152982 -0.08615356
## 36_6-2_PredIdela_MAP014 -0.06604224 -0.11101164
## 37_1-1_Veh_MAP014       -0.05379684 -0.13760444
## 38_2-1_Dex_MAP014       -0.04873338 -0.10611438
## 39_3-1_Pred_MAP014      -0.05446721 -0.13037099
## 40_1-1_Veh_MAP031        0.24620542  0.06330853
## 41_2-1_Dex_MAP031        0.23362479  0.08880741
## 42_3-1_Pred_MAP031       0.24003156  0.07941765
## 43_4-1_Idela_MAP014     -0.05717464 -0.12889953
## 44_6-1_PredIdela_MAP014 -0.06016741 -0.11773046
## 45_4-1_Idela_MAP031      0.25346922  0.07130807
## 46_5-1_DexIdela_MAP031   0.23360915  0.09118074
## 47_6-1_PredIdela_MAP031  0.24656010  0.08902041
```

Plotting the factors estimated by RUV:

```
par(mfrow = c(2, 1), mar = c(3,5,3,1))
for (i in 1:2) {
  stripchart(pData(set)[, i] ~ dds$patient, vertical = TRUE, main = paste0("W", i))
  abline(h = 0)
 }
```

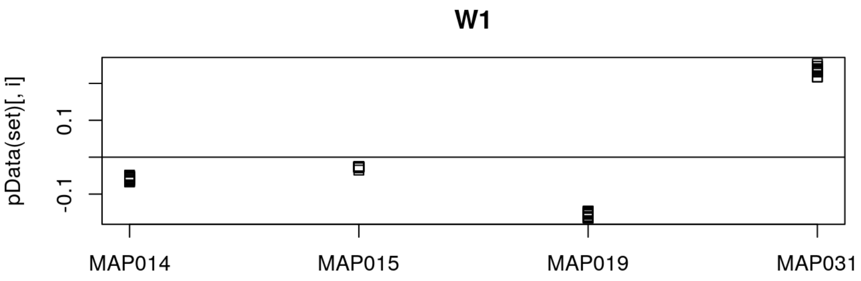

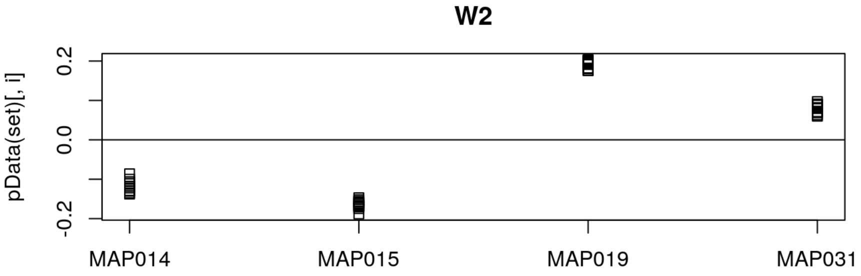

Adding the unwanted variation to the model design

Also adding in grouping variable at this stage and the interaction (additive vs. synergistic, with additive as the reference)

```
ddsruv <- dds

ddsruv$W1 <- set$W_1
ddsruv$W2 <- set$W_2

ddsruv$group <- factor(paste0(ddsruv$gc, ddsruv$idela))
ddsruv$group <- relevel(ddsruv$group, "VehVeh")

ddsruv$interaction <- relevel(ddsruv$interaction, "additive")

design(ddsruv) <- ~ W1 + W2 + interaction*group
```

Now filter out genes with < 2 reads on average per sample.

if there are 47 samples, that'd be 2 * 47 or 94

```
ddsruv <- ddsruv[ rowSums(counts(ddsruv)) > 94, ]
nrow(ddsruv)
```

```
## [1] 24881
```

Perform differential gene expression testing using the new model

```
ddsruv <- DESeq(ddsruv)
```

```
## using pre-existing normalization factors
```

```
## estimating dispersions
```

```
## found already estimated dispersions, replacing these
```

```
## gene-wise dispersion estimates
```

```
## mean-dispersion relationship
```

```
## final dispersion estimates
```

```
## fitting model and testing
```

```
## 1 rows did not converge in beta, labelled in mcols(object)$betaConv. Use larger maxit argument with nbinomWaldTest
```

```
resultsNames(ddsruv)
```

```
##  [1] "Intercept"
##  [2] "W1"
##  [3] "W2"
##  [4] "interaction_synergistic_vs_additive"
##  [5] "group_DexIdela_vs_VehVeh"
##  [6] "group_DexVeh_vs_VehVeh"
##  [7] "group_PredIdela_vs_VehVeh"
##  [8] "group_PredVeh_vs_VehVeh"
##  [9] "group_VehIdela_vs_VehVeh"
## [10] "interactionsynergistic.groupDexIdela"
## [11] "interactionsynergistic.groupDexVeh"
## [12] "interactionsynergistic.groupPredIdela"
## [13] "interactionsynergistic.groupPredVeh"
## [14] "interactionsynergistic.groupVehIdela"
```

##### Let’s look at numbers of genes regulated

###### Dex Alone

```
# this one shows the effect of dex alone on the additive samples
ddsruv %>%
  results(contrast = list(c("group_DexVeh_vs_VehVeh")), alpha = 0.01) %>%
  summary()
```

```
##
## out of 24881 with nonzero total read count
## adjusted p-value < 0.01
## LFC > 0 (up)       : 3192, 13%
## LFC < 0 (down)     : 3003, 12%
## outliers [1]       : 0, 0%
## low counts [2]     : 1930, 7.8%
## (mean count < 4)
## [1] see 'cooksCutoff' argument of ?results
## [2] see 'independentFiltering' argument of ?results
```

```
dex_res <- ddsruv %>%
  results(contrast = list(c("group_DexVeh_vs_VehVeh")), alpha = 0.01)

# this one shows the effect of dex alone plus the interaction (aka synergistic samples)
ddsruv %>%
  results(contrast = list(c("group_DexVeh_vs_VehVeh", "interactionsynergistic.groupDexVeh")), alpha = 0.01) %>%
  summary()
```

```
##
## out of 24881 with nonzero total read count
## adjusted p-value < 0.01
## LFC > 0 (up)       : 2042, 8.2%
## LFC < 0 (down)     : 1972, 7.9%
## outliers [1]       : 0, 0%
## low counts [2]     : 2895, 12%
## (mean count < 5)
## [1] see 'cooksCutoff' argument of ?results
## [2] see 'independentFiltering' argument of ?results
```

```
dex_syn_res <- ddsruv %>%
  results(contrast = list(c("group_DexVeh_vs_VehVeh", "interactionsynergistic.groupDexVeh")), alpha = 0.01)

# this one shows just the interaction genes which are different between the additive and synergistic samples for dex
ddsruv %>%
  results(contrast = list(c( "interactionsynergistic.groupDexVeh")), alpha = 0.01) %>%
  summary()
```

```
##
## out of 24881 with nonzero total read count
## adjusted p-value < 0.01
## LFC > 0 (up)       : 946, 3.8%
## LFC < 0 (down)     : 938, 3.8%
## outliers [1]       : 0, 0%
## low counts [2]     : 6754, 27%
## (mean count < 16)
## [1] see 'cooksCutoff' argument of ?results
## [2] see 'independentFiltering' argument of ?results
```

###### Pred Alone

```
# this one shows the effect of pred alone on the additive samples
ddsruv %>%
  results(contrast = list(c("group_PredVeh_vs_VehVeh")), alpha = 0.01) %>%
  summary()
```

```
##
## out of 24881 with nonzero total read count
## adjusted p-value < 0.01
## LFC > 0 (up)       : 605, 2.4%
## LFC < 0 (down)     : 842, 3.4%
## outliers [1]       : 0, 0%
## low counts [2]     : 9648, 39%
## (mean count < 43)
## [1] see 'cooksCutoff' argument of ?results
## [2] see 'independentFiltering' argument of ?results
```

```
pred_res <- ddsruv %>%
  results(contrast = list(c("group_PredVeh_vs_VehVeh")), alpha = 0.01)

# this one shows the effect of pred alone plus the interaction (aka synergistic samples)
ddsruv %>%
  results(contrast = list(c("group_PredVeh_vs_VehVeh", "interactionsynergistic.groupPredVeh")), alpha = 0.01) %>%
  summary()
```

```
##
## out of 24881 with nonzero total read count
## adjusted p-value < 0.01
## LFC > 0 (up)       : 583, 2.3%
## LFC < 0 (down)     : 532, 2.1%
## outliers [1]       : 0, 0%
## low counts [2]     : 3859, 16%
## (mean count < 7)
## [1] see 'cooksCutoff' argument of ?results
## [2] see 'independentFiltering' argument of ?results
```

```
pred_syn_res <- ddsruv %>%
  results(contrast = list(c("group_PredVeh_vs_VehVeh", "interactionsynergistic.groupPredVeh")), alpha = 0.01)

# this one shows just the interaction genes which are different between the additive and synergistic samples for pred
ddsruv %>%
  results(contrast = list(c( "interactionsynergistic.groupPredVeh")), alpha = 0.01) %>%
  summary()
```

```
##
## out of 24881 with nonzero total read count
## adjusted p-value < 0.01
## LFC > 0 (up)       : 27, 0.11%
## LFC < 0 (down)     : 38, 0.15%
## outliers [1]       : 0, 0%
## low counts [2]     : 7236, 29%
## (mean count < 18)
## [1] see 'cooksCutoff' argument of ?results
## [2] see 'independentFiltering' argument of ?results
```

Idela alone

```
# this one shows the effect of idela alone on the additive samples
ddsruv %>%
  results(contrast = list(c("group_VehIdela_vs_VehVeh")), alpha = 0.01) %>%
  summary()
```

```
##
## out of 24881 with nonzero total read count
## adjusted p-value < 0.01
## LFC > 0 (up)       : 100, 0.4%
## LFC < 0 (down)     : 198, 0.8%
## outliers [1]       : 0, 0%
## low counts [2]     : 11577, 47%
## (mean count < 92)
## [1] see 'cooksCutoff' argument of ?results
## [2] see 'independentFiltering' argument of ?results
```

```
idela_res <- ddsruv %>%
  results(contrast = list(c("group_VehIdela_vs_VehVeh")), alpha = 0.01)

# this one shows the effect of idela alone plus the interaction (aka synergistic samples)
ddsruv %>%
  results(contrast = list(c("group_VehIdela_vs_VehVeh", "interactionsynergistic.groupVehIdela")), alpha = 0.01) %>%
  summary()
```

```
##
## out of 24881 with nonzero total read count
## adjusted p-value < 0.01
## LFC > 0 (up)       : 2, 0.008%
## LFC < 0 (down)     : 8, 0.032%
## outliers [1]       : 0, 0%
## low counts [2]     : 0, 0%
## (mean count < 2)
## [1] see 'cooksCutoff' argument of ?results
## [2] see 'independentFiltering' argument of ?results
```

```
idela_syn_res <- ddsruv %>%
  results(contrast = list(c("group_VehIdela_vs_VehVeh", "interactionsynergistic.groupVehIdela")), alpha = 0.01)

# this one shows just the interaction genes which are different between the additive and synergistic samples for idela
ddsruv %>%
  results(contrast = list(c( "interactionsynergistic.groupVehIdela")), alpha = 0.01) %>%
  summary()
```

```
##
## out of 24881 with nonzero total read count
## adjusted p-value < 0.01
## LFC > 0 (up)       : 0, 0%
## LFC < 0 (down)     : 0, 0%
## outliers [1]       : 0, 0%
## low counts [2]     : 0, 0%
## (mean count < 2)
## [1] see 'cooksCutoff' argument of ?results
## [2] see 'independentFiltering' argument of ?results
```

Dex+Idela

```
# this one shows the effect of dex+idela on the additive samples
ddsruv %>%
  results(contrast = list(c("group_DexIdela_vs_VehVeh")), alpha = 0.01) %>%
  summary()
```

```
##
## out of 24881 with nonzero total read count
## adjusted p-value < 0.01
## LFC > 0 (up)       : 3336, 13%
## LFC < 0 (down)     : 3063, 12%
## outliers [1]       : 0, 0%
## low counts [2]     : 1930, 7.8%
## (mean count < 4)
## [1] see 'cooksCutoff' argument of ?results
## [2] see 'independentFiltering' argument of ?results
```

```
di_res <- ddsruv %>%
  results(contrast = list(c("group_DexIdela_vs_VehVeh")), alpha = 0.01)

# this one shows the effect of dex+idela plus the interaction (aka synergistic samples)
ddsruv %>%
  results(contrast = list(c("group_DexIdela_vs_VehVeh", "interactionsynergistic.groupDexIdela")), alpha = 0.01) %>%
  summary()
```

```
##
## out of 24881 with nonzero total read count
## adjusted p-value < 0.01
## LFC > 0 (up)       : 2106, 8.5%
## LFC < 0 (down)     : 2087, 8.4%
## outliers [1]       : 0, 0%
## low counts [2]     : 2412, 9.7%
## (mean count < 4)
## [1] see 'cooksCutoff' argument of ?results
## [2] see 'independentFiltering' argument of ?results
```

```
di_syn_res <- ddsruv %>%
  results(contrast = list(c("group_DexIdela_vs_VehVeh", "interactionsynergistic.groupDexIdela")), alpha = 0.01)

# this one shows just the interaction genes which are different between the additive and synergistic samples for dex+idela
ddsruv %>%
  results(contrast = list(c( "interactionsynergistic.groupDexIdela")), alpha = 0.01) %>%
  summary()
```

```
##
## out of 24881 with nonzero total read count
## adjusted p-value < 0.01
## LFC > 0 (up)       : 1135, 4.6%
## LFC < 0 (down)     : 1185, 4.8%
## outliers [1]       : 0, 0%
## low counts [2]     : 5789, 23%
## (mean count < 12)
## [1] see 'cooksCutoff' argument of ?results
## [2] see 'independentFiltering' argument of ?results
```

##### Pred+Idela

```
# this one shows the effect of pred+idela on the additive samples
ddsruv %>%
  results(contrast = list(c("group_PredIdela_vs_VehVeh")), alpha = 0.01) %>%
  summary()
```

```
##
## out of 24881 with nonzero total read count
## adjusted p-value < 0.01
## LFC > 0 (up)       : 1657, 6.7%
## LFC < 0 (down)     : 1911, 7.7%
## outliers [1]       : 0, 0%
## low counts [2]     : 5307, 21%
## (mean count < 10)
## [1] see 'cooksCutoff' argument of ?results
## [2] see 'independentFiltering' argument of ?results
```

```
pi_res <- ddsruv %>%
  results(contrast = list(c("group_PredIdela_vs_VehVeh")), alpha = 0.01)

# this one shows the effect of pred+idela plus the interaction (aka synergistic samples)
ddsruv %>%
  results(contrast = list(c("group_PredIdela_vs_VehVeh", "interactionsynergistic.groupPredIdela")), alpha = 0.01) %>%
  summary()
```

```
##
## out of 24881 with nonzero total read count
## adjusted p-value < 0.01
## LFC > 0 (up)       : 725, 2.9%
## LFC < 0 (down)     : 706, 2.8%
## outliers [1]       : 0, 0%
## low counts [2]     : 4824, 19%
## (mean count < 9)
## [1] see 'cooksCutoff' argument of ?results
## [2] see 'independentFiltering' argument of ?results
```

```
pi_syn_res <- ddsruv %>%
  results(contrast = list(c("group_PredIdela_vs_VehVeh", "interactionsynergistic.groupPredIdela")), alpha = 0.01)

# this one shows just the interaction genes which are different between the additive and synergistic samples for pred+idela
ddsruv %>%
  results(contrast = list(c( "interactionsynergistic.groupPredIdela")), alpha = 0.01) %>%
  summary()
```

```
##
## out of 24881 with nonzero total read count
## adjusted p-value < 0.01
## LFC > 0 (up)       : 205, 0.82%
## LFC < 0 (down)     : 178, 0.72%
## outliers [1]       : 0, 0%
## low counts [2]     : 10613, 43%
## (mean count < 62)
## [1] see 'cooksCutoff' argument of ?results
## [2] see 'independentFiltering' argument of ?results
```

##### Make tables to help with plotting comparisons of the treatments in additive vs. synergistic specimens

Prep and merge tables - wrote a function to help with this

```
results_table <- function(res_name, deseq_obj, new_name) {
  df <- results(deseq_obj, name = res_name)
  df <- as.data.frame(df)
  df <- df[,c(1:3, 5:6)]
  colnames(df) <- c("base_mean", paste0(new_name, "_log2FC"), paste0(new_name,"_lfcse"), paste0(new_name, "_pval"), paste0(n
ew_name, "_adjp"))
  new_name <- df
  return(new_name)
}

idela <- results_table("group_VehIdela_vs_VehVeh", ddsruv, "idela")
dex_only <- results_table("group_DexVeh_vs_VehVeh", ddsruv, "dex")
pred_only <- results_table("group_PredVeh_vs_VehVeh", ddsruv, "pred")
dex_idela <- results_table("group_DexIdela_vs_VehVeh", ddsruv, "di")
pred_idela <- results_table("group_PredIdela_vs_VehVeh", ddsruv, "pi")

# modify the function so that it can take into account the extra condition effect
results_table_syn <- function(res_name, deseq_obj, new_name) {
  df <- results(deseq_obj, res_name)
  df <- as.data.frame(df)
  df <- df[,c(1:3, 5:6)]
  colnames(df) <- c("base_mean", paste0(new_name, "_log2FC"), paste0(new_name,"_lfcse"), paste0(new_name, "_pval"), paste0(n
ew_name, "_adjp"))
  new_name <- df
  return(new_name)
}

dex_only_syn <- results_table_syn(list(c("group_DexVeh_vs_VehVeh", "interactionsynergistic.groupDexVeh")), ddsruv, "dex_sy
n")
idela_syn <- results_table_syn(list(c("group_VehIdela_vs_VehVeh", "interactionsynergistic.groupVehIdela")), ddsruv, "idela_s
yn")
pred_only_syn <- results_table_syn(list(c("group_PredVeh_vs_VehVeh", "interactionsynergistic.groupPredVeh")), ddsruv, "pred_
syn")
dex_idela_syn <- results_table_syn(list(c("group_DexIdela_vs_VehVeh", "interactionsynergistic.groupDexIdela")), ddsruv, "di_
syn")
pred_idela_syn <- results_table_syn(list(c("group_PredIdela_vs_VehVeh", "interactionsynergistic.groupPredIdela")), ddsruv,
"pi_syn")

sum_table <-
  cbind(idela, dex_only[, c(2:5)]) %>%
  cbind(., pred_only[, c(2:5)]) %>%
  cbind(., dex_idela[, c(2:5)]) %>%
  cbind(., pred_idela[, c(2:5)])

# make a separate sum table for synergistic samples
sum_table_syn <-
  cbind(idela_syn, dex_only_syn[, c(2:5)]) %>%
  cbind(., pred_only_syn[, c(2:5)]) %>%
  cbind(., dex_idela_syn[, c(2:5)]) %>%
  cbind(., pred_idela_syn[, c(2:5)])

# make a table with both additive and synergistic samples together to graph synergistic vs. additive treatments against each
other
sum_table_all <-
  cbind(idela, dex_only[, c(2:5)]) %>%
  cbind(., pred_only[, c(2:5)]) %>%
  cbind(., dex_idela[, c(2:5)]) %>%
  cbind(., pred_idela[, c(2:5)]) %>%
  cbind(., idela_syn[, c(2:5)]) %>%
  cbind(., dex_only_syn[, c(2:5)]) %>%
  cbind(., pred_only_syn[, c(2:5)]) %>%
  cbind(., dex_idela_syn[, c(2:5)]) %>%
  cbind(., pred_idela_syn[, c(2:5)])

add_geneids <- function(genelist) {
  genelist$symbol <- mapIds(org.Hs.eg.db, keys=substr(row.names(genelist), 1, 15), column="SYMBOL", keytype="ENSEMBL", multi
Vals="first")
  genelist$entrez <- mapIds(org.Hs.eg.db, keys=substr(row.names(genelist), 1, 15), column="ENTREZID", keytype="ENSEMBL", mul
tiVals="first")
  genelist$genename <- mapIds(org.Hs.eg.db, keys=substr(row.names(genelist), 1, 15), column="GENENAME", keytype="ENSEMBL", m
ultiVals="first")
  #genelist <- genelist %>% drop_na(log2FoldChange)
  return(genelist)
}
```

```
sum_table <- add_geneids(sum_table)
```

```
## 'select()' returned 1:many mapping between keys and columns
## 'select()' returned 1:many mapping between keys and columns
## 'select()' returned 1:many mapping between keys and columns
```

```
sum_tbl_add <- sum_table %>%
  dplyr::select(0,(length(sum_table)-2):length(sum_table), everything()) %>%
  rownames_to_column(var = "Ensembl_geneid") %>%
  as_tibble()

sum_table_syn <- add_geneids(sum_table_syn)
```

```
## 'select()' returned 1:many mapping between keys and columns
## 'select()' returned 1:many mapping between keys and columns
## 'select()' returned 1:many mapping between keys and columns
```

```
sum_tbl_syn <- sum_table_syn %>%
  dplyr::select(0,(length(sum_table_syn)-2):length(sum_table_syn), everything()) %>%
  rownames_to_column(var = "Ensembl_geneid") %>%
  as_tibble()

sum_table_all <- add_geneids(sum_table_all)
```

```
## 'select()' returned 1:many mapping between keys and columns
## 'select()' returned 1:many mapping between keys and columns
## 'select()' returned 1:many mapping between keys and columns
```

```
sum_tbl_all <- sum_table_all %>%
  dplyr::select(0,(length(sum_table_all)-2):length(sum_table_all), everything()) %>%
  rownames_to_column(var = "Ensembl_geneid") %>%
  as_tibble()
```

Making bar plots of the total number of genes regulated in each treatment condition for the additive specimens

```
sum_lng_add <- sum_tbl_add %>%
  pivot_longer(cols = !c(1:5), names_to = c("treat", "stat"), names_sep = "_", values_to = "value") %>%
  pivot_wider(names_from = "stat", values_from = "value") %>%
  replace_na(list(pval = 1, adjp = 1)) %>%
  mutate(treat = factor(treat, c("idela", "pred", "pi", "dex", "di")))

sum_lng_add %>%
  group_by(treat) %>%
  summarise(Up = sum(adjp <= 0.01 & log2FC > 0), Down = sum(adjp <= 0.01 & log2FC < 0)) %>%
  pivot_longer(cols = c("Up", "Down"), names_to = "Regulation", values_to = "Number") %>%
  ggplot(aes(treat, Number, fill = Regulation)) +
  geom_col(width = 0.8, position=position_dodge(0.9)) +
  scale_fill_manual(values=c('blue','red')) +
  scale_x_discrete(breaks=c("idela", "pred", "pi", "dex", "di"), labels=c("Idela", "Pred", "Pred +\nIdela", "Dex", "Dex +\nI
dela")) +
  theme_bw() +
  ylab("Number of Genes Regulated") +
  theme(axis.title.x=element_blank(), axis.title.y = element_text(face = "bold"), legend.position = c(0.12, 0.85))
```

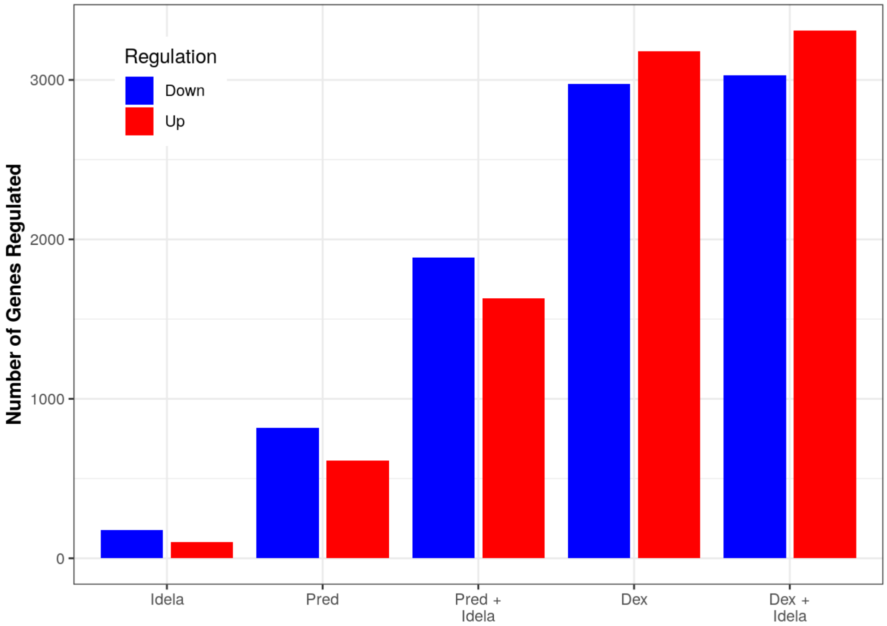

```
# ggsave("pts_add_up_down_summary_20221123.pdf", width = 5, height = 4)
# ggsave("pts_add_up_down_summary_20221123.png", width = 5, height = 4)
# ggsave("pts_add_up_down_summary_20221123.svg", width = 5, height = 4)
```

Making bar plots of the total number of genes regulated in each treatment condition for the synergistic specimens

```
sum_lng_syn <- sum_tbl_syn %>%
  pivot_longer(cols = !c(1:5), names_to = c("treat", "combo", "stat"), names_sep = "_", values_to = "value") %>%
  pivot_wider(names_from = "stat", values_from = "value") %>%
  replace_na(list(pval = 1, adjp = 1)) %>%
  mutate(treat = factor(treat, c("idela", "pred", "pi", "dex", "di")))

sum_lng_syn %>%
  group_by(treat) %>%
  summarise(Up = sum(adjp <= 0.01 & log2FC > 0), Down = sum(adjp <= 0.01 & log2FC < 0)) %>%
  pivot_longer(cols = c("Up", "Down"), names_to = "Regulation", values_to = "Number") %>%
  ggplot(aes(treat, Number, fill = Regulation)) +
  geom_col(width = 0.8, position=position_dodge(0.9)) +
  scale_fill_manual(values=c('blue','red')) +
  scale_x_discrete(breaks=c("idela", "pred", "pi", "dex", "di"), labels=c("Idela", "Pred", "Pred +\nIdela", "Dex", "Dex +\nI
dela")) +
  theme_bw() +
  ylab("Number of Genes Regulated") +
  theme(axis.title.x=element_blank(), axis.title.y = element_text(face = "bold"), legend.position = c(0.12, 0.85))
```

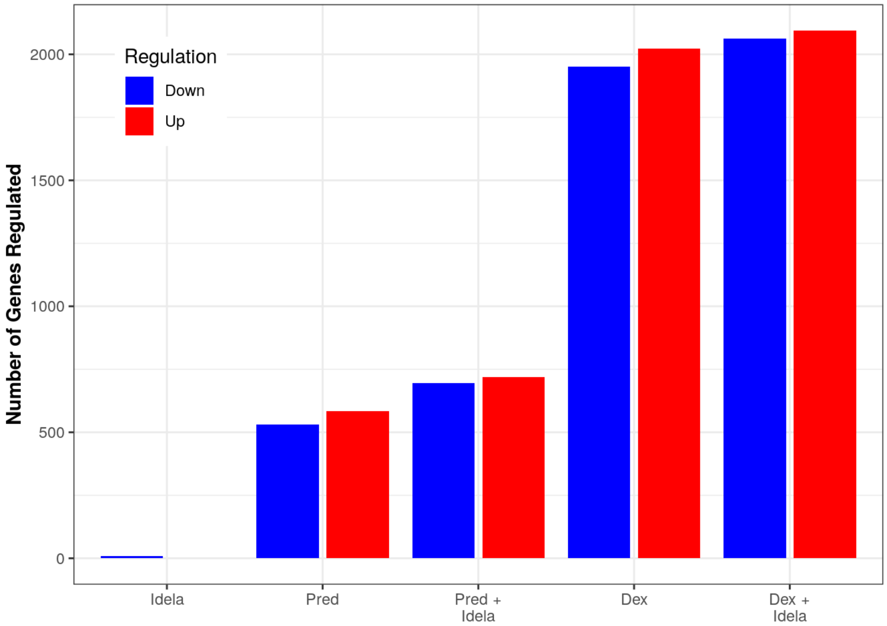

```
# ggsave("pts_syn_up_down_summary_20221123.pdf", width = 5, height = 4)
# ggsave("pts_syn_up_down_summary_20221123.png", width = 5, height = 4)
# ggsave("pts_syn_up_down_summary_20221123.svg", width = 5, height = 4)
```

#### Making plots of the treatment conditions

##### Dex vs. Dex+idela for additive samples

```
add_dex_idela_plot <- sum_tbl_add %>%
  dplyr::filter(dex_adjp <= 0.01 | di_adjp <= 0.01) %>%
  dplyr::filter(abs(dex_log2FC) < 10 & abs(di_log2FC) < 10) %>%
  ggplot(aes(x = dex_log2FC, y = di_log2FC)) +
  geom_point() +
  xlim(-8, 10) + ylim(-10, 10) +
  geom_abline(slope = 1, intercept = 0, color = "red") +
  geom_smooth(method = "lm", color = "black") +
  stat_cor(label.x.npc = "left", label.y.npc = "top") +
  stat_regline_equation(label.y = 7.5, aes(label = after_stat(eq.label))) +
  xlab("Dex Log2FC Additive") +
  ylab("Dex+Idela Log2FC Additive") +
  theme_bw()

add_dex_idela_plot
```

```
## `geom_smooth()` using formula = 'y ~ x'
```

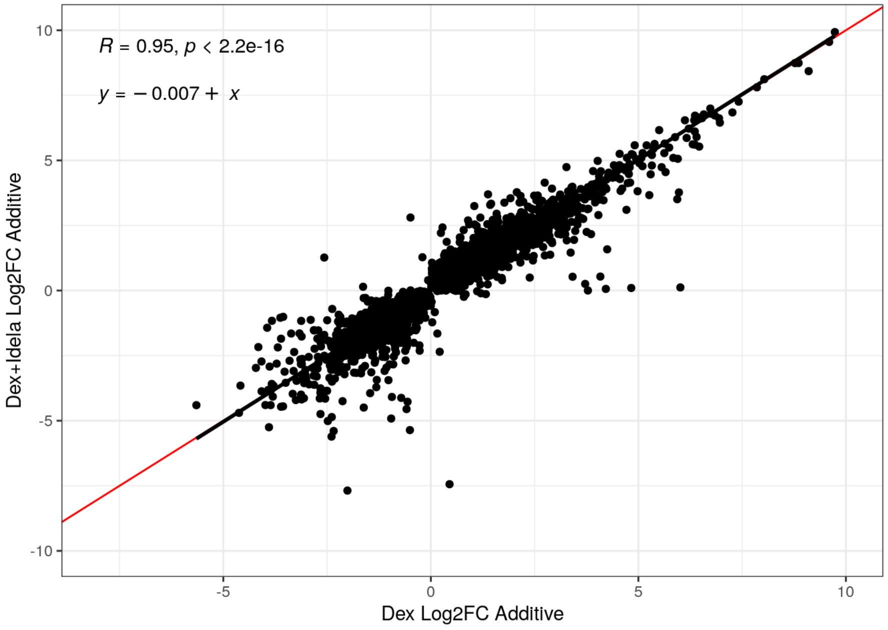

##### Dex vs. Dex+Idela for synergistic samples

```
syn_dex_idela_plot <- sum_tbl_syn %>%
  dplyr::filter(dex_syn_adjp <= 0.01 | di_syn_adjp <= 0.01) %>%
  dplyr::filter(abs(dex_syn_log2FC) < 10 & abs(di_syn_log2FC) < 10) %>%
  ggplot(aes(x = dex_syn_log2FC, y = di_syn_log2FC)) +
  geom_point() +
  xlim(-8, 10) + ylim(-10, 10) +
  geom_abline(slope = 1, intercept = 0, color = "red") +
  geom_smooth(method = "lm", color = "black") +
  stat_cor(label.x.npc = "left", label.y.npc = "top") +
  stat_regline_equation(label.y = 7.5, aes(label = after_stat(eq.label))) +
  xlab("Dex Log2FC Synergistic") +
  ylab("Dex+Idela Log2FC Synergistic") +
  theme_bw()

syn_dex_idela_plot
```

```
## `geom_smooth()` using formula = 'y ~ x'
```

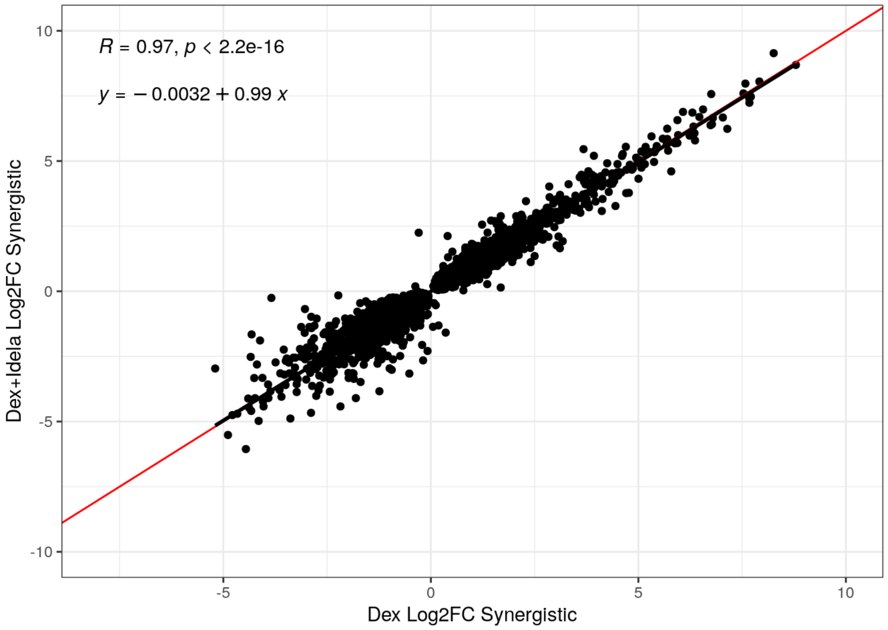

Pred vs. Pred+idela for additive samples

```
add_pred_idela_plot <- sum_tbl_add %>%
  dplyr::filter(pred_adjp <= 0.01 | pi_adjp <= 0.01) %>%
  dplyr::filter(abs(pred_log2FC) < 10 & abs(pi_log2FC) < 10) %>%
  ggplot(aes(x = pred_log2FC, y = pi_log2FC)) +
  geom_point() +
  xlim(-5, 10) + ylim(-5, 10) +
  geom_abline(slope = 1, intercept = 0, color = "red") +
  geom_smooth(method = "lm", color = "black") +
  stat_cor(label.x.npc = "left", label.y.npc = "top") +
  stat_regline_equation(label.y = 8, aes(label = after_stat(eq.label))) +
  xlab("Pred Log2FC Additive") +
  ylab("Pred+Idela Log2FC Additive") +
  theme_bw()

add_pred_idela_plot
```

```
## `geom_smooth()` using formula = 'y ~ x'
```

```
## Warning: Removed 1 rows containing non-finite values (`stat_smooth()`).
```

```
## Warning: Removed 1 rows containing non-finite values (`stat_cor()`).
```

```
## Warning: Removed 1 rows containing non-finite values
## (`stat_regline_equation()`).
```

```
## Warning: Removed 1 rows containing missing values (`geom_point()`).
```

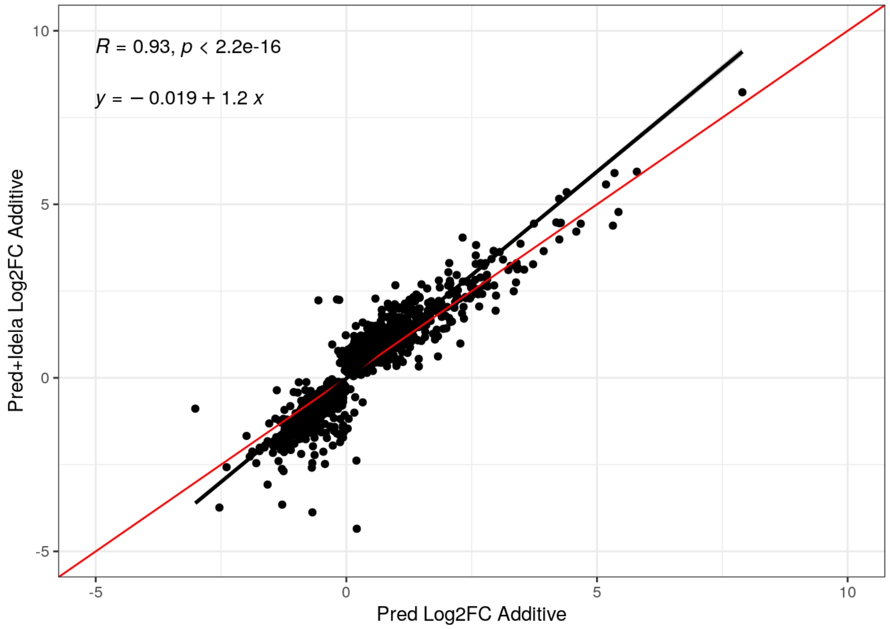

Pred vs. Pred+idela for synergistic samples

```
syn_pred_idela_plot <- sum_tbl_syn %>%
  dplyr::filter(pred_syn_adjp <= 0.01 | pi_syn_adjp <= 0.01) %>%
  dplyr::filter(abs(pred_syn_log2FC) < 10 & abs(pi_syn_log2FC) < 10) %>%
  ggplot(aes(x = pred_syn_log2FC, y = pi_syn_log2FC)) +
  geom_point() +
  xlim(-5, 10) + ylim(-5, 10) +
  geom_abline(slope = 1, intercept = 0, color = "red") +
  geom_smooth(method = "lm", color = "black") +
  stat_cor(label.x.npc = "left", label.y.npc = "top") +
  stat_regline_equation(label.y = 8, aes(label = after_stat(eq.label))) +
  xlab("Pred Log2FC Synergistic") +
  ylab("Pred+Idela Log2FC Synergistic") +
  theme_bw()

syn_pred_idela_plot
```

```
## `geom_smooth()` using formula = 'y ~ x'
```

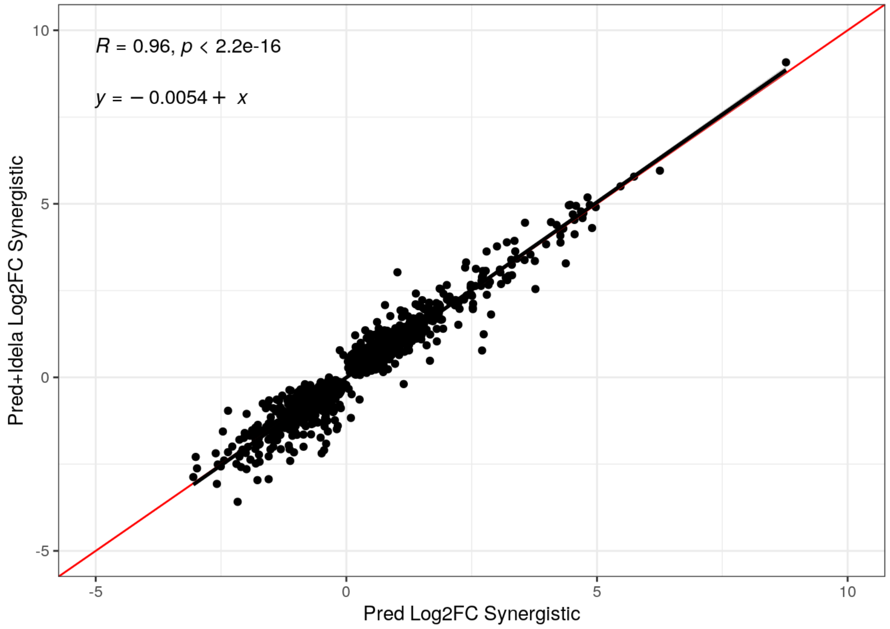

#### Saving the scatter plots as png and svg

Dex vs. dex+idela

```
#di_scatter_plots <- grid.arrange(add_dex_idela_plot, syn_dex_idela_plot, nrow = 1)

#ggsave("di_add_vs_syn_scatter_bw_20221123.png", plot = di_scatter_plots, width = 8, height = 4, units = "in")
#ggsave("di_add_vs_syn_scatter_bw_20221123.svg", plot = di_scatter_plots, width = 8, height = 4, units = "in")

# ggsave("di_add_scatter_bw_20221123.png", plot = add_dex_idela_plot, width = 4, height = 4, units = "in")
# ggsave("di_add_scatter_bw_20221123.svg", plot = add_dex_idela_plot, width = 4, height = 4, units = "in")

# ggsave("di_syn_scatter_bw_20221123.png", plot = syn_dex_idela_plot, width = 4, height = 4, units = "in")
# ggsave("di_syn_scatter_bw_20221123.svg", plot = syn_dex_idela_plot, width = 4, height = 4, units = "in")
```

Pred vs. Pred+idela

```
# pi_scatter_plots <- grid.arrange(add_pred_idela_plot, syn_pred_idela_plot, nrow = 1)

# ggsave("pi_add_vs_syn_scatter_bw.png", plot = pi_scatter_plots, width = 8, height = 4, units = "in")
# ggsave("pi_add_vs_syn_scatter_bw.svg", plot = pi_scatter_plots, width = 8, height = 4, units = "in")

# ggsave("pi_add_scatter_bw_20221123.png", plot = add_pred_idela_plot, width = 4, height = 4, units = "in")
# ggsave("pi_add_scatter_bw_20221123.svg", plot = add_pred_idela_plot, width = 4, height = 4, units = "in")

# ggsave("pi_syn_scatter_bw_20221123.png", plot = syn_pred_idela_plot, width = 4, height = 4, units = "in")
# ggsave("pi_syn_scatter_bw_20221123.svg", plot = syn_pred_idela_plot, width = 4, height = 4, units = "in")
```

#### Let's compare dex and pred alone for additive vs. synergistic samples to take a look at how GC-induced gene regulation compares in the additive vs. synergistic specimens

Dex alone, additive vs. synergistic specimens

```
dex_add_vs_syn_plot <- sum_tbl_all %>%
  dplyr::filter(dex_adjp <= 0.01 & dex_syn_adjp <= 0.01) %>%
  dplyr::filter(abs(dex_log2FC) < 10 & abs(dex_syn_log2FC) < 10) %>%
  ggplot(aes(x = dex_log2FC, y = dex_syn_log2FC)) +
  geom_point() +
  xlim(-8, 10) + ylim(-8, 10) +
  geom_abline(slope = 1, intercept = 0, color = "red") +
  geom_smooth(method = "lm", color = "black") +
  stat_cor(label.x.npc = "left", label.y.npc = "top") +
  stat_regline_equation(label.y = 7.5, aes(label = after_stat(eq.label))) +
  xlab("Dex Log2FC Additive") +
  ylab("Dex Log2FC Synergistic") +
  theme_bw()

dex_add_vs_syn_plot
```

```
## `geom_smooth()` using formula = 'y ~ x'
```

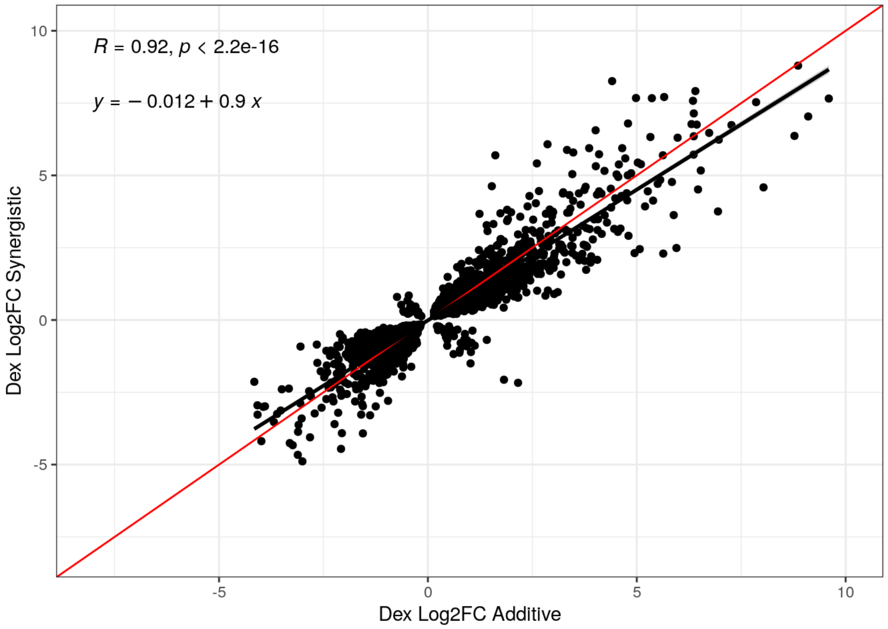

Pred alone, additive vs. synergistic specimens

```
pred_add_vs_syn_plot <- sum_tbl_all %>%
  dplyr::filter(pred_adjp <= 0.01 & pred_syn_adjp <= 0.01) %>%
  dplyr::filter(abs(pred_log2FC) < 10 & abs(pred_syn_log2FC) < 10) %>%
  ggplot(aes(x = pred_log2FC, y = pred_syn_log2FC)) +
  geom_point() +
  xlim(-5, 10) + ylim(-5, 10) +
  geom_abline(slope = 1, intercept = 0, color = "red") +
  geom_smooth(method = "lm", color = "black") +
  stat_cor(label.x.npc = "left", label.y.npc = "top") +
  stat_regline_equation(label.y = 8, aes(label = after_stat(eq.label))) +
  xlab("Pred Log2FC Additive") +
  ylab("Pred Log2FC Synergistic") +
  theme_bw()

pred_add_vs_syn_plot
```

```
## `geom_smooth()` using formula = 'y ~ x'
```

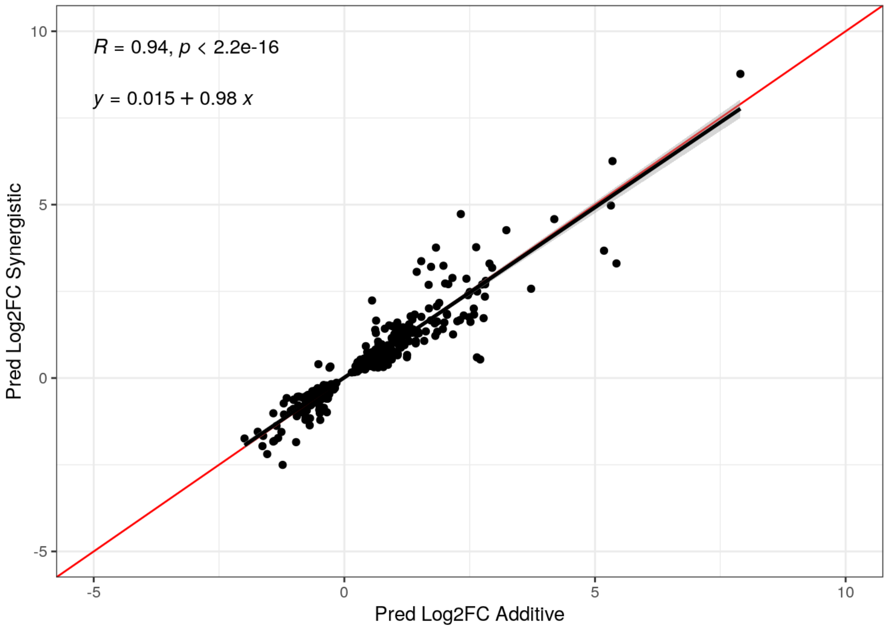

Dex/pred alone for additive vs. synergistic

```
# dexpred_scatter_plots <- grid.arrange(dex_add_vs_syn_plot, pred_add_vs_syn_plot, nrow = 1)

# ggsave("dexpred_add_vs_syn_scatter_bw.png", plot = dexpred_scatter_plots, width = 8, height = 4, units = "in")
# ggsave("dexpred_add_vs_syn_scatter_bw.svg", plot = dexpred_scatter_plots, width = 8, height = 4, units = "in")

# ggsave("dex_add_vs_syn_scatter_bw_20230119.png", plot = dex_add_vs_syn_plot, width = 4, height = 4, units = "in")
# ggsave("dex_add_vs_syn_scatter_bw_20230119.svg", plot = dex_add_vs_syn_plot, width = 4, height = 4, units = "in")

# ggsave("pred_add_vs_syn_scatter_bw_20230119.png", plot = pred_add_vs_syn_plot, width = 4, height = 4, units = "in")
# ggsave("pred_add_vs_syn_scatter_bw_20230119.svg", plot = pred_add_vs_syn_plot, width = 4, height = 4, units = "in")
```

Summary - the only condition where idela seems to enhance gene regulation is additive specimens with pred.

I'll repeat the analysis with all 4 specimens, not splitting into additive vs. synergistic, to compare these results to the GC sensitive specimen results (since all 4 of these specimens are GC sensitive) and ensure that the results are consistent with when I analyzed all specimens together and subset into GC sensitive vs. resistant.

```
ddsruv <- dds

ddsruv$W1 <- set$W_1
ddsruv$W2 <- set$W_2

ddsruv$group <- factor(paste0(ddsruv$gc, ddsruv$idela))
ddsruv$group <- relevel(ddsruv$group, "VehVeh")

design(ddsruv) <- ~ W1 + W2 + group
```

Re-run DESeq with this new design to re-estimate parameters and results.

```
ddsruv <- DESeq(ddsruv)
```

```
## using pre-existing normalization factors
```

```
## estimating dispersions
```

```
## found already estimated dispersions, replacing these
```

```
## gene-wise dispersion estimates
```

```
## mean-dispersion relationship
```

```
## final dispersion estimates
```

```
## fitting model and testing
```

```
resultsNames(ddsruv)
```

```
## [1] "Intercept"                 "W1"
## [3] "W2"                        "group_DexIdela_vs_VehVeh"
## [5] "group_DexVeh_vs_VehVeh"    "group_PredIdela_vs_VehVeh"
## [7] "group_PredVeh_vs_VehVeh"   "group_VehIdela_vs_VehVeh"
```

##### Now I will filter out genes with < 2 reads on average per sample.

if there are 65 samples, that'd be 2 * 47 or 94

```
ddsruv <- ddsruv[ rowSums(counts(ddsruv)) > 94, ]
nrow(ddsruv)
```

```
## [1] 24881
```

#### Now attempting to create results tables

##### Dex Alone

```
dex_res <- results(ddsruv, name = "group_DexVeh_vs_VehVeh", independentFiltering = TRUE, alpha = 0.01)

summary(dex_res)
```

```
##
## out of 24881 with nonzero total read count
## adjusted p-value < 0.01
## LFC > 0 (up)       : 2220, 8.9%
## LFC < 0 (down)     : 1987, 8%
## outliers [1]       : 0, 0%
## low counts [2]     : 483, 1.9%
## (mean count < 2)
## [1] see 'cooksCutoff' argument of ?results
## [2] see 'independentFiltering' argument of ?results
```

##### Pred Alone

```
pred_res <- results(ddsruv, name = "group_PredVeh_vs_VehVeh", independentFiltering = TRUE, alpha = 0.01)

summary(pred_res)
```

```
##
## out of 24881 with nonzero total read count
## adjusted p-value < 0.01
## LFC > 0 (up)       : 341, 1.4%
## LFC < 0 (down)     : 318, 1.3%
## outliers [1]       : 0, 0%
## low counts [2]     : 6271, 25%
## (mean count < 14)
## [1] see 'cooksCutoff' argument of ?results
## [2] see 'independentFiltering' argument of ?results
```

##### Idela Alone

```
idela_res <- results(ddsruv, name = "group_VehIdela_vs_VehVeh", independentFiltering = TRUE, alpha = 0.01)

summary(idela_res)
```

```
##
## out of 24881 with nonzero total read count
## adjusted p-value < 0.01
## LFC > 0 (up)       : 1, 0.004%
## LFC < 0 (down)     : 4, 0.016%
## outliers [1]       : 0, 0%
## low counts [2]     : 0, 0%
## (mean count < 2)
## [1] see 'cooksCutoff' argument of ?results
## [2] see 'independentFiltering' argument of ?results
```

Dex + Idela

```
di_res <- results(ddsruv, name = "group_DexIdela_vs_VehVeh", independentFiltering = TRUE, alpha = 0.01)

summary(di_res)
```

```
##
## out of 24881 with nonzero total read count
## adjusted p-value < 0.01
## LFC > 0 (up)       : 2254, 9.1%
## LFC < 0 (down)     : 2033, 8.2%
## outliers [1]       : 0, 0%
## low counts [2]     : 0, 0%
## (mean count < 2)
## [1] see 'cooksCutoff' argument of ?results
## [2] see 'independentFiltering' argument of ?results
```

Pred + Idela

```
pi_res <- results(ddsruv, name = "group_PredIdela_vs_VehVeh", independentFiltering = TRUE, alpha = 0.01)

summary(pi_res)
```

```
##
## out of 24881 with nonzero total read count
## adjusted p-value < 0.01
## LFC > 0 (up)       : 801, 3.2%
## LFC < 0 (down)     : 877, 3.5%
## outliers [1]       : 0, 0%
## low counts [2]     : 3859, 16%
## (mean count < 7)
## [1] see 'cooksCutoff' argument of ?results
## [2] see 'independentFiltering' argument of ?results
```

##### Continue with making tables and plots to compare gene regulation between conditions

###### Prep and merge tables - wrote a function to help with this

```
results_table <- function(res_name, deseq_obj, new_name) {
  df <- results(deseq_obj, name = res_name)
  df <- as.data.frame(df)
  df <- df[,c(1:3, 5:6)]
  colnames(df) <- c("base_mean", paste0(new_name, "_log2FC"), paste0(new_name,"_lfcse"), paste0(new_name, "_pval"), paste0(n
ew_name, "_adjp"))
  new_name <- df
  return(new_name)
}

idela <- results_table("group_VehIdela_vs_VehVeh", ddsruv, "idela")
dex_only <- results_table("group_DexVeh_vs_VehVeh", ddsruv, "dex")
pred_only <- results_table("group_PredVeh_vs_VehVeh", ddsruv, "pred")
dex_idela <- results_table("group_DexIdela_vs_VehVeh", ddsruv, "di")
pred_idela <- results_table("group_PredIdela_vs_VehVeh", ddsruv, "pi")

sum_table <-
  cbind(idela, dex_only[, c(2:5)]) %>%
  cbind(., pred_only[, c(2:5)]) %>%
  cbind(., dex_idela[, c(2:5)]) %>%
  cbind(., pred_idela[, c(2:5)])

add_geneids <- function(genelist) {
  genelist$symbol <- mapIds(org.Hs.eg.db, keys=substr(row.names(genelist), 1, 15), column="SYMBOL", keytype="ENSEMBL", multi
Vals="first")
  genelist$entrez <- mapIds(org.Hs.eg.db, keys=substr(row.names(genelist), 1, 15), column="ENTREZID", keytype="ENSEMBL", mul
tiVals="first")
  genelist$genename <- mapIds(org.Hs.eg.db, keys=substr(row.names(genelist), 1, 15), column="GENENAME", keytype="ENSEMBL", m
ultiVals="first")
  #genelist <- genelist %>% drop_na(log2FoldChange)
  return(genelist)
}

sum_table <- add_geneids(sum_table)
```

```
## 'select()' returned 1:many mapping between keys and columns
## 'select()' returned 1:many mapping between keys and columns
## 'select()' returned 1:many mapping between keys and columns
```

```
sum_tbl <- sum_table %>%
  dplyr::select(0,(length(sum_table)-2):length(sum_table), everything()) %>%
  rownames_to_column(var = "Ensembl_geneid") %>%
  as_tibble()
```

Making bar charts of the number of genes which are regulated in each treatment condition for visualization of results

```
sum_lng <- sum_tbl %>%
  pivot_longer(cols = !c(1:5), names_to = c("treat", "stat"), names_sep = "_", values_to = "value") %>%
  pivot_wider(names_from = "stat", values_from = "value") %>%
  replace_na(list(pval = 1, adjp = 1)) %>%
  mutate(treat = factor(treat, c("idela", "pred", "pi", "dex", "di")))

sum_lng %>%
  group_by(treat) %>%
  summarise(Up = sum(adjp <= 0.01 & log2FC > 0), Down = sum(adjp <= 0.01 & log2FC < 0)) %>%
  pivot_longer(cols = c("Up", "Down"), names_to = "Regulation", values_to = "Number") %>%
  ggplot(aes(treat, Number, fill = Regulation)) +
  geom_col(width = 0.8, position=position_dodge(0.9)) +
  scale_fill_manual(values=c('blue','red')) +
  scale_x_discrete(breaks=c("idela", "pred", "pi", "dex", "di"), labels=c("Idela", "Pred", "Pred +\nIdela", "Dex", "Dex +\nI
dela")) +
  theme_bw() +
  ylab("Number of Genes Regulated") +
  theme(axis.title.x=element_blank(), axis.title.y = element_text(face = "bold"), legend.position = c(0.12, 0.85))
```

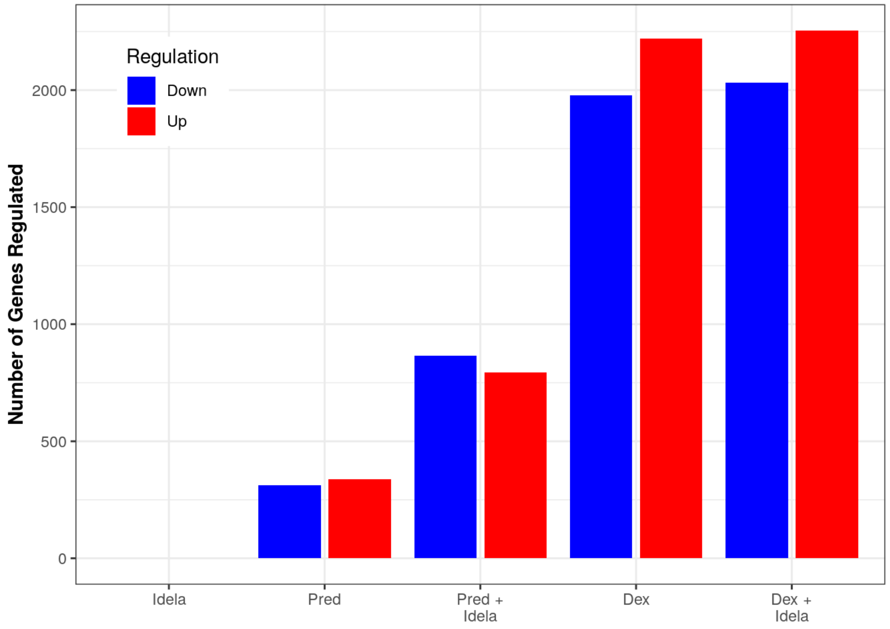

```
# ggsave("pt_samples_up_down_summary.pdf", width = 5, height = 4)
# ggsave("pt_samples_up_down_summary.png", width = 5, height = 4)
# ggsave("pt_samples_up_down_summary.svg", width = 5, height = 4)
```

Plotting Dex vs. Dex + Idela

```
dex_vs_di_all <- sum_tbl %>%
  dplyr::filter(dex_adjp <= 0.01 | di_adjp <= 0.01) %>%
  dplyr::filter(abs(dex_log2FC) < 10 & abs(di_log2FC) < 10) %>%
  ggplot(aes(x = dex_log2FC, y = di_log2FC)) +
  geom_point() +
  xlim(-6, 10) + ylim(-8, 10) +
  geom_abline(slope = 1, intercept = 0, color = "red") +
  geom_smooth(method = "lm", color = "black") +
  stat_cor(label.x.npc = "left", label.y.npc = "top") +
  stat_regline_equation(label.y = 8, aes(label = after_stat(eq.label))) +
  xlab("Dex Log2FC") +
  ylab("Dex+Idela Log2FC") +
  theme_bw()

dex_vs_di_all
```

```
## `geom_smooth()` using formula = 'y ~ x'
```

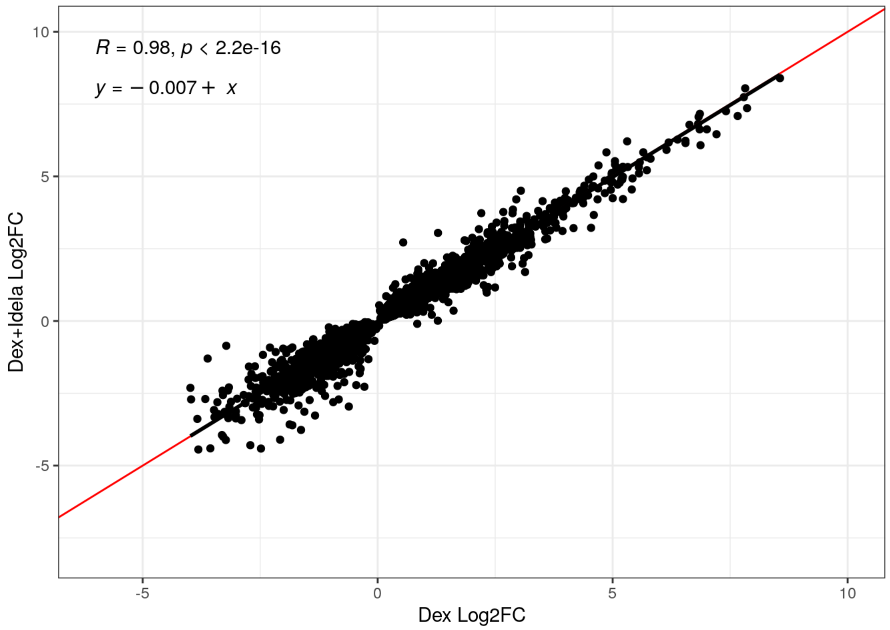

Plotting Pred vs Pred + Idela

```
pred_vs_pi_all <- sum_tbl %>%
  dplyr::filter(pred_adjp <= 0.01 | pi_adjp <= 0.01) %>%
  dplyr::filter(abs(pred_log2FC) < 10 & abs(pi_log2FC) < 10) %>%
  ggplot(aes(x = pred_log2FC, y = pi_log2FC)) +
  geom_point() +
  xlim(-4, 8) + ylim(-7, 10) +
  geom_abline(slope = 1, intercept = 0, color = "red") +
  geom_smooth(method = "lm", color = "black") +
  stat_cor(label.x.npc = "left", label.y.npc = "top") +
  stat_regline_equation(label.y = 8, aes(label = after_stat(eq.label))) +
  xlab("Pred Log2FC") +
  ylab("Pred+Idela Log2FC") +
  theme_bw()

pred_vs_pi_all
```

```
## `geom_smooth()` using formula = 'y ~ x'
```

```
## Warning: Removed 1 rows containing non-finite values (`stat_smooth()`).
```

```
## Warning: Removed 1 rows containing non-finite values (`stat_cor()`).
```

```
## Warning: Removed 1 rows containing non-finite values
## (`stat_regline_equation()`).
```

```
## Warning: Removed 1 rows containing missing values (`geom_point()`).
```

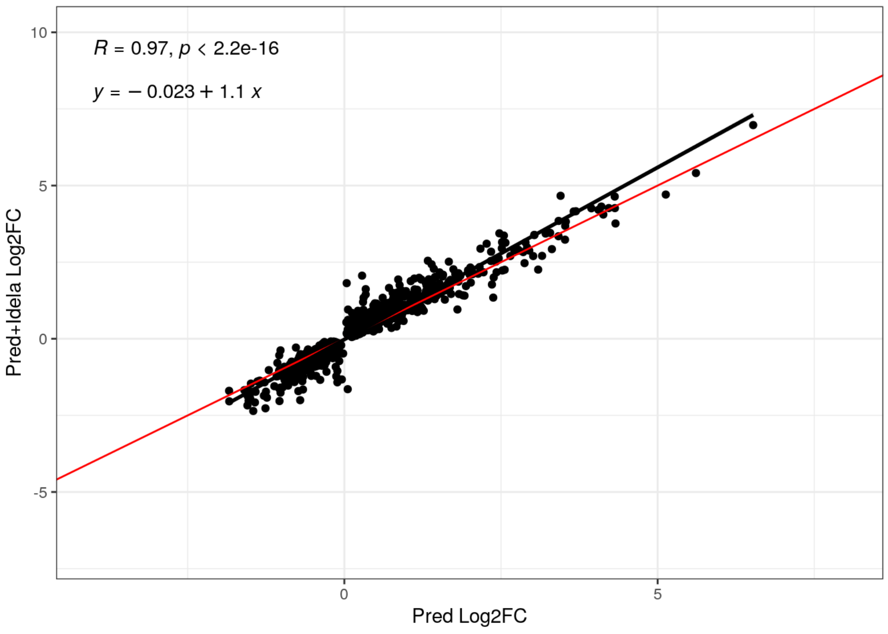

This is consistent with GC sensitive specimens (which these 4 all are) - idela enhances pred but not dex

#### I will look at effector genes again and try to create figures like figure 2E (effector gene regulation in NALM6 cells). I will look at all 4 of these GC sensitive specimens together (since they all contain 2 biological replicates) and also at additive vs. synergistic specimens to see if there are any effector genes which stick out as being enhanced by idela.

First, read in screen data from NALM6 cells:

```
full_rhos <- readxl::read_excel("./full_rhos_180815.xlsx")
sig_rhos <- dplyr::filter(full_rhos, Rho.P.value < 0.05)
full_gammas <- read_csv("./full_gammas_180815.csv")
```

```
## New names:
## Rows: 19132 Columns: 9
## ── Column specification
## ──────────────────────────────────────────────────────── Delimiter: "," chr
## (2): Symbol, GeneInfo dbl (7): ...1, Entrez_Gene_ID, Gamma...shRNAs,
## Gamma...shRNAs.with.sufficien...
## ℹ Use `spec()` to retrieve the full column specification for this data. ℹ
## Specify the column types or set `show_col_types = FALSE` to quiet this message.
## • `` -> `...1`
```

```
sig_gammas <- dplyr::filter(full_gammas, Gamma.P.value < 0.01)

cagek_rhos <- readxl::read_excel("./CAGEK_rhos_1508.xlsx")
c_sig_rhos <- dplyr::filter(cagek_rhos, `Rho P value` < 0.05)
cagek_gammas <- readxl::read_excel("./CAGEK_rhos_1508.xlsx", sheet = 2)
c_sig_gammas <-dplyr::filter(cagek_gammas, `Gamma P value` < 0.01)
```

##### Make tables of results filtered for significance and whether genes are upregulated or down regulated

```
sig_additive_dex <- dplyr::filter(sum_tbl_add, dex_adjp <= 0.01 | di_adjp <= 0.01)
sig_additive_pred <- dplyr::filter(sum_tbl_add, pred_adjp <= 0.01 | pi_adjp <= 0.01)

sig_synergistic_dex <- dplyr::filter(sum_tbl_syn, dex_syn_adjp <= 0.01 | di_syn_adjp <= 0.01)
sig_synergistic_pred <- dplyr::filter(sum_tbl_syn, pred_syn_adjp <= 0.01 | pi_syn_adjp <= 0.01)

sig_all_dex <- dplyr::filter(sum_tbl, dex_adjp <= 0.01 | di_adjp <= 0.01)
sig_all_pred <- dplyr::filter(sum_tbl, pred_adjp <= 0.01 | pi_adjp <= 0.01)

sig_add_up_dex <- dplyr::filter(sig_additive_dex, dex_log2FC > 0)
sig_add_down_dex <- dplyr::filter(sig_additive_dex, dex_log2FC < 0)

sig_add_up_pred <- dplyr::filter(sig_additive_pred, pred_log2FC > 0)
sig_add_down_pred <- dplyr::filter(sig_additive_pred, pred_log2FC < 0)

sig_syn_up_dex <- dplyr::filter(sig_synergistic_dex, dex_syn_log2FC > 0)
sig_syn_down_dex <- dplyr::filter(sig_synergistic_dex, dex_syn_log2FC < 0)

sig_syn_up_pred <- dplyr::filter(sig_synergistic_pred, pred_syn_log2FC > 0)
sig_syn_down_pred <- dplyr::filter(sig_synergistic_pred, pred_syn_log2FC < 0)

sig_all_up_dex <- dplyr::filter(sig_all_dex, dex_log2FC > 0)
sig_all_down_dex <- dplyr::filter(sig_all_dex, dex_log2FC < 0)

sig_all_up_pred <- dplyr::filter(sig_all_pred, pred_log2FC > 0)
sig_all_down_pred <- dplyr::filter(sig_all_pred, pred_log2FC < 0)

dim(sig_add_up_dex)
```

```
## [1] 3830   25
```

```
dim(sig_add_down_dex)
```

```
## [1] 3538   25
```

```
dim(sig_add_up_pred)
```

```
## [1] 1690   25
```

```
dim(sig_add_down_pred)
```

```
## [1] 1943   25
```

```
dim(sig_syn_up_dex)
```

```
## [1] 2426   25
```

```
dim(sig_syn_down_dex)
```

```
## [1] 2456   25
```

```
dim(sig_syn_up_pred)
```

```
## [1] 869  25
```

```
dim(sig_syn_down_pred)
```

```
## [1] 864  25
```

```
dim(sig_all_up_dex)
```

```
## [1] 2591   25
```

```
dim(sig_all_down_dex)
```

```
## [1] 2372   25
```

```
dim(sig_all_up_pred)
```

```
## [1] 834  25
```

```
dim(sig_all_down_pred)
```

```
## [1] 908  25
```

### The way to figure out which effector genes are most strongly regulated would be to take the significant rho genes, and overlap them with the signifcantly regulated genes. For those, we then want to look at those with the biggest fold change difference between dex and dex + idela (and also pred and pred+idela)

Will also want to do this process for additive and synergistic samples and all 4 samples together (GC sensitive samples)

```
# full screen with additive samples and dex
olap_rhos_add <- sig_rhos %>%
  inner_join(sig_additive_dex, by = c("Symbol" = "symbol")) %>%
  mutate(diff_dex_add = dex_log2FC - di_log2FC)
n_distinct(olap_rhos_add$Symbol)
```

```
## [1] 651
```

```
# full screen with additive samples and pred
olap_rhos_add_pred <- sig_rhos %>%
  inner_join(sig_additive_pred, by = c("Symbol" = "symbol")) %>%
  mutate(diff_pred_add = pred_log2FC - pi_log2FC)
n_distinct(olap_rhos_add_pred$Symbol)
```

```
## [1] 357
```

```
# cagek screen with additive samples and dex
c_olap_rhos_add <- c_sig_rhos %>%
  inner_join(sig_additive_dex, by = c("Symbol" = "symbol")) %>%
  mutate(diff_dex_add = dex_log2FC - di_log2FC)
n_distinct(c_olap_rhos_add$Symbol)
```

```
## [1] 286
```

```
# cagek screen with additive samples and pred
c_olap_rhos_add_pred <- c_sig_rhos %>%
  inner_join(sig_additive_pred, by = c("Symbol" = "symbol")) %>%
  mutate(diff_pred_add = pred_log2FC - pi_log2FC)
n_distinct(c_olap_rhos_add_pred$Symbol)
```

```
## [1] 156
```

##### Combine the results of both screens for the additive samples

```
doub_imp_genes_add_dex <- intersect(olap_rhos_add$Symbol, c_olap_rhos_add$Symbol)
doub_imp_genes_add_dex
```

```
##  [1] "ACADM"    "ADNP"     "AFF1"     "ARID1A"   "BCL2"     "BCL2L11"
##  [7] "BCOR"     "BIRC5"    "BMF"      "BOP1"     "BRD2"     "BRD4"
## [13] "C17orf49" "CARM1"    "CD79A"    "CELF1"    "CHAMP1"   "CTCF"
## [19] "DLGAP5"   "DOLPP1"   "EHMT2"    "EIF2B1"   "EIF3I"    "EIF3L"
## [25] "EP300"    "ETV6"     "GPS2"     "GSK3A"    "HIF1A"    "IRAK4"
## [31] "ITPKB"    "KAT6A"    "LARP1"    "MAML2"    "MAPK1"    "MBNL1"
## [37] "MED11"    "MED13"    "MEF2A"    "MMP14"    "MSI2"     "NCK1"
## [43] "NCOA2"    "NCOR2"    "NELFCD"   "NLE1"     "NOL6"     "NR3C1"
## [49] "PAX5"     "PDCD5"    "PHC3"     "PIK3CD"   "PLAGL2"   "POLG"
## [55] "POU2F1"   "PPP1R12A" "PPP5C"    "PRC1"     "PRDM1"    "PREX1"
## [61] "PRR12"    "PTBP1"    "RASSF4"   "RAVER1"   "RBMX2"    "RRP12"
## [67] "RUVBL1"   "SAFB"     "SAFB2"    "SETD1A"   "SPEN"     "SRRM1"
## [73] "SSRP1"    "SUPT16H"  "TADA3"    "THOC2"    "WIZ"      "YTHDC1"
## [79] "ZBED4"    "ZMIZ1"    "ZMYM4"    "ZMYND8"   "ZNF320"   "ZNF592"
## [85] "ZNF638"   "ZNF671"
```

```
doub_imp_genes_add_pred <- intersect(olap_rhos_add_pred$Symbol, c_olap_rhos_add_pred$Symbol)
doub_imp_genes_add_pred
```

```
##  [1] "ACADM"   "AFF1"    "ARID1A"  "BCL2"    "BMF"     "BOP1"    "CARM1"
##  [8] "CHAMP1"  "CREBBP"  "DLGAP5"  "DOLPP1"  "EHMT2"   "EIF3I"   "EP300"
## [15] "IRAK4"   "LARP1"   "MBNL1"   "NLE1"    "NOL6"    "NR3C1"   "PAX5"
## [22] "PDCD5"   "PIK3CD"  "PLAGL2"  "POU2F1"  "PPP5C"   "PRDM1"   "PREX1"
## [29] "PRR12"   "PTBP1"   "RRP12"   "RUVBL1"  "SAFB2"   "SPEN"    "SRRM1"
## [36] "SSRP1"   "SUPT16H" "YTHDC1"  "ZNF320"  "ZNF638"  "ZNF671"
```

##### And now for the synergistic samples

```
# full screen with synergistic samples and dex
olap_rhos_syn <- sig_rhos %>%
  inner_join(sig_synergistic_dex, by = c("Symbol" = "symbol")) %>%
  mutate(diff_dex_syn = dex_syn_log2FC - di_syn_log2FC)
n_distinct(olap_rhos_syn$Symbol)
```

```
## [1] 362
```

```
# full screen with synergistic samples and pred
olap_rhos_syn_pred <- sig_rhos %>%
  inner_join(sig_synergistic_pred, by = c("Symbol" = "symbol")) %>%
  mutate(diff_pred_syn = pred_syn_log2FC - pi_syn_log2FC)
n_distinct(olap_rhos_syn_pred$Symbol)
```

```
## [1] 135
```

```
# cagek screen with synergistic samples and dex
c_olap_rhos_syn <- c_sig_rhos %>%
  inner_join(sig_synergistic_dex, by = c("Symbol" = "symbol")) %>%
  mutate(diff_dex_syn = dex_syn_log2FC - di_syn_log2FC)
n_distinct(c_olap_rhos_syn$Symbol)
```

```
## [1] 188
```

```
# cagek screen with synergistic samples and pred
c_olap_rhos_syn_pred <- c_sig_rhos %>%
  inner_join(sig_synergistic_pred, by = c("Symbol" = "symbol")) %>%
  mutate(diff_pred_syn = pred_syn_log2FC - pi_syn_log2FC)
n_distinct(c_olap_rhos_syn_pred$Symbol)
```

```
## [1] 66
```

##### Combine the results of both screens for the synergistic samples

```
doub_imp_genes_syn_dex <- intersect(olap_rhos_syn$Symbol, c_olap_rhos_syn$Symbol)
doub_imp_genes_syn_dex
```

```
##  [1] "AFF1"     "ANKRD11"  "ARID1A"   "BBX"      "BCL2L11"  "BCOR"    
##  [7] "BMF"      "BRD2"     "BRD4"     "C17orf49" "CARS2"    "CD79A"   
## [13] "CDC42"    "CHAMP1"   "CNOT2"    "CPEB3"    "CREBBP"   "EBF1"    
## [19] "EHMT2"    "EIF4E2"   "EP300"    "ETV6"     "GPS2"     "GSK3A"   
## [25] "HIF1A"    "IRAK4"    "KAT6A"    "LEF1"     "MBNL1"    "MED23"   
## [31] "MEF2A"    "MMP14"    "MSI2"     "MTMR4"    "NCK1"     "NCOA1"   
## [37] "NCOR2"    "NUP214"   "PARD6B"   "PAX5"     "PHF6"     "PIK3CD"  
## [43] "POLG"     "POU2F1"   "PRDM1"    "PREX1"    "PRKAB1"   "PRR12"   
## [49] "RGS9"     "RRP12"    "RUVBL1"   "SAFB2"    "SESN3"    "SPEN"    
## [55] "SPI1"     "SRRM1"    "SSRP1"    "SYK"      "TAF3"     "ZMIZ1"   
## [61] "ZNF592"   "ZNF608"
```

```
doub_imp_genes_syn_pred <- intersect(olap_rhos_syn_pred$Symbol, c_olap_rhos_syn_pred$Symbol)
doub_imp_genes_syn_pred
```

```
##  [1] "AFF1"     "BCOR"     "C17orf49" "EP300"    "ETV6"     "IRAK4"   
##  [7] "LEF1"     "MBNL1"    "MTMR4"    "NCOA1"    "NUP214"   "PAX5"    
## [13] "POU2F1"   "PRR12"    "RUVBL1"   "SPEN"     "SPI1"     "SRRM1"   
## [19] "SSRP1"    "SYK"      "ZNF608"
```

Combine the results of both screens for the synergistic samples

```
doub_imp_genes_syn_dex <- intersect(olap_rhos_syn$Symbol, c_olap_rhos_syn$Symbol)
doub_imp_genes_syn_dex
```

```
##  [1] "AFF1"     "ANKRD11"  "ARID1A"   "BBX"      "BCL2L11"  "BCOR"    
##  [7] "BMF"      "BRD2"     "BRD4"     "C17orf49" "CARS2"    "CD79A"   
## [13] "CDC42"    "CHAMP1"   "CNOT2"    "CPEB3"    "CREBBP"   "EBF1"    
## [19] "EHMT2"    "EIF4E2"   "EP300"    "ETV6"     "GPS2"     "GSK3A"   
## [25] "HIF1A"    "IRAK4"    "KAT6A"    "LEF1"     "MBNL1"    "MED23"   
## [31] "MEF2A"    "MMP14"    "MSI2"     "MTMR4"    "NCK1"     "NCOA1"   
## [37] "NCOR2"    "NUP214"   "PARD6B"   "PAX5"     "PHF6"     "PIK3CD"  
## [43] "POLG"     "POU2F1"   "PRDM1"    "PREX1"    "PRKAB1"   "PRR12"   
## [49] "RGS9"     "RRP12"    "RUVBL1"   "SAFB2"    "SESN3"    "SPEN"    
## [55] "SPI1"     "SRRM1"    "SSRP1"    "SYK"      "TAF3"     "ZMIZ1"   
## [61] "ZNF592"   "ZNF608"
```

```
doub_imp_genes_syn_pred <- intersect(olap_rhos_syn_pred$Symbol, c_olap_rhos_syn_pred$Symbol)
doub_imp_genes_syn_pred
```

```
##  [1] "AFF1"     "BCOR"     "C17orf49" "EP300"    "ETV6"     "IRAK4"   
##  [7] "LEF1"     "MBNL1"    "MTMR4"    "NCOA1"    "NUP214"   "PAX5"    
## [13] "POU2F1"   "PRR12"    "RUVBL1"   "SPEN"     "SPI1"     "SRRM1"   
## [19] "SSRP1"    "SYK"      "ZNF608"
```

Comparing effector genes for additive and synergistic specimens with dexamethasone:

```
dex_effector_comp <- intersect(doub_imp_genes_add_dex, doub_imp_genes_syn_dex)

dex_effector_comp
```

```
##  [1] "AFF1"     "ARID1A"   "BCL2L11"  "BCOR"     "BMF"      "BRD2"    
##  [7] "BRD4"     "C17orf49" "CD79A"    "CHAMP1"   "EHMT2"    "EP300"   
## [13] "ETV6"     "GPS2"     "GSK3A"    "HIF1A"    "IRAK4"    "KAT6A"   
## [19] "MBNL1"    "MEF2A"    "MMP14"    "MSI2"     "NCK1"     "NCOR2"   
## [25] "PAX5"     "PIK3CD"   "POLG"     "POU2F1"   "PRDM1"    "PREX1"   
## [31] "PRR12"    "RRP12"    "RUVBL1"   "SAFB2"    "SPEN"     "SRRM1"   
## [37] "SSRP1"    "ZMIZ1"    "ZNF592"
```

```
n_distinct(dex_effector_comp)
```

```
## [1] 39
```

```
n_distinct(doub_imp_genes_add_dex)
```

```
## [1] 86
```

```
n_distinct(doub_imp_genes_syn_dex)
```

```
## [1] 62
```

##### Comparing effector genes for additive and synergistic specimens with prednisolone:

```
pred_effector_comp <- intersect(doub_imp_genes_add_pred, doub_imp_genes_syn_pred)

pred_effector_comp
```

```
##  [1] "AFF1"   "EP300"  "IRAK4"  "MBNL1"  "PAX5"   "POU2F1" "PRR12"  "RUVBL1"
##  [9] "SPEN"   "SRRM1"  "SSRP1"
```

```
n_distinct(pred_effector_comp)
```

```
## [1] 11
```

```
n_distinct(doub_imp_genes_add_pred)
```

```
## [1] 41
```

```
n_distinct(doub_imp_genes_syn_pred)
```

```
## [1] 21
```

Make a table of effector gene lists in each comparison

```
combined_effectors_dex <- qpcR:::cbind.na(doub_imp_genes_add_dex, doub_imp_genes_syn_dex, dex_effector_comp)

# write.csv(combined_effectors_dex, file="effector_gene_dex_comparison_20221128.csv")
```

```
combined_effectors_pred <- qpcR:::cbind.na(doub_imp_genes_add_pred, doub_imp_genes_syn_pred, pred_effector_comp)

# write.csv(combined_effectors_pred, file="effector_gene_pred_comparison_20221128.csv")
```

##### Now combining results of screen with all samples together (GC sensitive samples)

```
# full screen with GC sensitive samples and dex
olap_rhos_all <- sig_rhos %>%
  inner_join(sig_all_dex, by = c("Symbol" = "symbol")) %>%
  mutate(diff_dex_all = dex_log2FC - di_log2FC)
n_distinct(olap_rhos_all$Symbol)
```

```
## [1] 392
```

```
# full screen with GC sensitive samples and pred
olap_rhos_all_pred <- sig_rhos %>%
  inner_join(sig_all_pred, by = c("Symbol" = "symbol")) %>%
  mutate(diff_pred_all = pred_log2FC - pi_log2FC)
n_distinct(olap_rhos_all_pred$Symbol)
```

```
## [1] 162
```

```
# cagek screen with GC sensitive samples and dex
c_olap_rhos_all <- c_sig_rhos %>%
  inner_join(sig_all_dex, by = c("Symbol" = "symbol")) %>%
  mutate(diff_dex_all = dex_log2FC - di_log2FC)
n_distinct(c_olap_rhos_all$Symbol)
```

```
## [1] 198
```

```
# cagek screen with GC sensitive samples and pred
c_olap_rhos_all_pred <- c_sig_rhos %>%
  inner_join(sig_all_pred, by = c("Symbol" = "symbol")) %>%
  mutate(diff_pred_all = pred_log2FC - pi_log2FC)
n_distinct(c_olap_rhos_all_pred$Symbol)
```

```
## [1] 91
```

Combine the results of both screens for the four GC sensitive samples

```
doub_imp_genes_all_dex <- intersect(olap_rhos_all$Symbol, c_olap_rhos_all$Symbol)
doub_imp_genes_all_dex
```

```
##  [1] "AFF1"     "ANKRD11"  "ARID1A"   "BCL2"     "BCL2L11"  "BCOR"
##  [7] "BMF"      "BRD2"     "BRD4"     "C17orf49" "CD79A"    "CHAMP1"
## [13] "CREBBP"   "CTCF"     "DDX46"    "EBF1"     "EHMT2"    "EIF2B1"
## [19] "EP300"    "ETV6"     "FOXJ3"    "GPS2"     "GSK3A"    "HIF1A"
## [25] "IRAK4"    "KAT6A"    "LARP1"    "LEF1"     "MAML2"    "MAPK1"
## [31] "MBNL1"    "MED23"    "MSI2"     "MTMR4"    "NCK1"     "NCOA1"
## [37] "NCOR2"    "NLE1"     "NOL6"     "PARD6B"   "PHC3"     "PIK3CD"
## [43] "POLG"     "POU2F1"   "PPP5C"    "PRDM1"    "PREX1"    "PRR12"
## [49] "RRP12"    "RUVBL1"   "SAFB"     "SAFB2"    "SRRM1"    "SSRP1"
## [55] "SUPT16H"  "TADA3"    "YTHDC1"   "ZBED4"    "ZMIZ1"    "ZNF592"
## [61] "ZNF608"   "ZNF671"
```

```
doub_imp_genes_all_pred <- intersect(olap_rhos_all_pred$Symbol, c_olap_rhos_all_pred$Symbol)
doub_imp_genes_all_pred
```

```
##  [1] "AFF1"    "BCL2"    "CHAMP1"  "CREBBP"  "CTCF"    "EP300"   "IRAK4"
##  [8] "LEF1"    "MBNL1"   "NOL6"    "PIK3CD"  "POU2F1"  "PPP5C"   "PRDM1"
## [15] "PREX1"   "RRP12"   "RUVBL1"  "SRRM1"   "SSRP1"   "SUPT16H" "YTHDC1"
## [22] "ZNF638"  "ZNF671"
```

Make bar charts of the effector genes:

First, additive samples:

```
# sum_lng_add <- mutate(treat = factor(treat, levels = c("idela", "pred", "pi", "dex", "di")))

additive_effectors_dex <- sum_lng_add %>%
  dplyr::filter(symbol %in% doub_imp_genes_add_dex) %>%
  ggplot(aes(treat, log2FC, fill = treat)) +
  labs(x = "", y = "log2FoldChange") +
  geom_col(position = "dodge") +
  scale_fill_viridis(discrete = T, option = "E") +
  facet_wrap(~symbol, scales = "free") +theme_bw() +
  theme(legend.position="none", axis.text.x = element_text(angle = 45, hjust = 1)) +
  geom_errorbar(aes(ymin=log2FC-lfcse, ymax=log2FC+lfcse), position = position_dodge(width = 0.9), width=0.5, colour="blac
k", size = 0.5)
```

```
## Warning: Using `size` aesthetic for lines was deprecated in ggplot2 3.4.0.
## ℹ Please use `linewidth` instead.
```

```
additive_effectors_dex
```

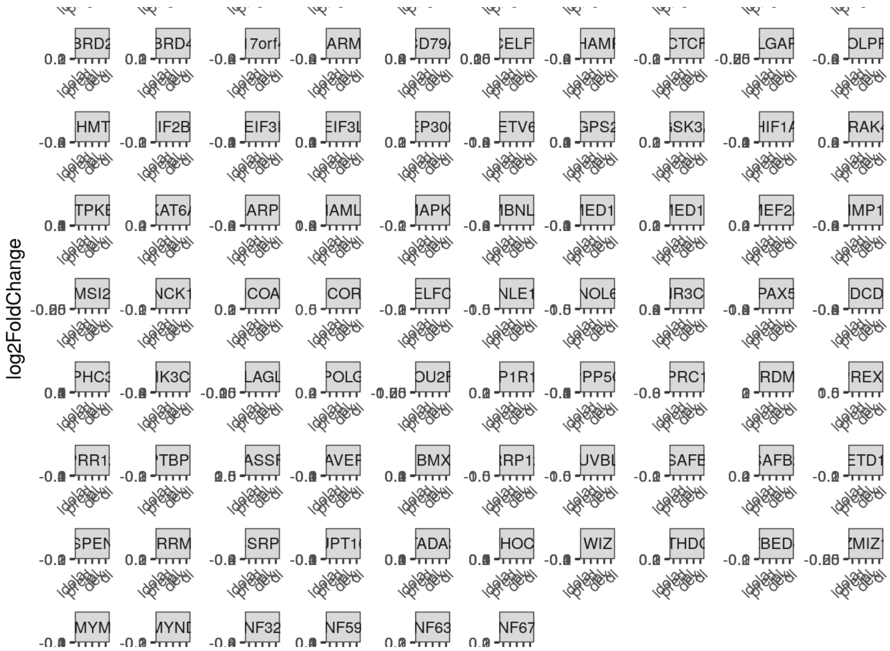

```
# ggsave("additive_effectors_dex.png", plot = additive_effectors_dex, width = 10, height = 6, units = "in")
```

```
additive_effectors_pred <- sum_lng_add %>%
  dplyr::filter(symbol %in% doub_imp_genes_add_pred) %>%
  ggplot(aes(treat, log2FC, fill = treat)) +
  labs(x = "", y = "log2FoldChange") +
  geom_col(position = "dodge") +
  scale_fill_viridis(discrete = T, option = "E") +
  facet_wrap(~symbol, scales = "free") +theme_bw() +
  theme(legend.position="none", axis.text.x = element_text(angle = 45, hjust = 1)) +
  geom_errorbar(aes(ymin=log2FC-lfcse, ymax=log2FC+lfcse), position = position_dodge(width = 0.9), width=0.5, colour="blac
k", size = 0.5)

additive_effectors_pred
```

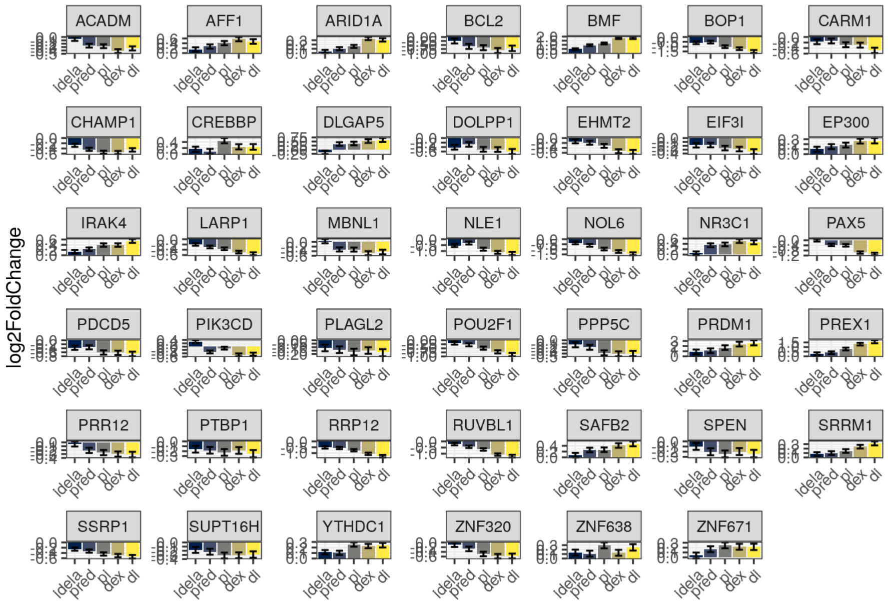

```
# ggsave("additive_effectors_pred.png", plot = additive_effectors_pred, width = 2, height = 2, units = "in")
```

Now synergistic samples:

```
# sum_lng_syn <- mutate(treat = factor(treat, levels = c("idela", "pred", "pi", "dex", "di")))

synergistic_effectors_dex <- sum_lng_syn %>%
  dplyr::filter(symbol %in% doub_imp_genes_syn_dex) %>%
  ggplot(aes(treat, log2FC, fill = treat)) +
  labs(x = "", y = "log2FoldChange") +
  geom_col(position = "dodge") +
  scale_fill_viridis(discrete = T, option = "E") +
  facet_wrap(~symbol, scales = "free") +theme_bw() +
  theme(legend.position="none", axis.text.x = element_text(angle = 45, hjust = 1)) +
  geom_errorbar(aes(ymin=log2FC-lfcse, ymax=log2FC+lfcse), position = position_dodge(width = 0.9), width=0.5, colour="blac
k", size = 0.5)

synergistic_effectors_dex
```

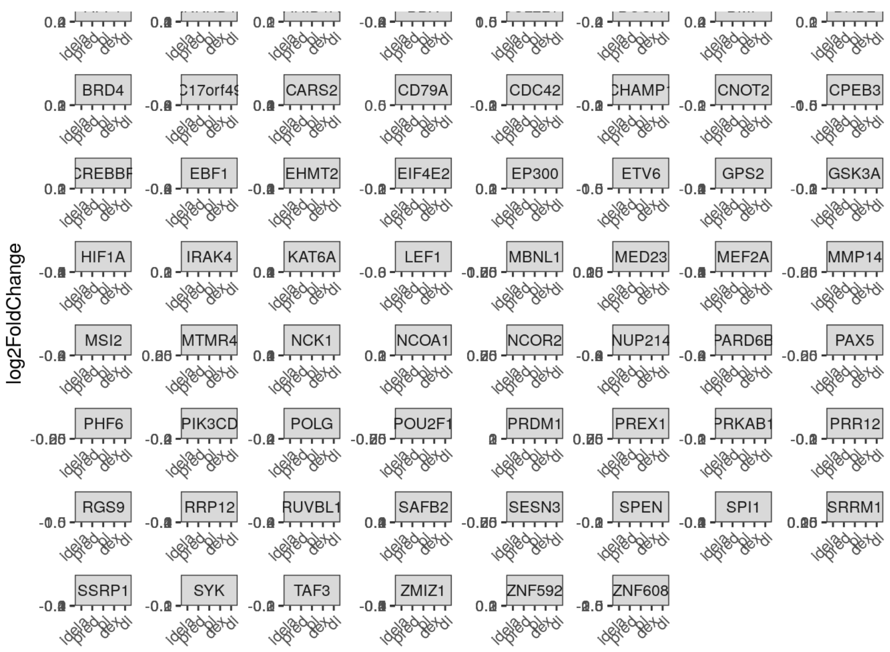

```
# ggsave("synergistic_effectors_dex.png", plot = synergistic_effectors_dex, width = 10, height = 6, units = "in")
```

```
synergistic_effectors_pred <- sum_lng_syn %>%
  dplyr::filter(symbol %in% doub_imp_genes_syn_pred) %>%
  ggplot(aes(treat, log2FC, fill = treat)) +
  labs(x = "", y = "log2FoldChange") +
  geom_col(position = "dodge") +
  scale_fill_viridis(discrete = T, option = "E") +
  facet_wrap(~symbol, scales = "free") +theme_bw() +
  theme(legend.position="none", axis.text.x = element_text(angle = 45, hjust = 1)) +
  geom_errorbar(aes(ymin=log2FC-lfcse, ymax=log2FC+lfcse), position = position_dodge(width = 0.9), width=0.5, colour="blac
k", size = 0.5)

synergistic_effectors_pred
```

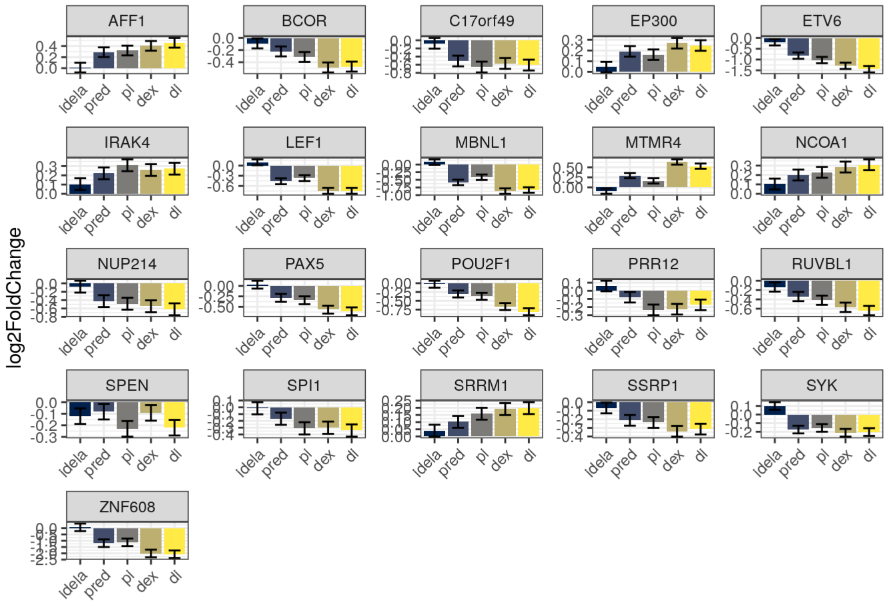

```
# ggsave("synergistic_effectors_pred.png", plot = synergistic_effectors_pred, width = 10, height = 6, units = "in")
```

Last, GC sensitive samples:

```
# sum_lng <- mutate(treat = factor(treat, levels = c("idela", "pred", "pi", "dex", "di")))

GCsens_effectors_dex <- sum_lng %>%
  dplyr::filter(symbol %in% doub_imp_genes_all_dex) %>%
  ggplot(aes(treat, log2FC, fill = treat)) +
  labs(x = "", y = "log2FoldChange") +
  geom_col(position = "dodge") +
  scale_fill_viridis(discrete = T, option = "E") +
  facet_wrap(~symbol, scales = "free") +theme_bw() +
  theme(legend.position="none", axis.text.x = element_text(angle = 45, hjust = 1)) +
  geom_errorbar(aes(ymin=log2FC-lfcse, ymax=log2FC+lfcse), position = position_dodge(width = 0.9), width=0.5, colour="blac
k", size = 0.5)

GCsens_effectors_dex
```

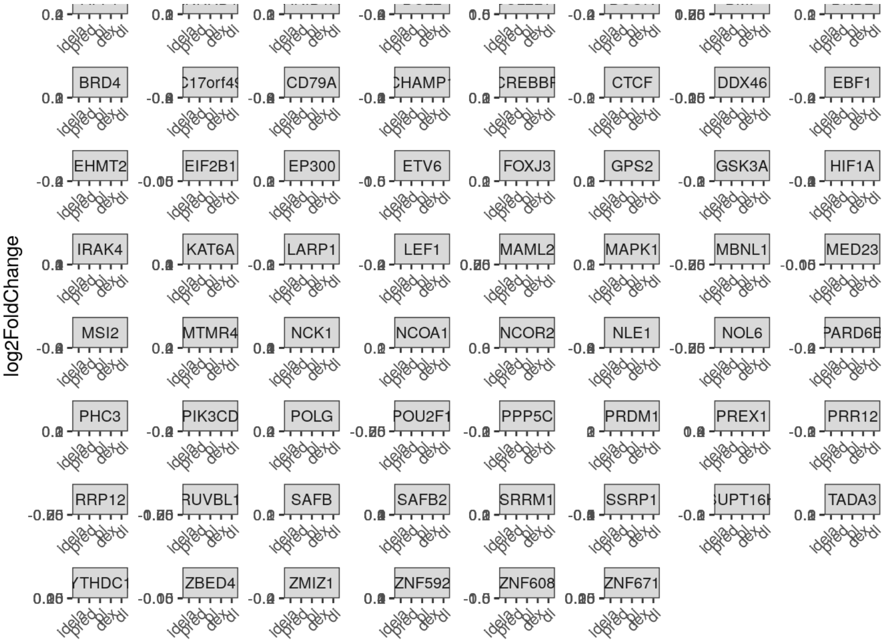

```
# ggsave("GCsens_effectors_dex.png", plot = GCsens_effectors_dex, width = 10, height = 6, units = "in")
```

```
GCsens_effectors_pred <- sum_lng %>%
  dplyr::filter(symbol %in% doub_imp_genes_all_pred) %>%
  ggplot(aes(treat, log2FC, fill = treat)) +
  labs(x = "", y = "log2FoldChange") +
  geom_col(position = "dodge") +
  scale_fill_viridis(discrete = T, option = "E") +
  facet_wrap(~symbol, scales = "free") +theme_bw() +
  theme(legend.position="none", axis.text.x = element_text(angle = 45, hjust = 1)) +
  geom_errorbar(aes(ymin=log2FC-lfcse, ymax=log2FC+lfcse), position = position_dodge(width = 0.9), width=0.5, colour="blac
k", size = 0.5)

GCsens_effectors_pred
```

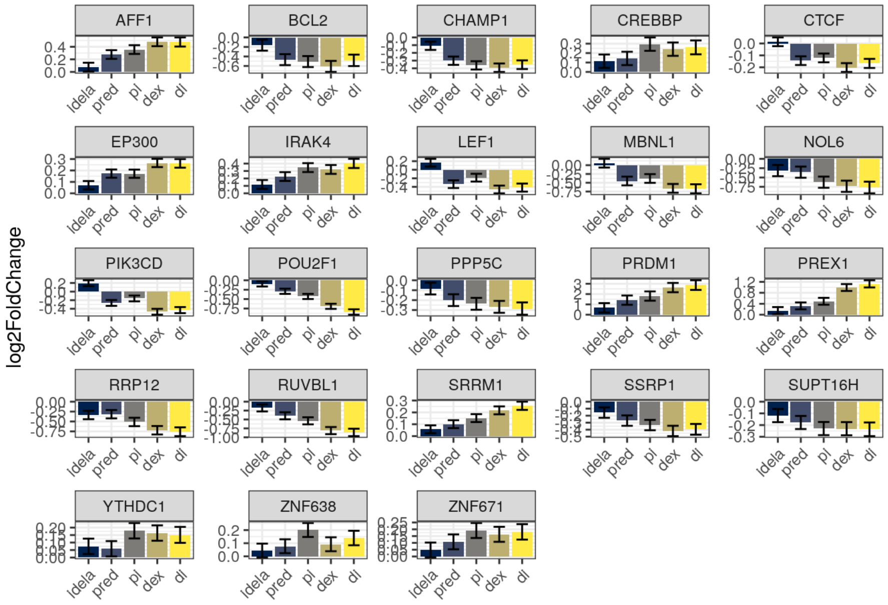

```
# ggsave("GCsens_effectors_pred.png", plot = GCsens_effectors_pred, width = 10, height = 6, units = "in")
```

Making a similar plot of effector genes that was done for NALM6 (figure 2E)

additive samples - dex

```
additive_dex_effector_plot <- sum_lng_add %>%
  dplyr::filter(symbol %in% doub_imp_genes_add_dex) %>%
  ggplot(aes(x= log2FC, y= reorder(symbol, log2FC))) +
  geom_line() +
  geom_point(aes(color=treat), size=2) +
  scale_color_viridis(discrete = T, option = "E") +
  theme_bw() + ylab("Effector Gene")+
  theme(legend.position="top")

additive_dex_effector_plot
```

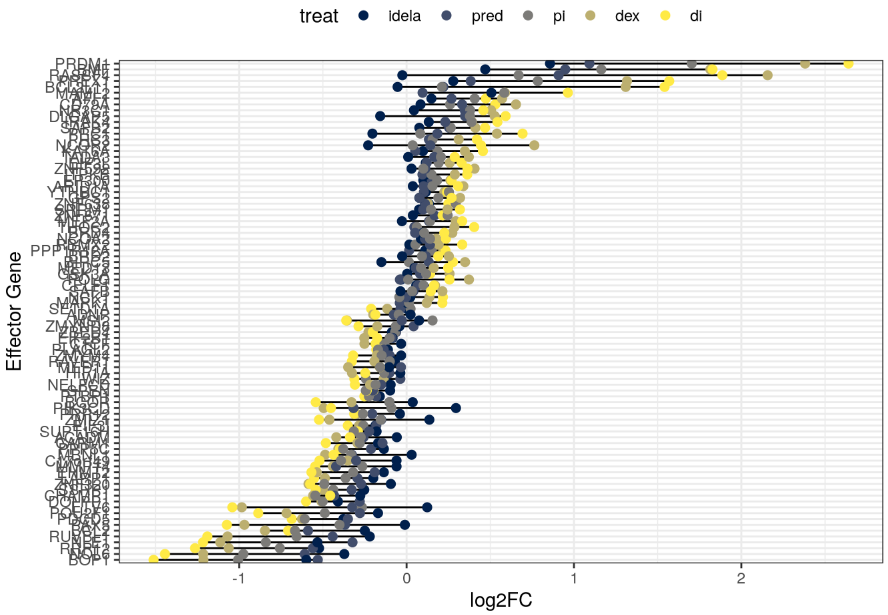

additive samples - pred

```
additive_pred_effector_plot <- sum_lng_add %>%
  dplyr::filter(symbol %in% doub_imp_genes_add_pred) %>%
  ggplot(aes(x= log2FC, y= reorder(symbol, log2FC))) +
  geom_line() +
  geom_point(aes(color=treat), size=2) +
  scale_color_viridis(discrete = T, option = "E") +
  theme_bw() + ylab("Effector Gene")+
  theme(legend.position="top")

additive_pred_effector_plot
```

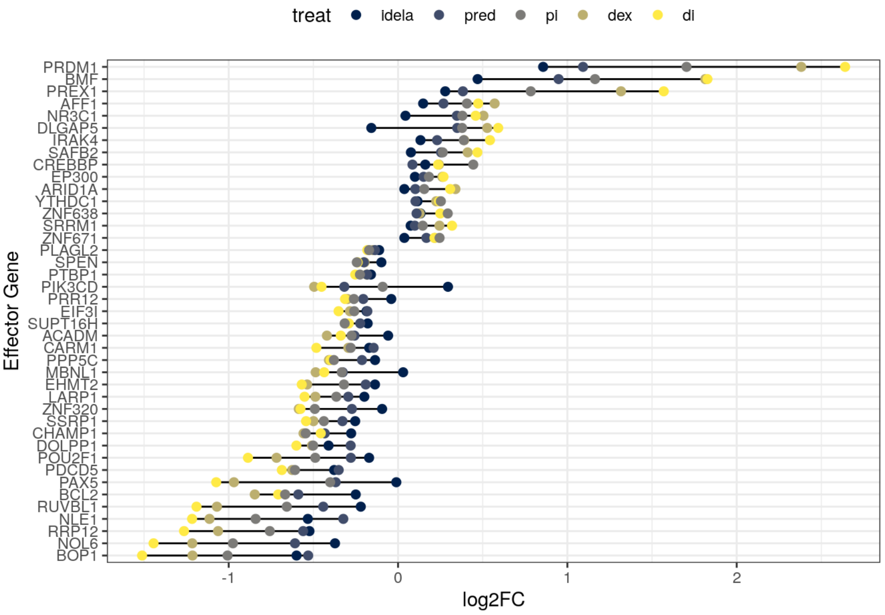

#### synergistic samples - dex

```
synergistic_dex_effector_plot <- sum_lng_syn %>%
  dplyr::filter(symbol %in% doub_imp_genes_syn_dex) %>%
  ggplot(aes(x= log2FC, y= reorder(symbol, log2FC))) +
  geom_line() +
  geom_point(aes(color=treat), size=2) +
  scale_color_viridis(discrete = T, option = "E") +
  theme_bw() + ylab("Effector Gene")+
  theme(legend.position="top")

synergistic_dex_effector_plot
```

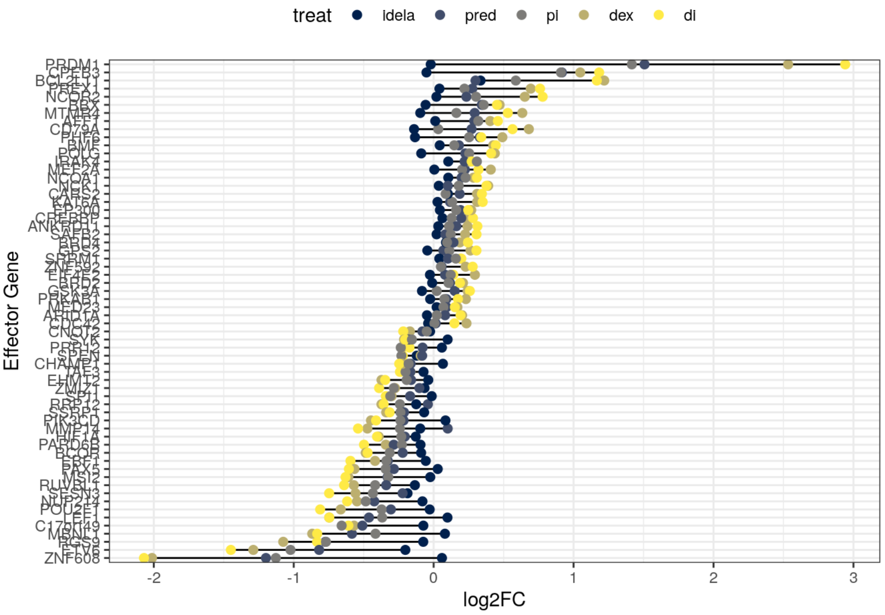

#### synergistic samples - pred

```
synergistic_pred_effector_plot <- sum_lng_syn %>%
  dplyr::filter(symbol %in% doub_imp_genes_syn_pred) %>%
  ggplot(aes(x= log2FC, y= reorder(symbol, log2FC))) +
  geom_line() +
  geom_point(aes(color=treat), size=2) +
  scale_color_viridis(discrete = T, option = "E") +
  theme_bw() + ylab("Effector Gene")+
  theme(legend.position="top")

synergistic_pred_effector_plot
```

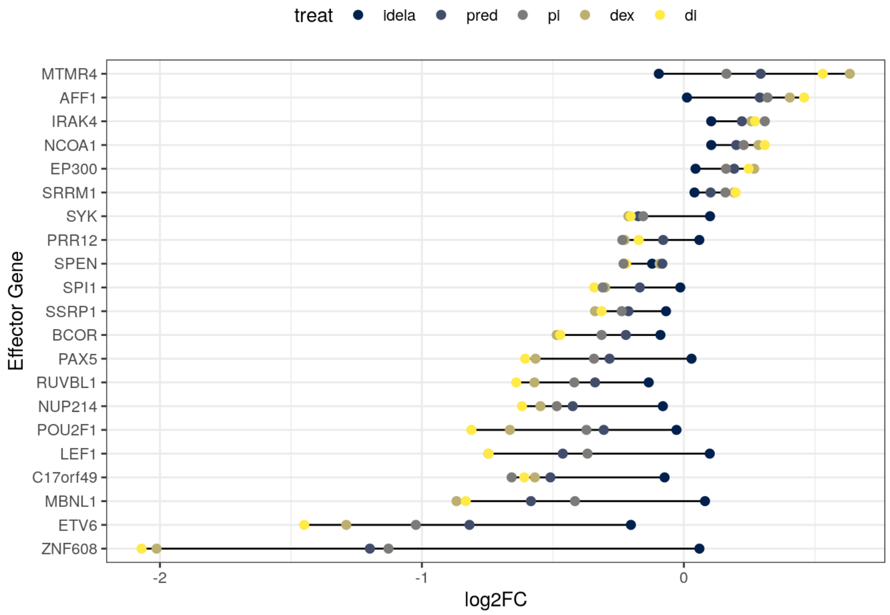

#### GC sensitive samples - dex

```
GCsens_dex_effector_plot <- sum_lng %>%
  dplyr::filter(symbol %in% doub_imp_genes_all_dex) %>%
  ggplot(aes(x= log2FC, y= reorder(symbol, log2FC))) +
  geom_line() +
  geom_point(aes(color=treat), size=2) +
  scale_color_viridis(discrete = T, option = "E") +
  theme_bw() + ylab("Effector Gene")+
  theme(legend.position="top")

GCsens_dex_effector_plot
```

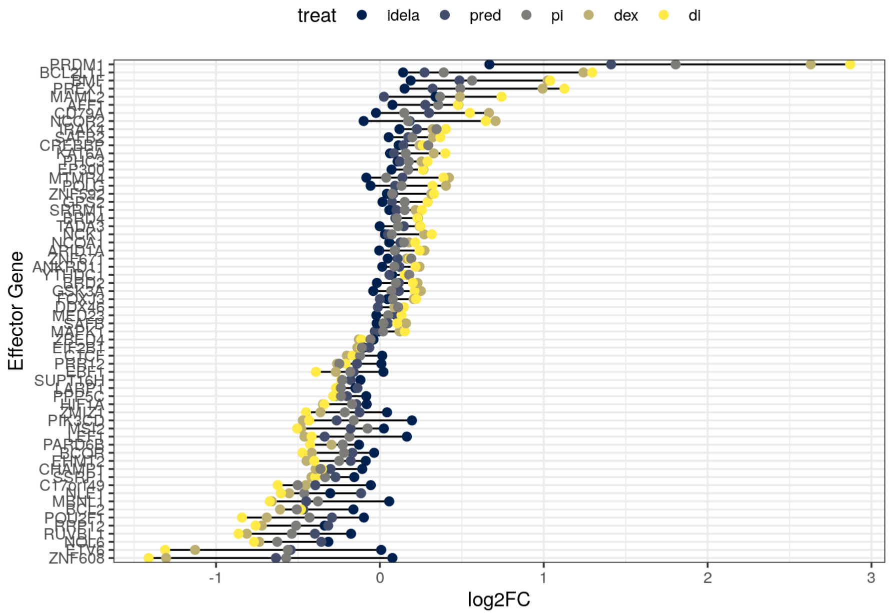

#### GC sensitive samples - pred

```
GCsens_pred_effector_plot <- sum_lng %>%
  dplyr::filter(symbol %in% doub_imp_genes_all_pred) %>%
  ggplot(aes(x= log2FC, y= reorder(symbol, log2FC))) +
  geom_line() +
  geom_point(aes(color=treat), size=2) +
  scale_color_viridis(discrete = T, option = "E") +
  theme_bw() + ylab("Effector Gene")+
  theme(legend.position="top")

GCsens_pred_effector_plot
```

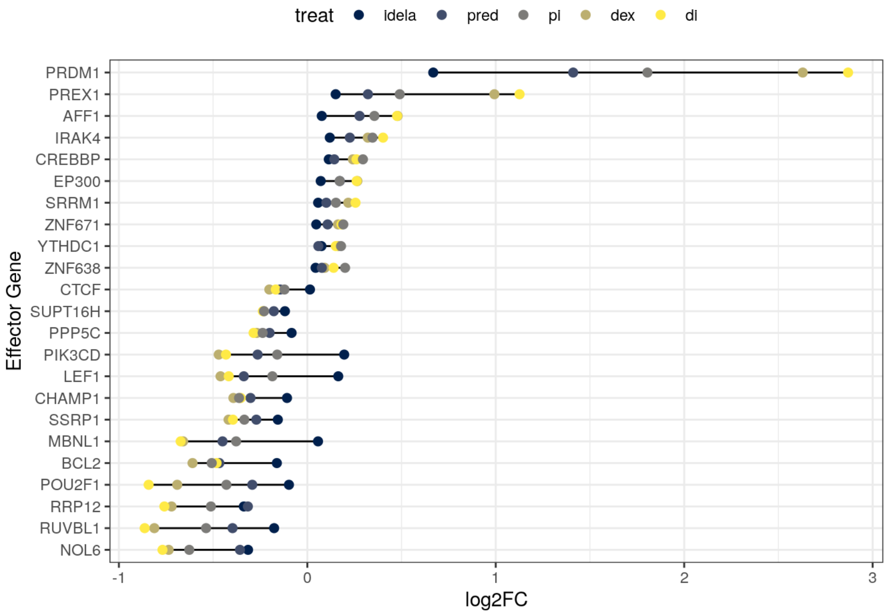

Save the graphs:

```
# ggsave("additive_effectors_dex_plot_20221123.pdf", additive_dex_effector_plot, width = 6, height = 10, units = "in")

# ggsave("synergistic_effectors_dex_plot_20221123.pdf", synergistic_dex_effector_plot, width = 6, height = 10, units = "in")

# ggsave("GCsens_effectors_dex_plot_20221123.pdf", GCsens_dex_effector_plot, width = 6, height = 10, units = "in")

# ggsave("additive_effectors_pred_plot_20221123.pdf", additive_pred_effector_plot, width = 6, height = 8, units = "in")

# ggsave("synergistic_effectors_pred_plot_20221123.pdf", synergistic_pred_effector_plot, width = 6, height = 8, units = "i
n")

# ggsave("GCsens_effectors_pred_plot_20221123.pdf", GCsens_pred_effector_plot, width = 6, height = 8, units = "in")
```

Session info:

```
sessionInfo()
```

```
## R version 4.1.1 (2021-08-10)
## Platform: x86_64-pc-linux-gnu (64-bit)
## Running under: Ubuntu 18.04.6 LTS
##
## Matrix products: default
## BLAS/LAPACK: /opt/OpenBLAS/lib/libopenblasp-r0.3.3.so
##
## locale:
##  [1] LC_CTYPE=en_US.UTF-8       LC_NUMERIC=C
##  [3] LC_TIME=en_US.UTF-8        LC_COLLATE=en_US.UTF-8
##  [5] LC_MONETARY=en_US.UTF-8    LC_MESSAGES=en_US.UTF-8
##  [7] LC_PAPER=en_US.UTF-8       LC_NAME=C
##  [9] LC_ADDRESS=C               LC_TELEPHONE=C
## [11] LC_MEASUREMENT=en_US.UTF-8 LC_IDENTIFICATION=C
##
## attached base packages:
## [1] parallel  stats4    stats     graphics  grDevices utils     datasets
## [8] methods   base
##
## other attached packages:
##  [1] qpcR_1.4-1                  Matrix_1.3-4
##  [3] robustbase_0.95-0           rgl_0.110.2
##  [5] minpack.lm_1.2-2            MASS_7.3-54
##  [7] RUVSeq_1.26.0               edgeR_3.34.1
##  [9] limma_3.48.3                EDASeq_2.26.1
## [11] ShortRead_1.50.0            GenomicAlignments_1.28.0
## [13] Rsamtools_2.8.0             Biostrings_2.60.2
## [15] XVector_0.32.0              BiocParallel_1.26.2
## [17] ggpubr_0.4.0                qvalue_2.24.0
## [19] viridis_0.6.2               viridisLite_0.4.1
## [21] forcats_0.5.2               stringr_1.5.0
## [23] dplyr_1.0.10                purrr_1.0.1
## [25] readr_2.1.3                 tidyr_1.2.1
## [27] tibble_3.1.8                ggplot2_3.4.0
## [29] tidyverse_1.3.2             ReportingTools_2.32.1
## [31] knitr_1.41                  org.Hs.eg.db_3.13.0
## [33] genefilter_1.74.1           apeglm_1.14.0
## [35] PoiClaClu_1.0.2.1           RColorBrewer_1.1-3
## [37] pheatmap_1.0.12             vsn_3.60.0
## [39] ensembldb_2.16.4            AnnotationFilter_1.16.0
## [41] GenomicFeatures_1.44.2      AnnotationDbi_1.54.1
## [43] rhdf5_2.36.0                DESeq2_1.32.0
## [45] SummarizedExperiment_1.22.0 Biobase_2.52.0
## [47] MatrixGenerics_1.4.3        matrixStats_0.62.0
## [49] GenomicRanges_1.44.0        GenomeInfoDb_1.28.4
## [51] IRanges_2.26.0              S4Vectors_0.30.2
## [53] BiocGenerics_0.38.0         tximeta_1.10.0
##
## loaded via a namespace (and not attached):
##   [1] rappdirs_0.3.3                rtracklayer_1.52.1
##   [3] AnnotationForge_1.34.1        GGally_2.1.2
##   [5] R.methodsS3_1.8.1             coda_0.19-4
##   [7] bit64_4.0.5                   aroma.light_3.22.0
##   [9] DelayedArray_0.18.0           R.utils_2.11.0
##  [11] PFAM.db_3.13.0                data.table_1.14.6
##  [13] rpart_4.1-15                  hwriter_1.3.2.1
##  [15] KEGGREST_1.32.0               RCurl_1.98-1.6
##  [17] generics_0.1.3                preprocessCore_1.54.0
##  [19] RSQLite_2.2.20                bit_4.0.5
##  [21] tzdb_0.3.0                    xml2_1.3.3
##  [23] lubridate_1.9.0               httpuv_1.6.8
##  [25] assertthat_0.2.1              gargle_1.2.1
##  [27] xfun_0.36                     tximport_1.20.0
##  [29] hms_1.1.2                     jquerylib_0.1.4
##  [31] evaluate_0.19                 promises_1.2.0.1
##  [33] DEoptimR_1.0-11               fansi_1.0.3
##  [35] restfulr_0.0.14               progress_1.2.2
##  [37] dbplyr_2.2.1                  readxl_1.4.1
##  [39] Rgraphviz_2.36.0              DBI_1.1.3
##  [41] geneplotter_1.70.0            htmlwidgets_1.6.1
##  [43] reshape_0.8.9                 googledrive_2.0.0
##  [45] ellipsis_0.3.2                backports_1.4.1
##  [47] annotate_1.70.0               biomaRt_2.48.3
##  [49] vctrs_0.5.1                   abind_1.4-5
##  [51] cachem_1.0.6                  withr_2.5.0
##  [53] BSgenome_1.60.0               vroom_1.6.0
##  [55] bdsmatrix_1.3-6               checkmate_2.1.0
##  [57] prettyunits_1.1.1             cluster_2.1.2
```

```
##  [59] lazyeval_0.2.2                crayon_1.5.2
##  [61] labeling_0.4.2                pkgconfig_2.0.3
##  [63] nlme_3.1-152                  ProtGenerics_1.24.0
##  [65] nnet_7.3-16                   rlang_1.0.6
##  [67] lifecycle_1.0.3               filelock_1.0.2
##  [69] affyio_1.62.0                 BiocFileCache_2.0.0
##  [71] GOstats_2.58.0                modelr_0.1.10
##  [73] AnnotationHub_3.0.2           dichromat_2.0-0.1
##  [75] cellranger_1.1.0              graph_1.70.0
##  [77] carData_3.0-5                 Rhdf5lib_1.14.2
##  [79] reprex_2.0.2                  base64enc_0.1-3
##  [81] googlesheets4_1.0.1           png_0.1-7
##  [83] rjson_0.2.21                  bitops_1.0-7
##  [85] R.oo_1.24.0                   rhdf5filters_1.4.0
##  [87] blob_1.2.3                    jpeg_0.1-9
##  [89] rstatix_0.7.0                 ggsignif_0.6.3
##  [91] scales_1.2.1                  memoise_2.0.1
##  [93] GSEABase_1.54.0               magrittr_2.0.3
##  [95] plyr_1.8.8                    zlibbioc_1.38.0
##  [97] compiler_4.1.1                BiocIO_1.2.0
##  [99] bbmle_1.0.25                  cli_3.6.0
## [101] affy_1.70.0                   Category_2.58.0
## [103] htmlTable_2.4.0               Formula_1.2-4
## [105] mgcv_1.8-36                   tidyselect_1.2.0
## [107] stringi_1.7.12                highr_0.10
## [109] emdbook_1.3.12                yaml_2.3.6
## [111] locfit_1.5-9.5                latticeExtra_0.6-29
## [113] grid_4.1.1                    sass_0.4.4
## [115] VariantAnnotation_1.38.0      polynom_1.4-1
## [117] tools_4.1.1                   timechange_0.2.0
## [119] rstudioapi_0.14               foreign_0.8-81
## [121] gridExtra_2.3                 farver_2.1.1
## [123] digest_0.6.31                 BiocManager_1.30.18
## [125] shiny_1.7.4                   Rcpp_1.0.9
## [127] car_3.0-13                    broom_1.0.2
## [129] BiocVersion_3.13.1            later_1.3.0
## [131] OrganismDbi_1.34.0            httr_1.4.4
## [133] ggbio_1.40.0                  biovizBase_1.40.0
## [135] colorspace_2.0-3              rvest_1.0.3
## [137] XML_3.99-0.9                  fs_1.5.2
## [139] splines_4.1.1                 RBGL_1.68.0
## [141] xtable_1.8-4                  jsonlite_1.8.4
## [143] R6_2.5.1                      Hmisc_4.7-0
## [145] pillar_1.8.1                  htmltools_0.5.4
## [147] mime_0.12                     glue_1.6.2
## [149] fastmap_1.1.0                 interactiveDisplayBase_1.30.0
## [151] mvtnorm_1.1-3                 utf8_1.2.2
## [153] lattice_0.20-44               bslib_0.4.2
## [155] numDeriv_2016.8-1.1           curl_5.0.0
## [157] GO.db_3.13.0                  survival_3.2-11
## [159] rmarkdown_2.19                munsell_0.5.0
## [161] GenomeInfoDbData_1.2.6        haven_2.5.1
## [163] reshape2_1.4.4                gtable_0.3.1
```
